# Take out the rubbish – Removing NUMTs and pseudogenes from the *Bemisia tabaci* cryptic species mtCOI database

**DOI:** 10.1101/724765

**Authors:** Daniele Kunz, Wee Tek Tay, Samia Elfekih, Karl Heinrichs Julius Gordon, Paul John De Barro

**Author notes:** These authors contributed equally (Daniele Kunz, Wee Tek Tay).

## Abstract

Identification of *Bemisia tabaci* cryptic whitefly species complex currently relies on molecular characterisation of the mitochondrial DNA cytochrome oxidase subunit I (mtCOI) partial gene, however, nuclear mitochondrial sequences (NUMTs), PCR-derived pseudogenes and/or poor sequence editing have hindered this effort. To-date, *ca.* 5,175 partial (≥ 300bp) mtCOI sequences for species identification purposes have been reported. We reviewed *ca.* 10% of sequences representing the standard *B. tabaci* species complex mtCOI dataset. We found that 333 sequences (64.9%) were NUMTs, pseudogenes and/or affected by poor sequence quality. Amino acid pattern analyses of high throughput sequencing-derived mtCOI gene from 24 ‘*tabaci*’ and ‘non-*tabaci*’ species enabled differentiation between NUMTs/pseudogene-affected and likely real mtCOI sequences, and that the SSA4, SSA5/SSA8, AsiaII-2 and AsiaII_4 species were NUMTs/pseudogenes artefacts. Intra-specific uncorrected nucleotide distances (*p*-dist) from our up-dated dataset ranged from 0-1.98%, inter-specific *p*-dist within phylogenetic clades ranged between *ca.* 2.5 and 8%, and 8 and >19% for species between phylogenetic clades. Differentiating between closely related species could therefore utilise an ‘average’ *p*-dist of 2.5%. Despite the smaller *B. tabaci* mtCOI dataset, six putative new species were identified. Adoption of our standardised workflow and up-dated mtCOI clean dataset could facilitate better diagnostics of *B. tabaci* and ‘non-*tabaci*’ cryptic species.

## Introduction

Accurate delimitation of cryptic insect species complex, such as agriculturally significant pests, underpins the development of effective management strategies and biosecurity preparedness. The whitefly *B. tabaci* is a cryptic species complex of over 38 polyphagous sa*p*-sucking species^1-3^, with members that are indigenous to all continents except Antarcti*ca.* Species such as the sub-Saharan African (SSA) cassava whitefly species, the *B. tabaci* Mediterranean (‘MED’) species, and *B. tabaci* Middle East Asian Minor 1 (‘MEAM1’) have been responsible for major crop losses around the world^3-6^. The key challenge in identifying the different members of the complex is that they lack any reliable morphological differences. As a result, molecular markers have been used to identify the different members of the complex. The most widely used of these is mitochondrial COI and for the most part, the 3’ portion of the gene. This has seen a massive amount of research effort that resulted in 5,175 partial (*ca.* 300 to >657bp) mtCOI sequences being deposited in public DNA databases (e.g., in GenBank, accessed 22-June-2019).

While these efforts have contributed to diverse global species being surveyed and characterised for their partial mtCOI genes, these publicly available sequences (e.g., from GenBank; also^7, 8^) are also often affected by poor quality issues. For example, NUMTs (nuclear mitochondrial sequences) have been misidentified and published in high frequencies^9^ which resulted in conflicting estimates of inter-species nucleotide distances^2,10^ and erroneous molecular species diagnosis (e.g., MEAM2, see Dinsdale et al^10^ *cf*. Tay et al.^11^; SSA4 see Berry et al.^12^ and Wosula et al.^13^ *cf*. Elfekih et al.^14^).

Amino acid composition of various mitochondrial DNA genes (e.g., cytochrome oxidase *b* (cyt *b*)^15^, the mtCOI) have been shown to remain relatively conserved across evolutionary time at both intra- and inter-species (especially between closely related species) levels (e.g., in Lepidoptera^16^; between various *Bemisia* species^9^; in crustacean^17^; see also Hebert et al.^18^). The patterns of gene region amino acid conservation, codon usage patterns, and nucleotide position composition mutation biases^17,19,20^ can be used to identify NUMTs. Increasing availability of high-throughput sequencing (HTS)-derived mitochondrial DNA genomes (mitogenomes)^9,11,21-24^ from globally diverged cryptic *B. tabaci* species complex represent valuable genomic resources that could assist with the identification of NUMTs^11^. Significant substitutional changes of amino acid residues, especially among related species across highly conserved protein structural locations, could also indicate poor sequence quality due to either NUMTs, PCR-artefacts (i.e., pseudogenes due to chimeric/assembly errors) and/or sequences with poor editing quality.

Difficulties to identify NUTMs and pseudogenes have significantly and negatively impacted research efforts to better manage the *Bemisia* cryptic species pest complex^25^, leading to misinterpretations of genome-wide SNP data^13^ and understanding of evolutionary genomics involving whole genome sequencing^26^. Although the recently published mtCOI dataset with 513 *B. tabaci* sequences^8^ aimed to address molecular diagnostic needs and species identification with efforts to remove insertion-deletion (indel)-affected sequences from the original dataset^8^, indel-affected sequences nevertheless remained present, while amino acid substitution patterns in sequences unaffected by indels were not examined. With the confusion surrounding the *B. tabaci* cryptic species nomenclature^1,4,27,28^, the global research community will benefit from this well-curated mtCOI dataset and a standardised workflow to assist with future molecular taxonomy, species diagnostics, and evolutionary genomics studies of the *Bemisia* whitefly species complex.

In this study, we propose a standardised analysis workflow to enable potential NUMTs beyond indel-associated PCR amplification and sequencing errors to be more easily identified. We demonstrate this on the *B. tabaci* mtCOI dataset that had previously been assessed for indels and premature stop codons^8^. We re-calculate the intra- and inter-species uncorrected pairwise nucleotide distances (*p*-dist), re-evaluate ‘genetic gaps’ for species boundary, and infer phylogenetic relationships for the ‘*Bemisia tabaci*’ cryptic species complex based on the curated dataset, and show that novel cryptic species could be overlooked in a dataset that is impacted by poor sequence data.

## Results

We detailed a workflow (Fig. 1) outlining a proposed standardised analytical procedure to identify and exclude potential NUMTs. We also provided an up-dated version of the partial mtCOI dataset as Suppl. Data 1. The dataset also included partial mtCOI of *B. tabaci* cryptic species that were from published mitochondrial DNA genomes obtained via HTS platform (i.e., Asia I^23^; MEAM1^11^; *B. emiliae* (AsiaII_7) and B. sp. ‘JpL’^24^; NW1, MED and MEAM1^21^; NW^22^; SSA1 and SSA2 (MK940753 and MK940754, this study), but included also mitogenomes obtained via Sanger long-sequencing approach (i.e., NW^29^; MED^30^; *B. afer*^31^), partial mtCOI sequences of novel Australian *Bemisia* species (MN056066, MN056067, MN056068), selected African *Bemisia* species^3^, and selected ‘AsiaII’ and ‘China’ *B. tabaci* cryptic species^2^. NUMTs-free outgroups were also included in the up-dated dataset. This up-dated dataset represents a conservative set of curated partial mtCOI sequences, i.e., sequence with at least one INDEL identified when aligned against the HTS-mtCOI reference dataset were excluded without further analysis of amino acid substitution patterns.

**Fig. 1:**
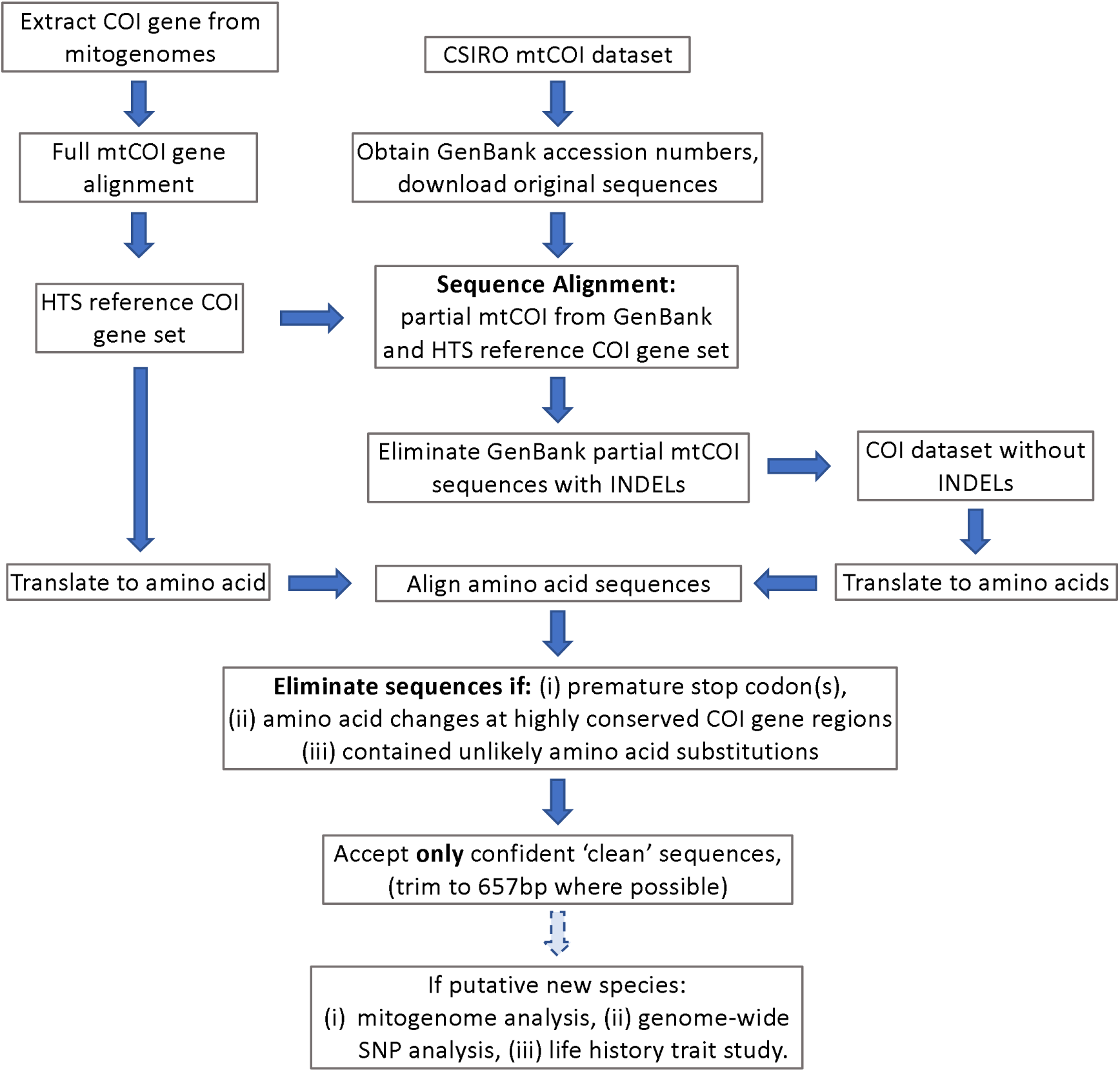
Schematic diagram outlining the proposed analysis workflow of utilizing high-throughput sequencing (HTS) generated *B. tabaci* cryptic species mitogenomes for identifying potential NUMTs and pseudogenes in partial mtCOI gene sequences. **Note:** Translation to amino acids uses the invertebrate mtDNA genetic codes. Faded blue arrow indicates suggested steps to further confirm putative *B. tabaci* cryptic species status. The HTS reference mtCOI gene set in this study involved *B. tabaci* and ‘non-*tabaci*’ cryptic species (i.e., ‘MEAM1’, ‘MED’, ‘MED-ASL’, ‘IO’, ‘AsiaI’, ‘AsiaII_7 (*B. emiliae*)’, ‘NW2’, ‘NW1’, ‘Australia’, ‘SSA1’, ‘SSA2’, ‘*B. afer*’, and ‘*B*. sp. JpL’; Suppl. Data 2; Suppl. Data 3). With the exception of the *B. afer*, one *B. tabaci* ‘NW’ (AY521259) and one *B. tabaci* ‘MED’ (JQ906700) mitogenomes which were generated via Sanger sequencing, all remaining mitogenomes were obtained via HTS platform. Analyses of amino acid changes between the HTS-mtCOI reference gene set and the Sanger sequenced partial mtCOI genes involved determining base substitutions that result in significant biochemical property changes of the amino acid residues (e.g., hydrophobic/hydrophilic, polar/non-polar, charged/un-charged). Partial mtCOI gene sequences from Sanger sequencing platform were not excluded if amino acid substitutions have not caused significant biochemical property changes.

Alignment of the HTS-generated partial mtCOI reference gene set (Fig. 2) showed that across evolutionary diverged *Bemisia* species including the cryptic *B. tabaci* species complex, highly conserved gene regions were evident. Comparisons between the various *B. tabaci* cryptic species showed that, with the exception of MEAM1 where five sequences (GU086346, GU086358, GU086351, GU086353, GU086357) showed a consistent amino acid change at an otherwise conserved amino acid position 97 in the mtCOI partial gene, all sequences that had amino acid substitutions occurring in the inter-species highly conserved COI gene regions were excluded, as shown for the MED species (Fig. 3). We acknowledge that amino acid changes could potentially occur at other conserved COI gene regions especially if changes were to occur between residues with similar biochemical properties, but without solid evidence such as from HTS-generated data we exercise caution by excluding such sequences. Table 1 provides a summary of the total number of input partial mtCOI sequences from the mtDNA dataset reported by Boykin et al.^8^ for each species, and the number of confirmed clean sequences, potential NUMTs and related observations from this study. A full list of related GenBank accession numbers for each of the category is also provided (see Suppl. Table 1).

**Table 1:**
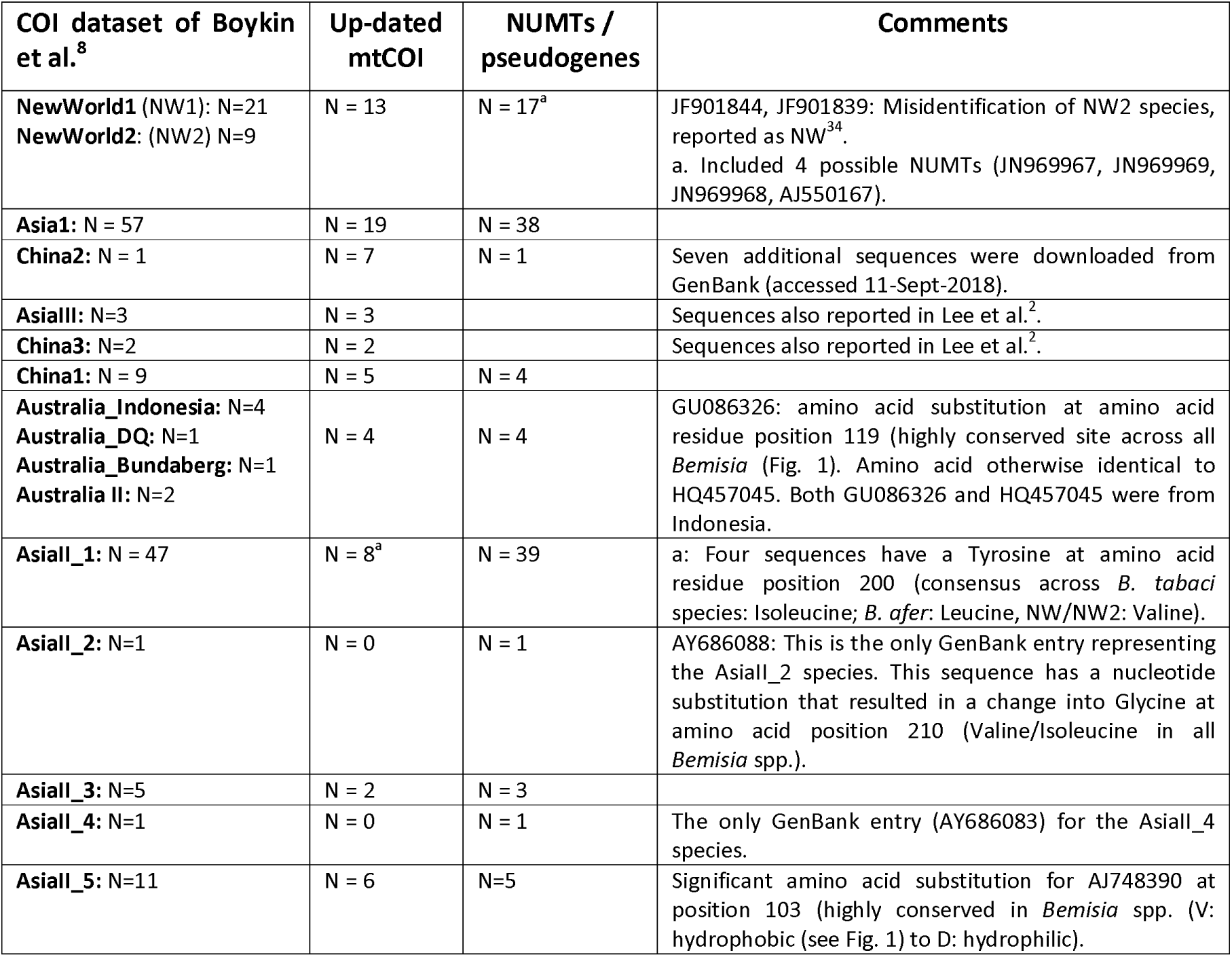
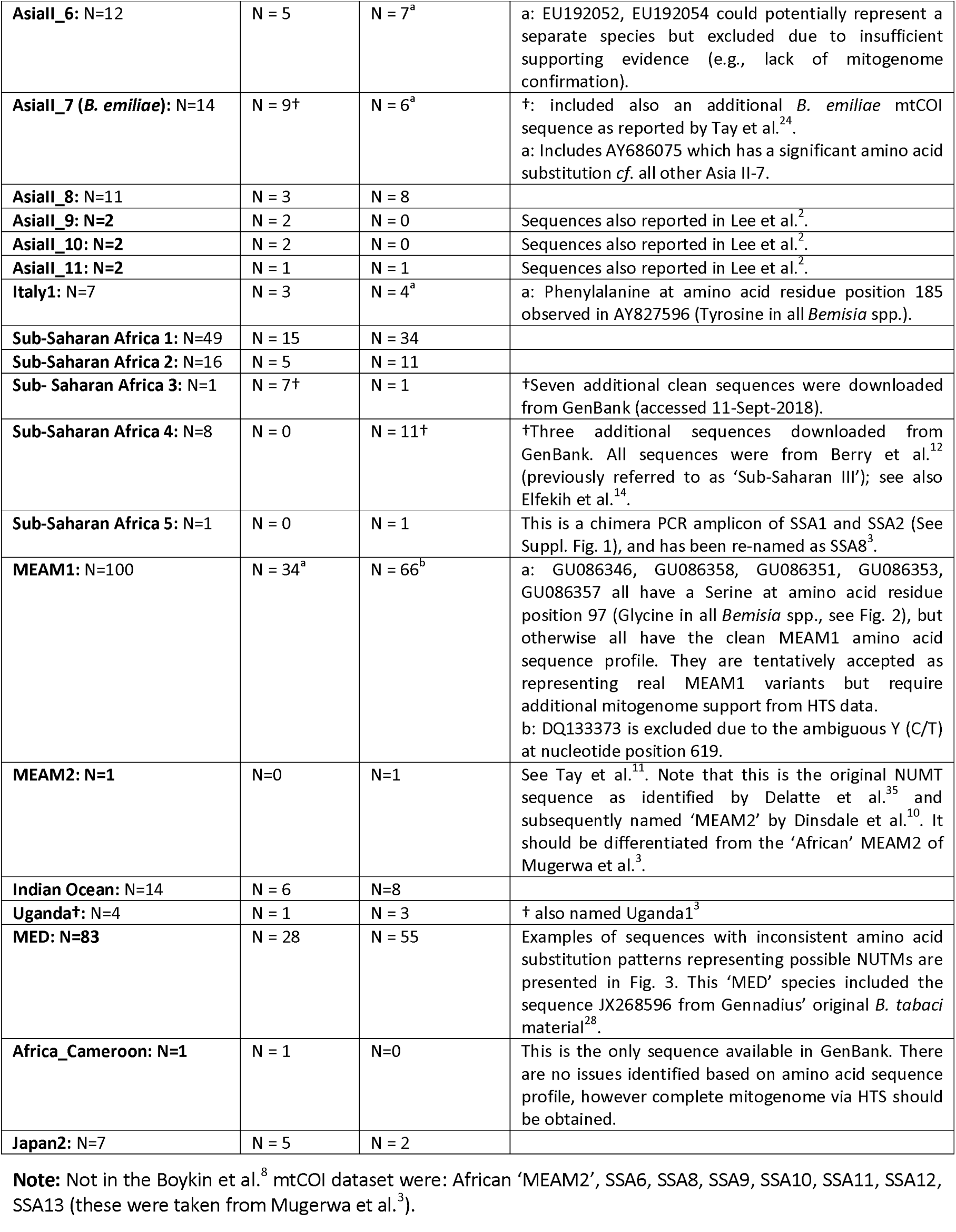
Summary of the *Bemisia tabaci* cryptic species partial mtCOI dataset v2^8^ against the current up-dated version of the dataset. Number of input sequences, up-dated partial mtCOI sequences with consistent amino acid profiles against mtCOI amino acids as obtained through HTS platform (see Fig. 2) and identified NUMTs and pseudogenes (i.e., chimeric genes identified as PCR artefacts due to contig assembly from amplification of two different *B. tabaci* cryptic species), and potential NUMTs. Relevant comments relating to observations of NUMTs and pseudogenes/potential NUMTs are also provided. For complete GenBank accession numbers see Suppl. Table 1.

**Fig. 2:**
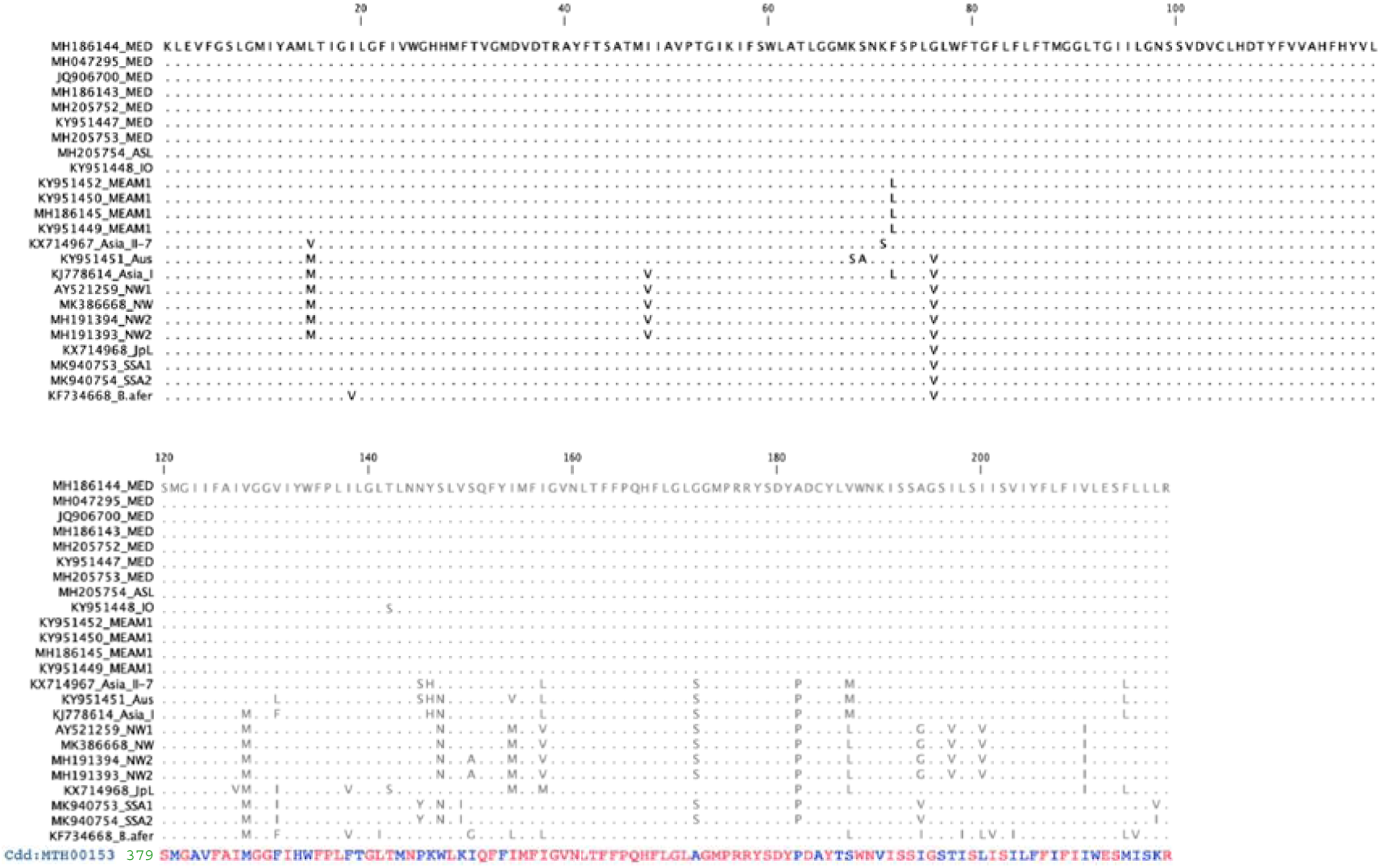
Partial (657bp) mitochondrial DNA cytochrome oxidase subunit I (mtCOI) gene translated amino acids alignment for all published *Bemisia tabaci* cryptic species including *B. afer* and *B*. sp. ‘JpL’ obtained through high throughput sequencing methods. Exceptions are *B. tabaci* ‘MED’ (JQ906700)^30^ and *B. tabaci* ‘NW’ (AY521259)^29^ which were obtained via long PCR and Sanger sequencing. Conserved domain for the heme_Cu_Oxidase_I superfamily for arthropods (MTH00153 cytochrome c oxidase subunit I; NCBI accessed 29-Dec-2018) was based on blastp search using the MED mtCOI amino acid sequence (MH186144; nucleotide positions 782-1438) as input query protein sequence. MTH00153 identified arthropod conserved and variable amino acid residues are in red and blue letter codes, respectively. The list of *B. tabaci* cryptic species mitogenomes/mtCOI gene used represent a divergent timeline of at least 45-168.5 MY^32,33^. **Note:** List of mitogenomes used: AY521259 NW^29^; JQ906700 MED^30^; MH191394 NW2, MH191393 NW2, MH186144 MED, MH047295 MED, MH186143 MED, MH186145 MEAM1^21^; MH205754 ASL, MH205753 MED, MH205752 MED^9^; KY951451 Aus, KY951448 IO, KY951447 MED, KY951452 MEAM1, KY951450 MEAM1, KY951449 MEAM1^11^; KX714967 *B. emiliae* (AsiaII_7)^24^, MK940753_SSA1; MK940754_SSA2; KX714968 *B.* sp. ‘JpL’^24^; KJ778614 AsiaI^23^; MK386668 NW^22^, and KF734668 *B. afer*^31^.

**Fig 3:**
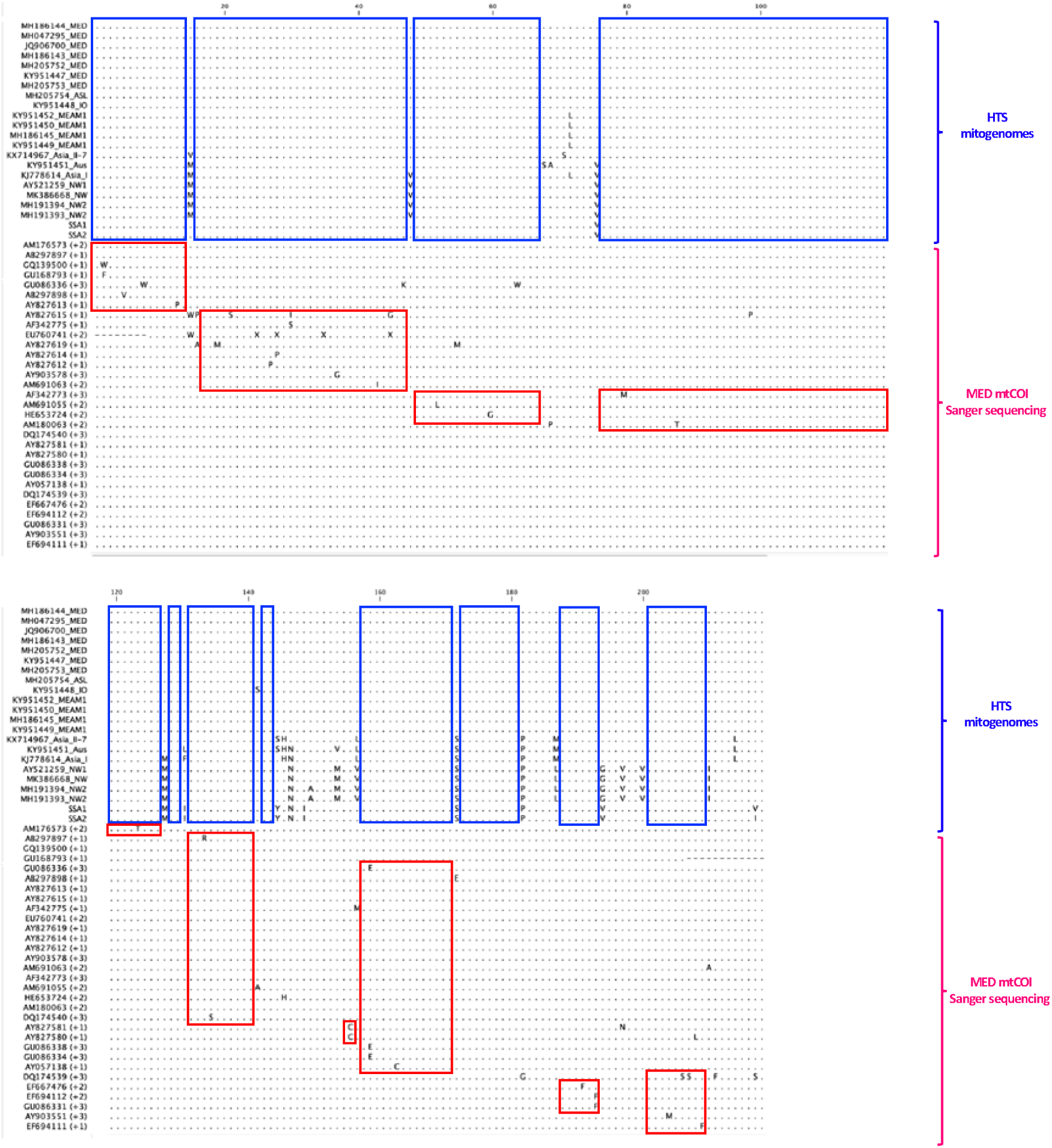
Alignments of all MED-NUMTs with no evidence of INDELs against corresponding partial mtCOI amino acid residues of related *B. tabaci* cryptic species as obtained via HTS methods. Blue boxes are conserved regions detected in the partial mtCOI region of the *B. tabaci* and ‘non-*tabaci*’ cryptic species as well as from the *Bemisia* ‘JpL’ species endemic to Japan and the Korean peninsula, as obtained by HTS methods. Red boxes are amino acid residue substitutions that were detected in putative NUMT sequences of *B. tabaci* MED species. These putative NUMT sequences showed greater intra-species amino acid changes than were present at the interspecific level that corresponded to over 45-168.5 MY^32,33^ of species divergence.

### Uncorrected intra- and inter-specific pairwise nucleotide distances (*p*-dist)

Intra-species *p*-dist was calculated to range from 0-1.9% in all phylogenetic species clades (Fig. 4; grey bars). Cut-off for intra-species diversity was therefore tentatively suggested at *ca.* 2% (light red arrow, Fig. 4). Three separate inter-species ‘genetic clusters’ (i.e., Clusters I, II, and III) were detected with Cluster I representing closely related species, such as between ‘MED’, ‘MED-ASL’ and ‘MED-Burkina Faso’ clades within the MED complex, and between the two ‘NW2’ sister clades (Fig. 5). For species within Cluster I, average *p*-dist of 2.5% should be used, and should incorporate the current up-dated dataset and any future data being generated, while setting the interspecies *p*-dist at *ca.* 3.5% as proposed by Dinsdale et al.^10^ should be considered for MED and NWs species complexes. Cluster II showed *p*-dist range of species at the *within* phylogenetic clade (i.e., *ca.* 3.5%-8%), and Cluster III contained species between sister clades (*ca.* 8 to ≥19%).

**Fig. 4:**
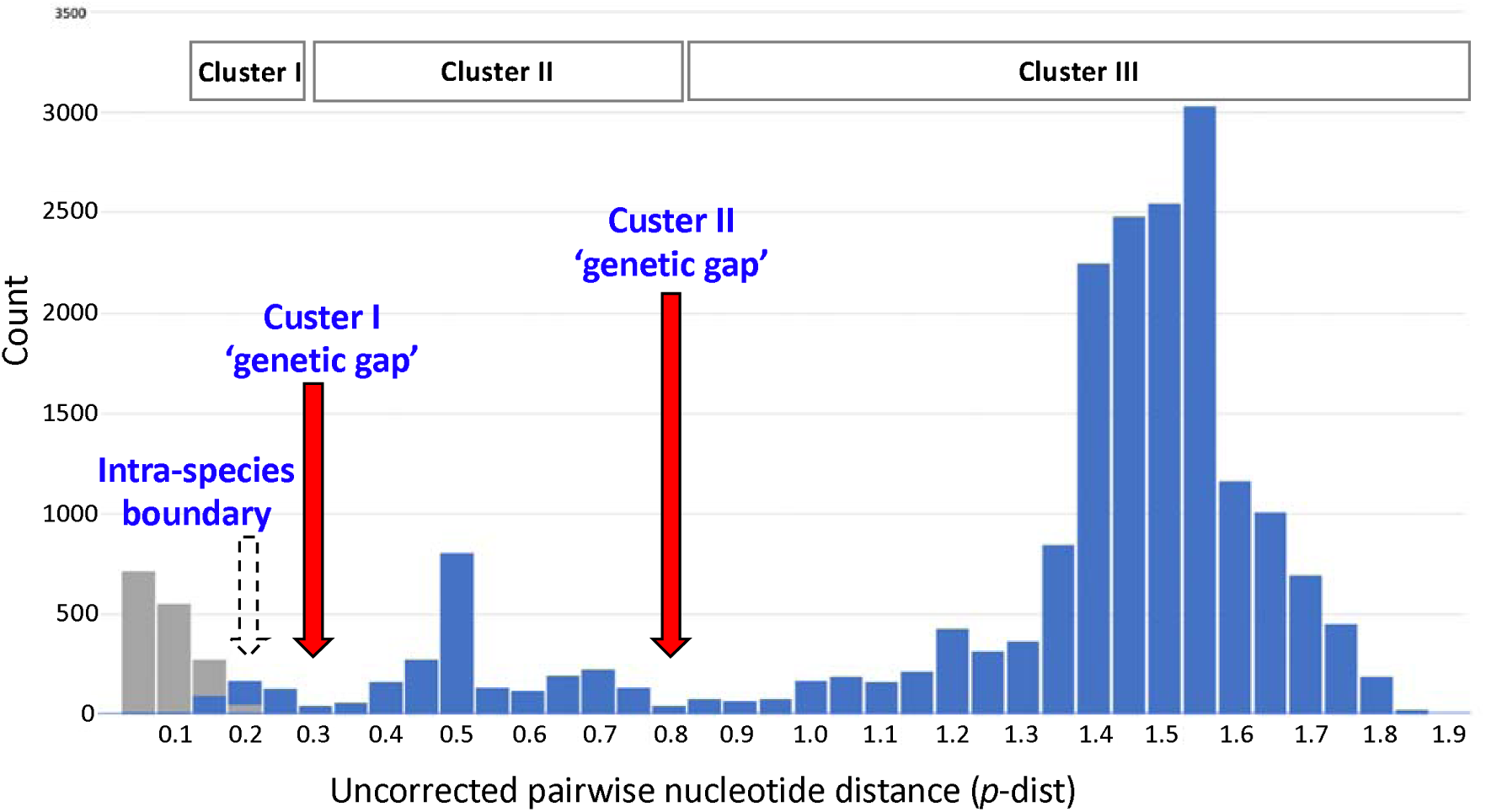
Uncorrected pairwise nucleotide distances (*p*-dist) of partial mtCOI (657bp) from the current up-dated *B. tabaci* cryptic species database. Grey bars are intra-specific comparisons which showed a maximum *p*-dist value of 0.198% as indicated by the lighter red arrow. Blue bars are inter-specific comparisons showing three genetic clusters (I, II, III). Cluster I included *p*-dist predominantly within the ‘MED/MED-Burkina Faso/MED-ASL’ species (i.e., 0.008 to 0.26 [MED vs. MED-Burkina Faso]; 0.0137 to 0.0274 [MED-Burkina Faso vs. MED-ASL]; 0.0152 to 0.032 [MED vs. MED-ASL]), and the ‘NW2/NW2-2’ species (0.0244 to 0.0304). Cluster II showed distribution of *p*-dist values between species within clades, and Cluster III showed *p*-dist values of species between clades. For highly related species within Cluster I we advocate that the ‘average’ *p*-dist’ should be used.

**Fig 5:**
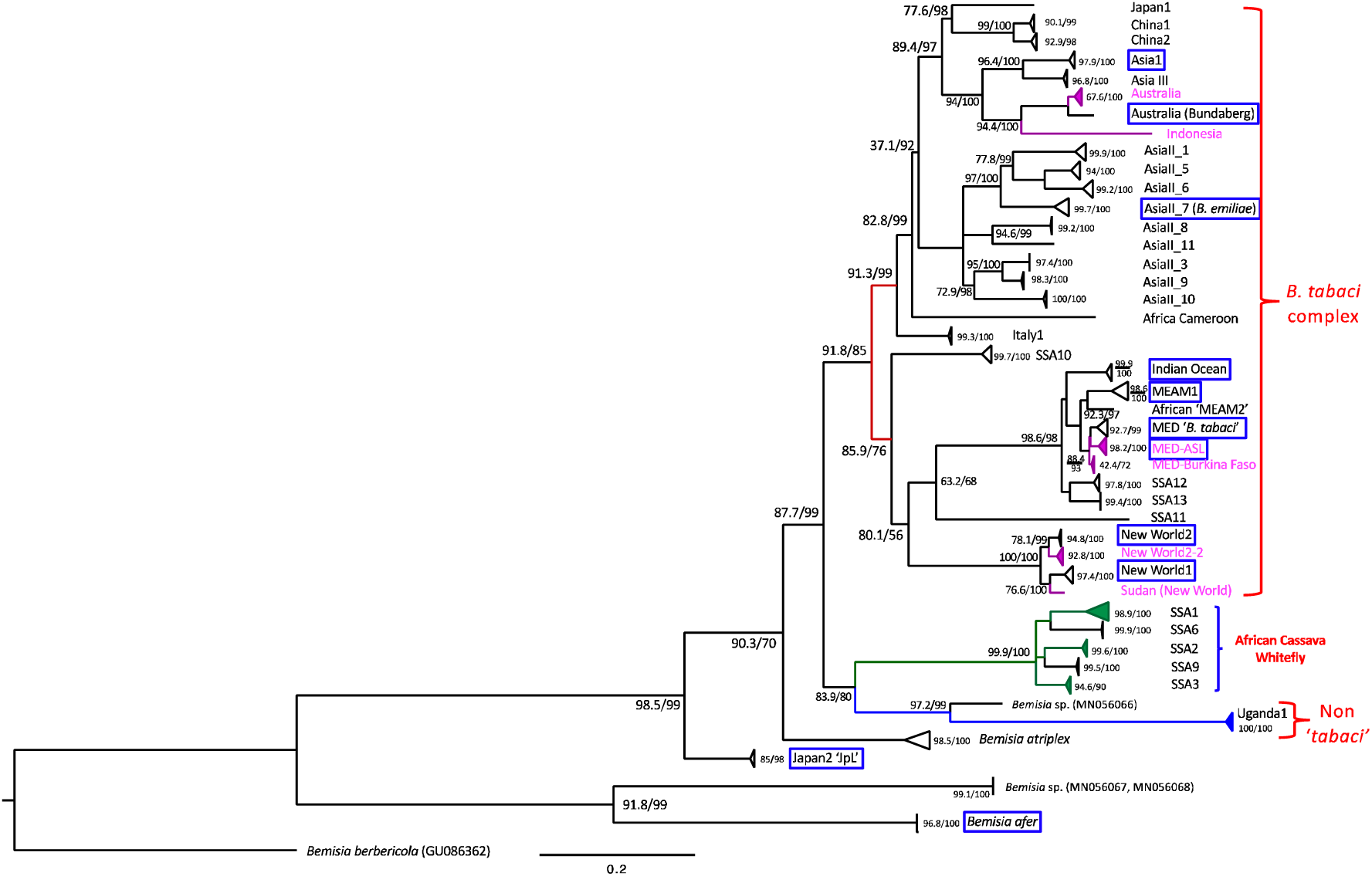
Maximum Likelihood (ML) phylogeny of *Bemisia tabaci* cryptic species of the up-dated mtCOI dataset after removal of potential NUMTs, as inferred using IQ-TREE. Based on the partial mtCOI gene, the ‘*B. tabaci* cryptic species complex’ potentially excludes the *B.* sp. ‘JpL’ and various sub-Saharan African (SSA) species (i.e., SSA1, SSA2, SSA3, SSA6, SSA9). Potential new species are in pink colour. With the exception of MED-ASL which has been shown to represent a cryptic species within the MED species complex through full mtDNA genome and mating compatibility analyses^9^, all other new species especially NW2-2, Sudan (New World), MED-Burkina Faso, Australia, and Indonesia, could remain possible PCR artefacts or NUMTs. Australia (Bundaberg), New World 2, and all species boxed in blue have had full mitogenomes reported. The AsiaII clade included six separate AsiaII sub-clades^1^ of AsiaII_1, AsiaII_5, AsiaII_6, AsiaII_7, Asia II_3 and Asia II_8 based on genetic gaps of >8% as separating between clades. Out groups are *B. berbericola* (HQ457046), *B. afer* (AF418673, KF734668) and two Australian ‘non-*tabaci*’ *Bemisia* species (*Bemisia*_sp_PDB_2-1 (MN056067); *Bemisia*_sp_PDB_2-2 (MN056068)).

### Identification of pseudogene/NUMTs-misidentified *B. tabaci* cryptic species

We excluded five species from the Boykin et al.^8^ dataset as being associated with NUMTs/pseudogenes. These five putative species included MEAM2 previously shown^11^ to be associated with MEAM1 NUMTs; SSA4, reported^14^ to be associated with SSA2 NUMTs; and SSA5, AsiaII_2, and AsiaII_4. The SSA5, AsiaII_2 and AsiaII-4 species is each being represented by a single sequence and have only been reported once^3,36^. In the case of SSA5, this sequence was shown to consist of 407bp of 5’ COI region of SSA1 and 327bp of 3’ COI region of SSA2 (see Suppl. Fig. 1). Both AsiaII_2 and AsiaII_4 sequences have multiple amino acid substitutions at conserved COI regions and were therefore excluded from the up-dated mtCOI dataset.

The identification of pseudogenes and NUMTs were based on comparisons with HTS-generated partial mtCOI sequences (Suppl. Data 2) and with conserved amino acid residues in mtCOI gene regions (Suppl. Data 3; noteworthy cases for a number of species are further described below.

#### I. NW, NW2 & Sudan

In the New World clade two species defined as New World (i.e., ‘NW’, also sometimes referred to as ‘NW1’) were previously reported^1^. We eliminated 11 sequences as potential NUMTs from the 30 partial mtCOI sequences (see Suppl. Table 1) due to significant amino acid changes at various positions. Two NW individuals (JF901844, JF901839) identified by Alemandri et al.^34^ represented possible misidentification with uncorrected nucleotide distances being closer to NW2 (2.54%) than to NW (4.581%). Our redefined species boundary based on *p*-dist nucleotide similarity values therefore also identified these two misidentified individuals (JF901844, JF901839) as belonging to a putative new NW2 species (Fig. 5, as ‘NW2-2’), although these two mtCOI sequences were incomplete at the 3’ end (i.e., only 633bp instead of 657bp), and further support based on HTS-derived mitogenomes will be needed. Diversity of *B. tabaci* cryptic species in the New World is therefore potentially higher than presently recognised. Presence of novel cryptic *B. tabaci* species in South America were previously also reported^37^. The remaining NW2 and NW1 species defined based on the up-dated mtCOI dataset are supported by the availability of draft and/or fully published mitochondrial DNA genomes^21,22^.

Reanalysis of the Sudan *B. tabaci* cryptic species partial mtCOI sequence (EU760727) did not exclude it as a potential NUMT sequence. Based on our *p*-dist criteria and inference from phylogenetic analysis (see below), our findings concurred with De Barro^38^ that this Sudan individual represented a putative novel species basal to the NW and NW2 species, supporting the African origins of New World *B. tabaci* cryptic species clade hypothesis^3^. Complete mitogenome and integrated studies involving life history traits will be needed to further support the novel species status of this Sudan *B. tabaci* cryptic species.

#### II. Australia & Australia/Indonesia

Of the 8 sequences that belonged to the Australia & Australia/Indonesia clade, four (GU086325, GU086326, GU086327, DQ130052) were ascertained as potential NUMTs due to significant amino acid substitutions. Removing these four sequences resulted in only one sequence with Indonesia origin (HQ457045) from those that originated only from Australia. The HQ457045 Indonesian origin sequence further matched one unpublished Indonesian *B. tabaci* cryptic species sequence (AB248263) deposited in GenBank at 100% nucleotide identity. Searches against GenBank using clean-up ‘Australia’ sequences resulted in five additional NUMT-free sequences (KY951451, KF452532, JX416166, JX416167, JX416160) being identified, of which one belonged to a HTS-generated draft complete mitogenome sequence (KY951451)^11^. Removal of NUMT sequences in this clade therefore separated Australian endemic species from a native Indonesia species. Phylogenetic reanalysis of the ‘Australia’ and ‘Australia/Indonesia’ partial mtCOI genes identified two sister clades within the Australia sequences with uncorrected *p*-dist between these two Australian species estimated at 3.35%, while the previous ‘Australia/Indonesia’ clade now contained only Indonesian ‘B. *tabaci*’ cryptic species and has been re-named ‘Indonesia’ (Fig. 5).

#### III. MED

Based on the uncorrected *p*-dist values, three genetic groups were detected in the current up-dated *B. tabaci* ‘MED’ sequences. These three genetic groups separated Gennadius’ (1889) B. *tabaci*^28^ from the ‘African Silver Leaf’ (ASL) ‘MED’ *B. tabaci* cryptic species^9^, and also a third group reported only from Burkina Faso (i.e., MED-Burkina Faso). The MED-Burkina Faso group currently does not have life history and mitogenome data to support its species status. A MED mitogenome from an individual that originated from Burkina Faso was previously reported^11^ however this individual (KY951447) has a mtCOI partial sequence that best matched Gennadius’ B. *tabaci*^28^ (JX268596) with sequence identity of 99.34%.

#### IV. MEAM2 and ‘Africa-MEAM2’

The MEAM2 sequence AJ550177, originally referred to as ‘Ms’ biotype^35^, and all subsequent MEAM2-related sequences in GenBank, i.e., (AB308110)^39^; sequences KK103B, KK104A, KK104B by Karut et al.^40^; KY951453, KY951454, KX234912, KX234913, KX234914^11^; KX679576 (GenBank unpublished sequence from Iraq); FJ939600, FJ939602 (GenBank unpublished sequence from Egypt)) have been shown^11^ to be NUMTs. The original AJ550177 MEAM2 should be differentiated from the ‘MEAM2’ from Africa (KX570778 – KX570784), so named because Mugerwa et al.^3^ determined a nucleotide distance of <3.5% when compared with the pseudogene MEAM2 AJ550177 instead of first excluding the MEAM2 pseudogene-misidentified species status prior to comparisons being carried out. Species status of this ‘African-MEAM2’ of Mugerwa et al.^3^ will need to be further confirmed by HTS-derived mitogenome.

#### V. SSA4 and SSA5

The SSA4 species originally reported by Berry et al.^12^ and named ‘Sub-Saharan III: Western Africa-Cameroon/Cassava’ has been shown^14^ as being of NUMTs origin from the genome of SSA2. The single SSA5 sequence (AM040598) recently renamed as SSA8^3^ showed strong signatures of being a chimeric contig (i.e., a pseudogene) from SSA1 and SSA2 (see Suppl. Fig. 1).

### Mitogenomes as resources to identify potential NUMTs and pseudogenes

A total of 20 *Bemisia tabaci* cryptic species mitogenomes, one B. sp. ‘JpL’ and one African *B. afer* mitogenomes have been published to-date. These 20 *B. tabaci* cryptic species included seven individuals of the *B. tabaci* MED species from Brazil^21^, China^30^, Spain, Israel^9^ and Africa^11^; four individuals of *B. tabaci* MEAM1 from Peru^11^ and Brazil^21^; two individuals of NW2 from Brazil^21^, two individuals of *B. tabaci* NW from the US^29^ and Barbados^22^; one *B. tabaci* Aus from Australia, and one *B. tabaci* Indian Ocean (IO) from La Réunion^11^, one *B. tabaci* Asia I^23^, one *B. emiliae* (*B. tabaci* Asia II-7) from Sri Lanka^24^; and one *B. tabaci* MED-ASL^9^; and the full mtCOI gene of the SSA1 (MK940753) and SSA2 (MK940754) species from Uganda.

These cryptic *B. tabaci* species represented species from phylogenetic clades of Africa/Middle East/Asia Minor; New World; Asia II-7; Asia 1, Australia, sub-Saharan African, and Asia I, and evolutionary divergence time line that were estimated at 44-66 million years based on partial mtCOI sequence analysis^32^ to 42.9 – 168 million years just between the ‘MED’ and ‘MEAM1’ species^33^ based on whole genome sequence analysis (*cf*. Chen et al.^26^). Alignments of translated amino acids from this 657bp region showed that significant conservation of amino acid residues occurred in the first 125 amino acids except for 8 residues (Fig. 2). In the region from amino acid 126 to 218 there were greater number of variable sites although regions of high amino acid residue conservation were still evident. Figure 2 included also alignment of corresponding mtCOI gene region of *B. afer* that similarly showed high level of amino acid conservation, suggesting that while various cryptic *B. tabaci* species were yet to be surveyed by HTS (e.g., the various AsiaII clades; the Italy clade, the Uganda Clade, the China clade) for the mtCOI gene region and their amino acid compositions, it would seem likely that similar high levels of amino acid conservation could be expected.

Variable sites across this partial mtCOI gene region was assessed for potential effects from observed substitutional changes (e.g., between hydrophobic amino acid residues; between amino acids with polar uncharged side chains, etc.). Significant changes resulted in amino acids with different biochemical properties likely reflected nucleotide substitution changes at the highly conserved 2^nd^ codon position. Examples of some of these changes (e.g., an unexplained amino acid substitution that resulted in a Methionine (M) to Lysine (K) change in GU086336, amino acid position 77; Fig. 3) that represent significant biochemical property changes, as well as greater intra-species amino acid substitution rates as compared with inter-species level changes, can been seen in Figure 3. Significantly for these sequences, majority of detected amino acid substitutions also occurred in the mtCOI gene regions that had remained conserved for over 40 to 168.5 million years^32,33^.

### Phylogenetic analysis from the up-dated mtCOI dataset

IQ-Tree phylogenetic analysis (best-fit model of substitution according to BIC: TIM2+F+I+G4; BIC score: 22638.5567; model for rate heterogeneity: I+G with four categories, proportions of invariable sites: 0.5218; Gamma-shape alpha: 0.9677) showed strong overall support of SH-aLRT and/or UFBoot for nodes separating putative cryptic species, supporting overall integrity of our up-dated mtCOI dataset. Weak node support values (SH-aLRT: < 80%; UFBoot: <95%) were also observed (e.g., branch node separating New World and SSA11; branch node separating SSA11 and MED/IO/MEAM/SSA12/SSA13) indicated lower probability that the corresponding clade is true, and likely reflected the short length of the barcoding region used as being less effective for inference of unbiased evolutionary relationships between targeted species. Phylogenetic analysis also found evidence to the likely presence of two Australian species in addition to the Indonesian species, with the Indonesian species previously being called ‘Australia/Indonesia’^1^. Note that the current up-dated mtCOI dataset contained only one haplotype from each of these species clades: Indonesia, Sudan, Australia (Bundaberg), Japan1, African Cameroon, ‘African MEAM2’, SSA11, and Sudan (New World). We were unable to confirm if these sequences were unaffected by NUMTs/pseudogenes although species such as the ‘Australia (Bundaberg)’ species (Fig. 5; Suppl. Data 2: GU086328) has been sequenced for its complete mitogenome (KY951451)^11^.

Current phylogeny of the *B. tabaci* cryptic species complex suggested Africa being the *B. tabaci* cryptic species complex’s ancestral range with African cryptic *Bemisia* species (e.g., Uganda/SSA1-3, SSA6, SSA9 clades) as a sister clade to all ‘B. *tabaci*’ cryptic species clades (Fig. 5). The African/Middle East/Asia Minor clade that included the Mediterranean species complex, MEAM1, ‘African MEAM2’, and Indian Ocean species, and the clade that included the SSA12 and SSA13 species, shared a most common recent ancestor with SSA11, and which in turn formed a sister clade with the New World/Sudan species, while the SSA10 species clade was shown to be most ancestral. The sub-Saharan African clade that included the SSA1, SSA2 and SSA3 species include also SSA6 and SSA9. The European (e.g., Italy1) clade is basal to the various AsiaII clades and the China/Indonesia/Australia/AsiaI evolutionary lineages. Its basal evolutionary relationship to the African Cameroon species (EU760739) is difficult to explain as also reflected by the weak SHaLRT node support value for this single African Cameroon sequence.

Node support for the placement of the Uganda1/*B*. species (MN056066) and SSA1, SSA2, SSA3, SSA6, SSA9 branch is low for the UFBoot value (i.e., <<95%) however the placement of these two clades as being sister clades to the *B. tabaci* cryptic species complex was high (91.8%) for the SHaLRT test, and approaching significant (i.e., 95%) for the UFBoot test (observed at 85%). The Uganda1 species has been reported as a ‘non-*tabaci*’ species but with adult morphology similar to B. *tabaci*^3^. Identification based on fourth instar pupa morphologies will be needed to further ascertain if the Uganda1 species belonged to the *B. tabaci* cryptic species complex. Current analysis therefore could not confirm if the Uganda1/*Bemisia* sp. from Queensland Australia (MN056066) and African cassava whitefly species complex (i.e., SSA1, SSA2, SSA3, SSA6, SSA9) should be considered as part of the *B. tabaci* species complex.

Species diversity within the MED complex have short branch lengths although complete mitogenomes and mating studies have supported the distinct species status between MED-ASL and MED^9^. Node support values for the ‘MED’ and ‘ASL’ were strong (92.7/99; 98.2/100, respectively) while node support values for clustering of the ‘MED-Burkina Faso’ group were weak.

## Discussion

Our proposed analyses of considering nucleotide composition biases (i.e., as reflecting potentially unlikely nucleotide substitutions at the highly conserved second codon position), by identifying synonymous and non-synonymous amino acid changes, as well as taking into consideration significant intra-species substitutions that occurred in evolutionary conserved mtCOI gene regions, will increase the toolset for identifying potential NUMTs in the *B. tabaci* cryptic species complex. We also demonstrated the value of HTS-generated mitogenomes as significant resources to help identify potential pseudogenes and NUMTs in the highly cryptic hemipteran insect *Bemisia tabaci* species complex. The HTS-generated partial mtCOI gene region of these diverse species enabled regions of significant amino acid conservation and patterns of unexpected amino acid substitutions at intra-species level to be identified. We have shown that 64.9% of the *B. tabaci* cryptic species mtCOI dataset^8^ as potential NUMTs and pseudogenes. Similar to findings for MEAM2^11^ and Elfekih et al.^14^ for SSA4^13,26^, NUMTs-affected ‘pseudo’ species included also SSA5, Asia II_2, and Asia II_4.

Based on our up-dated dataset, we recalculated intra-species uncorrected nucleotide distances (*p*-dist) to range from 0% to *ca.* 2%, beyond which would indicate separate species. However, for closely related species such as within the MED complex (MED, MED-ASL, MED-Burkina Faso), and the NW2/NW2-2 species complex, species boundary required average *p*-dist of 3.5% as proposed^10^. Potential novel species within the *B. tabaci* cryptic species complex were also identified, including: (i) a Burkina Faso ‘MED’ species basal to the ‘ASL’^9^ and the MED ‘real’ B. *tabaci*^28^ species; (ii) a new ‘Australia’ species that differed from the ‘Australia (Bundaberg)’ species, where its full mitogenome was previously reported^11^, (iii) a new ‘NW2’ species (i.e., NW2-2) that differed from the species where full mitogenomes had been sequenced^21^, and (iv) the removal of the Australia/Indonesia species clade that now includes only specimens from Indonesia (i.e., ‘Indonesia’ species). Our phylogenetic analysis of the new mtCOI dataset further supported the species status of the ‘Sudan’ species as being closely related to the NW/NW2 species clade, and queried the general acceptance of the cassava whitefly species complex (i.e., SSA1, SSA2, SSA3)^27^ as part of the ‘B. *tabaci*’ cryptic species complex.

Whilst our up-dated dataset identified four putative new species across the phylogenetic clades of the ‘B. *tabaci*’ cryptic species complex, confirmation of species status in these putative new species will require full mitogenome sequencing which would provide the first phylogenomic insights into species delimitation based on multigene inferences, as well as through life history traits and host plant preference studies^9,41-43^. Short branch lengths between species, such as between NW2/NW2-2, MED-Burkina Faso/MED/MED-ASL (Fig. 4) suggested that species status could be difficult to confirm, and genomic approaches including multigene phylogenies, full mitogenomes and genome-wide SNP analyses should be carried out in addition to life history studies. The usage of the short (657bp) partial mtCOI gene for species identification in the *B. tabaci* cryptic species complex remains highly attractive as tools for molecular diagnostics. Whilst the presence of INDEL’s and premature stop codons in the partial mtCOI gene enabled NUMTs and pseudogenes to be readily identified, NUMTs and pseudogenes could be missed if these signatures were located outside the sequenced region^11^, or where random nucleotide substitutions representing NUMTs have not led to pre-mature stop codons. Random nucleotide substitutions that might indicate potential NUMT, and that are associated with highly conserved functional genes, can also be detected based on codon positional bias, amino acid and nucleotide composition analyses^17,44,45^.

DNA barcoding among arthropods could lead to inaccurate spices richness estimation as NUMTs were coamplified^46,47^. A major contributing factor to the large proportions of NUMTs/pseudogenes in the *B. tabaci* mtCOI dataset potentially involved the widespread practice of utilising non-hemipteran COI primers (e.g., C-1-J2195/L2-N-3014)^12,48,49^. These ‘universal’ primers have not been designed for Hemiptera^3,11,50,51^, and the importance of using PCR primers of high efficacies developed for the *B. tabaci* species complex has been stated^3,50,52^. The inclusion of NUMTs/pseudogenes in evolutionary genetic studies^7,32^ could also potentially impact dating of species divergence when compared with estimation based on genome data^33^. Implications of failure to disentangle NUMTs/pseudogenes in these cryptic hemipteran whitefly species^13^ have also been shown to affect population genomic interpretations for the SSA cassava whitefly species in sub-Sharan Africa^14^, and failure to acknowledge^26^ could prolong further discrepancies.

Analyses of our up-dated mtCOI dataset demonstrated that in the majority of *B. tabaci* cryptic species, intra-species *p*-dist generally range from 0-1.9%, with individuals that differed by >2% to represent different species. However, in closely related species (i.e., the MED species complex; the NW2 species complex), the lower bounds of inter-species *p*-dist estimates overlapped the upper bounds *p*-dist of intra-species, with interspecies genetic gaps for these closely related species complex observed at *p*-dist of *ca.* 3%. The use of 4% genetic gap nucleotide distance^2^ represented an overly conservative species boundary that could undermine species delimitation, especially in life history trait studies by combining and treating of distinct species as one.

The advent of genome-based analytical options based on genome-wide SNP markers remained no panacea^13,26^ to solving the basic yet complex biological issues affecting the *Bemisia* cryptic species complex, when identification of NUMTs and pseudogenes remain significant challenging tasks to complicate result interpretation. With novel *B. tabaci* cryptic species still being identified, our workflow and dataset will represent an invaluable toolset to the global *Bemisia* research communities for improved mtCOI data management, molecular diagnostic, and development of biosecurity preparedness strategies for this highly invasive agricultural pest complex.

## Methods

We utilised the *Bemisia tabaci* mtCOI dataset of Boykin et al.^8^ (accessed 08-Sept-2017) for a proposed NUMT and pseudogene analysis workflow to generate a clean set mtCOI sequences. Original sequences from the National Centre for Biotechnology Information (NCBI) GenBank database were downloaded into the CLC Sequence Viewer 8 software (Qiagen Aarhus A/S). All published *B. tabaci* cryptic species mitochondrial DNA genomes were also downloaded (GenBank access date: 22-Oct. 2018) and the mtCOI gene extracted (see ‘HTS reference COI gene set’; Suppl. Data 2) followed by nucleotide sequence alignment. We also undertook HTS following the method described^23^ to generate the complete mtCOI gene for the SSA1 and SSA2 species in order to compare amino acid substitution patterns within the African cassava whitefly clade, based on Ugandan SSA1 and SSA2 individuals. The alignment parameters within the CLC Sequence Viewer 8 software for both nucleotides and amino acids were set as ‘Gap open cost: 10’, ‘Gap extension cost: 1.0’, and specifying ‘Alignment: very accurate (slow)’, followed by manual fine tuning where necessary.

We aligned all partial mtCOI gene sequences downloaded directly from GenBank against the HTS reference COI gene set, and eliminated all Sanger sequencing-generated sequences that contained indels. All indel-free sequences were translated to amino acid residues from appropriate codon positions using the invertebrate mitochondrial DNA genetic codes to: (i) identify potential premature stop codons, and (ii) enable amino acid residue alignment against the HTS reference COI amino acid dataset (Suppl. Data 3). We further eliminated all sequences where amino acid substitutions had occurred at highly conserved regions as identified within the trimmed HTS reference COI gene set. As a final check, all sequences that passed the INDEL and conserved amino acid region substitution tests were examined for presence of amino acid residues that might have violated permissible biochemical signatures, due typically to substitutions at the highly conserved second codon position. Sanger sequencing-generated nucleotides with such amino acid substitution patterns that were not present also in HTS-generated sequences were further eliminated as potential NUMTs. The overall purpose of utilising HTS reference mtCOI gene set was therefore to: (i) identify the mtDNA gene regions with highly-conserved amino acid signatures that have been retained across these cryptic hemipteran species complex, estimated to have diverged between 44.6 and 146 million years^32,33^, and (ii) to eliminate sequences representing potential ‘cryptic’ NUMTs where their amino acid substitution patterns have either violated permissible biochemical signatures, or that shared permissible biochemical signatures but violated the conserved regions amino acid signature rule. A schematic diagram outlining the analysis workflow is presented in Figure 1.

### Inclusion of additional GenBank sequences

Singleton partial mtCOI sequences representing a putative *B. tabaci* cryptic species (e.g., MEAM2, China2, AsiaII_2, AsiaII_4, SSA5, ‘Sudan1’) in the dataset of Boykin et al.^8^ were further compared to *Bemisia* cryptic species partial mtCOI sequences within GenBank (accessed 31-Jan-2019) using the NCBI blast search algorithm^53^ specifying the non-redundant (nr) sequence database. Sequences within GenBank that exhibited ≥98% sequence identity to these singleton partial mtCOI genes were downloaded to the CLC Sequence Viewer 8, followed by nucleotide and amino acid alignment to check for indels, premature stop codons, and amino acid conservation signatures as outlined above (Fig. 1). We further downloaded partial mtCOI sequences of related *Bemisia* species representing potential outgroups for phylogenetic analysis purposes. We assessed these outgroup sequences for presence of NUMTs and pseudogenes, and excluded sequences that could not be confirmed as free from NUMTs in all subsequent analyses.

### Phylogenetic analysis

Phylogenetic analysis using IQ-TREE web server <http://iqtree.cibiv.univie.ac.at>^54,55^ was carried out using the up-dated mtCOI database and included also partial mtCOI sequences extracted from published draft mitogenomes. We selected automatic model selection within ModelFinder^56^ and specified 2,000 bootstrap replications to assess for branch support, with maximum iterations for ultrafast bootstrap57 cut-off and a minimum correlation coefficient set at 1,000 and 0.99, respectively. IQ-TREE search parameter for perturbation strength and IQ-TREE stopping rule were as default values (i.e., 0.5 and 100, respectively). Visualisation of the phylogeny was by FigTree v1.4.4 (2006-2008, Andrew Rambaut, <http://tree.bio.ed.ac.uk>).

### Estimating uncorrected pairwise nucleotide distances (*p*-dist)

Studies to-date including analyses of global *B. tabaci* cryptic species complex partial mtCOI sequences have regularly used the Kimura-2-parameter (i.e., K2P) nucleotide substitution model to estimate species boundary^2,10,58^. The K2P model assumed equal base compositions (i.e., A=G=T=C)^59^ and violates the general mitochondrial DNA nucleotide compositions in Arthropods which is typically A-T rich (e.g., Lepidoptera^16,60,61^; Coleoptera^62,63^; *Bemisia* spp. e.g., Kunz et al.^22^; Tay et al.^23^). Instead, we used the uncorrected pairwise nucleotide distance (*p*-dist) for both within and between species comparison as recommended^64^. We used the multiple sequence alignment program MAFFT^65,66^ available in Geneious Prime and Geneious v11.1.5 with default settings (i.e., auto select for an appropriate strategy according to data size, scoring matrix set at 200PAM / K=2; and gap opening penalty and offset value set at 1.53 and 0.123, respectively). Estimated uncorrected *p*-dist were compared according to output phylogeny that identified specific species within major phylogenetic branches and between clades. All *p*-dist values of analysed individuals (excluding NUMTs/pseudogene-affected and misidentified species) were grouped in 0.01 (i.e., 1%) nucleotide distance increments to identify ‘genetic gap’ signatures^2,10^ for species boundary.

### Identification of PCR artefacts in SSA5

The sub-Saharan African 5 (SSA5; previously unpublished, renamed as SSA8^3^) partial mtCOI sequence (AM040598) was aligned against the up-dated CSIRO dataset and specifically against the SSA1, SSA2 and SSA3 sequences to ascertain nucleotide substitution patterns.

## Supporting information

Supplemental Tabel 1

## Availability of supporting data and materials

All sequences analysed are available from the NCBI. The clean mtCOI dataset and supporting data described in this paper is available as Supplementary Data1.

## Authors’ contributions

WTT, KHJG, SE, PJDB conceived the idea. WTT and DK coordinated the project and analysed the data. DK carried out laboratory experiments to generate HTS data. DK and WTT drafted the manuscript with input from all co-authors. PJDB provided the SSA1 and SSA2 samples. All authors have read and approved the manuscript.

## Acknowledgements

TK Walsh (CSIRO), S Vyskočilová (Natural Resources Institute, Greenwich University), and J Arnemann (Universidade Federal de Santa Maria, Brazil) provided helpful discussion.

---

**Suppl. Fig. 1:**
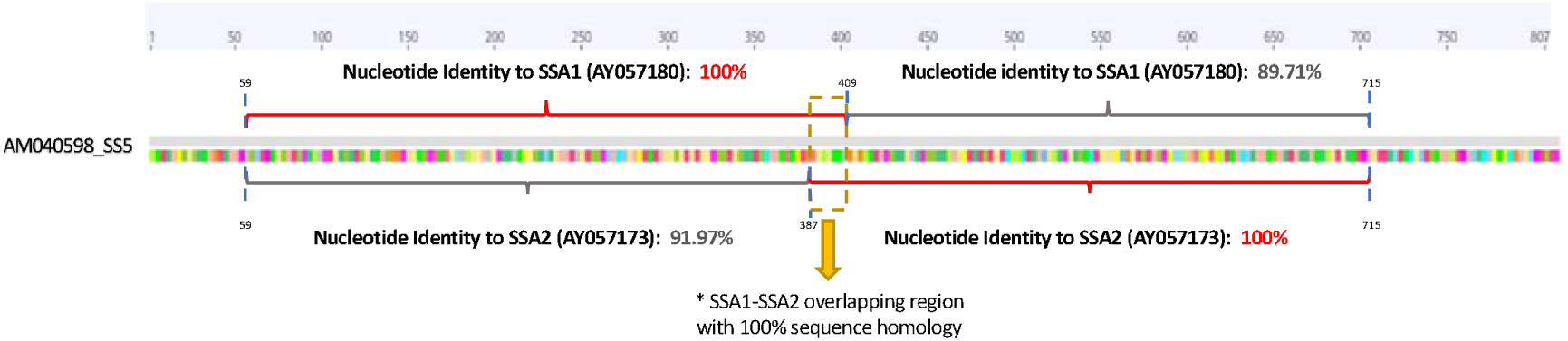
Chimeric amplicon signatures of SSA5 (renamed ‘SSA8’^3^). The shared nucleotide homology region (100%) between SSA1 and SSA2 (boxed section; nucleotide position 387-409) enabled the partial mtCOI sequences from these two different SSA species to be joined into a single contig that resulted in the misidentification of PCR chimeric products as a novel SSA species (i.e., SSA5). Note SSA1 (AY057180) and SSA2 (AY057173) are as entries in the current mtCOI dataset which have been assessed for NUMTs and trimmed to 657bp. Nucleotide positions are as SSA5 sequence (AM040598).

## References

1. De Barro, P. J., Liu, S. S., Boykin, L. M. & Dinsdale, A. B. Bemisia tabaci: a statement of species status. Annu Rev Entomol. 56, 1–19. doi:10.1146/annurev-ento-112408-085504 (2011).

2. Lee, W., Park, J., Lee, G. S., Lee, S. & Akimoto, S. Taxonomic status of the Bemisia tabaci complex (Hemiptera: Aleyrodidae) and reassessment of the number of its constituent species. PLoS ONE. 8, e63817. doi:10.1371/journal.pone.0063817 (2013).

3. Mugerwa, H., et al. African ancestry of New World, Bemisia tabaci-whitefly species. Sci Rep. 8, 2734. doi:10.1038/s41598-018-20956-3 (2018).

4. Macfadyen, S., et al. Cassava whitefly, Bemisia tabaci (Gennadius) (Hemiptera: Aleyrodidae) in East African farming landscapes: a review of the factors determining abundance. Bull Entomol Res. 108, 565–82. doi:10.1017/S0007485318000032 (2018).

5. Naranjo, S. E., Chu, C. C. & Henneberry, T. J. Economic injury levels for Bemisia tabaci (Homoptera: Aleyrodidae) in cotton: impact of crop price, control costs, and efficacy of control. Crop Protection 15, 779–788 (1996).

6. Otim-Nape, G. W. & Thresh, J. M. The current pandemic of cassava mosaic virus disease in Uganda. In The Epidemiology of Plant Diseases. Jones, D. G. (Ed.). 460 pp (1997).

7. Boykin, L. & De Barro, P. Global Bemisia dataset version 31 Dec 2012. CSIRO. (2012).

8. Boykin, L. M., Savill, A. & De Barro, P. Updated mtCOI reference dataset for the Bemisia tabaci species complex F1000Res. 6, 1835. doi:10.12688/f1000research.12858.1 (2017).

9. Vyskočilová, S., Tay, W. T., van Brunschot, S., Seal, S. & Colvin, J. An integrative approach to discovering cryptic species within the Bemisia tabaci whitefly species complex. Sci Rep. 8, 10886. doi:10.1038/s41598-018-29305-w (2018).

10. Dinsdale, A., Cook, L., Riginos, C., Buckley, Y. M. & De Barro, P. Refined Global Analysis of Bemisia tabaci (Hemiptera: Sternorrhyncha: Aleyrodoidea: Aleyrodidae) Mitochondrial Cytochrome Oxidase 1 to Identify Species Level Genetic Boundaries. Ann Entomol Soc Am. 103, 196–208. doi:10.1603/An09061 (2010).

11. Tay, W. T., Elfekih, S., Court, L. N., Gordon, K. H. J., Delatte, H. & De Barro, P. J. The Trouble with MEAM2: Implications of Pseudogenes on Species Delimitation in the Globally Invasive Bemisia tabaci (Hemiptera: Aleyrodidae) Cryptic Species Complex. Genome Biol E vol. 9, 2732–8. doi:10.1093/gbe/evx173 (2017).

12. Berry, S. D., et al. Molecular evidence for five distinct Bemisia tabaci (Homoptera : Aleyrodidae) geographic haplotypes associated with cassava plants in sub-Saharan Africa. Ann Entomol Soc Am. 97, 852–9. doi:10.1603/0013-8746(2004)097[0852:Meffdb]2.0.Co;2 (2004).

13. Wosula, E. N., Chen, W. B., Fei, Z. J. & Legg, J. P. Unravelling the Genetic Diversity among Cassava Bemisia tabaci Whiteflies Using NextRAD Sequencing. Genome Biol E vol. 9, 2958–73. doi:10.1093/gbe/evx219 (2017).

14. Elfekih, S., et al. On species delimitation, hybridization and population structure of cassava whitefly in Africa. Mol Ecol. Submitted (2019).

15. Griffiths, C. S. Correlation of functional domains and rates of nucleotide substitution in cytochrome b. Mol Phylogenet Evol. 7, 352–65. doi:10.1006/mpev.1997.0404 (1997).

16. Walsh. T. K., et al. Mitochondrial DNA genomes of five major Helicoverpa pest species from the Old and New Worlds (Lepidoptera: Noctuidae). Ecol E vol. 9, 2933–44. doi:10.1002/ece3.4971 (2019).

17. Buhay, J. E. “COI-Like” Sequences Are Becoming Problematic in Molecular Systematic and DNA Barcoding Studies. J Crustacean Biol. 29, 96–110. doi:10.1651/08-3020.1 (2009).

18. Hebert, P. D. N., Cywinska, A., Ball, S. L. & DeWaard, J. R. Biological identifications through DNA barcodes. P Roy Soc B-Biol Sci. 270(1512), 313–321. doi:10.1098/rspb.2002.2218 (2003).

19. Collins, T. M., Wimberger, P. H. & Naylor, G. J. P. Compositional bias, character-state bias, and charaterstate reconstruction using parsimony. Systematic Biol 43, 482–96 (1994).

20. Griffiths, B. S., et al. Community DNA hybridisation and %G+C profiles of microbial communities from heavy metal polluted soils. Fems Microbiol Ecol. 24, 103–12. DOI:10.1111/j.1574-6941.1997.tb00427.x (1997).

21. De Marchi, B. R., Kinene, T., Wainaina, J. M., Krause-Sakate, R. & Boykin, L. Comparative transcriptome analysis reveals genetic diversity in the endosymbiont Hamiltonella between native and exotic populations of Bemisia tabaci from Brazil. PLoS ONE 13, 7 doi:ARTN e020141110.1371/journal.pone.0201411 (2018).

22. Kunz, D., et al. Draft mitochondrial DNA genome of a 1920 Barbados cryptic Bemisia tabaci ‘New World’ species (Hemiptera: Aleyrodidae). Mitochondrial DNA B. 4, 1183–4. doi:10.1080/23802359.2019.1591197 (2019).

23. Tay, W. T., Elfekih, S., Court, L., Gordon, K. H. & De Barro, P. J. Complete mitochondrial DNA genome of Bemisia tabaci cryptic pest species complex Asia I (Hemiptera: Aleyrodidae). Mitochondrial DNA A. 27, 972–3. doi:10.3109/19401736.2014.926511 (2016).

24. Tay, W. T., et al. Novel molecular approach to define pest species status and tritrophic interactions from historical Bemisia specimens. Sci Rep 7, 439 doi:ARTN 42910.1038/s41598-017-00528-7 (2017).

25. Tay, W. T. & Gordon, K. H. J. Going global – genomic insights into insect invasions. Curr Opin Insect Sci. 31, 123–130. doi:10.1016/j.cois.2018.12.002 (2019).

26. Chen, W., et al. Genome of the African cassava whitefly Bemisia tabaci and distribution and genetic diversity of cassava-colonizing whiteflies in Africa. Insect Biochem Mol Biol. 110, 112–120. doi:10.1016/j.ibmb.2019.05.003 (2019).

27. Boykin, L. M., et al. Review and guide to a future naming system of African Bemisia tabaci species. Syst Entomol. 43, 427–33. doi:10.1111/syen.12294 (2018).

28. Tay, W. T., Evans, G. A., Boykin, L. M. & De Barro, P. J. Will the Real Bemisia tabaci Please Stand Up? PLoS ONE. 7, e50550. doi:ARTN e5055010.1371/journal.pone.0050550 (2012).

29. Thao, M. L., & Baumann, P. Evolutionary relationships of primary prokaryotic endosymbionts of whiteflies and their hosts. Appl Environ Microbiol. 70, 3401–6. doi:10.1128/AEM.70.6.3401-3406.2004 (2004).

30. Wang, H. L., et al. The characteristics and expression profiles of the mitochondrial genome for the Mediterranean species of the Bemisia tabaci complex. BMC Genomics. 14, 401. doi:10.1186/1471-2164-14-401 (2013).

31. Wang, H. L., Xiao, N., Yang, J., Wang, X. W., Colvin, J. & Liu, S. S. The complete mitochondrial genome of Bemisia afer (Hemiptera: Aleyrodidae). Mitochondiral DNA: DNA Mapping, Sequencing, and Analysis. 27, 98–99 (2013).

32. Boykin, L. M., Bell, C. D., Evans, G., Small, I. & De Barro, P. J. Is agriculture driving the diversification of the Bemisia tabaci species complex (Hemiptera: Sternorrhyncha: Aleyrodidae)?: Dating, diversification and biogeographic evidence revealed. BMC Evol Biol. 13, 228. doi:10.1186/1471-2148-13-228 (2013).

33. Xie, W., et al. The invasive MED/Q Bemisia tabaci genome: a tale of gene loss and gene gain. BMC Genomics. 19, 68. doi:10.1186/s12864-018-4448-9 (2018).

34. Alemandri, V., et al. Species within the Bemisia tabaci (Hemiptera: Aleyrodidae) complex in soybean and bean crops in Argentina. J Econ Entomol. 105, 48–53. doi:10.1603/ec11161 (2012).

35. Delatte H, et al. A new silverleaf-inducing biotype Ms of Bemisia tabaci (Hemiptera : Aleyrodidae) indigenous to the islands of the south-west Indian Ocean. Bull Entomol Res. 95, 29–35. doi:10.1079/Ber2004337 (2005).

36. Hu, J., et al. An extensive field survey combined with a phylogenetic analysis reveals rapid and widespread invasion of two alien whiteflies in China. PLoS ONE. 6:e16061. doi:10.1371/journal.pone.0016061 (2011).

37. Marubayashi, J. M., Rocha, K. C. G, Yuki, V. A. & Mituti, T. At least two indigenous species of the Bemisia tabaci complex are present in Brazil. J Appl Entomol. 137, 113–121 (2013).

38. De Barro PJ. The Bemisia tabaci Species Complex: Questions to Guide Future Research. J Integr Agr. 11, 187–196. doi:10.1016/S2095-3119(12)60003-3. (2012).

39. Ueda, S., Kitamura, T., Kijima, K., Honda, K. I. & Kanmiya, K. Distribution and molecular characterization of distinct Asian populations of Bemisia tabaci (Hemiptera: Aleyrodidae) in Japan. J Appl Entomol. 133, 355–366. doi:10.1111/j.1439-0418.2008.01379.x. (2009).

40. Karut, K., Kaydan, M. B., Tok, B., Doker, I. & Kazak, C. A new record for Bemisia tabaci (Gennadius) (Hemiptera: Aleyrodidae) species complex of Turkey. J Appl Entomol. 39, 158–60. doi:10.1111/jen.12169. (2015).

41. Liu, S. S., Colvin, J. & De Barro, P. J. Species Concepts as Applied to the Whitefly Bemisia tabaci Systematics: How Many Species Are There? J Integr Agr. 11, 176–86. doi:10.1016/S2095-3119(12)60002-1. (2012).

42. Vyskočilová, S., Seal, S. & Colvin, J. Relative polyphagy of “Mediterranean” cryptic Bemisia tabaci whitefly species and global pest status implications. J Pest Sci. 92, 1071–1088. doi:10.1007/s10340-019-01113-9. (2019).

43. Wang, P., Crowder, D. W. & Liu, S. S. Roles of mating behavioural interactions and life history traits in the competition between alien and indigenous whiteflies. Bull Entomol Res. 102, 395–405. doi:10.1017/S000748531100071x. (2012).

44. Bensasson, D., Zhang, D., Hartl, D. L. & Hewitt, G. M. Mitochondrial pseudogenes: evolution’s misplaced witnesses. Trends Ecol E vol. 16, 314–21 (2001).

45. Echols, N., et al. Comprehensive analysis of amino acid and nucleotide composition in eukaryotic genomes, comparing genes and pseudogenes. Nucleic Acids Res. 30, 2515–23. doi:10.1093/nar/30.11.2515 (2002).

46. Hazkani-Covo. E., Zeller. R. M. & Martin, W. Molecular poltergeists: mitochondrial DNA copies (numts) in sequenced nuclear genomes. PLoS Genet. 6, e1000834. doi:10.1371/journal.pgen.1000834 (2010).

47. Song, H., Buhay, J. E., Whiting, M. F. & Crandall, K. A. Many species in one: DNA barcoding overestimates the number of species when nuclear mitochondrial pseudogenes are coamplified. Proc Natl Acad Sci U S A. 105, 13486–13491. doi:10.1073/pnas.0803076105 (2008).

48. Roopa, H. K., et al. Prevalence of a new genetic group, MEAM-K, of the whitefly Bemisia tabaci (Hemiptera: Aleyrodidae) in Karnataka, India, as evident from mtCOI sequences. Fla Entomol. 98, 1062–1071 (2015).

49. Viscarret, M. M., et al. Mitochondrial DNA evidence for a distinct new world group of Bemisia tabaci (Gennadius) (Hemiptera : Aleyrodidae) indigenous to Argentina and Bolivia, and presence of the Old World B biotype in Argentina. Ann Entomol Soc Am. 96, 65–72. doi: 10.1603/0013-8746(2003)096[0065:Mdefad]2.0.Co;2. (2003).

50. Elfekih, S., Tay, W. T., Gordon, K., Court, L. N. & De Barro, P. J. Standardized molecular diagnostic tool for the identification of cryptic species within the Bemisia tabaci complex. Pest Manag Sci. 74, 170–3. doi:10.1002/ps.4676 (2018).

51. Simon, C., et al. Evolution, Weighting, and Phylogenetic Utility of Mitochondrial Gene-Sequences and a Compilation of Conserved Polymerase Chain-Reaction Primers. Ann Entomol Soc Am. 87, 651–701. DOI: 10.1093/aesa/87.6.651 (1994).

52. Shatters, R. G., Jr., Powell, C. A., Boykin, L. M., He, L. & McKenzie, C. L. Improved DNA barcoding method for Bemisia tabaci and related Aleyrodidae: development of universal and Bemisia tabaci biotype-specific mitochondrial cytochrome c oxidase I polymerase chain reaction primers. J Econ Entomol. 102, 750–8. doi:10.1603/029.102.0236. (2009).

53. Altschul, S. F., Gish, W., Miller, W., Myers, E. W. & Lipman, D. J. Basic local alignment search tool. J Mol Biol. 215, 403–10. doi:10.1016/S0022-2836(05)80360-2. (1990).

54. Nguyen, L.T., Schmidt, H. A., von Haeseler, A. & Minh, B. Q. IQ-TREE: a fast and effective stochastic algorithm for estimating maximum-likelihood phylogenies. Mol Biol E vol. 32, 268–74. doi:10.1093/molbev/msu300. (2015).

55. Trifinopoulos, J., Nguyen, L. T., von Haeseler, A. & Minh, B. Q. W-IQ-TREE: a fast online phylogenetic tool for maximum likelihood analysis. Nucleic Acids Res. 44 W1, W232–5. doi:10.1093/nar/gkw256. (2016).

56. Kalyaanamoorthy, S., Minh, B. Q., Wong, T. K. F., von Haeseler, A. & Jermiin, L. S. ModelFinder: fast model selection for accurate phylogenetic estimates. Nat Methods. 14, 587–9. doi:10.1038/nmeth.4285. (2017).

57. Hoang, D. T., Chernomor, O., von Haeseler, A., Minh, B. Q. & Vinh, L. S. UFBoot2: Improving the Ultrafast Bootstrap Approximation. Mol Biol E vol. 35, 518–22. doi:10.1093/molbev/msx281. (2018).

58. De Barro P. & Ahmed, M. Z. Genetic networking of the Bemisia tabaci cryptic species complex reveals pattern of biological invasions. PLoS ONE. 6, e25579. doi:10.1371/journal.pone.0025579. (2011).

59. Kimura, M. A simple method for estimating evolutionary rates of base substitutions through comparative studies of nucleotide sequences. J Mol Evol. 16, 111–120. (1980).

60. de Souza, B. R., Tay, W. T., Czepak, C., Elfekih, S. & Walsh, T. K. The complete mitochondrial DNA genome of a Chloridea (Heliothis) subflexa (Lepidoptera: Noctuidae) morpho-species. Mitochondrial DNA A: DNA Mapp Seq Anal. 27, 4532–3. doi:10.3109/19401736.2015.1101549. (2016).

61. Piper, M. C., van Helden, M., Court, L. N. & Tay, W. T. Complete mitochondrial genome of the European Grapevine moth (EGVM) Lobesia botrana (Lepidoptera: Tortricidae). Mitochondrial DNA A: DNA Mapp Seq Anal. 27, 3759–3760. doi:10.3109/19401736.2015.1079893. (2016).

62. Behere, G. T., Firake, D. M., Tay, W. T., Azad Thakur, N. S. & Ngachan, S. V. Complete mitochondrial genome sequence of a phytophagous ladybird beetle, Henosepilachna pusillanima (Mulsant) (Coleoptera: Coccinellidae). Mitochondrial DNA A: DNA Mapp Seq Anal. 27, 291–2. doi:10.3109/19401736.2014.892082. (2016).

63. Behere, G. T., et al. Characterization of draft mitochondrial genome of guava trunk borer, Aristobia reticulator (Fabricius, 1781) (Coleoptera: Cerambycidae: Lamiinae) from India. Mitochondrial DNA Part B. 4, 1592–3. DOI: 10.1080/23802359.2019.1602004. (2019).

64. Srivathsan, A. & Meier, R. On the inappropriate use of Kimura-2-parameter (K2P) divergences in the DNA-barcoding literature. Cladistics. 28, 190–194. doi:10.1111/j.1096-0031.2011.00370.x. (2012).

65. Katoh, K., Misawa, K., Kuma, K. & Miyata, T. MAFFT: a novel method for rapid multiple sequence alignment based on fast Fourier transform. Nucleic Acids Res. 30, 3059–3066. doi:10.1093/nar/gkf436. (2002).

66. Katoh, K. & Standley, D. M. MAFFT multiple sequence alignment software version 7: improvements in performance and usability. Mol Biol E vol. 30, 772–780. doi:10.1093/molbev/mst010. (2013).

