## Supplemental Data 1 for "Take out the rubbish – Removing NUMTs and pseudogenes from the *Bemisia tabaci* cryptic species mtCOI database"

>Japan2_Japan_Gunma_AB308111_AB308111.1_Bemisia_tabaci_mitochondrial_COI_gene_for_cytochrome_oxidase_subunit_I,_partial_cds,_country:_Japan:Gunma,_Ota

GAAAACTTGAAGTTTTTGGTAGACTAGGAATAATTTATACTATGTTAACT

ATTGGTATTTTAGGTTTTATTGTTTGAGGTCATCACATATTTACTGTTGG

GATAGATGTTGATACTCGGGCGTATTTTACTTCAGCTACAATAATTATTG

CTGTTCCTACGGGGATTAAAATCTTTAGTTGACTTGCTACTTTAGGAGGT

ATAAAGTCTAATAAGCTTAGTCCTCTTGTTCTTTGATTTACAGGGTTTTT

ATTTTTATTTACTATGGGTGGTTTAACTGGAATCATTTTGGGTAATTCTT

CTGTTGATGTTTGTTTACATGATACTTATTTTGTTGTTGCTCATTTTCAC

TATGTTTTATCTATGGGAATTATTTTTGCTGTTATGGGTGGGATTATTTA

TTGGTTTCCCCTGGTTTTAGGTCTGAGTTTAAATAATTATAGTTTAGTGT

CTCAATTTTATATGATATTTATGGGAGTAAATTTAACATTTTTTCCTCAA

CACTTTCTTGGCTTAGGGGGAATACCTCGTCGATATTCAGATTATCCTGA

TTGTTATCTTTTGTGGAACAAAATTTCCTCTGCGGGAAGCATCTTAAGTA

TTATTTCTGTCATTTATTTTTTATTTATTATTCTTGAATCTTTATTACTT

CTTCGAT

>Japan2_Japan_Kumamoto_AB308116_AB308116.1_Bemisia_tabaci_mitochondrial_COI_gene_for_cytochrome_oxidase_subunit_I,_partial_cds,_country:_Japan:Kumamoto,_Koshi,_Suya

GAAAACTTGAAGTTTTTGGTAGACTAGGAATAATTTATGCTATGTTAACT

ATTGGTATTTTAGGTTTTATTGTTTGAGGTCATCACATATTTACTGTTGG

GATAGATGTTGATACTCGGGCGTATTTTACTTCAGCTACAATAATTATTG

CTGTTCCTACGGGGATTAAAATCTTTAGTTGACTTGCTACTTTAGGAGGT

ATAAAGTCTAATAAGCTTAGTCCTCTTGTTCTTTGATTTACAGGGTTTTT

ATTTTTATTTACTATGGGTGGTTTAACTGGAATCATTTTGGGTAATTCTT

CTGTTGATGTTTGTTTACATGATACTTATTTTGTTGTTGCTCATTTTCAC

TATGTTTTATCTATGGGAATTATTTTTGCTGTTATGGGTGGGATTATTTA

TTGGTTTCCCCTGGTTTTAGGTCTGAGTTTAAATAATTATAGTTTAGTGT

CTCAATTTTATATGATATTCATGGGAGTAAATTTAACATTTTTTCCTCAA

CACTTTCTTGGCTTAGGGGGAATACCTCGTCGATATTCAGATTATCCTGA

TTGTTATCTTTTGTGGAACAAAATTTCCTCTGCGGGAAGCATCTTAAGTA

TTATTTCTGTCATTTATTTTTTATTTATTATTCTTGAATCTTTATTACTT

CTTCGAT

>Japan2_Japan_Kagoshima_AB308119_AB308119.1_Bemisia_tabaci_mitochondrial_COI_gene_for_cytochrome_oxidase_subunit_I,_partial_cds,_country:_Japan:Kagoshima,_Kamikawa

GAAAACTTGAAGTTTTTGGTAGACTAGGAATAATTTATGCTATGTTAACT

ATTGGTATTTTAGGTTTTATTGTTTGAGGTCATCACATATTTACTGTTGG

GATAGATGTTGATACTCGGGCGTATTTTACTTCAGCTACAATAATTATTG

CTGTTCCTACGGGGATTAAAATCTTTAGTTGACTTGCTACTTTAGGAGGT

ATAAAGTCTAATAAGCTTAGTCCTCTTGTTCTTTGATTTACAGGGTTTTT

ATTTTTATTTACTATGGGTGGTTTAACTGGAATCATTTTGGGTAATTCTT

CTGTTGATGTTTGTTTACATGATACTTATTTTGTTGTTGCTCATTTTCAC

TATGTTTTATCTATGGGAATTATTTTTGCTGTTATGGGTGGGATTATTTA

TTGGTTTCCCCTGGTTTTAGGTCTGAGTTTAAATAATTATAGTTTAGTGT

CTCAATTTTATATGATATTTATGGGAGTAAATTTAACATTTTTTCCTCAA

CACTTTCTTGGCTTAGGGGGAATACCTCGTCGATATTCAGATTATCCTGA

TTGTTATCTTTTGTGGAACAAAATTTCCTCTGCGGGAAGCATCTTAAGTA

TTATTTCTGTCATTTACTTTTTATTTATTATTCTTGAATCTTTATTACTT

CTTCGAT

>Japan2_Japan_Kumamoto_AB308115_AB308115.1_Bemisia_tabaci_mitochondrial_COI_gene_for_cytochrome_oxidase_subunit_I,_partial_cds,_country:_Japan:Kumamoto,_Yamaga

GAAAACTTGAAGTTTTTGGTAGACTAGGAATAATTTATGCTATGTTAACT

ATTGGTATTTTAGGTTTTATTGTTTGAGGTCATCATATATTTACTGTTGG

AATAGATGTTGATACTCGGGCGTATTTTACTTCAGCTACAATAATTATTG

CTGTTCCTACGGGGATTAAAATCTTTAGTTGACTTGCTACTTTAGGAGGT

ATAAAGTCTAATAAGCTTAGTCCTCTTGTTCTTTGATTTACAGGGTTTTT

ATTTTTATTTACTATGGGTGGTTTAACTGGAATCATTTTGGGTAATTCTT

CTGTTGATGTTTGTTTACATGATACTTATTTTGTTGTTGCTCATTTTCAC

TATGTTTTATCTATGGGAATTATTTTTGCTGTTATGGGTGGGATTATTTA

TTGGTTTCCCCTGGTTTTAGGTCTGAGTTTAAATAATTATAGTTTAGTGT

CTCAATTTTATATGATATTCATGGGAGTAAATTTAACATTTTTTCCTCAA

CACTTTCTTGGCTTAGGGGGAATACCTCGTCGATATTCAGATTATCCTGA

TTGTTATCTTTTGTGGAACAAAATTTCCTCTGCGGGAAGCATCTTAAGTA

TTATTTCTGTCATTTATTTTTTATTTATTATTCTTGAATCTTTATTACTT

CTTCGAT

>Japan2_Japan_Oita_AB308117_AB308117.1_Bemisia_tabaci_mitochondrial_COI_gene_for_cytochrome_oxidase_subunit_I,_partial_cds,_country:_Japan:Oita,_Usa

GAAAACTTGAAGTTTTTGGTAGACTAGGAATAATTTATGCTATGTTAACT

ATTGGTATTTTAGGTTTTATTGTTTGAGGTCATCACATATTTACTGTTGG

GATAGATGTTGATACTCGGGCGTATTTTACTTCAGCTACAATAATTATTG

CTGTTCCTACGGGGATTAAAATCTTTAGTTGACTTGCTACTTTAGGAGGT

ATAAAGTCTAATAAGCTTAGTCCTCTTGTTCTTTGATTTACAGGGTTTTT

ATTTTTATTTACTATGGGTGGTTTAACTGGAATCATTTTGGGTAATTCTT

CTGTTGATGTTTGTTTACATGATACTTATTTTGTTGTTGCTCATTTTCAC

TATGTTTTATCTATGGGAATTATTTTTGCTGTTATAGGTGGGATTATTTA

TTGGTTTCCCCTGGTTTTAGGTCTGAGTTTAAATAATTATAGTTTAGTGT

CTCAATTTTATATGATATTCATGGGAGTAAATTTAACATTTTTTCCTCAA

CACTTTCTTGGCTTAGGGGGAATACCTCGTCGATATTCAGATTATCCTGA

TTGTTATCTTTTGTGGAACAAAATTTCCTCTGCGGGAAGCATCTTAAGTA

TTATTTCTGTCATTTATTTTTTATTTATTATTCTTGAATCTTTATTACTT

CTTCGAT

>Japan1_Japan_Kagoshima_AB440786_AB440786.1_Bemisia_tabaci_mitochondrial_COI_gene_for_cytochrome_oxidase_subunit_I,_partial_cds,_isolate:_Kikai-2007-Wb

GAAAGCTTGAGGTATTTGGGAGATTAGGTATGATTTATGCTATAATAACT

ATTGGTATTTTGGGATTTATTGTTTGAGGTCATCATATATTTACTGTTGG

GATAGATGTTGACACTCGGGCTTATTTCACTTCAGCTACTATGATTATTG

CTGTTCCAACAGGAATTAAAATTTTTAGGTGACTTGCTACTTTAGGTGGA

ATAAAGTCTAATAAGTTTAGACCGCTTGTACTTTGATTTTCAGGATTTTT

ATTTTTATTTACTATAGGTGGATTAACTGGGATTATTCTTGGTAATTCTT

CTGTTGATGTTTGCTTGCATGATACCTATTTTGTTGTTGCTCATTTTCAT

TATGTTTTATCTATAGGAATTATTTTTGCTATTGTGGGAGGTTTTATTTA

TTGGTTTCCACTGATTTTAGGTATAACACTAAATAATCATAACCTGGTAT

CTCAGTTTTATATTATATTTTTGGGGGTAAATTTAACATTTTTTCCGCAA

CATTTTCTTGGGCTAGGGGGAATACCCCGCCGGTATTCAGACTATCCTGA

TTGTTATTTGATGTGAAATAAAATTTCTTCTGCAGGAAGCATCTTAAGTA

TTGTTTCTGTTATTTACTTTTTGTTTATTGTTTTAGAATCTTTACTTCTT

TTACGTT

>Asia1_Bangladesh_Chitagong_AJ748398_AJ748398.1 Bemisia tabaci mitochondrial partial coi gene for cytochrome oxidase I subunit, specific host okra, from Bangladesh, Chitagong

GGAAACTTGAGGTATTTGGCAGGTTAGGAATAATTTATGCTATAATAACTATTGGCATTTTGGGGTTTAT

TGTTTGAGGTCATCATATATTTACTGTTGGTATAGATGTTGATACTCGAGCTTATTTTACTTCAGCTACT

ATGGTTATTGCTGTTCCAACTGGGATTAAGATTTTCAGGTGGCTTGCTACTTTAGGTGGAATAAAATCCA

ATAAATTAAGGCCGTTAGTTCTTTGATTTACAGGATTTTTATTCTTATTTACCATGGGTGGACTAACTGG

GATTATTCTTGGTAATTCTTCTGTTGATGTTTGTTTGCATGATACTTATTTTGTTGTTGCTCATTTTCAT

TATGTTTTATCCATAGGAATCATTTTTGCTATCATAGGAGGTTTTATTTACTGATTTCCATTAATCTTAG

GTCTAACATTAAATAACCATAATTTGGTATCTCAGTTTTATATTATATTTTTGGGCGTTAACCTAACATT

TTTTCCACAACACTTTCTTGGATTAAGTGGAATACCTCGTCGGTATTCAGATTATCCTGATTGTTATCTC

ATATGAAATAAAATTTCTTCTGCGGGGAGTATCTTGAGAATTATTTCTGTTATCTATTTTTTATTTATTG

TTTTAGAATCTTTGCTTCTTTTACGGC

>Asia1_India_Indore_JN855572_JN855572.1 Bemisia tabaci isolate Indore cytochrome oxidase subunit 1 (co1) gene, partial cds; mitochondrial

GGAAACTTGAGGTATTTGGCAGGTTAGGAATAATTTATGCTATAATAACTATTGGCATTTTGGGGTTTAT

TGTTTGAGGTCATCATATATTTACTGTTGGTATAGATGTTGATACTCGAGCTTATTTTACTTCAGCTACT

ATGGTTATTGCTGTTCCAACTGGGATTAAGATTTTCAGGTGGCTTGCTACTTTAGGTGGAATAAAATCCA

ATAAATTAAGGCCGTTAGTTCTTTGATTTACAGGATTTTTATTCTTATTTACCATGGGTGGACTAACTGG

GATTATTCTTGGTAATTCTTCTGTTGATGTTTGTTTGCATGATACTTATTTTGTTGTTGCTCATTTTCAT

TATGTTTTATCCATAGGAATCATTTTTGCTATCATAGGAGGTTTTATTTACTGATTTCCATTAATCTTAG

GTCTAACATTAAATAACCATAATTTGGTATCTCAGTTTTATATTATATTTTTGGGCGTTAACCTAACATT

TTTTCCACAACACTTTCTTGGATTAAGCGGAATACCTCGTCGGTATTCAGATTATCCTGATTGTTATCTC

ATATGAAATAAAATTTCTTCTGCGGGGAGTATCTTGAGAATTATTTCTGTTATCTATTTTTTATTTATTG

TTTTAGAATCTTTGCTTCTTTTACGGC

>Asia1_China_Hainan_HM137323_HM137323.1 Bemisia tabaci isolate h11 cytochrome oxidase subunit 1 gene, partial cds; mitochondrial

GGAAACTTGAGGTATTTGGCAGGTTAGGAATAATTTATGCTATAATAACTATTGGCATTTTGGGGTTTAT

TGTTTGAGGTCATCATATATTTACTGTTGGTATAGATGTTGATACTCGAGCTTATTTTACTTCAGCTACT

ATGGTTATTGCTGTTCCAACTGGGATTAAGATTTTCAGGTGGCTTGCTACTTTAGGTGGAATAAAATCCA

ATAAATTAAGGCCGTTAGTTCTTTGATTTACAGGATTTTTATTCTTATTTACCATGGGTGGACTAACTGG

GATTATTCTTGGTAATTCTTCTGTTGATGTTTGTTTGCATGATACTTATTTTGTTGTTGCTCATTTTCAT

TATGTTTTATCCATAGGAATCATTTTTGCTATCATAGGAGGTTTTATTTACTGATTTCCATTAATCTTAG

GTCTAACATTAAATAACCATAATTTGGTATCTCAGTTTTATATTATATTTTTGGGCGTTAACCTAACATT

TTTTCCACAACACTTTCTTGGATTAAGCGGAATACCTCGTCGGTATTCAGATTATCCTGATTGTTATCTC

ATATGAAATAAAATTTCTTCTGCGGGGAGTATCTTGAGAATTATTTCTGTTATTTATTTTTTATTTATTG

TTTTAGAATCTTTGCTTCTTTTACGGC

>Asia1_India_AJ748359_AJ748359.1 Bemisia tabaci mitochondrial partial coi gene for cytochrome oxidase I subunit, specific host brinjal, from India, Karnataka, Chitradurga, Hiriyur

GGAAACTTGAGGTATTTGGCAGGTTAGGAATAATTTATGCTATAATAACTATTGGCATTTTGGGGTTTAT

TGTTTGAGGTCATCATATATTTACTGTTGGTATAGATGTTGATACTCGAGCTTATTTTACTTCAGCTACT

ATGGTTATTGCTGTTCCAACTGGGATTAAGATTTTCAGGTGGCTTGCTACTTTAGGTGGAATAAAATCCA

ATAAATTAAGGCCGTTAGTTCTTTGATTTACAGGATTTTTATTCTTATTTACCATGGGTGGACTAACTGG

GATTATTCTTGGTAATTCTTCTGTTGATGTTTGTTTGCATGATACTTATTTTGTTGTTGCTCATTTTCAT

TATGTTTTATCCATAGGAATCATTTTTGCTATCATGGGAGGTTTTATTTACTGATTTCCATTAATCTTAG

GTCTAACATTAAATAACCATAATTTGGTATCTCAGTTTTATATTATATTTTTGGGCGTTAACCTAACATT

TTTTCCACAACACTTTCTTGGATTAAGCGGAATACCTCGTCGGTATTCAGATTATCCTGATTGTTATCTC

ATATGAAATAAAATTTCTTCTGCGGGGAGTATCTTGAGAATTATTTCTGTTATCTATTTTTTATTTATTG

TTTTAGAATCTTTGCTTCTTTTACGGC

>Asia1_India_Karnataka_AJ748361_ AJ748361.1_ Bemisia tabaci mitochondrial partial coi gene for cytochrome oxidase I subunit, specific host brinjal, from India, Karnataka, Ranibennur, Sulibele

GGAAACTTGAGGTATTTGGCAGGTTAGGAATAATTTATGCTATAATAACTATTGGCATTTTGGGGTTTAT

TGTTTGAGGTCATCATATATTTACTGTTGGTATAGATGTTGATACTCGAGCTTATTTTACTTCAGCTACT

ATGGTTATTGCTGTTCCAACTGGGATTAAGATTTTCAGGTGGCTTGCTACTTTAGGTGGAATAAAATCCA

ATAAATTAAGGCCGTTAGTTCTTTGATTTACAGGATTTTTATTCTTATTTACCATGGGTGGACTAACTGG

GATTATTCTTGGTAATTCTTCTGTTGATGTTTGTTTGCATGATACTTATTTTGTTGTTGCTCATTTTCAT

TATGTTTTATCCATAGGAATCATTTTTGCTATCATAGGAGGTTTTATTTACTGATTTCCATTAATCTTAG

GTCTAACATTAAATAACCATAATTTGGTATCTCAGTTTTATATTATATTTTTGGGCGTTAACCTAACATT

TTTTCCACAACACTTTCTTGGATTAAGCGGAATACCTCGTCGGTATTCAGATTATCCTGATTGTTATCTC

ATATGAAATAAAATTTCTTCTGCGGGGAGTATCTTGAGAATTATTTCTGTTATCTATTTTTTATTTATTG

TTTTAGAATCTTTGCTTCTTTTACGGC

>Asia1_India_AJ748371_AJ748371.1 Bemisia tabaci mitochondrial partial coi gene for cytochrome oxidase I subunit, specific host Parthenium hysterophorus, from India, Karnataka, Chikmagalur, Mudigere, Kushalanagar

GGAAACTTGAGGTATTTGGCAGGTTAGGAATAATTTATGCTATAATAACTATTGGCATTTTGGGGTTTAT

TGTTTGAGGTCATCATATATTTACTGTTGGTATAGATGTTGATACTCGAGCTTATTTTACTTCAGCTACT

ATGGTTATTGCTGTTCCAACTGGGATTAAGATTTTCAGGTGGCTTGCTACTTTAGGTGGAATAAAATCCA

ATAAATTAAGGCCGTTAGTTCTTTGATTTACAGGATTTTTATTCTTATTTACCATGGGTGGACTGACTGG

GATTATTCTTGGTAATTCTTCTGTTGATGTTTGTTTGCATGATACTTATTTTGTTGTTGCTCATTTTCAT

TATGTTTTATCCATAGGAATCATTTTTGCTATCATAGGAGGTTTTATTTACTGATTTCCATTAATCTTAG

GTCTAACATTAAATAACCATAATTTGGTATCTCAGTTTTATATTATATTTTTGGGCGTTAACCTAACATT

TTTTCCACAACACTTTCTTGGATTAAGCGGAATACCTCGTCGGTATTCAGATTATCCTGATTGTTATCTC

ATATGAAATAAAATTTCTTCTGCGGGGAGTATCTTGAGAATTATTTCTGTTATCTATTTTTTATTTATTG

TTTTAGAATCTTTGCTTCTTTTACGGC

>Asia1_India_Andre_Pradesh_HM590171_HM590171.1 Bemisia tabaci isolate PEDAKURUPADU-3 cytochrome oxidase subunit 1 gene, partial cds; mitochondrial

GGAAACTTGAGGTATTTGGCAGGTTAGGAATAATTTATGCTATAATAACTATTGGCATTTTGGGGTTTAT

TGTTTGAGGTCATCATATATTTACTGTTGGTATAGATGTTGATACTCGAGCTTATTTTACTTCAGCTACT

ATGGTTATTGCTGTTCCAACTGGGATTAAGATTTTCAGGTGGCTTGCTACTTTAGGTGGAATAAAATCCA

ATAAATTAAGGCCGTTAGTTCTTTGATTTACAGGATTTTTATTCTTATTTACCATGGGTGGACTAACTGG

GATTATTCTTGGTAATTCTTCTGTTGATGTTTGTTTGCATGATACTTATTTTGTTGTTGCTCATTTTCAT

TATGTTTTATCCATAGGAATCATTTTTGCCATCATAGGAGGTTTTATTTACTGATTTCCATTAATCTTAG

GTCTAACATTAAATAACCATAATTTGGTATCTCAGTTTTATATTATATTTTTGGGCGTTAACCTAACATT

TTTTCCACAACACTTTCTTGGATTAAGCGGAATACCTCGTCGGTATTCAGATTATCCTGATTGTTATCTC

ATATGAAATAAAATTTCTTCTGCGGGGAGTATCTTGAGAATTATTTCTGTTATCTATTTTTTATTTATTG

TTTTAGAATCTTTGCTTCTTTTACGGC

>Asia1_India_JN703454_JN703454.1 Bemisia tabaci voucher F2028 cytochrome oxidase subunit I gene, partial cds; mitochondrial

GGAAACTTGAGGTATTTGGCAGGTTAGGAATAATTTATGCTATAATAACTATTGGCATTTTGGGGTTTAT

TGTTTGAGGTCATCATATATTTACTGTTGGTATAGATGTTGATACTCGAGCTTATTTTACTTCAGCTACT

ATGGTTATTGCTGTTCCAACTGGGATTAAGATTTTCAGGTGGCTTGCTACTTTAGGTGGAATAAAATCCA

ATAAATTAAGGCCGTTAGTTCTTTGATTTACAGGATTTTTATTCTTATTTACCATGGGTGGACTAACTGG

GATTATTCTTGGTAATTCTTCTGTTGATGTTTGTTTGCATGATACTTATTTTGTTGTTGCTCATTTTCAT

TATGTTTTATCCATAGGAATCATTTTTGCTATCATAGGGGGTTTTATTTACTGATTTCCATTAATCTTAG

GTCTAACATTAAATAACCATAATTTGGTATCTCAGTTTTATATTATATTTTTGGGCGTTAACCTAACATT

TTTTCCACAACACTTTCTTGGATTAAGCGGAATACCTCGTCGGTATTCAGATTATCCTGATTGTTATCTC

ATATGAAATAAAATTTCTTCTGCGGGGAGTATCTTGAGAATTATTTCTGTTATCTATTTTTTATTTATTG

TTTTAGAATCTTTGCTTCTTTTACGGC

>Asia1_India_Maharastra_HM590159_HM590159.1 Bemisia tabaci isolate AKOLA-2 cytochrome oxidase subunit 1 gene, partial cds; mitochondrial

GGAAACTTGAGGTATTTGGCAGGTTAGGAATAATTTATGCTATAATAACTATTGGCATTTTGGGGTTTAT

TGTTTGAGGTCATCATATATTTACTGTTGGTATAGATGTTGATACTCGAGCTTATTTTACTTCAGCTACT

ATGGTTATTGCTGTTCCAACTGGGATTAAGATTTTCAGGTGGCTTGCTACTTTAGGTGGAATAAAATCCA

ATAAATTAAGGCCGTTAGTTCTTTGATTTACAGGATTTTTATTCTTATTTACCATGGGTGGACTAACTGG

GATTATTCTTGGTAATTCTTCTGTTGATGTTTGTTTGCATGATACTTATTTTGTTGTTGCTCATTTTCAT

TATGTTTTATCCATAGGAATCATTTTTGCTATCATAGGAGGTTTTATTTACTGATTTCCATTAATCTTAG

GTCTAACATTAAATAACCATAATTTGGTATCTCAGTTTTATATTATATTTTTGGGCGTTAACCTAACATT

TTTTCCACAACACTTTCTTGGATTAAGCGGAATACCTCGTCGGTATTCAGATTATCCTGATTGTTATCTC

ATATGAAATAAAATTTCTTCTGCAGGGAGTATCTTGAGAATTATTTCTGTTATCTATTTTTTATTTATTG

TTTTAGAATCTTTGCTTCTTTTACGGC

>Asia1_India_Tamilnadu_HM590180_HM590180.1 Bemisia tabaci isolate PARCHURU-1 cytochrome oxidase subunit 1 gene, partial cds; mitochondrial

GGAAACTTGAAGTATTTGGCAGGTTAGGAATAATTTATGCTATAATAACTATTGGCATTTTGGGGTTTAT

TGTTTGAGGTCATCATATATTTACTGTTGGTATAGATGTTGATACTCGAGCTTATTTTACTTCAGCTACT

ATGGTTATTGCTGTTCCAACTGGGATTAAGATTTTCAGGTGGCTTGCTACTTTAGGTGGAATAAAATCCA

ATAAATTAAGGCCGTTAGTTCTTTGATTTACAGGATTTTTATTCTTATTTACCATGGGTGGACTAACTGG

GATTATTCTTGGTAATTCTTCTGTTGATGTTTGTTTGCATGATACTTATTTTGTTGTTGCTCATTTTCAT

TATGTTTTATCCATAGGAATCATTTTTGCTATCATAGGAGGTTTTATTTACTGATTTCCATTAATCTTAG

GTCTAACATTAAATAACCATAATTTGGTATCTCAGTTTTATATTATATTTTTGGGCGTTAACCTAACATT

TTTTCCACAACACTTTCTTGGATTAAGCGGAATACCTCGTCGGTATTCAGATTATCCTGATTGTTATCTC

ATATGAAATAAAATTTCTTCTGCGGGGAGTATCTTGAGAATTATTTCTGTTATCTATTTTTTATTTATTG

TTTTAGAATCTTTGCTTCTTTTACGGC

>Asia1_India_Andre_Pradesh_HM590156_HM590156.1 Bemisia tabaci isolate KOLLETIGUNTA-3 cytochrome oxidase subunit 1 gene, partial cds; mitochondrial

GGAAACTTGAGGTATTTGGCAGGTTAGGAATAATTTATGCTATAATAACTATTGGCATTTTGGGGTTTAT

TGTTTGAGGTCATCATATATTTACTGTTGGTATAGATGTTGATACTCGAGCTTATTTTACTTCAGCTACT

ATGGTTATTGCTGTTCCAACTGGGATTAAGATTTTCAGGTGGCTTGCTACTTTAGGTGGAATAAAATCCA

ATAAATTAAGGCCGTTAGTTCTTTGATTTACAGGATTTTTATTCTTATTTACCATGGGTGGACTAACTGG

GATTATTCTTGGTAATTCTTCTGTTGATGTTTGTTTGCATGATACTTATTTTGTTGTCGCTCATTTTCAT

TATGTTTTATCCATAGGAATCATTTTTGCTATCATAGGAGGTTTTATTTACTGATTTCCATTAATCTTAG

GTCTAACATTAAATAACCATAATTTGGTATCTCAGTTTTATATTATATTTTTGGGTGTTAACCTAACATT

TTTTCCACAACACTTTCTTGGATTAAGCGGAATACCTCGTCGGTATTCAGATTATCCTGATTGTTATCTC

ATATGAAATAAAATTTCTTCTGCGGGGAGTATCTTGAGAATTATTTCTGTTATCTATTTTTTATTTATTG

TTTTAGAATCTTTGCTTCTTTTACGGC

>Asia1_India_HM590154_HM590154.2 Bemisia tabaci isolate NAGPUR-3 cytochrome oxidase subunit 1 gene, partial cds; mitochondrial

GGAAACTTGAGGTATTTGGCAGGTTAGGAATAATTTATGCTATAATAACTATTGGCATTTTGGGGTTTAT

TGTTTGAGGTCATCATATATTTACTGTTGGTATAGATGTTGATACTCGAGCTTATTTTACTTCAGCTACT

ATGGTTATTGCTGTTCCAACTGGGATTAAGATTTTCAGGTGGCTTGCTACTTTAGGTGGAATAAAATCCA

ATAAATTAAGGCCGTTAGTTCTTTGATTTACAGGATTTTTATTCTTATTTACCATGGGTGGACTAACTGG

GATTATTCTTGGTAATTCTTCTGTTGATGTTTGTTTGCATGATACTTATTTTGTTGTTGCTCATTTTCAT

TATGTTTTATCCATAGGAATCATTTTTGCTATCATAGGAGGTTTTATTTACTGATTTCCATTAATCTTAG

GTCTAACATTAAATAACCATAATTTGGTATCTCAGTTTTATATTATATTTTTGGGTGTTAACCTAACATT

TTTTCCACAACACTTTCTTGGATTAAGCGGAATACCTCGTCGGTATTCAGATTATCCTGATTGTTATCTC

ATATGAAATAAAATTTCTTCTGCGGGGAGTATCTTGAGAATTATTTCTGTTATCTATTTTTTATTTATTG

TTTTAGAATCTTTGCTTCTTTTACGGC

>Asia1_Thailand_AF164671_AF164671.1 Bemisia tabaci biotype Thailand Euphorbia cytochrome oxidase I gene, partial sequence, mitochondrial gene for mitochondrial protein

GGAAACTTGAGGTATTTGGCAGGTTAGGAATAATTTATGCTATAATAACTATTGGCATTTTGGGGTTTAT

TGTTTGAGGTCATCATATATTTACTGTTGGTATAGATGTTGATACTCGAGCTTATTTTACTTCAGCTACT

ATGGTTATTGCTGTTCCAACTGGGATTAAGATTTTCAGGTGGCTTGCTACTTTAGGTGGAATAAAATCCA

ATAAATTAAGGCCGTTAGTTCTTTGATTTACAGGATTTTTATTCTTATTTACCATGGGTGGACTAACTGG

GATTATTCTTGGTAATTCTTCTGTTGATGTTTGTTTGCATGATACTTATTTTGTTGTTGCTCATTTTCAT

TATGTTTTATCCATAGGAATCATTTTTGCTATCATAGGAGGTTTTATTTACTGATTTCCATTAATCTTAG

GTCTAACATTAAATAACCATAATTTGGTATCTCAGTTTTATATTATATTTTTGGGTGTTAACCTAACATT

TTTTCCACAACACTTTCTTGGGTTAAGCGGAATACCTCGTCGGTATTCAGATTATCCTGATTGTTATCTC

ATATGAAATAAAATTTCTTCTGCGGGGAGTATCTTGAGAATTATTTCTGTTATCTATTTTTTATTTATTG

TTTTAGAATCTTTGCTTCTTTTACGGC

>Asia1_India_AJ748369_AJ748369.1 Bemisia tabaci mitochondrial partial coi gene for cytochrome oxidase I subunit, specific host Parthenium hysterophorus, from India, Karnataka, Bangalore, Hebbal

GGAAACTTGAGGTATTTGGTAGGTTAGGAATAATTTATGCTATAATAACTATTGGCATTTTGGGGTTTAT

TGTTTGAGGTCATCATATATTTACTGTTGGTATAGATGTTGATACTCGAGCTTATTTTACTTCAGCTACT

ATGGTTATTGCTGTTCCAACTGGGATTAAGATTTTCAGGTGGCTTGCTACTTTAGGTGGAATAAAATCCA

ATAAATTAAGGCCGTTAGTTCTTTGATTTACAGGATTTTTATTCTTATTTACCATGGGTGGACTAACTGG

GATCATTCTTGGTAATTCTTCTGTTGATGTTTGTTTGCATGATACTTATTTTGTTGTTGCTCATTTTCAT

TATGTTTTATCCATAGGAATCATTTTTGCTATCATAGGAGGTTTTATTTACTGATTTCCATTAATCTTAG

GTCTAACATTAAATAACCATAATTTGGTATCTCAGTTTTATATTATATTTTTGGGCGTTAACCTAACATT

TTTTCCACAACACTTTCTTGGATTAAGCGGAATACCTCGTCGGTATTCAGATTATCCTGATTGTTATCTC

ATATGAAATAAAATTTCTTCTGCGGGGAGTATCTTGAGAATTATTTCTGTTATCTATTTTTTATTTATTG

TTTTAGAATCTTTGCTTCTTTTACGGC

>Asia1_Indonesia_AB248260_AB248260.1 Bemisia tabaci mitochondrial COI gene for cytochrome oxidase subunit I, partial cds, isolate: BtCb2BJB

GGAAACTTGAGGTATTTGGCAGGTTAGGAATAATTTATGCTATAATAACTATTGGCATCTTGGGGTTTAT

TGTTTGAGGTCATCATATATTTACTGTTGGTATAGATGTTGATACTCGAGCTTATTTTACTTCAGCTACT

ATGGTTATTGCTGTTCCAACTGGGATTAAGATTTTCAGGTGGCTTGCTACTTTAGGTGGAATAAAATCCA

ATAAATTAAGGCCGTTAGTTCTTTGATTTACAGGATTTTTATTCTTATTTACCATGGGTGGACTAACTGG

GATTATTCTTGGTAATTCTTCTGTTGATGTTTGTTTGCATGATACTTATTTTGTTGTTGCTCATTTTCAT

TATGTTTTATCCATAGGAATCATTTTTGCTATCATAGGAGGTTTTATTTACTGATTTCCATTAATCTTAG

GTCTAACATTAAATAACCATAATTTAGTATCTCAGTTTTATATTATATTTTTGGGCGTTAACCTAACATT

TTTTCCACAACACTTTCTTGGATTAAGCGGAATACCTCGTCGGTATTCAGATTATCCTGATTGTTATCTC

ATATGAAATAAAATTTCTTCTGCGGGGAGTATCTTGAGAATTATTTCTGTTATCTATTTTTTATTTATTG

TTTTAGAATCTTTGCTTCTTTTACGGC

>Asia1_Indonesia_HE653706_HE653706.1 Bemisia tabaci mitochondrial partial COI gene for cytochrome oxidase subunit 1, isolate Indo#25

GGAAACTTGAGGTATTCGGCAGGTTAGGAATAATTTATGCTATAATAACTATTGGCATCTTGGGGTTTAT

TGTTTGAGGTCATCATATATTTACTGTTGGTATAGATGTTGATACTCGAGCTTATTTTACTTCAGCTACT

ATGGTTATTGCTGTTCCAACTGGGATTAAGATTTTCAGGTGGCTTGCTACTTTAGGTGGAATAAAATCCA

ATAAATTAAGGCCGTTAGTTCTTTGATTTACAGGATTTCTATTCTTATTTACCATGGGTGGACTAACTGG

GATTATTCTTGGTAATTCTTCTGTTGATGTTTGTTTGCATGATACTTATTTTGTTGTTGCTCATTTTCAT

TATGTTTTATCCATAGGAATCATTTTTGCTATCATAGGAGGTTTTATTTACTGATTTCCATTAATCTTAG

GTCTAACATTAAATAACCATAATTTGGTATCTCAGTTTTATATTATATTTTTGGGCGTTAACCTAACATT

TTTTCCACAACACTTTCTTGGATTAAGCGGAATACCTCGTCGGTATTCAGATTATCCTGATTGTTATCTC

ATATGAAATAAAATTTCTTCTGCGGGGAGTATCTTGAGAATTATTTCTGTTATCTATTTTTTATTTATTG

TTTTAGAATCTTTGCTTCTTTTACGGC

>Asia1_Indonesia_HE653710_HE653710.1 Bemisia tabaci mitochondrial partial COI gene for cytochrome oxidase subunit 1, isolate Indo#29

GGAAACTTGAGGTATTCGGCAGGTTAGGAATAATTTATGCTATAATAACTATTGGCATCTTGGGGTTTAT

TGTTTGAGGTCATCATATATTTACTGTTGGTATAGATGTTGATACTCGAGCTTATTTTACTTCAGCTACT

ATGGTTATTGCTGTTCCAACTGGGATTAAGATTTTCAGGTGGCTTGCTACTTTAGGTGGAATAAAATCCA

ATAAATTAAGGCCGTTAGTTCTTTGATTTACAGGATTTTTATTCTTATTTACCATGGGTGGACTAACTGG

GATTATTCTTGGTAATTCTTCTGTTGATGTTTGTTTGCATGATACTTATTTTGTTGTTGCTCATTTTCAT

TATGTTTTATCCATAGGAATCATTTTTGCTATCATAGGAGGTTTTATTTACTGATTTCCATTAATCTTAG

GTCTAACATTAAATAACCATAATTTGGTATCTCAGTTTTATATTATATTTTTGGGCGTTAACCTAACATT

TTTTCCACAACACTTTCTTGGATTAAGCGGAATACCTCGTCGGTATTCAGATTATCCTGATTGTTATCTC

ATATGAAATAAAATTTCTTCTGCGGGGAGTATCTTGAGAATTATTTCTGTTATCTATTTTTTATTTATTG

TTTTAGAATCTTTGCTTCTTTTACGGC

>Asia1_Indonesia_HE653683_HE653683.1 Bemisia tabaci mitochondrial partial COI gene for cytochrome oxidase subunit 1, isolate Indo#02

GGAAACTAGAGGTATTTGGCAGGTTAGGAATAATTTATGCTATAATAACTATTGGCATCTTGGGGTTTAT

TGTTTGAGGTCATCATATATTTACTGTTGGTATAGATGTTGATACTCGAGCTTATTTTACTTCAGCTACT

ATGGTTATTGCTGTTCCAACTGGGATTAAGATTTTCAGGTGGCTTGCTACTTTAGGTGGAATAAAATCCA

ATAAATTAAGGCCGTTAGTTCTTTGATTTACAGGATTTTTATTCTTATTTACCATGGGTGGACTAACTGG

GATTATTCTTGGTAATTCTTCTGTTGATGTTTGTTTGCATGATACTTATTTTGTTGTTGCTCATTTTCAT

TATGTTTTATCCATAGGAATCATTTTTGCTATCATAGGAGGTTTTATTTACTGATTTCCATTAATCTTAG

GTCTAACATTAAATAACCATAATTTGGTATCTCAGTTTTATATTATATTTTTGGGCGTTAACCTAACATT

TTTTCCACAACACTTTCTTGGATTAAGCGGAATACCTCGTCGGTATTCAGATTATCCTGATTGTTATCTC

ATATGAAATAAAATTTCTTCTGCGGGGAGTATCTTGAGAATTATTTCTGTTATCTATTTTTTATTTATTG

TTTTAGAATCTTTGCTTCTTTTACGGC

>Asia1_India_AJ748365_AJ748365.1 Bemisia tabaci mitochondrial partial coi gene for cytochrome oxidase I subunit, specific host field bean, from India, Karnataka, Bangalore, Hosakote, Jyotipura

GGAAACTTGAGGTATTTGGCAGGTTAGGAATAATTTATGCTATAATAACTATTGGCATTTTGGGGTTTAT

TGTTTGAGGTCATCATATATTTACTGTTGGTATAGATGTTGATACTCGAGCTTATTTTACTTCAGCTACT

ATGGTTATTGCTGTTCCAACTGGGATTAAGATTTTCAGGTGGCTTGCTACTTTAGGTGGAATAAAATCCA

ATAAATTAAGGCCGTTAGTTCTTTGATTTACAGGATTTTTATTCTTATTTACCATGGGTGGACTAACTGG

GATTATTCTTGGGAATTCTTCTGTTGATGTTTGTTTGCATGATACTTATTTTGTTGTTGCTCATTTTCAT

TATGTTTTATCCATAGGAATCATTTTTGCTATCATAGGAGGTTTTATTTACTGATTTCCATTAATCTTAG

GTCTAACATTAAATAACCATAATTTGGTATCTCAGTTTTATATTATATTTCTGGGCGTTAACCTAACATT

TTTTCCACAACACTTTCTTGGATTAAGCGGAATACCTCGTCGGTATTCAGATTATCCTGATTGTTATCTC

ATATGAAATAAAATTTCTTCTGCGGGGAGTATCTTGAGAATTATTTCTGTTATCTATTTTTTATTTATTG

TTTTAGAATCTTTGCTTCTTTTACGGC

>Australia-1 II JX416166.1 Australia Northern Territory AUSII Bemisia tabaci isolate W15 cytochrome c oxidase subunit 1 gene, partial cds; mitochondrial

GAAAACTTGAGGTATTTGGGAGACTAGGAATAATCTACGCTATAATAACTATTGGTATTCTCGGCTTTAT

TGTTTGGGGTCATCACATATTTACTGTTGGAATAGATGTTGATACTCGAGCTTATTTTACCTCAGCCACT

ATGATTATTGCTGTTCCAACTGGAATTAAAATTTTTAGGTGGCTTGCTACCTTAGGTGGAATAAGGGCTA

ACAAATTTAGTCCTCTTGTGCTTTGATTCACAGGATTTTTATTCTTATTTACCATGGGCGGATTAACTGG

GATTATTCTTGGTAATTCTTCTGTTGATGTCTGCTTGCACGATACTTATTTTGTTGTTGCTCATTTTCAT

TATGTTTTATCTATGGGAATTATTTTTGCTATCGTGGGAGGTCTTATTTATTGATTTCCATTAATTCTAG

GTTTAACATTAAATAGACATAATTTAGTGTCTCAGTTTTACGTTATATTTTTGGGAGTTAACTTAACTTT

TTTTCCACAACATTTTCTTGGTTTAAGCGGGATACCTCGTCGGTACTCAGATTATCCCGATTGCTATCTA

ATGTGAAATAAAATTTCTTCTGCGGGGAGTATCTTGAGTATTATTTCTGTTATCTATTTTCTATTTATTG

TTTTAGAATCTTTGCTGCTTTTGCGGT

>Australia II KC109797.1 Australia Western Australia Kununurra AUSII Bemisia tabaci isolate AN6 cytochrome c oxidase subunit 1 gene, partial cds; mitochondrial

GAAAACTTGAGGTATTTGGGAGACTAGGAATAATCTACGCTATAATAACTATTGGTATTCTCGGCTTTAT

TGTTTGGGGTCATCACATATTTACCGTTGGAATAGATGTTGATACTCGAGCTTATTTTACCTCCGCCACT

ATGATTATTGCTGTTCCAACTGGAATTAAAATTTTTAGGTGGCTTGCTACCTTAGGTGGAATAAGGGCTA

ACAAATTTAGTCCTCTTGTGCTTTGATTCACAGGATTTTTATTCTTATTTACCATGGGCGGGTTAACTGG

GATTATTCTTGGTAATTCTTCTGTTGATGTCTGCTTGCACGATACTTATTTTGTTGTTGCTCATTTTCAT

TATGTTTTATCTATGGGAATTATTTTTGCTATCGTGGGAGGTCTTATTTATTGATTTCCATTAATTCTAG

GTTTAACATTAAATAGACATAATTTAGTGTCTCAGTTTTACGTTATATTTTTGGGAGTTAACTTAACTTT

TTTCCCACAACATTTTCTTGGTTTAAGCGGGATACCTCGGCGGTACTCAGATTATCCCGATTGCTATTTA

ATGTGAAATAAAATTTCTTCTGCGGGGAGTATCTTGAGTATTATTTCTGTTATCTATTTTCTATTTATTG

TTTTAGAATCTTTGCTGCTTTTGCGGT

>Australia_Bundaberg_GU086328_GU086328.1 Bemisia tabaci isolate H14_Australia_Bundaberg cytochrome c oxidase subunit 1 gene, partial cds; mitochondrial

GAAAACTTGAGGTATTTGGGAGACTCGGAATAATTTACGCTATAATAACTATTGGTATTCTTGGTTTTAT

TGTTTGGGGTCATCATATATTTACTGTTGGGATAGATGTTGATACTCGAGCTTACTTTACTTCAGCCACT

ATAATTATTGCTGTTCCAACTGGAATCAAAATTTTTAGGTGGCTTGCTACCTTAGGTGGAATAAGGGCTA

ACAAATTTAGTCCTCTTGTGCTTTGATTCACAGGATTTTTATTCTTATTTACCATGGGTGGGTTAACTGG

GATTATTCTTGGTAATTCTTCTGTTGATGTCTGCTTGCACGATACTTATTTTGTTGTTGCTCATTTTCAT

TATGTTTTATCTATGGGAATTATTTTTGCTATCGTGGGAGGTCTCATTTATTGATTTCCATTAATTCTAG

GTTTAACATTAAATAGACATAATTTGGTTTCTCAGTTTTACGTTATATTTTTGGGAGTTAATTTAACGTT

TTTTCCACAACATTTCCTTGGTTTAAGCGGAATACCTCGCCGGTACTCAGATTATCCCGATTGCTATCTA

ATATGAAATAAAATTTCTTCTGCGGGGAGTATCTTGAGCATTATTTCTGTTATCTATTTTCTATTTATTG

TTTTAGAATCTTTGCTGCTTTTGCGGT

>Australia_Indonesia_HQ457045_HQ457045.1 Bemisia tabaci cytochrome oxidase subunit 1 gene, partial cds; mitochondrial

GGAAACTTGAAGTATTTGGGAGGCTAGGGTTAATCTATGCTATAATAACTATTGGTATTCTTGGTTTTAT

TGTCTGGGGTCATCATATATTTACTGTTGGGATAGATGTTGATACTCGAGCTTATTTTACTTCAGCCACT

ATAATTATTGCTGTTCCAACTGGAATCAAAATTTTTAGGTGACTTGCCACCTTAGGTGGGATAAAGACTA

ATAAATTTAGTCCTCTTGTTCTTTGATTTACAGGATTTTTATTTTTATTCACTATGGGTGGATTGACGGG

AATTATTCTTGGCAATTCGTCTGTTGATGTTTGTTTGCATGATACTTATTTTGTTGTTGCTCATTTTCAT

TATGTTTTATCCATAGGAATTATTTTTGCTATTATAGGAGGTCTTATTTATTGATTTCCATTAATCTGGG

GTCTAACATTAAATGAGCACAACTTAGTATCTCAATTTTATGTTATGTTCTTAGGAGTTAACTTAACATT

TTTCCCGCAGCACTTTCTTGGACTTAGTGGGATACCTCGGCGATACTCAGATTATCCCGATTGTTATCTA

ATGTGGAATAAAATTTCTTCTGCAGGAAGTATTTTAAGTATTATTTCGGTTATTTACTTTTTATTTATCG

TTTTAGAATCTTTACTTCTCTTGCGGT

>China1_China_Anhui_GQ139495_GQ139495.1 Bemisia tabaci isolate CN-AH-NB cytochrome oxidase subunit 1 (CO1) gene, partial cds; mitochondrial

GAAAACTTGAGGTATTTGGAAGGTTGGGCATGATTTATGCTATAATAACTATTGGTATTTTAGGTTTTAT

TGTTTGAGGTCATCATATATTTACTGTTGGGATAGATGTTGATACTCGGGCTTATTTTACTTCGGCTACC

ATGATTATCGCTGTCCCAACAGGAATTAAAATTTTTAGGTGGCTTGCTAGTTTAGGTGGAATAAAATCTA

ATAAATCTAGGCCGCTTGTTCTTTGGTTTACGGGATTTTTGTTTTTATTTACTATGGGTGGATTAACAGG

AATCATTCTTGGTAATTCTTCTGTTGATGTTTGTTTGCACGATACTTATTTTGTTGTTGCTCATTTTCAT

TATGTTCTATCTATGGGAATTATCTTTGCTATTATGGGAGGTTTTATTTATTGATTTCCACTGATTTTAG

GTCTAACACTAAATAGTCACAATTTAGTATCTCAGTTTTATATTATATTTTTGGGAGTAAACTTAACATT

TTTTCCGCAGCATTTTCTTGGGCTAGGGGGGATACCTCGACGATATTCTGATTATCCTGATTGTTATCTA

ATGTGAAATAAAATTGCTTCTGCGGGGAGTATCTTAAGTGTTATTTCTGTTATTTATTTTTTATTTATCG

TTTTAGAATCTTTACTTCTTTTGCGTT

>China1_China_Zhejiang_DQ309078_DQ309078.1 Bemisia tabaci haplotype ZHJ-3 cytochrome oxidase subunit 1 (COI) gene, partial cds; mitochondrial

GAAAACTTGAGGTATTTGGAAGGTTGGGTATGATTTATGCTATAATAACTATTGGTATTTTAGGTTTTAT

TGTTTGAGGTCATCATATATTTACTGTTGGGATAGATGTTGATACTCGGGCTTATTTTACTTCGGCTACC

ATGATTATCGCTGTCCCAACAGGAATTAAAATTTTTAGGTGGCTTGCTAGTTTAGGTGGAATAAAATCTA

ATAAATCTAGGCCGCTTGTTCTTTGGTTTACGGGATTTTTGTTTTTATTTACTATGGGTGGATTAACAGG

AATCATTCTTGGTAATTCTTCTGTTGATGTTTGTTTGCACGATACTTATTTTGTTGTTGCTCATTTTCAT

TATGTTCTATCTATGGGAATTATCTTTGCTATTATGGGAGGTTTTATTTATTGATTTCCACTGATTTTAG

GTCTAACACTAAATAGTCACAATTTAGTATCTCAGTTTTATATTATATTTTTGGGAGTAAACTTAACATT

TTTTCCGCAGCATTTTCTTGGGCTAGGGGGGATACCTCGACGATATTCTGATTATCCTGATTGTTATCTA

ATGTGAAATAAAATTGCTTCTGCGGGGAGTATCTTAAGTGTTATTTCTGTTATTTATTTTTTATTTATCG

TTTTAGAATCTTTACTTCTTTTGCGTT

>China1_China_Jiangxi_HM137328_HM137328.1 Bemisia tabaci isolate j16 cytochrome oxidase subunit 1 gene, partial cds; mitochondrial

GAAAACTTGAGGTATTTGGAAGGTTGGGTATGATTTATGCTATAATAACTATTGGTATTTTAGGTTTTAT

TGTTTGAGGTCATCATATATTTACTGTTGGAATAGATGTTGATACTCGGGCTTATTTTACTTCGGCTACC

ATGATTATCGCTGTCCCAACAGGAATTAAAATTTTTAGGTGGCTTGCTAGTTTAGGTGGAATAAAATCTA

ATAAATCTAGGCCGCTTGTTCTTTGGTTTACGGGATTTTTGTTTTTATTTACTATGGGTGGATTAACAGG

AATCATTCTTGGTAATTCTTCTGTTGATGTTTGTTTGCACGATACTTATTTTGTTGTTGCTCATTTTCAT

TATGTTCTATCTATGGGAATTATCTTTGCTATTATGGGAGGTTTTATTTATTGATTTCCACTGATTTTAG

GTCTAACACTAAATAGTCACAATTTAGTATCTCAGTTTTATATTATATTTTTGGGAGTAAACTTAACATT

TTTTCCGCAGCATTTTCTTGGGCTAGGGGGGATACCTCGACGATATTCTGATTATCCTGATTGTTATCTA

ATGTGAAATAAAATTGCTTCTGCGGGGAGTATCTTAAGTGTTATTTCTGTTATTTATTTTTTATTTATCG

TTTTAGAATCTTTACTTCTTTTGCGTT

>China1_China_Sichuan_HM137351_HM137351.1 Bemisia tabaci isolate ch39 cytochrome oxidase subunit 1 gene, partial cds; mitochondrial

GAAAACTTGAGGTATTTGGAAGGTTGGGTATGATTTATGCTATAATAACTATTGGTATTTTAGGTTTTAT

TGTTTGAGGTCATCATATATTTACTGTTGGGATAGATGTTGATACTCGGGCTTATTTTACTTCGGCTACC

ATGATTATCGCTGTCCCAACAGGAATTAAAATTTTTAGGTGGCTTGCTAGTTTAGGTGGAATAAAATCTA

ATAAATCTAGGCCGCTTGTTCTTTGGTTTACGGGATTTTTGTTTTTATTTACTATGGGTGGATTAACAGG

AATCATTCTTGGTAATTCTTCTGTTGATGTTTGTTTGCACGATACTTATTTTGTTGTTGCTCATTTTCAT

TATGTTCTATCTATGGGAATTATCTTTGCTATTATGGGAGGTTTTATTTATTGATTTCCACTGATTTTAG

GTCTAACACTAAATAGTCACAATTTAGTATCTCAGTTTTATATTATATTTTTGGGAGTAAACTTAACATT

TTTTCCGCAGCATTTTCTTGGGCTAGGGGGGATACCTCGACGATATTCTGATTATCCTGATTGTTATCTA

ATGTGAAATAAGATTGCTTCTGCGGGGAGTATCTTAAGTGTTATTTCTGTTATTTATTTTTTATTTATCG

TTTTAGAATCTTTACTTCTTTTGCGTT

>China1_China_Sichuan_HM137315_HM137315.1 Bemisia tabaci isolate s3 cytochrome oxidase subunit 1 gene, partial cds; mitochondrial

GAAAACTTGAGGTATTTGGAAGGTTGGGTATGATTTATGCTATAATAACTATTGGTATTTTAGGTTTTAT

TGTTTGAGGTCATCATATATTTACTGTTGGGATAGATGTTGATACTCGGGCTTATTTTACTTCGGCTACC

ATGATTATCGCTGTCCCAACAGGAATTAAAATTTTTAGGTGGCTTGCTAGTTTAGGTGGAATAAAATCTA

ATAAATCTAGGCCGCTTGTTCTTTGGTTTACGGGATTTTTGTTTTTATTTACTATGGGTGGATTAACAGG

AATCATTCTTGGTAATTCTTCTGTTGATGTTTGTTTGCACGATACTTATTTTGTTGTTGCTCATTTTCAT

TATGTTCTATCTATGGGAATTATCTTTGCTATTATGGGAGGTTTTATTTATTGATTTCCACTGATTTTAG

GTCTAACACTAAATAGTCACAATTTAGTATCTCAGTTTTATATTATATTTTTGGGAGTAAACTTAACATT

TTTTCCGCAGCATTTTCTTGGGCTAGGGGGGATACCTCGACGATATTCTGATTATCCTGATTGTTATCTA

ATGTGAAATAAAATTGCTTCTGCGGGGAGTATCTTAAGTGTTATCTCTGTTATTTATTTTTTATTTATCG

TTTTGGAATCTTTACTTCTTTTGCGTT

>China2_HQ916820 Bemisia tabaci isolate LC8 cytochrome oxidase subunit I (COI) gene,partial cds; mitochondrial

GAAAACTTGAGGTATTTGGAAGGTTGGGCATGATTTATGCTATAATAACTATTGGTATTTTAGGTTTTAT

TGTTTGAGGTCATCATATATTTACTGTTGGGATAGATGTGGATACTCGGGCTTATTTTACTTCGGCTACT

ATGATTATTGCTGTTCCAACAGGAATTAAAATTTTTAGGTGGCTTGCTAGTTTAGGTGGAATAAAATCTA

ATAAATCTAGGCCTCTTGTTCTTTGGTTTACAGGATTTTTGTTTTTATTTACTATAGGTGGATTAACAGG

AATTATTCTTGGTAATTCTTCTGTTGATGTTTGTTTGCACGATACTTATTTTGTTGTTGCTCATTTTCAT

TATGTTCTATCTATAGGAATTATTTTTGCTATTATGGGGGGTTTTATTTATTGATTTCCACTAATTTTAG

GTCTAACACTAAACAGTCACAATTTAGTATCTCAGTTTTATATTATATTTTTAGGAGTAAACTTAACATT

TTTTCCACAGCACTTTCTTGGGCTAAGGGGGATACCTCGACGATATTCTGATTATCCTGATTGTTACTTA

ATGTGAAATAAAATTGCTTCTGCGGGGAGTATCTTAAGCGTTATTTCTGTTATTTATTTTTTATTTATTG

TTTTAGAGTCTTTACTCCTTTTGCGTT

>China2_KC113546 Bemisia tabaci isolate M22 cytochrome oxidase subunit I (COI) gene,partial cds; mitochondrial

GAAAACTTGAGGTATTTGGAAGGTTGGGCATGATTTATGCTATAATAACTATTGGTATTTTAGGTTTTAT

TGTTTGAGGTCATCATATATTTACTGTTGGGATAGATGTGGATACTCGGGCTTATTTTACTTCGGCTACT

ATGATTATTGCTGTTCCAACAGGAATTAAAATTTTTAGGTGGCTTGCTAGTTTAGGTGGAATAAAATCTA

ATAAATCTAGGCCTCTTGTTCTTTGGTTTACAGGATTTTTGTTTTTATTTACTATAGGTGGATTAACAGG

AATTATTCTTGGTAATTCTTCTGTTGATGTTTGTTTGCACGATACTTATTTTGTTGTTGCTCATTTTCAT

TATGTTCTATCTATAGGAATTATTTTTGCTATTATGGGGGGTTTTATTTATTGATTTCCACTAATTTTAG

GTCTAACACTAAACAGTCACAATTTAGTATCTCAGTTTTATATTATATTTTTAGGAGTAAACTTAACATT

TTTTCCACAGCACTTTCTTGGGCTAAGGGGGATACCTCGACGATATTCTGATTATCCTGATTGTTACTTA

ATGTGAAATAAAATTGCTTCTGCGGGGAGTATCTTAAGCGTTATTTCTGTTATTTATTTTTTATTTATTG

TTTTAGAATCTTTACTCCTTTTGCGTT

>China2_KC959606 Bemisia tabaci isolate 56-KualaLipis cytochrome c oxidase subunit 1(CO1) gene, partial cds; mitochondrial

GAAAACTTGAGGTATTTGGAAGGTTGGGCATGATTTATGCTATAATAACTATTGGTATTTTAGGTTTTAT

TGTTTGAGGTCATCATATATTTACTGTTGGGATAGATGTGGATACTCGGGCTTATTTTACTTCGGCTACT

ATGATTATTGCTGTTCCAACAGGAATTAAAATTTTTAGGTGGCTTGCTAGTTTAGGTGGAATAAAATCTA

ATAAATCTAGGCCTCTTGTTCTTTGGTTTACAGGATTTTTGTTTTTATTTACTATAGGTGGATTAACAGG

AATTATTCTTGGTAATTCTTCTGTTGATGTTTGTTTGCACGACACTTATTTTGTTGTTGCTCATTTTCAT

TATGTTCTATCTATAGGAATTATTTTTGCTATTATGGGGGGTTTTATTTATTGATTTCCACTAATTTTAG

GTCTAACACTAAACAGTCACAATTTAGTATCTCAGTTTTATATTATATTTTTAGGAGTAAACTTAACATT

TTTTCCACAGCACTTTCTTGGGCTAAGGGGGATACCTCGACGATATTCTGATTATCCTGATTGTTACTTA

ATGTGAAATAAAATTGCTTCTGCGGGGAGTATCTTAAGCGTTATTTCTGTTATTTATTTTTTATTTATTG

TTTTAGAATCTTTACTCCTTTTGCGTT

>China2_KC959601 Bemisia tabaci isolate 23-KualaLipis cytochrome c oxidase subunit 1(CO1) gene, partial cds; mitochondrial

GAAAACTTGAGGTATTTGGGAGGTTGGGCATGATTTATGCTATAATAACTATTGGTATTTTAGGTTTTAT

TGTTTGAGGTCATCATATATTTACTGTTGGGATAGATGTGGATACTCGGGCTTATTTTACTTCGGCTACT

ATGATTATTGCTGTTCCAACAGGAATTAAAATTTTTAGGTGGCTTGCTAGTTTAGGTGGAATAAAATCTA

ATAAATCTAGGCCTCTTGTTCTTTGGTTTACAGGATTTTTGTTTTTATTTACTATAGGTGGATTAACAGG

AATTATTCTTGGTAATTCTTCTGTTGATGTTTGTTTGCACGACACTTATTTTGTTGTTGCTCATTTTCAT

TATGTTCTATCTATAGGAATTATTTTTGCTATTATGGGGGGTTTTATTTATTGATTTCCACTAATTTTAG

GTCTAACACTAAACAGTCACAATTTAGTATCTCAGTTTTATATTATATTTTTAGGAGTAAACTTAACATT

TTTTCCACAGCACTTTCTTGGGCTAAGGGGGATACCTCGACGATATTCTGATTATCCTGATTGTTACTTA

ATGTGAAATAAAATTGCTTCTGCGGGGAGTATCTTAAGCGTTATTTCTGTTATTTATTTTTTATTTATTG

TTTTAGAATCTTTACTCCTTTTGCGTT

>China2_KC959599 Bemisia tabaci isolate 29-KualaLipis cytochrome c oxidase subunit 1(CO1) gene, partial cds; mitochondrial

GAAAACTTGAGGTATTTGGGAGGTTGGGCATGATTTATGCTATAATAACTATTGGTATTTTAGGTTTTAT

TGTTTGAGGTCATCATATATTTACTGTTGGGATAGATGTGGATACTCGGGCTTATTTTACTTCGGCTACT

ATGATTATTGCTGTTCCAACAGGAATTAAAATTTTTAGGTGGCTTGCTAGTTTAGGTGGAATAAAATCTA

ATAAATCTAGGCCTCTTGTTCTTTGGTTTACAGGATTTTTGTTTTTATTTACTATAGGTGGATTAACAGG

AATTATTCTTGGTAATTCTTCTGTTGATGTTTGTTTGCACGACACTTATTTTGTTGTTGCTCATTTTCAT

TATGTTTTATCTATAGGAATTATTTTTGCTATTATGGGGGGTTTTATTTATTGATTTCCACTAATTTTAG

GTCTAACACTAAACAGTCACAATTTAGTATCTCAGTTTTATATTATATTTTTAGGAGTAAACTTAACATT

TTTTCCACAGCACTTTCTTGGGCTAAGGGGGATACCTCGACGATATTCTGATTATCCTGATTGTTACTTA

ATGTGAAATAAAATTGCTTCTGCGGGGAGTATCTTAAGCGTTATTTCTGTTATTTATTTTTTATTTATTG

TTTTAGAATCTTTACTCCTTTTGCGTT

>China2_KC959600 Bemisia tabaci isolate 31-KualaLipis cytochrome c oxidase subunit 1(CO1) gene, partial cds; mitochondrial

GAAAACTTGAGGTATTTGGAAGGTTGGGTATGATTTATGCTATAATAACTATTGGTATTTTAGGTTTTAT

TGTTTGAGGTCATCATATATTTACTGTTGGGATAGATGTGGATACTCGAGCTTATTTTACTTCGGCTACT

ATGATTATCGCTGTTCCAACAGGAATTAAAATTTTTAGGTGGCTTGCTAGTTTAGGTGGAATAAAATCTA

ATAAATCTAGGCCTCTTGTTCTTTGGTTTACAGGATTTTTGTTTTTATTTACTATAGGTGGATTAACAGG

AATTATTCTTGGTAATTCTTCTGTTGATGTTTGTTTGCACGATACTTATTTTGTTGTTGCTCATTTTCAT

TATGTTCTATCTATAGGAATTATTTTTGCTATTATGGGGGGTTTTATTTATTGATTTCCACTAATTTTAG

GTCTAACACTAAACAGTCACAATTTAGTATCTCAGTTTTATATTATATTTTTAGGAGTAAACTTAACATT

TTTTCCACAGCACTTTCTTGGGCTAAGGGGGATACCTCGACGATATTCTGATTATCCTGATTGTTACTTA

ATGTGAAATAAAATTGCTTCTGCGGGGAGTATCTTAAGCGTTATTTCTGTTATTTATTTTTTATTTATTG

TTTTAGAATCTTTACTCCTTTTGCGTT

>China2_JX678666 Bemisia tabaci isolate 12-F10-Gua Musang cytochrome c oxidasesubunit 1 (CO1) gene, partial cds; mitochondrial

GAAAACTTGAGGTATTTGGAAGGTTGGGTATGATTTATGCTATAATAACTATTGGTATTTTAGGTTTTAT

TGTTTGAGGTCATCATATATTTACTGTTGGGATAGATGTGGATACTCGAGCTTATTTTACTTCGGCTACT

ATAATTATCGCTGTTCCAACAGGAATTAAAATTTTTAGGTGGCTTGCTAGTTTAGGTGGAATAAAATCTA

ATAAATCTAGGCCTCTTGTTCTTTGGTTTACAGGATTTTTGTTTTTATTTACTATAGGTGGATTAACAGG

AATTATTCTTGGTAATTCTTCTGTTGATGTTTGTTTGCACGATACTTATTTTGTTGTTGCTCATTTTCAT

TATGTTCTATCTATAGGAATTATTTTTGCTATTATGGGGGGTTTTATTTATTGATTTCCACTAATTTTAG

GTCTAACACTAAACAGTCACAATTTAGTATCTCAGTTTTATATTATATTTTTAGGAGTAAACTTAACATT

TTTTCCACAGCACTTTCTTGGGCTAAGGGGGATACCTCGACGATATTCTGATTATCCTGATTGTTACTTA

ATGTGAAATAAAATTGCTTCTGCGGGGAGTATCTTAAGCGTTATTTCTGTTATTTATTTTTTATTTATTG

TTTTAGAATCTTTACTCCTTTTGCGTT

>AsiaII_1_India_Varanasi_JN855579_JN855579.1

GAAAGCTTGAAGTATTTGGCAGATTAGGTATAATTTATGCTATAGTAACGATTGGCATTCTAGGTTTTAT

TGTGTGAGGTCATCATATATTTACTGTTGGAATAGATGTGGACACTCGGGCTTATTTTACTTCAGCTACT

ATGATTATTGCTGTTCCGACTGGAATTAAAATCTTTAGGTGACTTGCTACTCTAGGTGGAATAAAGTTTA

ACATATTTAGTCCGCTTGGACTTTGGTTTGCCGGATTTCTTTTCTTATTTACTATGGGTGGATTAACTGG

AATTATTCTGGGTAATTCTTCTGTTGATGTCTGCTTACATGATACTTACTTTGTTGTTGCTCATTTTCAT

TATGTTTTATCTATAGGAATTATCTTTGCTATCGTGGGAGGAGTTATTTATTGATTTCCGGTAATCTTGG

GATTAACACTAAATAGACATAGTTTGGTATCACAGTTTTACATTATGTTTTTGGGAGTAAATTTAACGTT

TTTCCCGCAACATTTTCTTGGGCTGAGAGGTATACCTCGTCGCTATTCAGATTATCCTGACTGTTATCTA

ATATGAAATAAAATTTCTTCTGCGGGGAGAATTTTGAGCACTATTTCTGTTATTTATTTTTTATTTATTG

TTTTAGAGTCTTTGCTTCTTTTGCGGT

>AsiaII_1_India_JN703449_JN703449.1

GAAAGCTTGAAGTATTTGGCAGATTAGGTATAATTTATGCTATAGTAACGATTGGCATTCTAGGTTTTAT

TGTGTGAGGTCATCATATATTTACTGTTGGAATAGATGTGGACACTCGGGCTTATTTTACTTCAGCTACT

ATGATTATTGCTGTTCCGACTGGAATTAAAATCTTTAGGTGACTTGCTACTCTAGGTGGAATAAAGTTTA

ACATATTTAGTCCGCTTGGACTTTGGTTTGCTGGATTTCTTTTCTTATTTACTATGGGTGGATTAACTGG

AATTATTCTGGGTAATTCTTCTGTTGATGTCTGCTTACATGATACTTACTTTGTTGTTGCTCATTTTCAT

TATGTTTTATCTATAGGAATTATCTTTGCTATCGTGGGAGGAGTTATTTATTGATTTCCGGTAATCTTGG

GATTAACACTAAATAGACATAGTTTGGTATCACAGTTTTACATTATGTTTTTGGGAGTAAATTTAACGTT

TTTCCCGCAACATTTTCTTGGGCTGAGAGGTATACCTCGTCGCTATTCAGATTATCCTGACTGTTATCTA

ATATGAAATAAAATTTCTTCTGCGGGGAGAATTTTGAGCACTATTTCTGTTATTTATTTTTTATTTATTG

TTTTAGAGTCTTTGCTTCTTTTACGGT

>AsiaII_1_India_Punjab_HM590129_HM590129.1

GAAAGCTTGAAGTATTTGGCAGATTAGGTATAATTTATGCTATAGTAACGATTGGCATCCTAGGTTTTAT

TGTGTGAGGTCATCATATATTTACTGTTGGAATAGATGTGGACACTCGGGCTTATTTTACTTCAGCTACT

ATGATTATTGCTGTTCCGACTGGAATTAAAATCTTTAGGTGACTTGCTACTCTAGGTGGAATAAAGTTTA

ACATATTTAGTCCGCTTGGACTTTGGTTTGCTGGATTTCTTTTCTTATTTACTATGGGTGGATTAACTGG

AATTATTCTGGGTAATTCTTCTGTTGATGTCTGCTTACATGATACTTACTTTGTTGTTGCTCATTTTCAT

TATGTTTTATCTATAGGAATTATCTTTGCTATCGTGGGAGGAGTTATTTATTGATTTCCGGTAATCTTGG

GACTAACACTAAATAGACATAGTTTGGTATCACAGTTTTACATTATGTTTTTGGGAGTAAATTTAACGTT

TTTCCCGCAACATTTTCTTGGGCTGAGAGGTATACCTCGTCGCTATTCAGATTATCCTGACTGTTATCTA

ATATGAAATAAAATTTCTTCTGCGGGGAGAATTTTGAGCACTATTTCTGTTATTTATTTTTTATTTATTG

TTTTAGAGTCTTTGCTTCTTTTGCGGT

>AsiaII_1_China_Hainan_HM137337_HM137337.1

GAAAGCTTGAAGTATTTGGCAGATTAGGTATAATTTATGCTATAGTAACGATTGGCATTCTAGGTTTTAT

TGTGTGAGGTCATCATATATTTACTGTTGGAATAGATGTGGACACTCGGGCTTATTTTACTTCAGCTACT

ATGATTATTGCTGTTCCGACTGGAATTAAAATCTTTAGGTGACTTGCTACTCTAGGTGGAATAAAGTTTA

ACATATTTAGTCCGCTTGGACTTTGGTTTGCTGGATTTCTTTTCTTATTTACTATGGGTGGATTAACTGG

AATTATTCTGGGTAATTCTTCTGTTGATGTCTGCTTACATGATACTTACTTTGTTGTTGCTCATTTTCAT

TATGTTTTATCTATAGGAATTATCTTTGCTATCGTGGGAGGAGTTATTTATTGATTTCCGGTAATCTTGG

GATTAACGCTAAATAGACATAGTTTGGTATCACAGTTTTACATTATGTTTTTGGGAGTAAATTTAACGTT

TTTCCCGCAACATTTTCTTGGGCTGAGAGGTATACCTCGTCGCTATTCAGATTATCCTGACTGTTATCTA

ATATGGAATAAAATTTCTTCTGCGGGGAGAATTTTGAGCATTATTTCTGTTATTTATTTTTTATTTATTG

TTTTAGAGTCTTTGCTTCTTTTGCGGT

>AsiaII_1_China_Zhejiang_AJ867557_AJ867557.1

GAAAGCTTGAAGTATTTGGCAGATTAGGTATAATTTATGCTATAGTAACGATTGGCATTCTAGGTTTTAT

TGTGTGAGGTCATCATATATTTACTGTTGGAATAGATGTGGACACTCGGGCTTATTTTACTTCAGCTACT

ATGATTATTGCTGTTCCGACTGGAATTAAAATCTTTAGGTGACTTGCTACTCTAGGTGGAATAAAGTTTA

ACATATTTAGTCCGCTTGGACTTTGGTTTGCTGGATTTCTTTTCTTATTTACTATGGGTGGATTAACTGG

AATTATTCTGGGTAATTCTTCTGTTGATGTCTGCTTACATGATACTTACTTTGTTGTTGCTCATTTTCAT

TATGTTTTATCTATAGGAATTATCTTTGCTATCGTGGGAGGAGTTATTTATTGATTTCCGGTAATCTTGG

GATTAACACTAAATAGACATAGTTTGGTATCACAGTTTTACATTATGTTTTTGGGAGTAAATTTAACGTT

TTTCCCGCAACATTTTCTTGGGCTGAGAGGTATACCTCGTCGCTATTCAGATTATCCTGACTGTTATCTA

ATATGAAATAAAATTTCTTCTGCGGGGAGAATTTTGAGCATTATTTCTGTTATTTATTTTTTATTTATTG

TTTTAGAGTCTTTGCTTCTTTTGCGGT

>AsiaII_1_India_Akola_JN855567_JN855567.1

GAAAGCTTGAAGTATTTGGCAGATTAGGTATAATTTATGCTATAGTAACGATTGGCATTCTAGGTTTTAT

TGTGTGAGGTCATCATATATTTACTGTTGGAATAGATGTGGACACTCGGGCTTATTTTACTTCAGCTACT

ATGATTATTGCTGTTCCGACTGGAATTAAAATCTTTAGGTGACTTGCTACTCTAGGTGGAATAAAGTTTA

ACATATTTAGTCCGCTTGGACTTTGGTTTGCTGGATTTCTTTTCTTATTTACTATGGGTGGATTAACTGG

AATTATTCTGGGTAATTCTTCTGTTGATGTCTGCTTACATGATACTTACTTTGTTGTTGCTCATTTTCAT

TATGTTTTATCTATAGGAATTATCTTTGCTATCGTGGGAGGAGTTATTTATTGATTTCCGGTAATCTTGG

GATTAACACTAAATAGACATAGTCTGGTATCACAGTTTTACATTATGTTTTTGGGAGTAAATTTAACGTT

TTTCCCGCAACATTTTCTTGGGCTGAGAGGTATACCTCGTCGCTATTCAGATTATCCTGACTGTTATCTA

ATATGAAATAAAATTTCTTCTGCGGGGAGAATTTTGAGCATTATTTCTGTTATTTATTTTTTATTTATTG

TTTTAGAGTCTTTGCTTCTTTTGCGGT

>AsiaII_1_India_Punjab_HM590132_HM590132.1

GAAAGCTTGAAGTATTTGGCAGATTAGGTATAATTTATGCTATAGTAACGATTGGCATTCTAGGTTTTAT

TGTGTGAGGTCATCATATATTTACTGTTGGAATAGATGTGGACACTCGGGCTTATTTTACTTCAGCTACT

ATGATTATTGCTGTTCCGACTGGAATTAAAATCTTTAGGTGACTTGCTACTCTAGGTGGAATAAAGTTTA

ACATATTTAGCCCGCTTGGACTTTGGTTTGCTGGGTTTCTTTTCTTATTTACTATGGGTGGATTAACTGG

AATTATTCTGGGTAATTCTTCTGTTGATGTCTGCTTACATGATACTTACTTTGTTGTTGCTCATTTTCAT

TATGTTTTATCTATAGGAATTATCTTTGCTATCGTGGGAGGAGTTATTTATTGATTTCCGGTAATCTTGG

GATTAACACTAAATAGACATAGTTTGGTATCACAGTTTTACATTATGTTTTTGGGAGTAAATTTAACGTT

TTTCCCGCAACATTTTCTTGGGCTGAGAGGTATACCTCGTCGCTATTCAGATTATCCTGACTGTTATCTA

ATATGAAATAAAATTTCTTCTGCGGGGAGAATTTTGAGCATTATTTCTGTTATTTATTTTTTATTTATTG

TTTTAGAGTCTTTGCTTCTTTTGCGGT

>AsiaII_1_India_Rajasthan_HM590138_HM590138.1

GAAAGCTTGAAGTATTTGGCAGATTAGGTATAATTTATGCTATAGTAACGATTGGCATCCTAGGTTTTAT

TGTGTGAGGTCATCATATATTTACTGTTGGAATAGATGTGGACACTCGGGCTTATTTTACTTCAGCTACT

ATGATTATTGCTGTTCCGACTGGAATTAAAATCTTTAGGTGACTTGCCACTCTAGGTGGAATAAAGTTTA

ACATATTTAGTCCGCTTGGACTTTGGTTCGCTGGATTTCTTTTCTTATTTACTATGGGTGGATTAACTGG

AATTATTCTGGGTAATTCTTCTGTTGATGTCTGCTTACATGATACTTACTTTGTTGTTGCTCATTTTCAT

TATGTTTTATCTATAGGAATTATCTTTGCTATCGTGGGAGGAGTTATTTATTGATTTCCGGTAATCTTGG

GATTAACACTAAATAGACATAGTTTGGTATCACAGTTTTACATTATGTTTTTGGGAGTAAATTTAACGTT

TTTCCCGCAACATTTTCTTGGGTTGAGAGGTATACCTCGTCGCTATTCAGATTATCCTGACTGTTATCTA

ATATGAAATAAAATTTCTTCTGCGGGGAGAATTTTGAGCACTATTTCTGTTATTTATTTTTTATTTATTG

TTTTAGAGTCTTTGCTTCTTTTGCGGT

>AsiaII_7_China_Fujian_GQ139492_GQ139492.1 Bemisia tabaci isolate CN-FJ-NB cytochrome oxidase subunit 1 (CO1) gene, partial cds; mitochondrial

GAAAACTTGAGGTATTTGGCAGGTTAGGTATAATTTATGCTATAGTAACGATTGGTATTTTAGGTTTTAT

TGTTTGAGGTCATCATATATTTACTGTTGGGATGGATGTTGATACCCGGGCCTATTTTACCTCAGCTACT

ATGATTATTGCTGTTCCGACTGGGATTAAAATTTTTAGGTGACTTGCTACTCTAGGTGGGATAAAATCTA

ATAGATTTAGCCCCCTTGGGCTTTGGTTTACTGGATTTCTTTTTTTATTTACTATGGGTGGGTTAACTGG

GATTATTCTTGGTAATTCTTCTGTTGATGTCTGCTTACATGATACTTATTTTGTTGTTGCTCATTTTCAT

TATGTTTTATCTATAGGAATTATTTTTGCTATTGTGGGAGGGGTTATTTACTGATTTCCGTTAATCTTGG

GCTTAACACTGAATAGCCACAGTCTGGTATCACAGTTTTATATTATGTTTTTGGGAGTAAACTTAACTTT

TTTTCCACAACATTTTCTTGGGCTAAGAGGAATACCTCGCCGATATTCAGACTATCCTGATTGTTACTTG

ATATGAAATAAGATTTCTTCTGCGGGGAGAATTTTGAGCATTATTTCTGTTATTTATTTTTTATTTATTG

TTTTAGAGTCGTTGCTTCTTTTACGTC

>AsiaII_7_India_AJ748378_AJ748378.1 Bemisia tabaci mitochondrial partial coi gene for cytochrome oxidase I subunit, specific host watermelon, from India, Karnataka, Kolar, Chintamani, Talagavara

GAAAACTTGAGGTATTTGGCAGGTTAGGTATAATTTATGCTATAGTAACGATTGGTATTTTAGGTTTTAT

TGTTTGAGGTCATCATATATTTACTGTTGGGATGGATGTTGATACCCGGGCCTATTTTACCTCAGCTACT

ATGATTATTGCTGTTCCGACTGGGATTAAAATTTTTAGGTGACTTGCTACTCTAGGTGGGATAAAATCTA

ATAGATTTAGCCCCCTTGGGCTTTGGTTTACTGGATTTCTTTTTTTATTTACTATGGGTGGGTTAACTGG

GATTATTCTTGGTAATTCTTCTGTTGATGTCTGCTTACATGATACTTATTTTGTTGTTGCTCATTTTCAT

TATGTTTTATCTATAGGAATTATTTTTGCTATTGTGGGAGGGGTTATTTACTGATTTCCGTTAATCTTGG

GCTTAACACTGAATAGCCACAGTCTGGTATCACAGTTTTATATTATGTTTTTGGGAGTAAACTTAACTTT

TTTTCCACAACATTTTCTTGGGCTAAGAGGAATACCTCGCCGATATTCAGACTATCCTGATTGTTACTTG

ATATGAAATAAGATTTCTTCTGCGGGGAGAATTTTGAGCATTATTTCTGTTATTTATTTTTTATTTATTG

TTTTAGAGTCGTTGCTTCTTTTACGTC

>AsiaII_7_Taiwan_DQ174522_(Indonesia)_DQ174522.1 Bemisia tabaci from Gossypium hirsutum cytochrome oxidase subunit I gene, partial cds; mitochondrial

GAAAACTTGAGGTATTTGGCAGGTTAGGTATAATTTATGCTATAGTAACGATTGGTATTTTAGGTTTTAT

TGTTTGAGGTCATCATATATTTACTGTTGGGATGGATGTTGATACCCGGGCCTATTTTACCTCAGCTACT

ATGATTATTGCTGTTCCGACTGGGATTAAAATTTTTAGGTGACTTGCTACTCTAGGTGGGATAAAATCTA

ATAGATTTAGCCCCCTTGGGCTTTGGTTTACTGGATTTCTTTTTTTATTTACTATGGGTGGGTTAACTGG

GATTATTCTTGGTAATTCTTCTGTTGATGTCTGCTTACATGATACTTATTTTGTTGTTGCTCATTTTCAT

TATGTTTTATCTATAGGAATTATTTTTGCTATTGTGGGAGGGGTTATTTACTGATTTCCGTTAATCTTGG

GCTTAACACTGAATAGCCACAGTCTGGTATCACAGTTTTATATTATGTTTTTGGGAGTAAACTTAACTTT

TTTTCCACAACATTTTCTTGGGCTAAGAGGAATACCTCGCCGATATTCAGACTATCCTGATTGTTACTTG

ATATGAAATAAGATTTCTTCTGCGGGGAGAATTTTGAGCATTATTTCTGTTATTTATTTTTTATTTATTG

TTTTAGAGTCGTTGCTTCTTTTACGTC

>AsiaII_7_India_Hyderabad_DQ116650_DQ116650.1 Bemisia tabaci isolation-source cotton cytochrome oxidase subunit I (COI) gene, partial cds; mitochondrial

GAAAACTTGAGGTATTTGGTAGGTTAGGTATAATTTATGCTATAGTAACGATTGGTATTTTAGGTTTTAT

TGTTTGAGGTCATCATATATTTACTGTTGGGATGGATGTTGATACCCGGGCCTATTTTACCTCAGCTACT

ATGATTATTGCTGTTCCGACTGGGATTAAAATTTTTAGGTGACTTGCTACTCTAGGTGGGATAAAATCTA

ATAGATTTAGCCCCCTTGGGCTTTGGTTTACTGGATTTCTTTTTTTATTTACTATGGGTGGGTTAACTGG

GATTATTCTTGGTAATTCTTCTGTTGATGTCTGCTTACATGATACTTATTTTGTTGTTGCTCATTTTCAT

TATGTTCTATCTATAGGAATTATTTTTGCTATTGTGGGAGGGGTTATTTACTGATTTCCTTTAATCTTGG

GCTTAACATTGAATAGCCACAGTCTGGTATCACAGTTTTATATTATGTTTTTGGGAGTAAACTTAACTTT

TTTTCCACAACATTTTCTTGGGCTAAGAGGAATACCTCGCCGATATTCAGACTATCCTGATTGTTACTTG

ATATGAAATAAGATTTCTTCTGCGGGGAGAATTTTGAGCATTATTTCTGTTATTTATTTTTTATTTATTG

TTTTAGAGTCGTTGCTTCTTTTACGTC

>AsiaII_7_India_Madras_DQ116660_DQ116660.1 Bemisia tabaci isolation-source ipomoea cytochrome oxidase subunit I (COI) gene, partial cds; mitochondrial

GAAAACTTGAGGTATTTGGTAGGTTAGGTATAATTTATGCTATAGTAACTATTGGTATTTTAGGTTTTAT

TGTTTGAGGTCATCATATATTTACTGTTGGGATAGATGTTGATACCCGGGCTTATTTTACCTCAGCTACT

ATGATTATTGCTGTTCCGACTGGGATTAAAATTTTTAGGTGACTTGCTACTCTAGGTGGGATAAAATCTA

ATAGATTTAGCCCCCTTGGGCTTTGGTTTACTGGATTTCTTTTTTTATTTACTATGGGTGGGTTAACTGG

GATTATTCTTGGTAATTCTTCTGTTGATGTCTGCTTACATGATACTTATTTTGTTGTTGCTCATTTTCAT

TATGTTTTATCTATAGGAATTATTTTTGCTATTGTGGGAGGGGTTATTTACTGATTTCCGTTAATCTTGG

GCTTAACACTGAATAGCCACAGTCTGGTATCACAGTTTTATATTATGTTTTTGGGAGTAAACTTAACTTT

TTTCCCACAACATTTTCTTGGGCTAAGAGGAATACCTCGCCGATATTCAGATTATCCTGATTGTTACTTG

ATATGAAATAAGATTTCTTCTGCGGGGAGAATTTTGAGCATTATTTCTGTTATTTATTTTTTATTTATTG

TTTTAGAGTCGTTGCTGCTTTTACGTC

>AsiaII_7_India_Madras_DQ116661_DQ116661.1 Bemisia tabaci isolation-source poinsettia cytochrome oxidase subunit I (COI) gene, partial cds; mitochondrial

GAAAACTTGAGGTATTTGGTAGGTTAGGTATAATTTATGCTATAGTAACTATTGGTATTTTAGGTTTTAT

TGTTTGAGGTCATCATATATTTACTGTTGGGATAGATGTTGATACCCGGGCTTATTTTACCTCAGCTACT

ATGATTATTGCTGTTCCGACTGGGATTAAAATTTTTAGGTGACTTGCTACTCTAGGTGGGATAAAATCTA

ATAGATTTAGCCCCCTTGGGCTTTGGTTTACTGGATTTCTTTTTTTATTTACTATGGGTGGGTTAACTGG

GATTATTCTTGGTAATTCTTCTGTTGATGTCTGCTTACATGATACTTATTTTGTTGTTGCTCATTTTCAT

TATGTTTTATCTATAGGAATTATTTTTGCTATTGTGGGAGGGGTTATTTACTGATTTCCGTTAATCTTGG

GCTTAACACTGAATAGCCACAGTCTGGTATCACAGTTTTATATTATGTTTTTGGGAGTAAACTTAACTTT

TTTCCCACAACATTTTCTTGGGCTAAGAGGAATACCTCGCCGATATTCAGATTATCCTGATTGTTACTTG

ATATGAAATAAGATTTCTTCTGCGGGGAGAATTTTGAGCATTATTTCTGTTATTTATTTTTTATTTATTG

TTTTAGAGTCGTTGCTTCTTTTACGTC

>AsiaII_7_India_AJ748375_AJ748375.1 Bemisia tabaci mitochondrial partial coi gene for cytochrome oxidase I subunit, specific host soybean, from India, Karnataka, Bangalore, Hebbal

GAAAACTTGAGGTATTTGGTAGATTAGGTATAATTTATGCTATGGTAACGATTGGTATTTTAGGTTTTAT

TGTTTGAGGTCATCATATATTTACTGTTGGGATGGATGTTGATACTCGGGCCTATTTTACCTCAGCTACC

ATGATTATTGCTGTTCCGACTGGGATTAAAATTTTTAGGTGACTTGCTACTCTAGGTGGGATAAAATCTA

ATAGATTTAGCCCCCTTGGGCTTTGGTTTACTGGATTTCTTTTTTTATTTACTATGGGTGGGTTAACTGG

GATTATTCTTGGTAATTCTTCTGTTGATGTCTGCTTACATGATACTTATTTTGTTGTTGCTCATTTTCAT

TATGTTTTATCTATAGGAATTATTTTTGCTATTGTGGGAGGGGTTATTTACTGATTTCCGTTAATCTTGG

GCTTAACACTGAATAGTCACAGTCTGGTATCCCAGTTTTATATTATGTTTTTGGGAGTAAACTTAACTTT

TTTTCCACAACACTTTCTTGGGCTAAGAGGAATACCTCGCCGATATTCAGATTATCCTGATTGTTACTTG

ATATGAAATAAGATTTCTTCTGCGGGGAGAATTTTGAGCATTATTTCTGTTATTTATTTTTTATTTATTG

TTTTAGAGTCGTTGCTTCTTTTACGTC

>AsiaII_7_Taiwan_Kaohsiung_DQ174521_DQ174521.1 Bemisia tabaci from Euphorbia pulcherrima cytochrome oxidase subunit I gene, partial cds; mitochondrial

GAAAACTTGAGGTATTTGGTAGATTAGGTATAATTTATGCTATGGTAACGATTGGTATTTTAGGTTTTAT

TGTTTGAGGTCATCATATATTTACTGTTGGGATGGATGTTGATACTCGGGCCTATTTTACCTCAGCTACC

ATGATTATTGCTGTTCCGACTGGGATTAAAATTTTTAGGTGACTTGCTACTCTAGGTGGGATAAAATCTA

ATAGATTTAGCCCCCTTGGGCTTTGGTTTACTGGATTTCTTTTTTTATTTACTATGGGTGGGTTAACTGG

GATTATTCTTGGTAATTCTTCTGTTGATGTCTGCTTACATGATACTTATTTTGTTGTTGCTCATTTTCAT

TATGTTTTATCTATAGGAATTATTTTTGCTATTGTGGGAGGGGTTATTTACTGATTTCCGTTAATCTTGG

GCTTAACACTGAATAGTCACAGTCTGGTATCCCAGTTTTATATTATGTTTTTGGGAGTAAACTTAACTTT

TTTTCCACAACATTTTCTTGGGCTAAGAGGAATACCTCGCCGATATTCAGATTATCCTGATTGTTACTTG

ATATGAAATAAGATTTCTTCTGCGGGGAGAATTTTGAGCATTATTTCTGTTATTTATTTTTTATTTATTG

TTTTAGAGTCGTTGCTTCTTTTACGTC

>AsiaII_5_Bangladesh_Joydebpur_AJ748395_AJ748395.1 Bemisia tabaci mitochondrial partial coi gene for cytochrome oxidase I subunit, specific host sponge gourd, from Bangladesh, Joydebpur

GAAAACTTGAAGTATTTGGTAGGTTGGGAATAATTTATGCTATAGTAACTATTGGAATTCTAGGTTTTAT

TGTGTGAGGTCATCACATATTTACCGTTGGGATAGATGTTGATACTCGGGCTTATTTTACTTCAGCCACT

ATAATTATCGCTGTTCCGACCGGAATTAAAATCTTTAGGTGGCTTGCTACTCTAGGTGGAATAAAATCTA

ACAAGTTTAGTCCTCTTGGACTTTGGTTTACTGGATTTCTTTTTTTATTTACCATGGGTGGGTTAACTGG

GATCATTCTTGGTAATTCCTCTGTTGATGTCTGCTTACATGATACTTACTTCGTTGTTGCTCATTTTCAT

TATGTTCTATCTATAGGAATTATTTTTGCTATTGTGGGAGGTGTTATTTATTGATTTCCATTAATTTTGG

GCTTAACACTAAATAGTCATAGTTTGGTATCGCAGTTTTACATTATGTTTTTAGGAGTAAATTTAACGTT

TTTTCCACAACATTTCCTTGGGTTGAGGGGAATACCTCGACGGTACTCAGACTATCCCGATTGTTATCTG

ATATGAAATAAAATTTCTTCTGCGGGGAGAATTTTAAGTGTTATTTCCGTCATTTATTTTTTATTTATTG

TTTTGGAATCTTTACTCCTTTTGCGGT

>AsiaII_5_Bangladesh_Chapai_AJ748391_AJ748391.1 Bemisia tabaci mitochondrial partial coi gene for cytochrome oxidase I subunit, specific host brinjal, from Bangladesh, Chapai

GAAAACTTGAAGTATTTGGTAGGTTGGGAATAATTTATGCTATAGTAACTATTGGAATTCTAGGTTTTAT

TGTGTGAGGTCATCACATATTTACCGTTGGGATAGATGTTGATACTCGGGCTTATTTTACTTCAGCCACT

ATAATTATCGCTGTTCCGACCGGAATTAAAATCTTTAGGTGGCTTGCTACTCTAGGTGGAATAAAATCTA

ACAAGTTTAGTCCTCTTGGACTTTGGTTTACTGGATTTCTTTTTTTATTTACCATGGGTGGGTTAACTGG

GATCATTCTTGGTAATTCCTCTGTTGATGTCTGCTTACATGATACTTACTTTGTTGTTGCTCATTTTCAT

TATGTTCTATCTATAGGAATTATTTTTGCTATTGTGGGAGGTGTTATTTATTGATTTCCATTAATTTTGG

GCTTAACACTAAATAGTCATAGTTTGGTATCGCAGTTTTACATTATGTTTTTAGGAGTAAATTTAACGTT

TTTTCCACAACATTTCCTTGGGTTGAGGGGAATACCTCGACGGTACTCAGACTATCCCGATTGTTATCTG

ATATGAAATAAAATTTCTTCTGCGGGGAGAATTTTAAGTGTTATTTCCGTCATTTATTTTTTATTTATTG

TTTTGGAATCTTTACTCCTTTTGCGGT

>AsiaII_5_India_New_Delhi_AM408901_AM408901.1 Bemisia tabaci mitochondrial partial COI gene for cytochrome oxidase I, isolated from New Delhi, India

GAAAACTTGAAGTATTTGGTAGGTTGGGAATAATTTATGCTATAGTAACTATTGGAATTCTAGGTTTTAT

TGTGTGAGGTCATCACATATTTACCGTCGGGATAGATGTTGATACTCGGGCTTATTTTACTTCAGCCACT

ATAATTATCGCTGTTCCGACCGGAATTAAAATCTTTAGGTGGCTTGCTACTCTAGGTGGAATAAAATCTA

ACAAGTTTAGTCCTCTTGGACTTTGGTTTACTGGATTTCTTTTTTTATTTACCATGGGTGGGTTAACTGG

GATCATTCTTGGTAATTCCTCTGTTGATGTCTGCTTACATGATACTTACTTTGTTGTTGCTCATTTTCAT

TATGTTCTATCTATAGGAATTATTTTTGCTATTGTGGGAGGTGTTATTTATTGATTTCCATTAATTTTGG

GCTTAACACTAAATAGTCATAGTTTGGTATCGCAGTTTTACATTATGTTTTTAGGAGTAAATTTAACGTT

TTTTCCACAACATTTCCTTGGGTTGAGGGGAATACCTCGACGGTACTCAGACTATCCCGATTGTTATCTG

ATATGAAATAAAATTTCTTCTGCGGGGAGAATTTTAAGTGTTATTTCCGTCATTTATTTTTTATTTATTG

TTTTGGAATCTTTACTCCTTTTGCGGT

>AsiaII_5_Bangladesh_Joydebpur_AJ748393_AJ748393.1 Bemisia tabaci mitochondrial partial coi gene for cytochrome oxidase I subunit, specific host ridge gourd, from Bangladesh, Joydebpur

GAAAACTTGAAGTATTTGGTAGGTTGGGAATAATTTATGCTATAGTAACTATTGGAATTCTAGGTTTTAT

TGTGTGAGGTCATCACATATTTACCGTTGGGATAGATGTTGATACTCGGGCTTATTTTACTTCAGCCACT

ATAATTATCGCTGTTCCGACCGGAATTAAAATCTTTAGGTGGCTTGCTACTCTAGGTGGAATAAAATCTA

ACAAGTTTAGTCCTCTTGGACTTTGGTTTACTGGATTTCTTTTTTTATTTACCATGGGTGGGTTAACTGG

GATCATTCTTGGTAATTCCTCTGTTGATGTCTGCTTACATGATACTTACTTTGTTGTTGCTCATTTTCAT

TATGTTCTATCTATAGGAATTATCTTTGCTATTGTGGGAGGTGTTATTTATTGATTTCCATTAATTTTGG

GCTTAACACTAAATAGTCATAGTTTGGTATCGCAGTTTTACATTATGTTTTTAGGAGTAAATTTAACGTT

TTTTCCACAACATTTCCTTGGGTTGAGGGGAATACCTCGACGGTACTCAGACTATCCCGATTGTTATCTG

ATATGAAATAAAATTTCTTCTGCGGGGAGAATTTTAAGTGTTATTTCCGTCATTTATTTTTTATTTATTG

TTTTGGAATCTTTACTCCTTTTGCGGT

>AsiaII_5_India_AJ748376_AJ748376.1 Bemisia tabaci mitochondrial partial coi gene for cytochrome oxidase I subunit, specific host tomato, from India, Karnataka, Belgaum

GAAAACTTGAAGTATTTGGTAGGTTGGGAATAATTTATGCTATAGTAACTATTGGAATTCTAGGTTTTAT

TGTGTGAGGTCATCACATATTTACCGTTGGGATAGATGTTGATACTCGGGCTTATTTTACTTCAGCCACT

ATAATTATCGCTGTTCCGACCGGAATTAAAATCTTTAGGTGACTTGCTACTCTAGGTGGAATAAAATCTA

ACAAGTTTAGTCCTCTTGGACTTTGGTTTACTGGATTTCTTTTTTTATTTACCATGGGTGGGTTAACTGG

GATCATTCTTGGTAATTCCTCTGTTGATGTCTGCTTACATGATACTTACTTTGTTGTTGCTCATTTTCAT

TATGTTCTATCTATAGGAATTATCTTTGCTATTGTGGGAGGTGTTATTTATTGATTTCCATTAATTTTGG

GCTTAACACTAAATAGTCATAGTTTGGTATCGCAGTTTTACATTATGTTTTTAGGAGTAAATTTAACGTT

TTTTCCACAACATTTCCTTGGGTTGAGGGGAATACCTCGACGGTACTCAGACTATCCCGATTGTTATCTG

ATATGAAATAAAATTTCTTCTGCGGGGAGAATTTTAAGTGTTATTTCCGTCATTTATTTTTTATTTATTG

TTTTGGAATCTTTACTCCTTTTGCGGT

>AsiaII_5_India_AF418666_AF418666.2 Bemisia tabaci cytochrome oxidase subunit I (COI) gene, partial cds; mitochondrial

GAAAACTTGAAGTATTTGGTAGGTTGGGAATAATTTATGCTATAGTAACTATTGGAATTCTAGGTTTTAT

TGTGTGAGGTCATCACATATTTACCGTTGGGATAGATGTTGATACTCGGGCTTATTTTACTTCAGCCACT

ATAATTATCGCTGTTCCGACCGGAATTAAAATCTTTAGGTGACTTGCTACTCTAGGTGGAATAAAATCTA

ACAAGTTTAGTCCGCTTGGACTTTGGTTTACTGGATTTCTTTTTTTATTTACCATGGGTGGGTTAACTGG

GATCATTCTTGGTAATTCCTCTGTTGATGTCTGCTTACATGATACTTACTTTGTTGTTGCTCATTTTCAT

TATGTTCTATCTATAGGAATTATCTTTGCTATTGTGGGAGGTGTTATTTATTGATTTCCATTAATTTTGG

GCTTAACACTAAATAGTCATAGTCTGGTATCGCAGTTTTACATTATGTTTTTAGGAGTAAATTTAACGTT

TTTTCCACAACATTTCCTTGGGTTGAGGGGAATACCTCGACGGTACTCAGACTATCCCGATTGTTATCTG

ATATGAAATAAAATTTCTTCTGCGGGGAGAATTTTAAGTGTTATTTCCGTTATTTATTTTTTATTTATTG

TTTTGGAATCTTTACTTCTTTTGCGGT

>AsiaII_6_Taiwan_DQ174520_DQ174520.1 Bemisia tabaci from Ipomoea acuminata cytochrome oxidase subunit I gene, partial cds; mitochondrial

GAAAACTTGAAGTGTTTGGTAGGTTAGGTATAATTTATGCTATAGTGACTATTGGAATTCTAGGTTTTAT

TGTGTGAGGTCATCATATATTTACTGTTGGGATAGATGTTGATACTCGGGCTTATTTTACTTCAGCTACT

ATGATTATTGCTGTTCCGACCGGAATCAAAATCTTTAGGTGACTTGCTACCTTAGGTGGAATAAAATCTA

ATAGGTTTAGTCCGCTTAGACTTTGGTTTACCGGATTTCTTTTTTTATTTACTATGGGTGGGTTAACTGG

GATCATTCTTGGTAATTCTTCTGTTGATGTTTGTTTACACGACACTTATTTTGTTGTTGCTCATTTTCAT

TATGTTTTATCTATAGGAATTATCTTTGCTATTGTGGGAGGTCTTATCTACTGATTTCCATTAATTTTGG

GCTTAACACTTAATAGTCATAGCCTGGTATCACAGTTTTACATTATGTTTTTGGGAGTAAATTTAACGTT

TTTTCCACAGCATTTCCTTGGGTTGAGGGGAATGCCTCGCCGGTATTCAGACTATCCCGATTGTTATCTG

ATGTGAAATAAAATTTCTTCTGCGGGGAGAATTTTAAGTGTTATTTCTGTAATTTATTTTTTATTTATTG

TCTTGGAATCTTTACTTCTTTTACGGT

>AsiaII_6_Japan_Okinawa_AB308125_AB308125.1 Bemisia tabaci mitochondrial COI gene for cytochrome oxidase subunit I, partial cds, country: Japan:Okinawa, Kurima island

GAAAACTTGAAGTGTTTGGTAGGCTAGGTATAATTTATGCTATAGTGACTATTGGAATTCTAGGTTTTAT

TGTGTGAGGTCATCATATATTTACTGTTGGGATAGATGTTGATACTCGGGCTTATTTTACTTCAGCTACT

ATGATTATTGCTGTTCCGACCGGAATCAAAATCTTTAGGTGACTTGCTACCTTAGGTGGAATAAAATCTA

ATAGGTTTAGTCCGCTTAGACTTTGGTTTACCGGATTTCTTTTTTTATTTACTATGGGTGGGTTAACTGG

GATCATTCTTGGTAATTCTTCTGTTGATGTTTGTTTACACGACACTTATTTTGTTGTTGCTCATTTTCAT

TATGTTTTATCCATAGGAATTATCTTTGCTATTGTGGGAGGTCTTATTTACTGATTTCCATTAATTTTGG

GCTTAACACTTAATAGTCATAGCCTGGTATCACAGTTTTACATTATGTTTTTGGGAGTAAATTTAACGTT

TTTTCCACAGCATTTCCTTGGGTTGAGGGGAATGCCTCGCCGGTATTCAGACTATCCCGATTGTTATCTG

ATGTGAAATAAAATTTCTTCTGCGGGGAGAATTTTAAGTGTTATTTCTGTAATTTATTTTTTATTTATTG

TCTTGGAATCTTTACTTCTTTTACGGT

>AsiaII_6_Japan_Okinawa_AB308126_AB308126.1 Bemisia tabaci mitochondrial COI gene for cytochrome oxidase subunit I, partial cds, country: Japan:Okinawa, Ishigaki island

GAAAACTTGAAGTGTTTGGTAGGCTAGGTATAATTTATGCTATAGTGACTATTGGAATTCTAGGTTTTAT

TGTGTGAGGTCATCATATATTTACTGTTGGGATAGATGTTGATACTCGGGCTTATTTTACTTCAGCTACT

ATGATTATTGCTGTTCCGACCGGAATCAAAATCTTTAGGTGACTTGCTACCTTAGGTGGAATAAAATCTA

ATAGGTTTAGTCCGCTTAGACTTTGGTTTACCGGATTTCTTTTTTTATTTACTATGGGTGGGTTAACTGG

GATCATTCTTGGTAATTCTTCTGTTGATGTTTGTTTACACGACACTTATTTTGTTGTTGCTCATTTTCAT

TATGTTTTATCTATAGGAATTATCTTTGCTATTGTGGGAGGTCTTATTTACTGATTTCCATTAATTTTGG

GCTTAACACTTAATAGTCATAGCCTGGTATCACAGTTTTACATTATGTTTTTGGGAGTAAATTTAACGTT

TTTTCCACAGCATTTCCTTGGGTTGAGGGGAATGCCTCGCCGGTATTCAGACTATCCCGATTGTTATCTG

ATGTGAAATAAAATTTCTTCTGCGGGGAGAATTTTAAGTGTTATTTCTGTAATTTATTTTTTATTTATTG

TCTTGGAATCTTTACTTCTTTTACGGT

>AsiaII_6_Japan_Kagoshima_AB440788_AB440788.1 Bemisia tabaci mitochondrial COI gene for cytochrome oxidase subunit I, partial cds, isolate: Yoron-2008-Wb

GAAAACTTGAAGTGTTTGGCAGGTTAGGTATAATTTATGCTATAGTGACTATTGGAATTCTAGGTTTTAT

TGTGTGAGGTCATCATATATTTACTGTTGGGATAGATGTTGATACTCGGGCTTATTTTACTTCAGCTACT

ATGATTATTGCTGTTCCGACCGGAATCAAAATCTTTAGGTGACTTGCTACTTTAGGTGGAATAAAATCTA

ACAGGTTTAGTCCGCTTAGACTTTGGTTTACCGGATTTCTTTTTTTATTTACTATGGGTGGGTTAACTGG

GATCATTCTTGGCAATTCTTCTGTTGATGTCTGTTTACACGACACTTATTTTGTTGTTGCTCATTTTCAT

TATGTTTTATCTATAGGAATTATCTTTGCTATTGTGGGAGGTCTTATCTACTGATTTCCATTAATTTTGG

GCTTAACGCTTAATAGTCATAGCCTGGTATCACAGTTTTACATTATGTTTTTGGGAGTAAATTTAACGTT

TTTTCCACAGCATTTCCTTGGGTTGAGGGGAATGCCTCGCCGGTATTCAGACTATCCCGACTGTTATCTG

ATGTGAAATAAAATTTCTTCTGCGGGGAGAATTTTAAGTGTTATTTCTGTAATTTATTTTTTATTTATTG

TCTTGGAATCTTTACTTCTTTTACGGT

>AsiaII_6_Japan_Okinawa_AB308124_AB308124.1 Bemisia tabaci mitochondrial COI gene for cytochrome oxidase subunit I, partial cds, country: Japan:Okinawa, Yomitan

GAAAACTTGAAGTGTTTGGCAGGTTAGGTATAATTTATGCTATAGTGACTATTGGAATTCTAGGTTTTAT

TGTGTGAGGTCATCATATATTTACTGTTGGGATAGATGTTGATACTCGGGCTTATTTTACTTCAGCTACT

ATGATTATTGCTGTTCCGACCGGAATCAAAATCTTTAGGTGACTTGCTACTTTAGGTGGAATAAAATCTA

ACAGGTTTAGTCCGCTTAGACTTTGGTTTACCGGATTTCTTTTTTTATTTACTATGGGTGGGTTAACTGG

GATCATTCTTGGCAATTCTTCTGTTGATGTCTGTTTACACGACACTTATTTTGTTGTTGCTCATTTTCAT

TATGTTTTATCTATAGGAATTATCTTTGCTATTGTGGGAGGTCTTATCTACTGATTTCCATTAATTTTGG

GCTTAACACTTAATAGTCATAGCCTGGTATCACAGTTTTACATTATGTTTTTGGGAGTAAATTTAACGTT

TTTTCCACAGCATTTCCTTGGGTTGAGGGGAATGCCTCGCCGGTATTCAGACTATCCCGACTGTTATCTG

ATGTGAAATAAAATTTCTTCTGCGGGGAGAATTTTAAGTGTTATTTCTGTAATTTATTTTTTATTTATTG

TCTTGGAATCTTTACTTCTTTTACGGT

>AsiaII_3_China_AJ783706_AJ783706.2 Bemisia tabaci mitochondrial partial coi gene for cytochrome oxidase I, exon 1

GAAAGCTTGAAGTATTTGGAAGGCTGGGTATAATTTATGCTATAGTAACTATTGGCATTTTAGGTTTTAT

TGTCTGAGGTCATCATATATTTACTGTTGGGATAGATGTAGATACCCGAGCTTATTTTACCTCAGCTACC

ATAATTATTGCTGTTCCAACTGGAATTAAAATCTTTAGGTGACTTGCTACCCTAGGTGGAATAAAGTCTA

ACAAGCTCAGTCCGGTTGTTCTCTGATTTACTGGATTTTTGTTTTTATTTACTATGGGCGGGTTAACTGG

AATTATTCTTGGTAACTCTTCTGTCGATGTCTGCTTACACGATACTTATTTTGTTGTTGCTCACTTTCAT

TATGTTTTGTCTATAGGAATTATTTTTGCTATTGTGGGTGGGGCTATCTATTGATTTCCATTAATTTTAG

GATTAACATTGAATAGTCATAGTCTGGTATCTCAATTTTACGTCATATTTCTGGGAGTAAATTTAACATT

TTTTCCACAACATTTTCTTGGGCTAAGGGGGATACCTCGCCGGTATTCAGACTATCCTGATTGTTATTTG

ATATGAAATAAAATTTCTTCTGCAGGGAGAATTTTGAGTATTATTTCTGTTATTTATTTTTTATTTATCG

TTTTAGAGTCTTTACTTCTTTTGCGAT

>AsiaII_3_China_Zhejiang_DQ309074_DQ309074.1 Bemisia tabaci clone 1 haplotype ZHJ-1 cytochrome oxidase subunit 1 (COI) gene, partial cds; mitochondrial

GAAAGCTTGAAGTATTTGGAAGGCTGGGTATAATTTATGCTATAGTAACTATTGGCATTTTAGGTTTTAT

TGTCTGAGGTCATCATATATTTACTGTTGGGATAGATGTAGATACCCGAGCTTATTTTACCTCAGCTACC

ATAATTATTGCTGTTCCAACTGGAATTAAAATCTTTAGGTGACTTGCTACCCTAGGTGGAATAAAGTCTA

ACAAGCTCAGTCCGGTTGTTCTCTGATTTACTGGATTTTTGTTTTTATTTACTATGGGCGGGTTAACTGG

AATTATTCTTGGTAACTCTTCTGTCGATGTCTGCTTACACGATACTTATTTTGTTGTTGCTCACTTTCAT

TATGTTTTGTCTATAGGAATTATTTTTGCTATTGTGGGTGGGGCTATCTATTGATTTCCATTAATTTTAG

GATTAACATTGAATAGTCATAGTCTGGTATCTCAATTTTACGTCATATTTCTGGGAGTAAATTTAACATT

TTTTCCACAACATTTTCTTGGGCTAAGGGGGATACCTCGCCGGTATTCAGACTATCCTGATTGTTATTTG

ATATGAAATAAAATTTCTTCTGCAGGGAGAATTTTGAGTATTATTTCTGTTATTTATTTTTTATTTATCG

TTTTAGAGTCTTTACTTCTTTTGCGAT

>AsiaII_8_India_AJ748357_AJ748357.1 Bemisia tabaci mitochondrial partial coi gene for cytochrome oxidase I subunit, specific host Acanthospermum hispidium, from India, Karnataka, Belgaum

GAAAACTTGAAGTATTTGGAAGACTAGGGATAATTTATGCTATATTAACTATTGGCATTTTAGGTTTTAT

TGTTTGAGGTCATCATATATTTACTGTTGGGATAGATGTTGATACTCGGGCCTATTTTACCTCAGCTACT

ATAGTTATCGCTGTTCCAACTGGAATTAAAATCTTTAGTTGGCTTGCTACTTTGGGGGGAATAAAATCTA

ACAAGTTTAGTCCTCTTGTTCTTTGATTTACTGGATTTCTGTTTCTATTTACTATGGGCGGACTAACTGG

GATTATTCTTGGTAATTCTTCTGTTGATGTCTGTTTACATGACACTTATTTTGTTGTTGCTCATTTTCAT

TATGTTCTGTCTATGGGAATTATTTTTGCTATTGTAGGAGGGGCTATTTACTGATTTCCATTAATCTTGG

GCTTAACACTAAATAGTCACAGCTTGGTGTCTCAGTTTTATATGATGTTTATGGGAGTAAATTTAACGTT

TTTTCCACAACATTTTCTTGGACTAAGGGGGATACCTCGTCGGTATTCAGATTACCCTGATTGTTATTTG

ATATGAAATAAAATTTCTTCTGCGGGTAGAATTTTGAGTATTATTTCTGTTATTTATTTTTTATTTATTG

TTTTAGAGTCTTTCCTCCTTCTGCGAT

>AsiaII_8_India_Uttarahand_HM590188_HM590188.2 Bemisia tabaci isolate SAUNI-DDS12 cytochrome oxidase subunit 1 gene, partial cds; mitochondrial

GAAAACTTGAAGTATTTGGAAGACTAGGGATAATTTATGCTATATTAACTATTGGCATTTTAGGTTTTAT

TGTTTGAGGTCATCATATATTTACTGTTGGGATAGATGTTGATACTCGGGCCTATTTTACCTCAGCTACT

ATAGTTATCGCTGTTCCAACTGGAATTAAAATCTTTAGTTGGCTTGCTACTTTGGGGGGAATAAAATCTA

ACAAGTTTAGTCCTCTTGTTCTTTGATTTACTGGATTTCTGTTTCTATTTACTATGGGCGGACTAACTGG

GATTATTCTTGGTAATTCTTCTGTTGATGTCTGTTTACATGACACTTATTTTGTTGTTGCTCATTTTCAT

TATGTTCTGTCTATGGGAATTATTTTTGCTATTGTAGGAGGGGCTATTTACTGATTTCCATTAATCTTGG

GCTTAACACTAAATAGTCACAGCTTGGTGTCTCAGTTTTATATGATGTTTATGGGAGTAAATTTAACGTT

TTTTCCACAACATTTTCTTGGGCTAAGGGGGATACCTCGTCGGTATTCAGATTACCCTGATTGTTATTTG

ATATGAAATAAAATTTCTTCTGCGGGTAGAATTTTGAGTATTATTTCTGTTATTTATTTTTTATTTATTG

TTTTAGAGTCTTTCCTCCTTCTGCGAT

>AsiaII_8_India_AJ748362_AJ748362.1 Bemisia tabaci mitochondrial partial coi gene for cytochrome oxidase I subunit, specific host cluster bean, from India, Karnataka, Dharwad, Annebavi, Tumbadi

GAAAACTTGAAGTATTTGGAAGACTAGGGATAATTTATGCTATATTAACTATTGGCATTTTAGGTTTTAT

TGTTTGAGGTCATCATATATTTACTGTTGGGATAGATGTTGATACTCGGGCCTATTTTACCTCAGCTACT

ATAGTTATCGCTGTTCCAACTGGAATTAAAATCTTTAGTTGGCTTGCTACTTTGGGGGGAATAAAATCTA

ACAAGTTTAGTCCTCTTGTTCTTTGATTTACTGGATTTCTGTTTCTATTTACCATGGGCGGACTAACTGG

GATTATTCTTGGTAATTCTTCTGTTGATGTCTGTTTACATGACACTTATTTTGTTGTTGCTCATTTTCAT

TATGTTCTGTCTATGGGAATTATTTTTGCTATTGTAGGAGGGGCTATTTACTGATTTCCATTAATCTTGG

GCTTAACACTAAATAGTCACAGCTTGGTGTCTCAGTTTTATATGATGTTTATGGGAGTAAATTTAACGTT

TTTTCCACAACATTTTCTTGGGCTAAGGGGGATACCTCGTCGGTATTCAGATTACCCTGATTGTTATTTG

ATATGAAATAAAATTTCTTCTGCGGGTAGAATTTTGAGTATTATTTCTGTTATTTATTTTTTATTTATTG

TTTTAGAGTCTTTCCTCCTTCTGCGAT

>Italy1_Sicily_AY827601_AY827601.1-1 Bemisia tabaci voucher IMIDA-It2.3 cytochrome oxidase subunit I (COI) gene, partial cds; mitochondrial

GAAAACTTGAGGTGTTTGGAAGGCTGGGTATAATCTATGCTATAATAACTATTGGCATTTTAGGTTTTAT

TGTTTGGGGTCATCATATATTTACTGTTGGGATAGATGTTGATACTCGGGCTTATTTTACTTCAGCTACA

ATAATTATTGCTGTTCCTACGGGAATTAAAATTTTTAGGTGGCTTGCCACCTTAGGTGGGATAAAGTCTA

ATAAGTTTAGTCCTCTTGTTCTTTGATTTACAGGATTTTTATTTTTATTTACTATGGGTGGATTAACTGG

AATTATTCTTGGTAATTCTTCTGTTGATGTCTGTTTACACGATACTTATTTTGTTGTCGCTCATTTTCAT

TATGTTTTATCTATAGGAATTATTTTTGCTATTATAGGAGGGATTATTTATTGATTTCCACTAATTTTGG

GCATAACACTAAATGGTCATAATCTGGTATCTCAGTTTTATATTATGTTTATGGGAGTAAATTTAACATT

TTTCCCACAGCATTTTCTCGGGTTAAGAGGTATACCTCGCCGGTATTCAGATTATCCAGATTGTTATTTG

ATGTGAAATAAAATTTCTTCTGCGGGAAGAATCTTGAGTATTATTTCTGTTATTTATTTTTTATTTATTG

TTCTAGAATCTTTTCTTCTCTTGCGGT

>Italy1_Sicily_AY827598_AY827598.1-1 Bemisia tabaci voucher IMIDA-It1.4 cytochrome oxidase subunit I (COI) gene, partial cds; mitochondrial

GAAAACTTGAGGTGTTTGGAAGGCTGGGTATAATCTATGCTATAATAACTATTGGCATTTTAGGTTTTAT

TGTTTGGGGTCATCATATATTTACTGTTGGGATAGATGTTGATACTCGGGCTTATTTTACTTCAGCTACA

ATAATTATTGCTGTTCCTACGGGAATTAAAATTTTTAGGTGGCTTGCCACCTTAGGTGGGATAAAGTCTA

ATAAGTTTAGTCCTCTTGTTCTTTGATTTACAGGATTTTTATTTTTATTTACTATGGGTGGATTAACTGG

AATTATTCTTGGTAATTCTTCTGTTGATGTCTGTTTACACGATACTTATTTTGTTGTCGCTCATTTTCAT

TATGTTTTATCTATAGGAATTATTTTTGCTATTATAGGAGGGATTATTTATTGATTTCCACTAATTTTGG

GCATAACACTAAATGGTCATAATCTGGTATCTCAGTTTTATATTATGTTTATGGGAGTAAATTTAACATT

TTTCCCACAGCATTTTCTCGGGTTAAGAGGTATACCTCGCCGGTATTCAGATTATCCAGATTGTTATTTG

GTGTGAAATAAAATTTCTTCTGCGGGAAGAATCTTGAGTATTATTTCTGTTATTTATTTTTTATTTATTG

TTCTAGAATCTTTTCTTCTCCTGCGGT

>Italy1_Sicily_AY827600_AY827600.1-1 Bemisia tabaci voucher IMIDA-It2.2 cytochrome oxidase subunit I (COI) gene, partial cds; mitochondrial

GAAAACTTGAGGTGTTTGGAAGGCTGGGTATAATCTATGCTATAATAACTATTGGCATTTTAGGTTTTAT

TGTTTGGGGTCATCATATATTTACTGTTGGGATAGATGTTGATACTCGCGCTTATTTTACTTCAGCTACA

ATAATTATTGCTGTTCCTACGGGAATTAAAATTTTTAGGTGGCTTGCCACCTTAGGTGGGATAAAGTCTA

ATAAGTTTAGTCCTCTTGTTCTTTGATTTACAGGATTTTTATTTTTATTTACTATGGGTGGATTAACTGG

AATTATTCTTGGTAATTCTTCTGTTGATGTCTGTTTACACGATACTTATTTTGTTGTCGCTCATTTTCAT

TATGTTTTATCTATAGGAATTATTTTTGCTATTATAGGAGGGATTATTTATTGATTTCCACTAATTTTGG

GCATAACACTAAATGGTCATAATCTGGTATCTCAGTTTTATATTATGTTTATGGGAGTAAATTTAACATT

TTTCCCACAGCATTTTCTCGGGTTAAGAGGTATACCTCGCCGGTATTCAGATTATCCAGATTGTTATTTG

ATGTGAAATAAAATTTCTTCTGCGGGAAGAATCTTGAGTATTATTTCTGTTATTTATTTTTTATTTATTG

TTCTAGAATCTTTTCTTCTCTTGCGGT

>IndianOcean_MS_Madagascar_AJ550171_AJ550171.1 Bemisia tabaci mitochondrial partial coi gene for cytochrome oxidase subunit 1, isolate 20-0

GGAAATTGGAGGTATTTGGAAGGTTGGGTATAATTTATGCTATATTAACTATTGGCATCTTGGGGTTTAT

TGTTTGAGGTCACCATATATTTACAGTTGGAATAGATGTAGATACTCGAGCTTATTTTACCTCAGCTACT

ATGATTATTGCTGTTCCTACAGGAATTAAAATTTTCAGTTGGCTTGCTACTTTGGGTGGAATAAAGTCTA

ATAAATTCAGGCCTCTTGGCCTTTGGTTCACAGGATTTTTATTTTTATTTACTATGGGTGGATTAACTGG

AATTATTCTTGGTAACTCTTCTGTAGATGTATGTTTGCACGATACTTATTTTGTTGTTGCACATTTTCAT

TATGTTTTATCAATAGGAATCATTTTTGCTATTGTAGGAGGAGTTATTTATTGATTTCCATTAATCTTGG

GTTTAAGTTTAAATAATTATAGTTTGGTATCTCAATTTTATATTATGTTCATTGGAGTAAATTTAACTTT

TTTTCCTCAGCACTTTCTTGGTTTAGGGGGAATGCCTCGACGATATTCAGACTATGCTGACTGCTATCTA

GTATGAAACAAAATTTCCTCTGCGGGAAGAATTTTGAGTATCATTTCTGTTATTTATTTTTTATTTATTG

TTTTAGAATCTTTCCTTCTTCTGCGGT

>IndianOcean_Uganda_Busukuma_AY903538_AY903538.1 Bemisia tabaci clone UgdfBk13 cytochrome oxidase subunit I-like gene, complete sequence; mitochondrial

GGAAATTGGAGGTATTTGGAAGGTTGGGTATAATTTATGCTATATTAACTATTGGCATCTTGGGGTTTAT

TGTTTGAGGTCACCATATATTTACAGTTGGAATAGATGTAGATACTCGAGCTTATTTTACCTCAGCTACT

ATGATTATTGCTGTTCCTACAGGAATTAAAATTTTCAGTTGGCTTGCTACTTTGGGTGGAATAAAGTCTA

ATAAATTCAGGCCTCTTGGCCTTTGGTTCACAGGATTTTTATTTTTATTTACTATGGGTGGATTAACTGG

AATTATTCTTGGTAACTCTTCTGTAGATGTATGTTTGCACGATACTTATTTTGTTGTTGCACATTTTCAT

TATGTTTTATCAATAGGAATCATTTTTGCTATTGTAGGAGGAGTTATTTATTGATTTCCATTAATCTTGG

GTTTAAGTTTAAATAATTATAGTTTGGTATCTCAATTTTATATTATGTTCATTGGAGTAAATTTAACTTT

TTTTCCTCAGCACTTTCTTGGTTTAGGGGGAATGCCTCGACGATATTCAGACTACGCTGACTGCTATCTA

GTATGAAACAAAATTTCCTCTGCGGGAAGAATTTTGAGTATCATTTCTGTTATTTATTTTTTATTTATTG

TTTTAGAATCTTTCCTTCTTCTGCGGT

>IndianOcean_Uganda_Masindi_AY903522_AY903522.1 Bemisia tabaci clone UgBnMsc2 cytochrome oxidase subunit I-like gene, complete sequence; mitochondrial

GGAAATTGGAGGTATTTGGAAGGTTGGGTATAATTTATGCTATATTAACTATTGGCATCTTGGGGTTTAT

TGTTTGAGGTCACCATATATTTACAGTTGGAATAGATGTAGATACTCGAGCTTATTTTACCTCAGCTACT

ATGATTATTGCTGTTCCTACAGGAATTAAAATTTTCAGTTGGCTTGCTACTTTGGGTGGAATAAAGTCTA

ATAAATTCAGGCCTCTTGGCCTTTGGTTCACAGGATTTTTATTTTTATTTACTATGGGTGGATTAACTGG

AATTATTCTTGGTAACTCTTCTGTAGATGTATGTTTGCACGATACTTATTTTGTTGTTGCACATTTTCAT

TATGTTTTATCAATAGGAATCATTTTTGCTATTGTAGGAGGAGTTATTTATTGATTTCCATTAATCTTGG

GTTTAAGTTTAAATAATTATAGTTTGGTATCTCAATTTTATATTATGTTCATTGGAGTAAATTTAACTTT

TTTTCCTCAGCACTTTCTTGGTTTAGGGGGAATGCCTCGACGATATTCAGACTATGCTGACTGCTATCTA

GTATGAAACAAAATTTCCTCTGCGGGAAGAATTCTGAGTATCATTTCTGTTATTTATTTTTTATTTATTG

TTTTAGAATCTTTCCTTCTTCTGCGGT

>IndianOcean_Uganda_Busukuma_AY903539_AY903539.1 Bemisia tabaci clone UgDfBk77 cytochrome oxidase subunit I-like gene, complete sequence; mitochondrial

GGAAATTGGAGGTATTTGGAAGGTTGGGTATAATTTATGCTATATTAACTATTGGCATCTTGGGGTTTAT

TGTTTGAGGTCACCATATATTTACAGTTGGAATAGATGTAGATACTCGAGCTTATTTTACCTCAGCTACT

ATGATTATTGCTGTTCCTACAGGAATTAAAATTTTCAGTTGGCTTGCTACTTTGGGTGGAATAAAGTCTA

ATAAATTCAGGCCTCTTGGCCTTTGGTTCACAGGATTTTTATTTTTATTTACTATGGGTGGATTAACTGG

AATTATTCTTGGTAACTCTTCTGTAGATGTATGTTTGCACGATACTTATTTTGTTGTTGCACATTTTCAT

TATGTTTTATCAATAGGAATCATTTTTGCTATTGTAGGAGGAGTTATTTATTGATTTCCATTAATCTTGG

GTTTAAGTTTAAATAATTATAGTTTGGTATCTCAATTTTATATTATGTTCATTGGAGTAAATTTAACTTT

TTTTCCTCAGCACTTTCTTGGTTTAGGGGGAATGCCTCGACGATATTCAGACTATGCTGACTGCTATCTA

GTATGAAACAAAATTTCCTCTGCGGGAAGAATTTTGAGTATCATTTCTGTTATTTATTTTTTATTTATTG

TTTTAGAATCTTTCCTCCTTCTGCGGT

>IndianOcean_Uganda_Busukuma_AY903537_AY903537.1 Bemisia tabaci clone UgdfBk12 cytochrome oxidase subunit I-like gene, complete sequence; mitochondrial

GGAAATTGGAGGTATTTGGAAGGTTGGGTATAATTTATGCTATATTAACTATTGGCATCTTGGGGTTTAT

TGTCTGAGGTCACCATATATTTACAGTTGGAATAGATGTAGATACTCGAGCTTATTTTACCTCAGCTACT

ATGATTATTGCTGTTCCTACAGGAATTAAAATTTTCAGTTGGCTTGCTACTTTGGGTGGAATAAAGTCTA

ATAAATTCAGGCCTCTTGGCCTTTGGTTCACAGGATTTTTATTTTTATTTACTATGGGTGGATTAACTGG

AATTATTCTTGGTAACTCTTCTGTAGATGTATGTTTGCACGATACTTATTTTGTTGTTGCACATTTTCAT

TATGTTTTATCAATAGGAATCATTTTTGCTATTGTAGGAGGAGTTATTTATTGATTTCCATTAATCTTGG

GTTTAAGTTTAAATAATTATAGTTTGGTATCTCAATTTTATATTATGTTCATTGGAGTAAATTTAACTTT

TTTTCCTCAGCACTTTCTTGGTTTAGGGGGAATGCCTCGACGATATTCAGACTATGCTGACTGCTATCTA

GTATGAAACAAAATTTCCTCTGCGGGAAGAATTTTGAGTATCATTTCTGTTATTTATTTTTTATTTATTG

TTTTAGAATCTTTCCTTCTTCTGCGGT

>IndianOcean_Seychelles_AJ550182_AJ550182.1 Bemisia tabaci mitochondrial partial coi gene for cytochrome oxidase subunit 1, isolate 23-0

GGAAATTGGAGGTATTTGGAAGGTTGGGTATAATTTATGCTATATTAACTATTGGCATTTTGGGGTTTAT

TGTTTGAGGTCACCATATATTTACAGTTGGAATAGATGTAGATACTCGAGCTTATTTTACCTCAGCTACT

ATGATTATTGCTGTTCCTACAGGAATTAAAATTTTCAGTTGGCTTGCTACTTTGGGTGGAATAAAGTCTA

ATAAATTCAGGCCTCTTGGCCTTTGGTTCACAGGATTTTTATTTTTATTTACTATGGGTGGATTAACTGG

AATTATTCTTGGTAACTCTTCTGTAGATGTATGTTTACACGATACTTATTTTGTTGTTGCACATTTCCAT

TATGTTTTATCAATAGGAATCATTTTTGCTATTGTAGGAGGAGTTATTTATTGATTTCCATTAATCTTGG

GTTTAAGTTTAAATAATTATAGTTTGGTATCTCAATTTTATATTATGTTCATTGGAGTAAATTTAACTTT

TTTTCCTCAGCACTTTCTTGGTTTAGGGGGAATGCCTCGACGATATTCAGACTATGCTGACTGCTATCTA

GTATGAAACAAAATTTCCTCTGCGGGAAGAATTTTGAGTATTATTTCTGTTATTTATTTTTTATTTATTG

TTTTAGAATCTTTCCTTCTTCTGCGGT

>MidEastAm1_Australia_Emerald_HM070414_HM070414.1

GAAAATTAGAGGTATTTGGAAGGTTGGGTATAATTTATGCTATATTGACTATTGGTATTCTAGGGTTTAT

TGTTTGAGGTCATCATATATTCACAGTTGGAATAGATGTAGATACTCGAGCTTATTTCACTTCAGCCACT

ATAATTATTGCTGTTCCCACAGGAATTAAAATTTTTAGTTGGCTTGCTACTTTGGGTGGAATAAAGTCTA

ATAAATTAAGGCCTCTTGGCCTTTGATTTACAGGATTTTTATTTTTATTTACTATAGGTGGGTTAACTGG

AATTATTCTTGGTAATTCTTCTGTAGATGTGTGTCTGCATGACACTTATTTTGTTGTTGCACATTTTCAT

TATGTTTTATCAATAGGAATTATTTTTGCTATTGTAGGAGGAGTTATCTATTGATTTCCACTAATTTTAG

GTTTAACCTTAAATAATTATAGATTGGTGTCTCAATTTTATATCATGTTTATTGGAGTAAATTTAACTTT

TTTTCCTCAGCATTTTCTTGGTTTAGGGGGAATGCCTCGTCGATATTCAGATTATGCTGATTGCTATCTA

GTATGAAATAAAATTTCTTCTGCGGGAAGGATTCTGAGTATTATTTCTGTTATTTATTTTTTATTTATTG

TTTTAGAATCCTTTCTTCTTCTGCGGT

>MidEastAm1_Brazil_Capao_Bonito_JN689356_JN689356.1

GAAAATTAGAGGTATTTGGAAGGTTGGGTATAATTTATGCTATATTGACTATTGGTATTCTAGGGTTTAT

TGTTTGAGGTCATCATATATTCACAGTTGGAATAGATGTAGATACTCGAGCTTATTTCACTTCAGCCACT

ATAATTATTGCTGTTCCCACAGGAATTAAAATTTTTAGTTGGCTTGCTACTTTGGGTGGAATAAAGTCTA

ATAAATTAAGGCCTCTTGGCCTTTGATTTACAGGATTTTTATTTTTATTTACTATAGGTGGGTTAACTGG

AATTATTCTTGGTAATTCTTCTGTAGATGTGTGTCTGCATGACACTTATTTTGTTGTTGCACATTTTCAT

TATGTTTTATCAATAGGAATTATTTTTGCTATTGTAGGAGGAGTTATCTATTGATTTCCACTAATCTTAG

GTTTAACCTTAAATAATTATAGATTGGTGTCTCAATTTTATATCATGTTTATTGGAGTAAATTTAACTTT

TTTTCCTCAGCATTTTCTTGGTTTAGGGGGAATGCCTCGTCGATATTCAGATTATGCTGATTGCTATCTA

GTATGAAATAAAATTTCTTCTGCGGGAAGGATTCTGAGTATTATTTCTGTTATTTATTTTTTATTTATTG

TTTTAGAATCCTTTCTTCTTCTGCGGT

>MidEastAm1_Japan_Ehime_Hap1_AB204577_AB204577.1

GAAAATTAGAGGTATTTGGAAGGTTGGGTATAATTTATGCTATATTGACTATTGGTATTCTAGGGTTTAT

TGTTTGAGGTCATCATATATTCACAGTTGGAATAGATGTAGATACTCGAGCTTATTTCACTTCAGCCACT

ATAATTATTGCTGTTCCCACAGGAATTAAAATTTTTAGTTGGCTTGCTACTTTGGGTGGAATAAAGTCTA

ATAAATTAAGGCCTCTTGGCCTTTGATTTACAGGATTTTTATTTTTATTTACTATAGGTGGGTTAACTGG

AATTATTCTTGGTAATTCTTCTGTAGATGTGTGTCTGCATGACACTTATTTTGTTGTTGCACATTTTCAT

TATGTTTTATCAATAGGAATTATTTTTGCTATTGTAGGAGGAGTTATCTATTGATTTCCACTAATCTTAG

GTTTAACCTTAAATAATTATAGATTGGTGTCTCAATTTTATATCATGTTTATTGGAGTAAATTTAACTTT

TTTTCCTCAGCATTTTCTTGGTTTAGGGGGAATGCCTCGTCGATATTCAGATTATGCTGATTGCTATCTA

GTATGAAATAAAATTTCTTCTGCGGGAAGGATTCTGAGTATTATTTCTGTTATTTATTTTTTATTTATTG

TTTTAGAATCCTTTCTTCTTCTGCGGT

>MidEastAm1_USA_Arizona_HM070411_HM070411.1

GAAAATTAGAGGTATTTGGAAGGTTGGGTATAATTTATGCTATATTGACTATTGGTATTCTAGGGTTTAT

TGTTTGAGGTCATCATATATTCACAGTTGGAATAGATGTAGATACTCGAGCTTATTTCACTTCAGCCACT

ATAATTATTGCTGTTCCCACAGGAATTAAAATTTTTAGTTGGCTTGCTACTTTGGGTGGAATAAAGTCTA

ATAAATTAAGGCCTCTTGGCCTTTGATTTACAGGATTTTTATTTTTATTTACTATAGGTGGGTTAACTGG

AATTATTCTTGGTAATTCTTCTGTAGATGTGTGTCTGCATGACACTTATTTTGTTGTTGCACATTTTCAT

TATGTTTTATCAATAGGAATTATTTTTGCTATTGTAGGAGGAGTTATCTATTGATTTCCACTAATCTTAG

GTTTAACCTTAAATAATTATAGATTGGTGTCTCAATTTTATATCATGTTTATTGGAGTAAATTTAACTTT

TTTTCCTCAGCATTTTCTTGGTTTAGGGGGAATGCCTCGTCGATATTCAGATTATGCTGATTGCTATCTA

GTATGAAATAAAATTTCTTCTGCGGGAAGGATTCTGAGTATTATTTCTGTTATTTATTTTTTATTTATTG

TTTTAGAATCCTTTCTTCTTCTGCGGT

>MidEastAm1_USA_Florida_GU086340_GU086340.1

GAAAATTAGAGGTATTTGGAAGGTTGGGTATAATTTATGCTATATTGACTATTGGTATTCTAGGGTTTAT

TGTTTGAGGTCATCATATATTCACAGTTGGAATAGATGTAGATACTCGAGCTTATTTCACTTCAGCCACT

ATAATTATTGCTGTTCCCACAGGAATTAAAATTTTTAGTTGGCTTGCTACTTTGGGTGGAATAAAGTCTA

ATAAATTAAGGCCTCTTGGCCTTTGATTTACAGGATTTTTATTTTTATTTACTATAGGTGGGTTAACTGG

AATTATTCTTGGTAATTCTTCTGTAGATGTGTGTCTGCATGACACTTATTTTGTTGTTGCACATTTTCAT

TATGTTTTATCAATAGGAATTATTTTTGCTATTGTAGGAGGAGTTATCTATTGATTTCCACTAATCTTAG

GTTTAACCTTAAATAATTATAGATTGGTGTCTCAATTTTATATCATGTTTATTGGAGTAAATTTAACTTT

TTTTCCTCAGCATTTTCTTGGTTTAGGGGGAATGCCTCGTCGATATTCAGATTATGCTGATTGCTATCTA

GTATGAAATAAAATTTCTTCTGCGGGAAGGATTCTGAGTATTATTTCTGTTATTTATTTTTTATTTATTG

TTTTAGAATCCTTTCTTCTTCTGCGGT

>MidEastAm1_Egypt_DQ133373_DQ133373.1

GAAAATTAGAGGTATTTGGAAGGTTGGGTATAATTTATGCTATATTGACTATTGGTATTCTAGGGTTTAT

TGTTTGAGGTCATCATATATTCACAGTTGGAATAGATGTAGATACTCGAGCTTATTTCACTTCAGCCACT

ATAATTATTGCTGTTCCCACAGGAATTAAAATTTTTAGTTGGCTTGCTACTTTGGGTGGAATAAAGTCTA

ATAAATTAAGGCCTCTTGGCCTTTGATTTACAGGATTTTTATTTTTATTTACTATAGGTGGGTTAACTGG

AATTATTCTTGGTAATTCTTCTGTAGATGTGTGTCTGCATGACACTTATTTTGTTGTTGCACATTTTCAT

TATGTTTTATCAATAGGAATTATTTTTGCTATTGTAGGAGGAGTTATCTATTGATTTCCACTAATCTTAG

GTTTAACCTTAAATAATTATAGATTGGTGTCTCAATTTTATATCATGTTTATTGGAGTAAATTTAACTTT

TTTTCCTCAGCATTTTCTTGGTTTAGGGGGAATGCCTCGTCGATATTCAGATTATGCTGATTGCTATCTA

GTATGAAATAAAATTTCTTCTGCGGGAAGGATTCTGAGTATTATTTCTGTTATTTATTYTTTATTTATTG

TTTTAGAATCCTTTCTTCTTCTGCGGT

>MidEastAm1_Taiwan_Taidung_DQ174530_DQ174530.1

GAAAATTAGAGGTATTTGGAAGGTTGGGTATAATTTATGCTATATTGACTATTGGTATTCTAGGGTTTAT

TGTTTGAGGTCATCATATATTCACAGTTGGAATAGATGTAGATACTCGAGCTTATTTCACTTCAGCCACT

ATAATTATTGCTGTTCCCACAGGAATTAAAATTTTTAGTTGGCTTGCTACTTTGGGTGGAATAAAGTCTA

ATAAATTAAGGCCTCTTGGCCTTTGATTTACAGGATTTTTATTTTTATTTACTATAGGTGGGTTAACTGG

AATTATTCTTGGTAATTCTTCTGTAGATGTGTGTCTGCATGACACTTATTTTGTTGTTGCACATTTTCAT

TATGTTTTATCAATAGGAATTATTTTTGCTATTGTAGGGGGAGTTATCTATTGATTTCCACTAATCTTAG

GTTTAACCTTAAATAATTATAGATTGGTGTCTCAATTTTATATCATGTTTATTGGAGTAAATTTAACTTT

TTTTCCTCAGCATTTTCTTGGTTTAGGGGGAATGCCTCGTCGATATTCAGATTATGCTGATTGCTATCTA

GTATGAAATAAAATTTCTTCTGCGGGAAGGATTCTGAGTATTATTTCTGTTATTTATTTTTTATTTATTG

TTTTAGAATCCTTTCTTCTTCTGCGGT

>MidEastAm1_China_Zhejiang_GQ332577_GQ332577.1

GAAAATTAGAGGTATTTGGAAGGTTGGGTATAATTTATGCTATATTGACTATTGGTATTCTAGGGTTTAT

TGTTTGAGGTCATCATATATTCACAGTTGGAATAGATGTAGATACTCGAGCTTATTTCACTTCAGCCACT

ATAATTATTGCTGTTCCCACAGGAATTAAAATTTTTAGTTGGCTTGCTACTTTGGGTGGAATAAAGTCTA

ATAAATTAAGGCCTCTTGGCCTTTGATTTACAGGATTTTTATTTTTATTTACTATAGGTGGATTAACTGG

AATTATTCTTGGTAATTCTTCTGTAGATGTGTGTCTGCATGACACTTATTTTGTTGTTGCACATTTTCAT

TATGTTTTATCAATAGGAATTATTTTTGCTATTGTAGGAGGAGTTATCTATTGATTTCCACTAATCTTAG

GTTTAACCTTAAATAATTATAGATTGGTGTCTCAATTTTATATCATGTTTATTGGAGTAAATTTAACTTT

TTTTCCTCAGCATTTTCTTGGTTTAGGGGGAATGCCTCGTCGATATTCAGATTATGCTGATTGCTATCTA

GTATGAAATAAAATTTCTTCTGCGGGAAGGATTCTGAGTATTATTTCTGTTATTTATTTTTTATTTATTG

TTTTAGAATCCTTTCTTCTTCTGCGGT

>MidEastAm1_Saudi_Arabia_GU086344_GU086344.1

GAAAATTAGAGGTATTTGGAAGGTTGGGTATAATTTATGCTATACTGACTATTGGTATTCTAGGGTTTAT

TGTTTGAGGTCATCATATATTCACAGTTGGAATAGATGTAGATACTCGAGCTTATTTCACTTCAGCCACT

ATAATTATTGCTGTTCCCACAGGAATTAAAATTTTTAGTTGGCTTGCTACTTTGGGTGGAATAAAGTCTA

ATAAATTAAGGCCTCTTGGCCTTTGATTTACAGGATTTTTATTTTTATTTACTATAGGTGGGTTAACTGG

AATTATTCTTGGTAATTCTTCTGTAGATGTGTGTCTGCATGACACTTATTTTGTTGTTGCACATTTTCAT

TATGTTTTATCAATAGGAATTATTTTTGCTATTGTAGGAGGAGTTATCTATTGATTTCCACTAATCTTAG

GTTTAACCTTAAATAATTATAGATTGGTGTCTCAATTTTATATCATGTTTATTGGAGTAAATTTAACTTT

TTTTCCTCAGCATTTTCTTGGTTTAGGGGGAATGCCTCGTCGATATTCAGATTATGCTGATTGCTATCTA

GTATGAAATAAAATTTCTTCTGCGGGAAGGATTCTGAGTATTATTTCTGTTATTTATTTTTTATTTATTG

TTTTAGAATCCTTTCTTCTTCTGCGGT

>MidEastAm1_Japan_Kumamoto_AB204581_AB204581.1

GAAAATTAGAGGTATTTGGAAGGTTGGGTATAATTTATGCTATATTGACTATTGGTATTCTAGGGTTTAT

TGTTTGAGGTCATCATATATTCACAGTTGGAATAGATGTAGATACTCGAGCTTATTTCACTTCAGCCACT

ATAATTATTGCTGTTCCCACAGGAATTAAAATTTTTAGTTGGCTTGCTACTTTGGGTGGAATAAAGTCTA

ATAAATTAAGGCCTCTTGGCCTTTGATTTACAGGATTTTTATTTTTATTTACTATAGGTGGGTTAACTGG

AATTATTCTTGGTAATTCTTCTGTAGATGTGTGTCTGCATGACACTTATTTTGTTGTTGCACATTTTCAT

TATGTTTTATCAATAGGAATTATTTTTGCTATTGTAGGAGGAGTCATCTATTGATTTCCACTAATCTTAG

GTTTAACCTTAAATAATTATAGATTGGTGTCTCAATTTTATATCATGTTTATTGGAGTAAATTTAACTTT

TTTTCCTCAGCATTTTCTTGGTTTAGGGGGAATGCCTCGTCGATATTCAGATTATGCTGATTGCTATCTA

GTATGAAATAAAATTTCTTCTGCGGGAAGGATTCTGAGTATTATTTCTGTTATTTATTTTTTATTTATTG

TTTTAGAATCCTTTCTTCTTCTGCGGT

>MidEastAm1_Indonesia_Bogor_HM070410_HM070410.1

GAAAATTAGAGGTATTTGGAAGGTTGGGTATAATTTATGCTATATTGACTATTGGTATTCTAGGGTTTAT

TGTTTGAGGTCATCATATATTCACAGTTGGAATAGATGTAGATACTCGAGCTTATTTCACTTCAGCCACT

ATAATTATTGCTGTTCCCACAGGGATTAAAATTTTTAGTTGGCTTGCTACTTTGGGTGGAATAAAGTCTA

ATAAATTAAGGCCTCTTGGCCTTTGATTTACAGGATTTTTATTTTTATTTACTATAGGTGGGTTAACTGG

AATTATTCTTGGTAATTCTTCTGTAGATGTGTGTCTGCATGACACTTATTTTGTTGTTGCACATTTTCAT

TATGTTTTATCAATAGGAATTATTTTTGCTATTGTAGGAGGAGTTATCTATTGATTTCCACTAATCTTAG

GTTTAACCTTAAATAATTATAGATTGGTGTCTCAATTTTATATCATGTTTATTGGAGTAAATTTAACTTT

TTTTCCTCAGCATTTTCTTGGTTTAGGGGGAATGCCTCGTCGATATTCAGATTATGCTGATTGCTATCTA

GTATGAAATAAAATTTCTTCTGCGGGAAGGATTCTGAGTATTATTTCTGTTATTTATTTTTTATTTATTG

TTTTAGAATCCTTTCTTCTTCTGCGGT

>MidEastAm1_Cuba_Habana_FN821798_FN821798.1

GAAAATTAGAGGTATTTGGAAGGTTGGGTATAATTTATGCTATATTGACTATTGGTATTCTAGGGTTTAT

TGTTTGAGGTCATCATATATTCACAGTTGGAATAGATGTAGATACTCGAGCTTATTTCACTTCAGCCACT

ATAATTATTGCTGTTCCCACAGGAATTAAAATTTTTAGTTGGCTTGCTACTTTGGGTGGAATAAAGTCTA

ATAAATTAAGGCCTCTTGGCCTTTGATTTACAGGATTTTTATTTTTATTTACTATAGGTGGGTTAACTGG

AATTATTCTTGGTAATTCTTCTGTAGATGTGTGTCTGCATGACACTTATTTTGTTGTTGCACATTTTCAT

TATGTTTTATCGATAGGAATTATTTTTGCTATTGTAGGAGGAGTTATCTATTGATTTCCACTAATCTTAG

GTTTAACCTTAAATAATTATAGATTGGTGTCTCAATTTTATATCATGTTTATTGGAGTAAATTTAACTTT

TTTTCCTCAGCATTTTCTTGGTTTAGGGGGAATGCCTCGTCGATATTCAGATTATGCTGATTGCTATCTA

GTATGAAATAAAATTTCTTCTGCGGGAAGGATTCTGAGTATTATTTCTGTTATTTATTTTTTATTTATTG

TTTTAGAATCCTTTCTTCTTCTGCGGT

>MidEastAm1_Taiwan_GU086342_GU086342.1

GAAAATTAGAGGTATTTGGAAGGTTGGGTATAATTTATGCTATATTGACTATTGGTATTCTAGGGTTTAT

TGTTTGAGGTCATCATATATTCACAGTTGGAATAGATGTAGATACTCGAGCTTATTTCACTTCAGCCACT

ATAATTATTGCTGTTCCCACAGGAATTAAAATTTTTAGTTGGCTTGCTACTTTGGGTGGAATAAAGTCTA

ATAAATTAAGGCCTCTTGGCCTTTGATTTACAGGATTTTTATTTTTATTTACTATAGGTGGGTTAACTGG

AATTATTCTTGGTAATTCTTCTGTAGATGTGTGTCTGCATGACACTTATTTTGTTGTTGCACATTTTCAT

TATGTTTTATCAATAGGAATTATTTTTGCTATTGTAGGAGGAGTTATCTATTGATTTCCACTAATCTTAG

GTTTAACCTTAAATAATTATAGATTGGTGTCTCAATTTTATATCATGTTTATTGGAGTAAATTTAACTTT

TTTTCCTCAGCATTTTCTTGGTTTAGGGGGAATGCCTCGTCGATATTCAGATTATGCTGATTGCTATCTA

GTATGAAATAAAATTTCTTCTGCGGGAAGGATTCTGAGTATTATTTCCGTTATTTATTTTTTATTTATTG

TTTTAGAATCCTTTCTTCTTCTGCGGT

>MidEastAm1_Japan_Kumamoto_AB204580_AB204580.1

GAAAATTAGAGGTATTTGGAAGGTTGGGTATAATTTATGCTATATTGACTATTGGTATTCTAGGGTTTAT

TGTTTGAGGTCATCATATATTCACAGTTGGAATAGATGTAGATACTCGAGCTTATTTCACTTCAGCCACT

ATAATTATTGCTGTTCCCACAGGAATTAAAATTTTTAGTTGGCTTGCTACTTTGGGTGGAATAAAGTCTA

ATAAATTAAGACCTCTTGGCCTTTGATTTACAGGATTTTTATTTTTATTTACTATAGGTGGGTTAACTGG

AATTATTCTTGGTAATTCTTCTGTAGATGTGTGTCTGCATGACACTTATTTTGTTGTTGCACATTTTCAT

TATGTTTTATCAATAGGAATTATTTTTGCTATTGTAGGAGGAGTTATCTATTGATTTCCACTAATCTTAG

GTTTAACCTTAAATAATTATAGATTGGTGTCTCAATTTTATATCATGTTTATTGGAGTAAATTTAACTTT

TTTTCCTCAGCATTTTCTTGGTTTAGGGGGAATGCCTCGTCGATATTCAGATTATGCTGATTGCTATCTA

GTATGAAATAAAATTTCTTCTGCGGGAAGGATTCTGAGTATTATTTCTGTTATTTATTTTTTATTTATTG

TTTTAGAATCCTTTCTTCTTCTGCGGT

>MidEastAm1_Morocco_Nador_AM176570_AM176570.1

GAAAATTAGAGGTATTTGGAAGGTTGGGTATAATTTATGCTATATTGACTATTGGTATTCTGGGGTTTAT

TGTTTGAGGTCATCATATATTCACAGTTGGAATAGATGTAGATACTCGAGCTTATTTCACTTCAGCCACT

ATAATTATTGCTGTTCCCACAGGAATTAAAATTTTTAGTTGGCTTGCTACTTTGGGTGGAATAAAGTCTA

ATAAATTAAGGCCTCTTGGCCTTTGATTTACAGGATTTTTATTTTTATTTACTATAGGTGGGTTAACTGG

AATTATTCTTGGTAATTCTTCTGTAGATGTGTGTCTGCATGACACTTATTTTGTTGTTGCACATTTTCAT

TATGTTTTATCAATAGGAATTATTTTTGCTATTGTAGGAGGAGTTATCTATTGATTTCCACTAATCTTAG

GTTTAACCTTAAATAATTATAGATTGGTGTCTCAATTTTATATCATGTTTATTGGAGTAAATTTAACTTT

TTTTCCTCAGCATTTTCTTGGTTTAGGGGGAATGCCTCGTCGATATTCAGATTATGCTGATTGCTATCTA

GTATGAAATAAAATTTCTTCTGCGGGAAGGATTCTGAGTATTATTTCTGTTATTTATTTTTTATTTATTG

TTTTAGAATCCTTTCTTCTTCTGCGGT

>MidEastAm1_Egypt_Ismailia_GU977249_GU977249.1

GAAAATTAGAGGTATTTGGAAGGTTGGGTATAATTTATGCTATATTGACTATTGGTATTCTAGGGTTTAT

TGTTTGAGGTCATCATATATTCACAGTTGGAATAGATGTAGATACTCGAGCTTATTTCACTTCAGCCACT

ATAATTATTGCTGTTCCCACAGGAATTAAAATTTTTAGTTGGCTTGCTACTTTGGGTGGAATAAAGTCTA

ATAAATTAAGGCCTCTTGGCCTTTGATTTACAGGATTTTTATTTTTATTTACTATAGGTGGGTTAACAGG

AATTATTCTTGGTAATTCTTCTGTAGATGTGTGTCTGCATGACACTTATTTTGTTGTTGCACATTTTCAT

TATGTTTTATCAATAGGAATTATTTTTGCTATTGTAGGAGGAGTTATCTATTGATTTCCACTAATCTTAG

GTTTAACCTTAAATAATTATAGATTGGTGTCTCAATTTTATATCATGTTTATTGGAGTAAATTTAACTTT

TTTTCCTCAGCATTTTCTTGGTTTAGGGGGAATGCCTCGTCGATATTCAGATTATGCTGATTGCTATCTA

GTATGAAATAAAATTTCTTCTGCGGGAAGGATTCTGAGTATTATTTCTGTTATTTATTTTTTATTTATTG

TTTTAGAATCCTTTCTTCTTCTGCGGT

>MidEastAm1_Cuba_PinardelRio_FN821801_FN821801.1

GAAAATTAGAGGTATTTGGAAGGTTGGGTATAATTTATGCTATATTGACTATTGGTATTCTAGGGTTTAT

TGTTTGAGGTCATCATATATTCACAGTTGGAATAGATGTAGATACTCGAGCTTATTTCACTTCAGCCACT

ATAATTATTGCTGTTCCCACAGGAATTAAAATTTTTAGTTGGCTTGCTACTTTGGGTGGAATAAAGTCTA

ATAAATTAAGGCCTCTTGGCCTTTGATTTACAGGATTTTTATTTTTATTTACTATAGGTGGGTTAACTGG

AATTATTCTTGGTAATTCTTCTGTAGATGTGTGTCTGCATGACACTTATTTTGTTGTTGCACATTTTCAT

TATGTTTTATCAATAGGAATCATTTTTGCTATTGTAGGAGGAGTTATCTATTGATTTCCACTAATCTTAG

GTTTAACCTTAAATAATTATAGATTGGTGTCTCAATTTTATATCATGTTTATTGGAGTAAATTTAACTTT

TTTTCCTCAGCATTTTCTTGGTTTAGGGGGAATGCCTCGTCGATATTCAGATTATGCTGATTGCTATCTA

GTATGAAATAAAATTTCTTCTGCGGGAAGGATTCTGAGTATTATTTCTGTTATTTATTTTTTATTTATTG

TTTTAGAGTCCTTTCTTCTTCTGCGGT

>MidEastAm1_Cuba_Camaguey_FN821808_FN821808.1

GAAAATTAGAGGTATTTGGAAGGTTGGGTATAATTTATGCTATATTGACTATTGGTATTCTAGGGTTTAT

TGTTTGAGGTCATCATATATTCACAGTTGGAATAGATGCAGATACTCGAGCTTATTTCACTTCAGCCACT

ATAATTATTGCTGTTCCCACAGGAATTAAAATTTTTAGTTGGCTTGCTACTTTGGGTGGAATAAAGTCTA

ATAAATTAAGGCCTCTTGGCCTTTGATTTACAGGATTTTTATTTTTATTTACTATAGGTGGGTTAACTGG

AATTATTCTTGGTAATTCTTCCGTAGATGTGTGTCTGCATGACACTTATTTTGTTGTTGCACATTTTCAT

TATGTTTTATCAATAGGAATTATTTTTGCTATTGTAGGAGGAGTTATCTATTGATTTCCACTAATCTTAG

GTTTAACCTTAAATAATTATAGATTGGTGTCTCAATTTTATATCATGTTTATTGGAGTAAATTTAACTTT

TTTTCCTCAGCATTTTCTTGGTTTAGGGGGAATGCCTCGTCGATATTCAGATTATGCTGATTGCTATCTA

GTATGAAATAAAATTTCTTCTGCGGGAAGGATTCTGAGTATTATTTCTGTTATTTATTTTTTATTTATTG

TTTTAGAATCCTTTCTTCTTCTGCGGT

>MidEastAm1_Iraq_Al_A_Zamiyah_HM070413

GAAAATTAGAGGTATTTGGAAGGTTGGGTATAATTTATGCTATACTGACTATTGGTATTCTAGGGTTTAT

TGTTTGAGGTCATCATATATTCACAGTTGGAATAGATGTAGATACTCGAGCTTATTTCACTTCAGCCACT

ATAATTATTGCTGTTCCCACAGGAATTAAAATTTTTAGTTGGCTTGCTACTTTGGGTGGAATAAAGTCTA

ATAAATTAAGGCCTCTTGGCCTTTGATTTACAGGATTTTTATTTTTATTTACTATAGGTGGGTTAACTGG

AATTATCCTTGGTAATTCTTCTGTAGATGTGTGTCTGCATGACACTTATTTTGTTGTTGCACATTTTCAT

TATGTCTTATCAATAGGAATCATTTTTGCTATTGTAGGAGGAGTTATCTATTGATTTCCACTAATCTTAG

GTTTAACCTTAAATAATTATAGATTGGTGTCTCAATTTTATATCATGTTTATTGGAGTAAATTTAACTTT

TTTTCCTCAGCATTTTCTTGGTTTAGGGGGAATGCCTCGTCGATACTCAGATTATGCTGATTGCTATCTA

GTATGAAATAAAATTTCTTCTGCGGGAAGGATTTTGAGTATTATTTCTGTTATTTATTTTTTATTTATTG

TTTTAGAATCCTTTCTTCTTCTGCGGT

>MidEastAm1_United_Arab_Emirates_DQ133382_DQ133382.1

GAAAATTAGAGGTATTTGGAAGGTTGGGTATAATTTATGCTATACTGACTATTGGTATTCTAGGGTTTAT

TGTTTGAGGTCATCATATGTTCACAGTTGGAATAGATGTAGATACTCGAGCTTATTTCACTTCAGCCACT

ATAATTATTGCTGTTCCCACAGGAATTAAAATTTTTAGTTGGCTTGCTACTTTGGGTGGAATAAAGTCTA

ATAAATTAAGGCCTCTTGGCCTTTGATTTACAGGATTTTTATTTTTATTTACTATAGGTGGGTTAACTGG

AATTATTCTTGGTAATTCTTCTGTAGATGTGTGTCTGCATGACACTTATTTTGTTGTTGCACATTTTCAT

TATGTCTTATCAATAGGAATCATTTTTGCTATTGTAGGAGGAGTTATCTATTGATTTCCACTAATCTTAG

GTTTAACCTTAAATAATTATAGATTGGTGTCTCAATTTTATATCATGTTTATTGGAGTAAATTTAACTTT

TTTTCCTCAGCATTTTCTTGGTTTAGGGGGAATGCCTCGTCGATATTCAGATTATGCTGATTGCTATCTA

GTATGAAATAAAATTTCTTCTGCGGGAAGGATTTTGAGTATTATTTCTGTTATTTATTTTTTATTTATTG

TTTTAGAATCCTTTCTTCTTCTGCGGT

>MidEastAm1_Syria_Al_Hasakeh_Hap2_AB473559_AB473559.1

GAAAATTAGAGGTATTTGGAAGGTTGGGTATAATTTATGCTATATTGACTATTGGTATTCTAGGGTTTAT

TGTTTGAGGTCATCATATATTCACAGTTGGAATAGATGTAGATACTCGAGCTTATTTCACTTCAGCCACT

ATAATTATTGCTGTTCCCACAGGAATTAAAATTTTTAGTTGGCTTGCTACTTTGGGTGGAATAAAGTCTA

ATAAATTAAGGCCTCTTGGCCTTTGATTTACAGGATTTTTATTTTTATTTACTATAGGTGGGTTAACTGG

AATTATTCTTGGTAATTCTTCTGTAGATGTGTGTCTGCATGACACTTATTTTGTTGTTGCACATTTTCAT

TATGTCTTATCAATAGGAATCATTTTTGCTATTGTAGGAGGAGTTATCTATTGATTTCCACTAATCTTAG

GTTTAACCTTAAATAATTATAGATTGGTGTCTCAATTTTATATCATGTTTATTGGAGTAAATTTAACTTT

TTTTCCTCAGCATTTTCTTGGTTTAGGGGGAATGCCTCGTCGATACTCAGATTATGCTGATTGCTATCTA

GTATGAAATAAAATTTCTTCTGCGGGAAGGATTCTGAGTATTATTTCTGTTATTTATTTTTTATTTATTG

TTTTAGAATCCTTTCTTCTTCTGCGGT

>MidEastAm1_Iran_GU086352_GU086352.1

GAAAATTAGAGGTATTTGGAAGGTTGGGTATAATTTATGCTATACTGACTATTGGTATTCTAGGGTTTAT

TGTTTGAGGTCATCATATATTCACAGTTGGAATAGATGTAGATACTCGAGCTTATTTCACTTCAGCCACT

ATAATTATTGCTGTTCCCACAGGAATTAAAATTTTTAGTTGGCTTGCTACTTTGGGTGGAATAAAGTCTA

ATAAATTAAGGCCTCTTGGCCTTTGATTTACAGGATTTTTATTTTTATTTACTATAGGTGGGTTAACTGG

AATTATCCTTGGGAATTCTTCTGTAGATGTGTGTCTGCATGACACTTATTTTGTTGTTGCACATTTTCAT

TATGTTTTATCAATAGGAATTATTTTTGCTATTGTAGGAGGAGTTATCTATTGATTTCCACTAATCTTAG

GTTTAACCTTAAATAATTATAGATTGGTGTCTCAATTTTATATCATGTTTATTGGAGTAAATTTAACTTT

TTTTCCTCAGCATTTTCTTGGTTTAGGGGGAATGCCTCGTCGATATTCAGATTATGCTGATTGCTATCTA

GTATGAAATAAAATTTCTTCTGCGGGAAGGATTCTGAGTATTATTTCTGTTATTTATTTTTTATTTATTG

TTTTAGAATCCTTTCTTCTTCTGCGGT

>MidEastAm1_Kuwait_GU086356_GU086356.1

GAAAATTAGAGGTATTTGGAAGGTTGGGTATAATTTATGCTATACTGACTATTGGTATTCTAGGGTTTAT

TGTTTGAGGTCATCATATATTCACAGTTGGAATAGATGTAGATACTCGAGCTTATTTCACTTCAGCCACT

ATAATTATTGCTGTTCCCACAGGAATTAAAATTTTTAGTTGGCTTGCTACTTTGGGTGGAATAAAGTCTA

ATAAATTAAGGCCTCTTGGCCTTTGATTTACAGGATTTTTATTTTTATTTACTATAGGTGGGTTAACTGG

AATTATCCTTGGGAATTCTTCTGTAGATGTGTGTCTGCATGACACTTATTTTGTTGTTGCACATTTTCAT

TATGTCTTATCAATAGGAATTATTTTTGCTATTGTAGGAGGAGTTATCTATTGATTTCCACTAATCTTAG

GTTTAACCTTAAATAATTATAGATTGGTGTCTCAATTTTATATCATGTTTATTGGAGTAAATTTAACTTT

TTTTCCTCAGCATTTTCTTGGTTTAGGGGGAATGCCTCGTCGATATTCAGATTATGCTGATTGCTATCTA

GTATGAAATAAAATTTCTTCTGCGGGAAGGATTCTGAGTATTATTTCTGTTATTTATTTTTTATTTATTG

TTTTAGAATCCTTTCTTCTTCTGCGGT

>MidEastAm1_Saudi_Arabia_GU086358_GU086358.1

GAAAATTAGAGGTATTTGGAAGGTTGGGTATAATTTATGCTATATTGACTATTGGTATTCTAGGGTTTAT

TGTTTGAGGTCATCATATGTTCACAGTTGGAATAGATGTAGATACTCGAGCTTATTTCACTTCAGCCACT

ATAATTATTGCTGTTCCCACAGGAATTAAAATTTTTAGTTGGCTTGCTACTTTGGGTGGAATAAAGTCTA

ATAAATTAAGGCCTCTTGGCCTTTGATTTACAGGATTTTTATTTTTATTTACTATAGGTGGGTTAACTGG

AATTATCCTTAGGAATTCTTCTGTAGATGTGTGTCTGCATGACACTTATTTTGTTGTTGCACATTTTCAT

TATGTCTTATCAATAGGAATTATTTTTGCTATTGTAGGAGGAGTTATCTATTGATTTCCACTAATCTTAG

GTTTAACCTTAAATAATTATAGATTGGTGTCTCAATTTTATATCATGTTTATTGGAGTAAATTTAACTTT

TTTTCCTCAGCATTTTCTTGGTTTAGGGGGAATGCCTCGTCGATATTCAGATTATGCTGATTGCTATCTA

GTATGAAATAAAATTTCTTCTGCGGGAAGGATTCTGAGTATTATTTCTGTTATTTATTTTTTATTTATTG

TTTTAGAATCCTTTCTTCTTCTGCGGT

>MidEastAm1_Yemen_GU086359_GU086359.1

GAAAATTAGAGGTATTTGGAAGGTTGGGTATAATTTATGCTATATTGACTATTGGTATTCTAGGGTTTAT

TGTTTGAGGTCATCATATGTTCACAGTTGGAATAGATGTAGATACTCGAGCTTATTTCACTTCAGCCACT

ATAATTATTGCTGTTCCCACAGGAATTAAAATTTTTAGTTGGCTTGCTACTTTGGGTGGAATAAAGTCTA

ATAAATTAAGGCCTCTTGGCCTTTGATTTACAGGATTTTTATTTTTATTTACTATAGGTGGGTTAACTGG

AATTATCCTTGGGAATTCTTCTGTAGATGTGTGTCTGCATGACACTTATTTTGTTGTTGCACATTTTCAT

TATGTCTTATCAATAGGAATTATTTTTGCTATTGTAGGAGGAGTTATCTATTGATTTCCACTAATCTTAG

GTTTAACCTTAAATAATTATAGATTGGTGTCTCAATTTTATATCATGTTTATTGGAGTAAATTTAACTTT

TTTTCCTCAGCATTTTCTTGGTTTAGGGGGAATGCCTCGTCGATATTCAGATTATGCTGATTGCTATCTA

GTATGAAATAAAATTTCTTCTGCGGGAAGGATTCTGAGTATTATTTCTGTTATTTATTTTTTATTTATTG

TTTTAGAATCCTTTCTTCTTCTGCGGT

>MidEastAm1_Iran_GU086350_GU086350.1

GAAAATTAGAGGTATTTGGAAGGTTGGGTATAATTTATGCTATACTGACTATTGGTATTCTAGGGTTTAT

TGTTTGAGGTCATCATATGTTCACAGTTGGAATAGATGTAGATACTCGAGCTTATTTCACTTCAGCCACT

ATAATTATTGCTGTTCCCACAGGAATTAAAATTTTTAGTTGGCTTGCTACTTTGGGTGGAATAAAGTCTA

ATAAATTAAGGCCTCTTGGCCTTTGATTTACAGGATTTTTATTTTTATTTACTATAGGTGGGTTAACTGG

AATTATCCTTGGGAATTCTTCTGTAGATGTGTGTCTGCATGACACTTATTTTGTTGTTGCACATTTTCAT

TATGTCTTATCAATAGGAATTATTTTTGCTATTGTAGGAGGAGTTATCTATTGATTTCCACTAATCTTAG

GTTTAACCTTAAATAATTATAGATTGGTGTCTCAATTTTATATCATGTTTATTGGAGTAAATTTAACTTT

TTTTCCTCAGCATTTTCTTGGTTTAGGGGGAATGCCTCGTCGATATTCAGATTATGCTGATTGCTATCTA

GTATGAAATAAAATTTCTTCTGCGGGAAGGATTCTGAGTATTATTTCTGTTATTTATTTTTTATTTATTG

TTTTAGAATCCTTTCTTCTTCTGCGGT

>MidEastAm1_Kuwait_GU086346_GU086346.1

GAAAATTAGAGGTATTTGGAAGGTTGGGTATAATTTATGCTATACTGACTATTGGTATTCTAGGGTTTAT

TGTTTGAGGTCATCATATGTTCACAGTTGGAATAGATGTAGATACTCGAGCTTATTTCACTTCAGCCACT

ATAATTATTGCTGTTCCCACAGGAATTAAAATTTTTAGTTGGCTTGCTACTTTGGGTGGAATAAAGTCTA

ATAAATTAAGGCCTCTTGGCCTTTGATTTACAGGATTTTTATTTTTATTTACTATAGGTGGGTTAACTGG

AATTATCCTTAGGAATTCTTCTGTAGATGTGTGTCTGCATGACACTTATTTTGTTGTTGCACATTTTCAT

TATGTCTTATCAATAGGAATTATTTTTGCTATTGTAGGAGGAGTTATCTATTGATTTCCACTAATCTTAG

GTTTAACCTTAAATAATTATAGATTGGTGTCTCAATTTTATATCATGTTTATTGGAGTAAATTTAACTTT

TTTTCCTCAGCATTTTCTTGGTTTAGGGGGAATGCCTCGTCGATATTCAGATTATGCTGATTGCTATCTA

GTATGAAATAAAATTTCTTCTGCGGGAAGGATTCTGAGTATTATTTCTGTTATTTATTTTTTATTTATTG

TTTTAGAATCCTTTCTTCTTCTGCGGT

>MidEastAm1_Saudi_Arabia_GU086357_GU086357.1

GAAAATTAGAGGTATTTGGAAGGTTGGGTATAATTTATGCTATATTGACTATTGGTATTCTAGGGTTTAT

TGTTTGAGGTCATCATATATTCACAGTTGGAATAGATGTAGATACTCGAGCTTATTTCACTTCAGCCACT

ATAATTATTGCTGTTCCCACAGGAATTAAAATTTTTAGTTGGCTTGCTACTTTGGGTGGAATAAAGTCTA

ATAAATTAAGGCCTCTTGGCCTTTGATTTACAGGATTTTTATTTTTATTTACTATAGGTGGGTTAACTGG

AATTATCCTTAGGAATTCTTCTGTAGATGTGTGTCTGCATGACACTTATTTTGTTGTTGCACATTTTCAT

TATGTCTTATCAATAGGAATTATTTTTGCTATTGTAGGAGGAGTTATCTATTGATTTCCACTAATCTTAG

GTTTAACCTTAAATAATTATAGATTGGTGTCTCAATTTTATATCATGTTTATTGGAGTAAATTTAACTTT

TTTTCCTCAGCATTTTCTTGGTTTAGGGGGAATGCCTCGTCGATATTCAGATTATGCTGATTGCTATCTA

GTATGAAATAAAATTTCTTCTGCGGGAAGGATTTTGAGTATTATTTCTGTTATTTATTTTTTATTTATTG

TTTTAGAATCCTTTCTTCTTCTGCGGT

>MidEastAm1_Iran_GU086353_GU086353.1

GAAAATTAGAGGTATTTGGAAGGTTGGGTATAATTTATGCTATATTGACTATTGGTATTCTAGGGTTTAT

TGTTTGAGGTCATCATATGTTCACAGTTGGAATAGATGTAGATACTCGAGCTTATTTCACTTCAGCCACT

ATAATTATTGCTGTTCCCACAGGAATTAAAATTTTTAGTTGGCTTGCTACTTTGGGTGGAATAAAGTCTA

ATAAATTAAGGCCTCTTGGCCTTTGATTTACAGGATTTTTATTTTTATTTACTATAGGTGGGTTAACTGG

AATTATCCTTAGGAATTCTTCTGTAGATGTGTGTCTGCATGACACTTATTTTGTTGTTGCACATTTTCAT

TATGTCTTATCAATAGGAATTATTTTTGCTATTGTAGGAGGAGTTATCTATTGATTTCCACTAATCTTAG

GTTTAACCTTAAATAATTATAGATTGGTGTCTCAATTTTATATCATGTTTATTGGAGTAAATTTAACTTT

TTTTCCTCAGCATTTTCTTGGTTTAGGGGGAATGCCTCGTCGATATTCAGATTATGCTGATTGCTATCTA

GTATGAAATAAAATTTCTTCTGCGGGAAGGATTTTGAGTATTATTTCTGTTATTTATTTTTTATTTATTG

TTTTAGAATCCTTTCTTCTTCTGCGGT

>MidEastAm1_Iran_GU086351_GU086351.1

GAAAATTAGAGGTATTTGGAAGGTTGGGTATAATTTATGCTATACTGACTATTGGTATTCTAGGGTTTAT

TGTTTGAGGTCATCATATGTTCACAGTTGGAATAGATGTAGATACTCGAGCTTATTTCACTTCAGCCACT

ATAATTATTGCTGTTCCCACAGGAATTAAAATTTTTAGTTGGCTTGCTACTTTGGGTGGAATAAAGTCTA

ATAAATTAAGGCCTCTTGGCCTTTGATTTACAGGATTTTTATTTTTATTTACTATAGGTGGGTTAACTGG

AATTATCCTTAGGAATTCTTCTGTAGATGTGTGTCTGCATGACACTTATTTTGTTGTTGCACATTTTCAT

TATGTCTTATCAATAGGAATTATTTTTGCTATTGTAGGAGGAGTTATCTATTGATTTCCACTAATCTTAG

GTTTAACCTTAAATAATTATAGATTGGTGTCTCAATTTTATATCATGTTTATTGGAGTAAATTTAACTTT

TTTTCCTCAGCATTTTCTTGGTTTAGGGGGAATGCCTCGTCGATATTCAGATTATGCTGATTGCTATCTA

GTATGAAATAAAATTTCTTCTGCGGGAAGGATTTTGAGTATTATTTCTGTTATTTATTTTTTATTTATTG

TTTTAGAATCCTTTCTTCTTCTGCGGT

>MidEastAm1_Saudi_Arabia_GU086345_GU086345.1

GAAAATTAGAGGTATTTGGAAGGTTGGGTATAATTTATGCTATACTGACTATTGGTATTCTAGGGTTTAT

TGTTTGAGGTCATCATATGTTCACAGTTGGAATAGATGTAGATACTCGAGCTTATTTCACTTCAGCCACT

ATAATTATTGCTGTTCCCACAGGAATTAAAATTTTTAGTTGGCTTGCTACTTTGGGTGGAATAAAGTCTA

ATAAATTAAGGCCTCTTGGCCTTTGATTTACAGGATTTTTATTTTTATTTACTATAGGTGGGTTAACTGG

AATTATCCTTAGGAATTCTTCTGTAGATGTGTGTCTGCATGACACTTATTTTGTTGTTGCACATTTTCAT

TATGTCTTATCAATAGGAATCATTTTTGCTATTGTAGGAGGAGTTATCTATTGATTTCCACTAATCTTAG

GTTTAACCTTAAATAATTATAGATTGGTGTCTCAATTTTATATCATGTTTATTGGAGTAAATTTAACTTT

TTTTCCTCAGCATTTTCTTGGTTTAGGGGGAATGCCTCGTCGATATTCAGATTATGCTGATTGCTATCTA

GTATGAAATAAAATTTCTTCTGCGGGAAGGATTTTGAGTATTATTTCTGTTATTTATTTTTTATTTATTG

TTTTAGAATCCTTTCTTCTTCTGCGGT

>MidEastAm1_Kuwait_GU086355_GU086355.1

GAAAATTAGAGGTATTTGGAAGGTTGGGTATAATTTATGCTATATTGACTATTGGTATTCTAGGGTTTAT

TGTTTGAGGTCATCATATGTTCACAGTTGGAATAGATGTAGATACTCGAGCTTATTTCACTTCAGCCACT

ATAATTATTGCTGTTCCCACAGGAATTAAAATTTTTAGTTGGCTTGCTACTTTGGGTGGAATAAAGTCTA

ATAAATTAAGGCCTCTTGGCCTTTGATTTACAGGATTTTTATTTTTATTTACTATAGGTGGGTTAACTGG

AATTATCCTTGGGAATTCTTCTGTAGATGTGTGTCTGCATGACACTTATTTTGTTGTTGCACATTTTCAT

TATGTCTTATCAATAGGAATCATTTTTGCTATTGTAGGAGGAGTTATCTATTGATTTCCACTAATCTTAG

GTTTAACCTTAAATAATTATAGATTGGTGTCTCAATTTTATATCATGTTTATTGGAGTAAATTTAACTTT

TTTTCCTCAGCATTTTCTTGGTTTAGGGGGAATGCCTCGTCGATATTCAGATTATGCTGATTGCTATCTA

GTATGAAATAAAATTTCTTCTGCGGGAAGGATTTTGAGTATTATTTCTGTTATTTATTTTTTATTTATTG

TTTTAGAATCCTTTCTTCTTCTGCGGT

>MidEastAm1_Kuwait_GU086354_GU086354.1

GAAAATTAGAGGTATTTGGAAGGTTGGGTATAATTTATGCTATACTGACTATTGGTATTCTAGGGTTTAT

TGTTTGAGGTCATCATATGTTCACAGTTGGAATAGATGTAGATACTCGAGCTTATTTCACTTCAGCCACT

ATAATTATTGCTGTTCCCACAGGAATTAAAATTTTTAGTTGGCTTGCTACTTTGGGTGGAATAAAGTCTA

ATAAATTAAGGCCTCTTGGCCTTTGATTTACAGGATTTTTATTTTTATTTACTATAGGTGGGTTAACTGG

AATTATCCTTGGGAATTCTTCTGTAGATGTGTGTCTGCATGACACTTATTTTGTTGTTGCACATTTTCAT

TATGTCTTATCAATAGGAATCATTTTTGCTATTGTAGGAGGAGTTATCTATTGATTTCCACTAATCTTAG

GTTTAACCTTAAATAATTATAGATTGGTGTCTCAATTTTATATCATGTTTATTGGAGTAAATTTAACTTT

TTTTCCTCAGCATTTTCTTGGTTTAGGGGGAATGCCTCGTCGATATTCAGATTATGCTGATTGCTATCTA

GTATGAAATAAAATTTCTTCTGCGGGAAGGATTTTGAGTATTATTTCTGTTATTTATTTTTTATTTATTG

TTTTAGAATCCTTTCTTCTTCTGCGGT

>MidEastAm1_Yemen_GU086360_GU086360.1

GAAAATTAGAGGTATTTGGAAGGTTGGGTATAATTTATGCTATATTGACTATTGGTATTCTAGGGTTTAT

TGTTTGAGGTCATCATATATTCACAGTTGGAATAGATGCAGATACTCGAGCTTATTTCACTTCAGCCACT

ATAATTATTGCTGTTCCCACAGGAATTAAAATTTTTAGTTGGCTTGCTACTTTGGGTGGAATAAAGTCTA

ATAAATTAAGGCCTCTTGGCCTTTGATTTACAGGATTTTTATTTTTATTTACTATAGGTGGGTTAACTGG

AATTATCCTTGGGAATTCTTCTGTAGATGTGTGTCTGCATGACACTTATTTTGTTGTTGCACATTTTCAT

TATGTCTTATCAATAGGAATTATTTTTGCTATTGTAGGAGGAGTTATCTATTGATTTCCACTAATCTTAG

GTTTAACCTTAAATAATTATAGATTGGTGTCTCAATTTTATATCATGTTTATTGGAGTAAATTTAACTTT

TTTTCCTCAGCATTTTCTTGGTTTAGGGGGAATGCCTCGTCGATATTCAGATTATGCTGATTGCTATCTA

GTATGAAATAAAATTTCTTCTGCGGGAAGGATTCTGAGTATTATTTCTGTTATTTATTTTTTATTTATTG

TTTTAGAATCCTTTCTTCTTCTGCGGT

>Mediterranean_Burkina_Faso_FJ766381_FJ766381.1

GAAAATTAGAGGTATTTGGAAGGTTGGGCATAATTTATGCTATACTGACTATTGGTATCTTAGGGTTTAT

TGTTTGAGGACATCATATATTTACAGTTGGAATAGATGTAGATACTCGAGCTTATTTCACTTCAGCTACT

ATGATTATTGCCGTTCCTACAGGAATTAAAATTTTTAGTTGGCTTGCTACTTTGGGTGGAATAAAGTCCA

ATAAATTCAGGCCCCTTGGCCTTTGATTTACAGGATTTTTATTTTTATTTACTATAGGTGGATTAACTGG

AATTATTCTTGGTAACTCTTCTGTAGATGTGTGTTTGCATGACACTTATTTTGTTGTTGCGCATTTTCAT

TATGTCTTATCAATAGGAATTATTTTTGCTATTGTAGGAGGAGTTATCTATTGATTTCCATTAATCTTGG

GCTTAACCTTAAATAATTATAGCTTGGTGTCTCAATTTTATATCATGTTCATTGGAGTAAATTTAACTTT

TTTTCCTCAGCATTTTCTTGGTTTGGGGGGAATGCCTCGCCGATATTCAGACTATGCTGATTGTTATCTA

GTATGGAACAAAATTTCTTCTGCGGGAAGGATTTTGAGTATCATTTCTGTTATTTATTTTTTATTTATTG

TTTTAGAATCTTTTCTTCTTTTGCGTT

>Mediterranean_China_Q_Hap1_GU086329_GU086329.1

GAAAATTAGAGGTATTTGGAAGGTTGGGCATAATTTATGCTATATTGACTATTGGTATCTTAGGGTTTAT

TGTTTGAGGACATCATATATTTACAGTTGGAATAGATGTAGATACTCGAGCTTATTTCACTTCAGCTACT

ATGATTATTGCCGTTCCTACAGGAATTAAAATTTTTAGTTGGCTTGCTACTTTGGGTGGAATAAAGTCCA

ATAAATTCAGGCCCCTTGGCCTTTGATTTACAGGATTTTTATTTTTATTTACTATAGGTGGATTAACTGG

AATTATTCTTGGTAACTCTTCTGTAGATGTGTGTTTGCATGACACTTATTTTGTTGTTGCGCATTTTCAT

TATGTCTTATCAATAGGAATTATTTTTGCTATTGTAGGAGGAGTTATCTATTGATTTCCATTAATCTTGG

GCTTAACCTTAAATAATTATAGCTTGGTGTCTCAATTTTATATCATGTTCATTGGAGTAAATTTAACTTT

TTTTCCTCAGCATTTTCTTGGTTTGGGGGGAATGCCTCGCCGATATTCAGATTATGCTGATTGTTATCTA

GTATGGAACAAAATTTCTTCTGCGGGAAGGATTTTGAGTATCATTTCTGTTATTTATTTTTTATTTATTG

TTTTAGAATCTTTTCTTCTTTTGCGTT

>Mediterranean_France_AM691067_AM691067.1

GAAAATTAGAGGTATTNGGAAGGTTGGGCATAATTTATGCTATATTGACTATTGGTATNTTAGGGTTTAT

TGTTTGAGGACATCATATATTTACAGTTGGAATAGATGTAGATACTCGAGCTTATTTCACTTCAGCTACT

ATGATTATTGCCGTTCCTACAGGAATTAAAATTTTTAGTTGGCTTGCTACTTTGGGTGGAATAAAGTCCA

ATAAATTCAGGCCCCTTGGCCTTTGATTTACAGGATTTTTATTTTTATTTACTATAGGTGGATTAACTGG

AATTATTCTTGGTAACTCTTCTGTAGATGTGTGTTTGCATGACACTTATTTTGTTGTTGCGCATTTTCAT

TATGTCTTATCAATAGGAATTATTTTTGCTATTGTAGGAGGAGTTATCTATTGATTTCCATTAATCTTGG

GCTTAACCTTAAATAATTATAGCTTGGTGTCTCAATTTTATATCATGTTCATTGGAGTAAATTTAACTTT

TTTTCCTCAGCATTTTCTTGGTTTGGGGGGAATGCCTCGCCGATATTCAGATTATGCTGATTGTTATCTA

GTATGGAACAAAATTTCTTCTGCGGGAAGGATTTTGAGTATCATTTCTGTTATTTATTTTTTATTTATTG

TTTTAGAATCTTTTCTTCTTTTGCGTT

>Mediterranean_Taiwan_AOP_GU086339_GU086339.1

GAAAATTAGAGGTATTTGGAAGGTTGGGCATAATTTATGCTATATTGACTATTGGTATCTTAGGGTTTAT

TGTTTGAGGACATCATATATTTACAGTTGGGATAGATGTAGATACTCGAGCTTATTTCACTTCAGCTACT

ATGATTATTGCCGTTCCTACAGGAATTAAAATTTTTAGTTGGCTTGCTACTTTGGGTGGAATAAAGTCCA

ATAAATTCAGGCCCCTTGGCCTTTGATTTACAGGATTTTTATTTTTATTTACTATAGGTGGATTAACTGG

AATTATTCTTGGTAACTCTTCTGTAGATGTGTGTTTGCATGACACTTATTTTGTTGTTGCGCATTTTCAT

TATGTCTTATCAATAGGAATTATTTTTGCTATTGTAGGAGGAGTTATCTATTGATTTCCATTAATCTTGG

GCTTAACCTTAAATAATTATAGCTTGGTGTCTCAATTTTATATCATGTTCATTGGAGTAAATTTAACTTT

TTTTCCTCAGCATTTTCTTGGTTTGGGGGGAATGCCTCGCCGATACTCAGATTATGCTGATTGTTATCTA

GTATGGAACAAAATTTCTTCTGCGGGAAGGATTTTGAGTATCATTTCTGTTATTTATTTTTTATTTATTG

TTTTAGAATCTTTTCTTCTTTTGCGTT

>Mediterranean_France_Hap21_AM691062_AM691062.1

GAAAATTAGAGGTATTTGGAAGGTTGGGCATAATTTATGCTATATTGACTATTGGTATCTTAGGGTTTAT

TGTTTGAGGACATCATATATTTACAGTTGGAATAGATGTAGATACTCGAGCTTATTTCACTTCAGCTACT

ATGATTATTGCCGTTCCTACAGGAATTAAAATTTTTAGTTGGCTTGCTACTTTGGGTGGAATAAAGTCCA

ATAAATTCAGGCCCCTTGGCCTTTGATTTACAGGATTTTTATTTTTATTTACTATAGGTGGATTAACTGG

AATTATTCTTGGTAACTCTTCTGTAGATGTGTGTTTGCATGATACTTATTTTGTTGTTGCGCATTTTCAT

TATGTCTTATCAATAGGAATTATTTTTGCTATTGTAGGAGGAGTTATTTATTGATTTCCATTAATCTTGG

GCTTAACCTTAAATAATTATAGCTTGGTGTCTCAATTTTATATCATGTTCATTGGAGTAAATTTAACTTT

TTTTCCTCAGCATTTTCTTGGTTTGGGGGGAATGCCTCGCCGATATTCAGATTATGCTGATTGTTATCTA

GTATGGAACAAAATTTCTTCTGCGGGAAGGATTTTGAGTATTATTTCTGTTATTTATTTTTTATTTATTG

TTTTAGAATCTTTTCTTCTTTTGCGTT

>Mediterranean_Cameroon_Hap20_EU760721_EU760721.1

GAAAATTAGAGGTATTTGGAAGGTTGGGCATAATTTATGCTATATTGACTATTGGTATCTTAGGGTTTAT

TGTTTGAGGACATCATATATTTACAGTTGGAATAGATGTAGATACTCGAGCTTATTTCACTTCAGCTACT

ATGATTATTGCCGTTCCTACAGGAATTAAAATTTTTAGTTGGCTTGCTACTTTGGGTGGAATAAAGTCCA

ATAAATTCAGGCCCCTTGGCCTTTGATTTACAGGATTTTTATTTTTATTTACTATAGGTGGATTAACTGG

AATTATTCTTGGTAACTCTTCTGTAGATGTGTGTTTGCATGACACTTATTTTGTTGTTGCACATTTTCAT

TATGTCTTATCAATAGGAATTATTTTTGCTATTGTAGGAGGAGTTATCTATTGATTTCCATTAATCTTGG

GCTTAACCTTAAATAATTATAGCTTGGTGTCTCAATTTTATATCATGTTCATTGGAGTAAATTTAACTTT

TTTTCCTCAGCATTTTCTTGGTTTGGGGGGAATGCCTCGCCGATATTCAGATTATGCTGATTGTTATCTA

GTATGGAACAAAATTTCTTCTGCGGGAAGGATTTTGAGTATTATTTCTGTTATTTATTTTTTATTTATTG

TTTTAGAATCTTTTCTTCTTTTGCGTT

>Mediterranean_China_4_Hap2_GU086333_GU086333.1

GAAAATTAGAGGTATTTGGAAGGTTGGGGATAATTTATGCTATATTGACTATTGGTATCTTAGGGTTTAT

TGTTTGAGGACATCATATATTTACAGTTGGAATAGATGTAGATACTCGAGCTTATTTCACTTCAGCTACT

ATGATTATTGCCGTTCCTACAGGAATTAAAATTTTTAGTTGGCTTGCTACTTTGGGTGGAATAAAGTCCA

ATAAATTCAGGCCCCTTGGCCTTTGATTTACAGGATTTTTATTTTTATTTACTATAGGTGGATTAACTGG

AATTATTCTTGGTAACTCTTCTGTAGATGTGTGTTTGCATGACACTTATTTTGTTGTTGCGCATTTTCAT

TATGTCTTATCAATAGGAATTATTTTTGCTATTGTAGGAGGAGTTATCTATTGATTTCCATTAATCTTGG

GCTTAACCTTAAATAATTATAGCTTGGTGTCTCAATTTTATATCATGTTCATTGGAGTAAATTTAACTTT

TTTTCCTCAGCATTTTCTTGGTTTGGGGGGAATGCCTCGCCGATATTCAGATTATGCTGATTGTTATCTA

GTATGGAACAAAATTTCTTCTGCGGGAAGGATTTTGAGTATCATTTCTGTTATTTATTTTTTATTTATTG

TTTTAGAATCTTTTCTTCTTTTGCGTT

>Mediterranean_France_Hap19_AM691059_AM691059.1

GAAAATTAGAGGTATTTGGAAGGTTGGGGATAATTTATGCTATATTGACTATTGGTATCTTAGGGTTTAT

TGTTTGAGGACATCATATATTTACAGTTGGAATAGATGTAGATACTCGAGCTTATTTCACTTCAGCTACT

ATGATTATTGCCGTTCCTACAGGAATTAAAATTTTTAGTTGGCTTGCTACTTTGGGTGGAATAAAGTCCA

ATAAATTCAGGCCCCTTGGCCTTTGATTTACAGGATTTTTATTTTTATTTACTATAGGTGGATTAACTGG

AATTATTCTTGGTAACTCTTCTGTAGATGTGTGTTTGCATGACACTTATTTTGTTGTTGCGCATTTTCAT

TATGTCTTATCAATAGGAATTATTTTTGCTATTGTAGGAGGAGTTATTTATTGATTTCCATTAATCTTGG

GCTTAACCTTAAATAATTATAGCTTGGTGTCTCAATTTTATATCATGTTCATTGGAGTAAATTTAACTTT

TTTTCCTCAGCATTTTCTTGGTTTGGGGGGAATGCCTCGCCGATATTCAGATTATGCTGATTGTTATCTA

GTATGGAACAAAATTTCTTCTGCGGGAAGGATTTTGAGTATCATTTCTGTTATTTATTTTTTATTTATTG

TTTTAGAATCTTTTCTTCTTTTGCGTT

>Mediterranean_China_Beijing_Hap7_GU086332_GU086332.1

GAAAATTAGAGGTATTTGGGAGGTTGGGGATAATTTATGCTATATTGACTATTGGTATCTTAGGGTTTAT

TGTTTGAGGACATCATATATTTACAGTTGGAATAGATGTAGATACTCGAGCTTATTTCACTTCAGCTACT

ATGATTATTGCCGTTCCTACAGGAATTAAAATTTTTAGTTGGCTTGCTACTTTGGGTGGAATAAAGTCCA

ATAAATTCAGGCCCCTTGGCCTTTGATTTACAGGATTTTTATTTTTATTTACTATAGGTGGATTAACTGG

AATTATTCTTGGTAACTCTTCTGTAGATGTGTGTTTGCATGACACTTATTTTGTTGTTGCGCATTTTCAT

TATGTCTTATCAATAGGAATTATTTTTGCTATTGTAGGAGGAGTTATCTATTGATTTCCATTAATCTTGG

GCTTAACCTTAAATAATTATAGCTTGGTGTCTCAATTTTATATCATGTTCATTGGAGTAAATTTAACTTT

TTTTCCTCAGCATTTTCTTGGTTTGGGGGGAATGCCTCGCCGATATTCAGATTATGCTGATTGTTATCTA

GTATGGAACAAAATTTCTTCTGCGGGAAGGATTTTGAGTATCATTTCTGTTATTTATTTTTTATTTATTG

TTTTAGAATCTTTTCTTCTTTTGCGTT

>Mediterranean_Morocco_AM176571_AM176571.1

GAAAATTAGAGGTATTTGGAAGGTTGGGGATAATTTATGCTATATTGACTATTGGTATCTTGGGGTTTAT

TGTTTGAGGACATCATATATTTACAGTTGGAATAGATGTAGATACTCGAGCTTATTTCACTTCAGCTACT

ATGATTATTGCCGTTCCTACAGGAATTAAAATTTTTAGTTGGCTTGCTACTTTGGGTGGAATAAAGTCCA

ATAAATTCAGGCCCCTTGGCCTTTGATTTACAGGATTTTTATTTTTATTTACTATAGGTGGATTAACTGG

AATTATTCTTGGTAACTCTTCTGTAGATGTGTGTTTGCATGACACTTATTTTGTTGTTGCGCATTTTCAT

TATGTCTTATCAATAGGAATTATTTTTGCTATTGTAGGAGGAGTTATCTATTGGTTTCCATTAATCTTGG

GCTTAACCTTAAATAATTATAGCTTGGTGTCTCAATTTTATATCATGTTCATTGGAGTAAATTTAACTTT

TTTTCCTCAGCATTTTCTTGGTTTGGGGGGAATGCCTCGCCGATATTCAGATTATGCTGATTGTTATCTA

GTATGGAACAAAATTTCTTCTGCGGGAAGGATTTTGAGTATCATTTCTGTTATTTATTTTTTATTTATTG

TTTTAGAATCTTTTCTTCTTTTGCGTT

>Mediterranean_Burkina_Faso_Hap5_FJ766432_FJ766432.1

GAAAATTAGAGGTATTTGGAAGGTTGGGCATAATTTATGCTATATTGACTATTGGTATCTTAGGGTTTAT

TGTTTGAGGACATCATATATTTACAGTTGGAATAGATGTAGATACTCGAGCTTATTTCACTTCAGCTACT

ATGATTATTGCCGTTCCTACAGGAATTAAAATTTTTAGTTGGCTTGCTACTTTGGGTGGAATAAAGTCCA

ATAAATTCAGGCCCCTTGGCCTTTGATTTACAGGATTTTTATTTTTATTTACTATAGGTGGATTAACTGG

AATTATTCTTGGTAACTCTTCTGTAGATGTGTGTTTGCATGACACTTATTTTGTTGTTGCGCATTTTCAT

TATGTCTTATCAATAGGAATTATTTTTGCTATTGTAGGAGGAGTTATCTATTGATTTCCATTAATCTTGG

GCTTAACCTTAAATAATTATAGCTTGGTTTCTCAATTTTATATCATGTTCATTGGAGTAAATTTAACTTT

TTTTCCTCAGCATTTTCTTGGTTTGGGGGGAATGCCTCGCCGATATTCAGACTATGCTGATTGTTATCTA

GTATGGAACAAAATTTCTTCTGCGGGAAGGATTTTGAGTATCATTTCTGTTATTTATTTTTTATTTATTG

TTTTAGAATCTTTTCTTCTTTTGCGTT

>Mediterranean_Ghana_AM040609_AM040609.1

GAAAATTAGAGGTATTTGGAAGGTTGGGCATAATTTATGCTATATTGACTATTGGTATCTTAGGGTTTAT

TGTTTGAGGACATCATATATTTACAGTTGGAATAGATGTAGATACTCGAGCTTATTTCACTTCAGCTACT

ATGATTATTGCCGTTCCTACAGGAATTAAAATTTTTAGTTGGCTTGCTACTTTGGGTGGAATAAAGTCCA

ATAAATTCAGGCCCCTTGGCCTTTGATTCACAGGATTTTTATTTTTATTTACTATAGGTGGATTAACTGG

AATTATTCTTGGTAACTCTTCTGTAGATGTGTGTTTGCATGACACTTATTTTGTTGTTGCGCATTTTCAT

TATGTCTTATCAATAGGAATTATTTTTGCTATTGTAGGAGGAGTTATCTATTGATTTCCATTAATCTTGG

GCTTAACCTTAAATAATTATAGCTTGGTTTCTCAATTTTATATCATGTTCATTGGAGTAAATTTAACTTT

TTTTCCTCAGCATTTTCTTGGTTTGGGGGGAATGCCTCGCCGATATTCAGACTATGCTGATTGTTATCTA

GTATGGAACAAAATTTCTTCTGCGGGAAGGATTTTGAGTATCATTTCTGTTATTTATTTTTTATTTATTG

TTTTAGAATCTTTTCTTCTTTTGCGTT

>Mediterranean_Czech_B1_Hap4_GU086330_GU086330.1

GAAAATTAGAGGTATTTGGAAGGTTGGGTATAATTTATGCTATATTGACTATTGGTATCTTAGGGTTTAT

TGTTTGAGGACATCATATATTTACAGTTGGAATAGATGTAGATACTCGAGCTTATTTCACTTCAGCTACT

ATGATTATTGCCGTTCCTACAGGAATTAAAATTTTTAGTTGGCTTGCTACTTTGGGTGGAATAAAGTCCA

ATAAATTCAGGCCTCTTGGCCTTTGATTTACAGGATTTTTATTTTTATTTACTATAGGTGGATTAACTGG

AATTATTCTTGGTAACTCTTCTGTAGATGTGTGTTTGCATGACACTTATTTTGTTGTTGCGCATTTTCAT

TATGTCTTATCAATAGGAATTATTTTTGCTATTGTAGGAGGAGTTATTTATTGATTTCCATTAATCTTGG

GCTTAACCTTAAATAATTATAGCTTGGTGTCTCAATTTTATATCATGTTCATTGGAGTAAATTTAACTTT

TTTTCCTCAGCATTTTCTTGGTTTGGGGGGAATGCCTCGCCGATATTCAGATTATGCTGATTGCTATCTA

GTATGGAACAAAATTTCTTCTGCGGGAAGGATTTTGAGTATTATTTCTGTTATTTATTTTTTATTTATTG

TTTTAGAATCTTTTCTTCTTTTGCGTT

>Mediterranean_Croatia_II_b_GU086335_GU086335.1

GAAAATTAGAGGTATTTGGAAGGTTGGGCATAATTTATGCTATATTGACTATTGGTATCTTAGGGTTTAT

TGTTTGAGGACATCATATATTTACAGTTGGAATAGATGTAGATACTCGAGCTTATTTCACTTCAGCTACT

ATGATTATTGCTGTTCCTACAGGAATTAAAATTTTTAGTTGGCTTGCTACTTTGGGTGGAATAAAGTCCA

ATAAATTCAGGCCCCTTGGCCTTTGATTTACAGGATTTTTATTTTTATTTACTATAGGTGGATTAACTGG

AATTATTCTTGGTAACTCTTCTGTAGATGTGTGTTTGCATGACACTTATTTTGTTGTTGCGCATTTTCAT

TATGTCTTATCAATAGGAATTATTTTTGCTATTGTAGGAGGAGTTATCTATTGATTTCCATTAATCTTGG

GTTTAACCTTAAATAATTATAGTTTGGTGTCTCAATTTTATATCATGTTCATTGGAGTAAATTTAACTTT

TTTTCCTCAGCATTTTCTTGGTTTGGGGGGAATGCCTCGCCGATATTCAGATTATGCTGATTGTTATCTA

GTATGGAACAAAATTTCTTCTGCGGGAAGGATTTTGAGTATCATTTCTGTTATTTATTTTTTATTTATTG

TTTTAGAATCTTTTCTTCTTTTGCGTT

>Mediterranean_Burkina_Faso_FJ766385_FJ766385.1

GAAAATTAGAGGTATTTGGAAGGTTGGGTATAATTTATGCTATATTGACTATTGGTATCTTAGGGTTTAT

TGTTTGAGGACATCATATATTTACAGTTGGAATAGATGTAGATACTCGAGCTTATTTCACTTCAGCTACT

ATGATTATTGCTGTTCCTACGGGAATTAAAATTTTTAGTTGGCTTGCTACTTTGGGTGGAATAAAGTCTA

ATAAATTCAGGCCTCTTGGCCTTTGATTTACGGGATTTTTATTTTTATTTACTATAGGTGGATTAACTGG

AATTATTCTTGGTAACTCTTCTGTAGATGTGTGTTTGCATGACACTTATTTTGTTGTTGCGCATTTTCAT

TATGTCTTATCAATAGGAATTATTTTTGCTATTGTAGGAGGAGTTATCTATTGATTTCCATTAATCTTGG

GTTTAACCTTAAATAATTATAGTTTGGTGTCTCAATTTTATATCATGTTTATTGGAGTAAACTTAACTTT

TTTTCCTCAGCATTTTCTTGGTTTGGGGGGAATGCCTCGCCGATATTCAGATTATGCTGATTGTTATCTA

GTATGGAACAAAATTTCTTCTGCGGGAAGGATTTTGAGTATTATTTCTGTTATTTATTTTTTATTTATTG

TTTTAGAATCTTTTCTTCTTCTGCGTT

>Mediterranean_Burkina_Faso_Hap10_FJ766400_FJ766400.1

GAAAATTAGAGGTATTTGGAAGGTTGGGTATAATTTATGCTATATTGACTATTGGTATCTTAGGGTTTAT

TGTTTGAGGACATCATATATTTACAGTTGGAATAGATGTAGATACTCGAGCTTATTTCACTTCAGCTACT

ATGATTATTGCTGTTCCTACGGGAATTAAAATTTTTAGTTGGCTTGCTACTTTGGGTGGAATAAAGTCTA

ATAAATTCAGGCCTCTTGGCCTTTGATTTACAGGATTTTTATTTTTATTTACTATAGGTGGATTAACTGG

AATTATTCTTGGTAACTCTTCTGTAGATGTGTGTTTGCATGACACTTATTTTGTTGTTGCGCATTTTCAT

TATGTCTTATCAATAGGAATTATTTTTGCTATTGTAGGAGGAGTTATCTATTGATTTCCATTAATCTTGG

GTTTAACCTTAAATAATTATAGTTTGGTGTCTCAATTTTATATCATGTTTATTGGAGTAAACTTAACTTT

TTTTCCTCAGCATTTTCTTGGTTTGGGGGGAATGCCTCGCCGATATTCAGATTATGCTGATTGTTATCTA

GTATGGAACAAAATTTCTTCTGCGGGAAGGATTTTGAGTATTATTTCTGTTATTTATTTTTTATTTATTG

TTTTAGAATCTTTTCTTCTTCTGCGTT

>Mediterranean_Burkina_Faso_Hap12_FJ766405_FJ766405.1

GAAAATTAGAGGTATTTGGAAGGTTGGGTATAATTTATGCTATATTGACTATTGGTATCTTAGGGTTTAT

TGTTTGAGGACATCATATATTTACAGTTGGAATAGATGTAGATACTCGAGCTTATTTCACTTCAGCTACT

ATGATTATTGCTGTTCCTACGGGAATTAAAATTTTTAGTTGGCTTGCTACTTTGGGTGGAATAAAGTCTA

ATAAATTCAGGCCTCTTGGCCTTTGATTTACAGGATTTTTATTTTTATTTACTATAGGTGGATTAACTGG

AATTATTCTTGGTAACTCTTCTGTAGATGTGTGTTTGCATGACACTTATTTTGTTGTTGCGCATTTTCAT

TATGTTTTATCAATAGGAATTATTTTTGCTATTGTAGGAGGAGTTATCTATTGATTTCCATTAATCTTGG

GTTTAACCTTAAATAATTATAGTTTGGTGTCTCAATTTTATATCATGTTTATTGGAGTAAACTTAACTTT

TTTTCCTCAGCATTTTCTTGGTTTGGGGGGAATGCCTCGCCGATATTCAGATTATGCTGATTGTTATCTA

GTATGGAACAAAATTTCTTCTGCGGGAAGGATTTTGAGTATTATTTCTGTTATTTATTTTTTATTTATTG

TTTTAGAATCTTTTCTTCTTCTGCGTT

>Mediterranean_Burkina_Faso_Hap11_FJ766387_FJ766387.1

GAAAATTAGAGGTATTTGGAAGGTTGGGTATAATTTATGCTATATTGACTATTGGTATCTTAGGGTTTAT

TGTTTGAGGACATCATATATTTACAGTTGGAATAGATGTAGATACTCGAGCTTATTTCACTTCAGCTACT

ATGATTATTGCTGTTCCTACGGGAATTAAAATTTTTAGTTGGCTTGCTACTTTGGGTGGAATAAAGTCTA

ATAAATTCAGGCCTCTTGGCCTTTGATTTACAGGATTTTTATTTTTATTTACTATAGGTGGATTAACTGG

AATTATTCTTGGTAACTCTTCTGTAGATGTGTGTTTGCATGACACTTATTTTGTTGTTGCGCATTTTCAT

TATGTTTTATCAATAGGAATTATTTTTGCTATTGTAGGAGGAGTTATCTATTGATTTCCATTAATCTTGG

GTTTAACCTTAAATAATTATAGTTTGGTGTCTCAATTTTATATCATGTTTATTGGAGTAAACTTAACTTT

TTTTCCTCAGCATTTTCTTGGTTTGGGGGGAATGCCTCGCCGATATTCAGATTATGCTGATTGTTATCTA

GTATGGAACAAAATTTCTTCTGCGGGAAGGATTTTGAGTATTATTTCTGTTATTTATTTTTTATTTATTG

TTTTAGAATCTTTTCTTCTTCTTCGTT

>Mediterranean_Spain_Almeria_DQ302946_DQ302946.1

GAAAATTAGAGGTATTTGGAAGGTTGGGGATAATTTATGCTATATTGACTATTGGTATCTTAGGGTTTAT

TGTTTGAGGACATCATATATTTACAGTTGGAATAGATGTAGATACTCGAGCTTATTTCACTTCAGCTACT

ATGATTATTGCCGTTCCTACAGGAATTAAAATTTTTAGCTGGCTTGCTACTTTGGGTGGAATAAAGTCCA

ATAAATTCAGGCCCCTTGGCCTTTGATTTACAGGATTTTTATTTTTATTTACTATAGGTGGATTAACTGG

AATTATTCTTGGTAACTCTTCTGTAGATGTGTGTTTGCATGACACTTATTTTGTTGTTGCGCATTTTCAT

TATGTCTTATCAATAGGAATTATTTTTGCTATTGTAGGAGGAGTTATCTATTGATTTCCATTAATCTTGG

GCTTAACCTTAAATAATTATAGCTTGGTGTCTCAATTTTATATCATGTTCATTGGAGTAAATTTAACTTT

TTTTCCTCAGCATTTTCTTGGTTTGGGGGGAATGCCTCGCCGATATTCAGATTATGCTGATTGTTATCTA

GTATGGAACAAAATTTCTTCTGCGGGAAGGATTTTGAGTATCATTTCTGTTATTTATTTTTTATTTATTG

TTTTAGAATCTTTTCTTCTTTTGCGTN

>Mediterranean_Burkina_Faso_Hap9_FJ766391_FJ766391.1

GAAAATTAGAGGTATTTGGAAGGTTGGGTATAATTTATGCTATATTGACTATTGGTATCTTAGGGTTTAT

TGTTTGAGGACATCACATATTTACAGTTGGAATAGATGTAGATACTCGAGCTTACTTCACTTCAGCTACT

ATGATTATTGCCGTTCCTACAGGAATTAAAATTTTTAGTTGGCTTGCTACTTTGGGTGGAATAAAGTCTA

ATAAATTTAGGCCTCTTGGCCTTTGATTTACAGGATTTTTATTTTTATTTACTATAGGTGGATTAACTGG

AATTATTCTTGGTAACTCTTCTGTAGATGTATGTTTGCATGACACTTATTTTGTTGTTGCGCATTTTCAT

TATGTCTTATCAATAGGAATTATTTTTGCTATTGTAGGAGGAGTTATCTATTGATTTCCATTAATCTTGG

GTTTAACCTTAAATAACTATAGTTTGGTGTCTCAATTTTATATCATGTTCATTGGAGTAAACTTAACTTT

TTTTCCTCAGCATTTTCTTGGTTTGGGGGGAATGCCTCGCCGATATTCAGATTATGCTGATTGTTATCTA

GTATGGAACAAGATTTCTTCTGCGGGAAGGATTTTGAGTATCATTTCTGTTATTTATTTTTTATTTATTG

TTTTAGAATCTTTTCTTCTTCTGCGTT

>Mediterranean_IvoryCoast_Hap6_SubSahAfSL_AY057136_AY057136.1

GAAAATTAGAGGTATTTGGAAGGTTGGGTATAATTTATGCTATATTGACTATTGGTATCTTAGGGTTTAT

TGTTTGAGGACATCACATATTTACAGTTGGAATAGATGTAGATACTCGAGCTTACTTCACTTCAGCTACT

ATGATTATTGCCGTTCCTACAGGAATTAAAATTTTTAGTTGGCTTGCTACTTTGGGTGGAATAAAGTCTA

ATAAATTTAGGCCTCTTGGCCTTTGATTTACAGGATTTTTATTTTTATTTACTATAGGTGGATTAACTGG

AATTATTCTTGGTAACTCTTCTGTAGATGTATGTTTGCATGACACTTATTTTGTTGTTGCGCATTTTCAT

TATGTCTTATCAATAGGAATTATTTTTGCTATTGTAGGAGGAGTTATCTATTGATTTCCATTAATCTTGG

GTTTAACCTTAAATAACTATAGTTTGGTGTCTCAATTTTATATCATGTTCATTGGAGTAAACTTAACTTT

TTTTCCTCAGCATTTTCTTGGCTTGGGGGGAATGCCTCGCCGATATTCAGATTATGCTGATTGTTATCTA

GTATGGAACAAGATTTCTTCTGCGGGAAGGATTTTGAGTATCATTTCTGTTATTTATTTTTTATTTATTG

TTTTAGAATCTTTTCTTCTTCTGCGTT

>Mediterranean_Ivory_Coast_FJ766383_FJ766383.1

GAAAATTAGAGGTATTTGGAAGGTTGGGTATAATTTATGCTATATTGACTATTGGTATCTTAGGGTTTAT

TGTTTGAGGACATCACATATTTACAGTTGGAATAGATGTAGATACTCGAGCTTACTTCACTTCAGCTACT

ATGATTATTGCTGTTCCTACAGGAATTAAAATTTTTAGTTGGCTTGCTACTTTGGGTGGAATAAAGTCTA

ATAAATTTAGGCCTCTTGGCCTTTGATTTACAGGATTTTTATTTTTATTTACTATAGGTGGATTAACTGG

AATTATTCTTGGTAACTCTTCTGTAGATGTATGTTTGCATGACACTTATTTTGTTGTTGCGCATTTTCAT

TATGTCTTATCAATAGGAATTATTTTTGCTATTGTAGGAGGAGTTATCTATTGATTTCCATTAATCTTGG

GTTTAACCTTAAATAACTATAGTTTGGTGTCTCAATTTTATATCATGTTCATTGGAGTAAACTTAACTTT

TTTTCCTCAGCATTTTCTTGGTTTGGGGGGAATGCCTCGCCGATATTCAGATTATGCTGATTGTTATCTA

GTATGGAACAAGATTTCTTCTGCGGGAAGGATTTTGAGTATCATTTCTGTTATTTATTTTTTATTTATTG

TTTTAGAATCTTTTCTTCTTCTGCGTT

>Mediterranean_Ghana_SubSahAfSL_AY827589_AY827589.1

GAAAATTAGAGGTATTTGGAAGGTTGGGTATAATTTATGCTATATTGACTATTGGTATCTTAGGGTTTAT

TGTTTGAGGACATCATATATTTACAGTTGGAATAGATGTAGATACTCGAGCTTACTTCACTTCAGCTACT

ATGATTATTGCCGTTCCTACAGGAATTAAAATTTTTAGTTGGCTTGCTACTTTGGGTGGAATAAAGTCTA

ATAAATTTAGGCCTCTTGGCCTTTGATTTACAGGATTTTTATTTTTATTTACTATAGGTGGATTAACTGG

AATTATTCTTGGTAACTCTTCTGTAGATGTATGTTTGCATGACACTTATTTTGTTGTTGCGCATTTTCAT

TATGTCTTATCAATAGGAATTATTTTTGCTATTGTAGGAGGAGTTATCTATTGATTTCCATTAATCTTGG

GTTTAACCTTAAATAACTATAGTTTGGTGTCTCAATTTTATATCATGTTCATTGGAGTAAACTTAACTTT

TTTTCCTCAGCATTTTCTTGGTTTGGGGGGAATGCCTCGCCGATATTCAGATTATGCTGATTGTTATCTA

GTATGGAACAAGATTTCTTCTGCGGGAAGGATTTTGAGTATCATTTCCGTTATTTATTTTTTATTTATTG

TTTTAGAATCTTTTCTTCTTCTGCGTT

>Mediterranean_Ghana_SubSahAfSL_AY827588_AY827588.1

GAAAATTAGAGGTATTTGGAAGGTTGGGTATAATTTATGCTATATTGACTATTGGTATCTTAGGGTTTAT

TGTTTGAGGACATCACATATTTACAGTTGGAATAGATGTAGATACTCGAGCTTACTTCACTTCAGCTACT

ATGATTATTGCCGTTCCCACAGGAATTAAAATTTTTAGTTGGCTTGCTACTTTGGGTGGAATAAAGTCTA

ATAAATTTAGGCCTCTTGGCCTTTGATTTACAGGATTTTTATTTTTATTTACTATAGGTGGATTAACTGG

AATTATTCTTGGTAACTCTTCTGTAGATGTATGTTTGCATGACACTTATTTTGTTGTTGCGCATTTTCAT

TATGTCTTATCAATAGGAATTATTTTTGCTATTGTAGGAGGAGTTATCTATTGATTTCCATTAATCTTGG

GTTTAACCTTAAATAACTATAGTTTGGTGTCTCAATTTTATATCATGTTCATTGGAGTAAACTTAACTTT

TTTTCCTCAGCATTTTCTTGGCTTGGGGGGAATGCCTCGCCGATATTCAGATTATGCTGATTGTTATCTA

GTATGGAACAAGATTTCTTCTGCGGGAAGGATTCTGAGTATCATTTCTGTTATTTATTTTTTATTTATTG

TTTTAGAATCTTTTCTTCTTCTGCGTT

>Mediterranean_Ghana_SubSahAfSL_AY827590_AY827590.1

GAAAATTAGAGGTATTTGGAAGGTTGGGTATAATTTATGCTATATTGACTATTGGCATCTTAGGGTTTAT

TGTTTGAGGACATCACATATTTACAGTTGGAATAGATGTAGATACTCGAGCTTACTTCACTTCAGCTACT

ATGATTATTGCCGTTCCCACAGGAATTAAAATTTTTAGTTGGCTTGCTACTTTGGGTGGAATAAAGTCTA

ATAAATTTAGGCCTCTTGGCCTTTGATTTACAGGATTTTTATTTTTATTTACTATAGGTGGATTAACTGG

AATTATTCTTGGTAACTCTTCTGTAGATGTATGTTTGCATGACACTTATTTTGTTGTTGCGCATTTTCAT

TATGTCTTATCAATAGGAATTATTTTTGCTATTGTAGGAGGAGTTATCTATTGATTTCCATTAATCTTGG

GTTTAACCTTAAATAACTATAGTTTGGTGTCTCAATTTTATATCATGTTCATTGGAGTAAACTTAACTTT

TTTTCCTCAGCATTTTCTTGGTTTGGGGGGAATGCCTCGCCGATATTCAGATTATGCTGATTGTTATCTA

GTATGGAACAAGATTTCTTCTGCGGGAAGGATTCTGAGTATCATTTCTGTTATTTATTTTTTATTTATTG

TTTTAGAATCTTTTCTTCTTCTGCGTT

>Mediterranean_Uganda_Hap24_AY903540_AY903540.1

GAAAATTAGAGGTATTTGGAAGGTTGGGTATAATTTATGCTATATTGACTATTGGTATTTTAGGGTTTAT

TGTTTGAGGACATCACATATTTACAGTTGGAATAGATGTAGATACTCGAGCTTACTTCACTTCAGCTACT

ATGATTATTGCCGTTCCTACAGGAATTAAAATTTTTAGTTGGCTTGCTACTTTGGGTGGAATAAAGTCTA

ATAAATTTAGGCCTCTTGGCCTTTGATTTACAGGATTTTTATTTTTATTTACTATAGGTGGATTAACTGG

GATTATTCTTGGTAACTCTTCTGTAGATGTATGTTTGCATGACACTTATTTTGTTGTTGCACATTTTCAT

TATGTCTTATCAATAGGAATTATTTTTGCTATTGTAGGAGGAGTTATCTATTGATTTCCATTAATCTTGG

GTTTAACCTTAAATAACTATAGTTTGGTGTCTCAATTTTATATCATGTTCATTGGAGTAAACTTAACTTT

TTTTCCTCAGCATTTTCTTGGTTTGGGGGGAATGCCTCGCCGATATTCGGATTACGCTGATTGTTATCTA

GTATGGAACAAGATTTCTTCTGCGGGAAGGATTTTGAGTATCATTTCTGTTATTTATTTTTTATTTATTG

TTTTAGAATCTTTTCTTCTTCTGCGTT

>Mediterranean_Uganda_Namulonge_AY903531_AY903531.1

GAAAATTAGAGGTATTTGGAAGGTTGGGTATAATTTATGCTATATTGACTATTGGTATTTTAGGGTTTAT

TGTTTGAGGACATCACATATTTACAGTTGGAATAGATGTAGATACTCGAGCTTATTTCACTTCAGCTACT

ATGATTATTGCTGTTCCTACAGGAATTAAAATTTTTAGTTGGCTTGCTACTTTGGGTGGAATAAAGTCTA

ATAAATTTAGGCCTCTTGGCCTTTGATTTACAGGATTTTTATTTTTATTTACTATAGGTGGATTAACTGG

AATTATTCTTGGTAACTCTTCTGTAGATGTATGTTTGCATGACACTTATTTTGTTGTTGCGCATTTTCAT

TATGTCTTATCAATAGGAATTATTTTTGCTATTGTAGGAGGAGTTATCTATTGATTTCCATTAATCTTGG

GTTTAACCTTAAATAACTATAGTTTGGTGTCTCAATTTTATATCATGTTCATTGGAGTAAACTTAACTTT

TTTTCCTCAGCATTTTCTTGGTTTGGGAGGAATGCCTCGCCGATATTCAGATTATGCTGATTGTTATCTA

GTATGGAACAAGATTTCTTCTGCGGGAAGGATTTTGAGTATCATTTCTGTTATTTATTTTTTATTTATTG

TTTTAGAATCCTTTCTTCTTCTGCGTT

>Mediterranean_Uganda_Mukono_AY903556_AY903556.1

GTAAATTAGAGGTATTTGGAAGGTTGGGTATAATTTATGCTATATTGACTATTGGTATTTTAGGGTTTAT

TGTTTGAGGACATCACATATTTACAGTTGGAATAGATGTAGATACTCGAGCTTACTTCACTTCAGCTACT

ATGATTATTGCTGTTCCTACAGGAATTAAAATTTTTAGTTGGCTTGCTACTTTGGGTGGAATAAAGTCTA

ATAAATTTAGGCCTCTTGGCCTTTGATTTACAGGATTTTTATTTTTATTTACTATAGGTGGATTAACTGG

AATTATTCTTGGTAACTCTTCTGTAGATGTATGTTTGCATGACACTTATTTTGTTGTTGCGCATTTTCAT

TATGTCTTATCAATAGGAATTATTTTTGCTATTGTAGGAGGAGTTATCTATTGATTTCCATTAATCTTGG

GTTTAACCTTAAATAACTATAGTTTGGTGTCTCAATTTTATATCATGTTCATTGGAGTAAACTTAACTTT

TTTTCCTCAGCATTTTCTTGGTTTGGGAGGAATGCCTCGCCGATATTCAGATTATGCTGATTGTTATCTA

GTATGGAACAAGATTTCTTCTGCGGGAAGGATTTTGAGTATCATTTCTGTTATTTATTTTTTATTTATTG

TTTTAGAATCCTTTCTTCTTCTGCGTT

>NewWorld2_Brazil_Bastos_JN689351_JN689351.1 Bemisia tabaci isolate Selviria haplotype BR-2 cytochrome oxidase subunit I (COI) gene, partial cds; mitochondrial

GAAAATTAGAAGTATTTGGTAGGTTGGGCATAATTTATGCCATGATAACTATTGGCATTTTGGGATTTAT

TGTTTGGGGCCATCACATGTTTACTGTCGGAATAGATGTTGACACCCGAGCTTATTTTACCTCAGCTACT

ATAGTTATTGCTGTTCCGACGGGAATTAAAATTTTTAGGTGGCTTGCTACTCTAGGTGGAATAAAGTCTA

ATAAGTTTAGACCCCTAGTTCTTTGATTTACAGGATTTCTATTTTTATTTACTATGGGTGGATTAACTGG

AATTATTCTTGGTAATTCCTCGGTAGATGTGTGTCTTCATGATACCTACTTTGTTGTTGCTCATTTTCAT

TATGTTTTATCGATAGGAATTATCTTTGCTATTATAGGCGGAGTAATTTATTGATTTCCTTTAATTTTAG

GCTTAACTCTAAACAATTATAATTTGGTGGCTCAATTTTATATAATATTTGTGGGGGTAAACCTGACATT

TTTTCCACAGCATTTTCTTGGTTTAAGTGGAATGCCTCGTCGGTATTCTGATTATCCTGATTGTTATTTA

CTATGAAATAAAATTTCCTCTGGGGGGAGGGTCTTGAGCGTTATTTCTGTTATTTATTTTTTATTTATTA

TTTTAGAATCTTTTCTTCTTTTACGGC

>NewWorld2_Brazil_Ilha_Solteira_JN689354_JN689354.1 Bemisia tabaci isolate Bastos haplotype BR-4 cytochrome oxidase subunit I (COI) gene, partial cds; mitochondrial

GAAAATTAGAAGTATTTGGTAGGTTGGGCATAATTTATGCCATGATAACTATTGGCATTTTGGGATTTAT

TGTTTGGGGCCATCACATGTTTACTGTCGGAATAGATGTTGACACCCGAGCTTATTTTACCTCAGCTACT

ATAGTTATTGCTGTTCCGACGGGAATTAAAATTTTTAGGTGGCTTGCTACTCTAGGTGGAATAAAGTCTA

ATAAGTTTAGACCCCTAGTTCTCTGATTTACAGGATTTCTATTTTTATTTACTATGGGTGGATTAACTGG

AATTATTCTTGGTAATTCCTCGGTAGATGTGTGTCTTCACGATACCTACTTTGTTGTTGCTCATTTTCAT

TATGTTTTATCGATAGGAATTATCTTTGCTATTATAGGCGGAGTAATTTATTGATTTCCTTTAATTTTAG

GCTTAACTCTAAACAATTATAATTTGGTGGCTCAATTTTATATAATATTTGTGGGGGTAAACCTGACATT

TTTTCCACAGCATTTTCTTGGTTTAAGTGGAATGCCTCGTCGGTATTCTGATTATCCTGATTGTTATTTA

CTATGAAATAAAATTTCCTCTGGGGGGAGGGTCTTGAGCGTTATTTCTGTTATTTATTTTTTATTTATTA

TTTTAGAATCTTTTCTTCTTTTACGGC

>NewWorld2_Brazil_Bastos_JN689350_JN689350.1 Bemisia tabaci isolate Ilha Solteira haplotype BR-1 cytochrome oxidase subunit I (COI) gene, partial cds; mitochondrial

GCAAATTAGAAGTATTTGGTAGGCTGGGTATAATTTATGCCATGATAACTATTGGCATTTTGGGATTTAT

TGTTTGGGGTCATCACATGTTTACTGTCGGAATAGATGTTGACACCCGGGCTTATTTTACCTCAGCTACC

ATAGTTATTGCTGTTCCGACGGGAATTAAAATTTTTAGGTGGCTTGCTACTCTGGGTGGAATAAAGTCTA

ATAAGTTTAGACCCCTAGTTCTCTGGTTTACAGGATTTCTATTTTTATTTACTATGGGTGGATTAACTGG

AATTATTCTTGGTAATTCCTCGGTAGATGTGTGTCTTCATGATACTTACTTTGTTGTTGCTCATTTTCAT

TATGTTTTATCGATAGGAATTATCTTTGCTATTATAGGCGGAGTAATTTATTGATTTCCTTTAATTTTAG

GCTTAACTTTAAACAATTATAATTTGGTGTCTCAATTTTATATAATATTTGTGGGAGTAAACCTGACATT

TTTTCCACAGCATTTTCTTGGTTTAAGTGGAATGCCTCGTCGGTATTCTGATTATCCTGATTGTTATTTA

TTATGAAATAAAATTTCCTCTGGGGGGAGGGTCTTGAGTGTTATTTCTGTTATTTACTTTTTATTTATTA

TTTTAGAATCTTTTCTTCTCTTACGGC

>NewWorld2_Brazil_Bastos_JN689352_JN689352.1 Bemisia tabaci isolate Sao Jose do Rio Preto haplotype BR-3 cytochrome oxidase subunit I (COI) gene, partial cds; mitochondrial

GCAAATTAGAAGTATTTGGTAGGCTGGGTATAATTTATGCCATGATAACTATTGGCATTTTGGGATTTAT

TGTTTGGGGCCATCACATGTTTACTGTCGGAATAGATGTTGACACCCGGGCTTATTTTACCTCAGCTACT

ATAGTTATTGCTGTTCCGACGGGAATTAAAATTTTTAGGTGGCTTGCTACTCTGGGTGGAATAAAGTCTA

ATAAGTTTAGACCCCTAGTTCTCTGGTTTACAGGATTTCTATTTTTATTTACTATGGGTGGATTAACTGG

AATTATCCTTGGTAATTCCTCGGTAGATGTGTGTCTTCACGATACTTACTTTGTTGTTGCTCATTTTCAT

TATGTTTTATCGATAGGAATTATCTTTGCTATTATAGGCGGAGTAATTTATTGATTTCCTTTAATTTTAG

GCCTAACTCTAAACAATTATAATTTGGTGTCTCAATTTTATATAATATTTGTGGGAGTAAACTTGACATT

TTTTCCACAGCATTTTCTTGGTTTAAGTGGGATGCCTCGTCGGTATTCTGATTATCCTGATTGTTATTTA

TTATGAAATAAAATTTCCTCTGGGGGGAGGGTCTTGAGTGTTATTTCTGTTATTTACTTTTTATTTATTA

TTTTAGAATCTTTTCTTCTCTTACGGT

>NewWorld2_Brazil_Selviria_JN689355_JN689355.1 Bemisia tabaci isolate Bastos haplotype BR-5 cytochrome oxidase subunit I (COI) gene, partial cds; mitochondrial

GCAAATTAGAAGTATTTGGTAGGCTGGGTATAATTTATGCCATGATAACTATTGGCATTTTGGGATTTAT

TGTTTGGGGCCATCACATGTTTACTGTTGGAATAGATGTTGACACCCGGGCTTATTTTACCTCAGCTACT

ATAGTTATTGCTGTTCCGACGGGAATTAAAATTTTTAGGTGGCTTGCTACTCTTGGTGGAATAAAGTCTA

ATAAGTTTAGACCCCTAGTTCTCTGGTTTACAGGATTTCTATTTTTATTTACTATGGGTGGATTAACTGG

AATTATTCTTGGTAATTCCTCGGTAGATGTGTGTCTTCACGATACTTACTTTGTTGTTGCTCATTTTCAT

TATGTTTTATCGATAGGAATTATCTTTGCTATTATAGGCGGAGTAATTTATTGATTCCCTTTAATTTTAG

GCTTAACTCTAAACAATTATAATTTGGTGTCTCAATTTTATATAATATTTGTGGGAGTAAACCTGACATT

TTTTCCACAGCATTTTCTTGGTTTAAGTGGGATGCCTCGTCGGTATTCTGATTATCCTGATTGTTATTTA

TTATGAAATAAAATTTCCTCTGGGGGGAGGGTCTTGAGTGTTATTTCTGTTATTTACTTTTTATTTATTA

TTTTAGAATCTTTTCTTCTCTTACGGC

>NewWorld2-2_ Argentina_JF901839_JF901839.1 Bemisia tabaci isolate AR7 cytochrome c oxidase subunit 1 (CO1) gene, partial cds; mitochondrial

GAAAATTAGAAGTATTTGGTAGGTTGGGCATAATTTATGCCATGATAACTATTGGCATTTTGGGATTTATTGTTTGGGGCCATCACATGTTTACTGTCGGAATAGATGTTGACACCCGAGCTTATTTTACCTCAGCTACTATAGTTATTGCTGTTCCGACGGGAATTAAAATTTTTAGGTGGCTTGCTACTCTAGGTGGAATAAAGTCTAATAAGTTTAGACCCCTAGTTCTCTGATTTACAGGATTTCTATTTTTATTTACTATGGGTGGATTAACTGGAATTATTCTTGGTAATTCCTCGGTAGATGTGTGTCTTCACGATACCTACTTTGTTGTTGCTCATTTTCATTATGTTTTATCGATAGGAATTATCTTTGCTATTATAGGCGGAGTAATTTATTGATTTCCTTTAATTTTAGGCTTAACTCTAAACAATTATAATTTGGTGGCTCAATTTTATATAATATTTGTGGGGGTAAACCTGACATTTTTTCCACAGCATTTTCTTGGTTTAAGTGGAATGCCTCGTCGGTATTCTGATTATCCTGATTGTTATTTACTATGAAATAAAATTTCCTCTGGGGGGAGGGTCTTGAGCGTTATTTCTGTTATTTATTTTTTATTTATTATTT

>NewWorld2-2_Argentina_JF901844_JF901844.1_Bemisia tabaci isolate AR12 cytochrome c oxidase subunit 1 (CO1) gene, partial cds; mitochondrial

GAAAATTAGAAGTATTTGGTAGGTTGGGCATAATTTATGCCATGATAACTATTGGCATTTTGGGATTTATTGTTTGGGGCCATCACATGTTTACTGTCGGAATAGATGTTGACACCCGAGCTTATTTTACCTCAGCTACTATAGTTATTGCTGTTCCGACGGGAATTAAAATTTTTAGGTGGCTTGCTACTCTAGGTGGAATAAAGTCTAATAAGTTTAGACCCCTAGTTCTCTGATTTACAGGATTTCTATTTTTATTTACTATGGGTGGATTAACTGGAATTATTCTTGGTAATTCCTCGGTAGATGTGTGTCTTCACGATACCTACTTTGTTGTTGCTCATTTTCATTATGTTTTATCGATAGGAATTATCTTTGCTATTATAGGCGGAGTAATTTATTGATTTCCTTTAATTTTAGGCTTAACTCTAAACAATTATAATTTGGTGGCTCAATTTTATATAATATTTGTGGGGGTAAACCTGACATTTTTTCCACAGCATTTTCTTGGTTTAAGTGGAATGCCTCGTCGGTATTCNGATTATCCTGATTGTTATTTACTATGAAATAAAATTTCCTCTGGGGGGAGGGTCTTGAGCGTTATTTCTGTTATTTATTTTTTATTTATTATTT

>NewWorld1_Belize_DQ130053_DQ130053.1 Bemisia tabaci isolate 1 from Belize cytochrome oxidase subunit I (COI) gene, partial cds; mitochondrial

GAAAATTAGAAGTATTTGGTAGGCTGGGCATAATTTATGCCATGATAACTATTGGTATCTTGGGATTTAT

TGTTTGGGGACATCACATGTTTACTGTTGGAATAGATGTTGACACTCGGGCTTATTTTACTTCAGCTACT

ATAGTTATTGCTGTTCCGACGGGAATTAAAATCTTTAGGTGGCTTGCTACTCTAGGTGGAATAAAGTCTA

ATAAGTTTAGACCCCTAGTTCTCTGATTTACAGGATTTCTATTTTTATTTACTATGGGCGGATTAACTGG

AATTATTCTTGGTAATTCTTCGGTAGATGTGTGTCTTCATGATACTTATTTTGTTGTTGCTCATTTTCAT

TATGTTTTATCTATAGGAATTATCTTTGCTATTATAGGCGGAGTAATTTATTGATTTCCTTTAATTTTAG

GTCTAACTTTAAACAATTATAATCTGGTGTCTCAATTTTATATAATATTTGTGGGAGTAAATCTGACATT

TTTTCCACAGCATTTTCTTGGTTTAAGTGGAATACCTCGTCGGTATTCTGACTACCCTGATTGTTATCTA

CTATGAAATAAAATTTCCTCTGGGGGGAGGGTCTTAAGTGTTATTTCTGTTATTTATTTTTTATTTATTA

TTTTAGAGTCTTTTCTTCTCTTACGGC

>NewWorld1_Guatemala_AY057129_AY057129.1 Bemisia tabaci clone tom GT 3 cytochrome oxidase subunit I (COI) gene, partial cds; mitochondrial gene for mitochondrial product

GAAAATTAGAAGTATTTGGTAGCCTGGGCATAATTTATGCCATGATAACTATTGGTATCTTGGGATTTAT

TGTTTGGGGACATCACATGTTTACTGTTGGAATAGATGTTGACACTCGGGCTTATTTTACTTCAGCTACT

ATAGTTATTGCTGTTCCGACGGGAATTAAAATCTTTAGGTGGCTTGCTACTCTAGGTGGAATAAAGTCTA

ATAAGTTTAGACCCCTAGTTCTCTGATTTACAGGATTTCTATTTTTATTTACTATGGGTGGATTAACTGG

AATTATTCTTGGTAATTCTTCGGTAGATGTGTGTCTTCATGATACTTATTTTGTTGTTGCTCATTTTCAT

TATGTTTTATCTATAGGAATTATCTTTGCTATTATAGGCGGAGTAATTTATTGATTTCCTTTAATTTTAG

GTCTAACTTTAAACAATTATAACCTGGTGTCTCAATTTTATATAATATTTGTGGGAGTAAACCTGACATT

TTTTCCACAGCATTTTCTTGGTTTAAGTGGAATACCTCGTCGGTATTCTGACTATCCTGATTGTTATTTA

CTATGAAATAAAATTTCCTCTGGGGGGAGGGTCTTAAGTGTTATTTCTGTTATTTATTTTTTATTTATTA

TTTTAGAGTCTTTTCTTCTCTTACGGC

>NewWorld1_USA_A_biotype_AY521259_AY521259.2 Bemisia tabaci mitochondrion, complete genome

GAAAATTAGAAGTATTTGGTAGCCTGGGTATAATTTATGCCATGATAACTATTGGTATCTTGGGATTTAT

TGTTTGGGGACATCACATGTTTACTGTTGGAATAGATGTTGACACTCGGGCTTATTTTACTTCAGCTACT

ATAGTTATTGCTGTTCCGACGGGAATTAAAATCTTTAGGTGGCTTGCTACTCTAGGTGGAATAAAGTCTA

ATAAGTTTAGACCCCTAGTTCTCTGATTTACAGGATTTCTATTTTTATTTACTATGGGTGGATTAACTGG

AATTATTCTTGGTAATTCTTCGGTAGATGTGTGTCTTCATGATACTTATTTTGTTGTTGCTCATTTTCAT

TATGTTTTATCTATAGGAATTATCTTTGCTATTATAGGCGGAGTAATTTATTGATTTCCTTTAATTTTAG

GTCTAACTTTAAACAATTATAACCTGGTGTCTCAATTTTATATAATATTTGTGGGAGTAAACCTGACATT

TTTTCCACAGCATTTTCTTGGTTTAAGTGGAATACCTCGTCGGTATTCTGACTACCCTGATTGTTATTTA

CTATGAAATAAAATTTCCTCTGGGGGGAGGGTCTTAAGTGTTATTTCTGTTATTTATTTTTTATTTATTA

TTTTAGAGTCTTTTCTTCTCTTACGGC

>NewWorld1_Mexico_EU427729_EU427729.1 Bemisia tabaci pop-variant CLM133-NW cytochrome oxidase subunit 1-like gene, partial sequence; mitochondrial

GAAAATTAGAAGTATTTGGTAGTCTGGGCATAATTTATGCCATGATAACTATTGGTATCTTGGGATTTAT

TGTTTGGGGACATCATATGTTTACTGTTGGAATAGATGTTGACACTCGGGCTTATTTTACTTCAGCTACT

ATAGTTATTGCTGTTCCGACGGGAATTAAAATCTTTAGGTGGCTTGCTACTCTAGGTGGAATAAAGTCTA

ATAAGTTTAGACCCCTAGTTCTCTGATTTACAGGATTTCTATTTTTATTTACTATGGGTGGATTAACTGG

AATTATTCTTGGTAATTCTTCGGTAGATGTGTGTCTTCATGATACTTATTTTGTTGTTGCTCATTTTCAT

TATGTTTTATCTATAGGAATTATCTTTGCTATTATGGGCGGAGTAATTTATTGATTTCCTTTAATTTTAG

GTCTAACTTTAAACAATTATAACCTGGTGTCTCAATTTTATATAATATTTGTGGGAGTAAACCTGACATT

TTTTCCACAGCATTTTCTTGGTTTAAGTGGAATACCTCGTCGGTATTCTGACTACCCTGATTGTTATCTA

CTATGAAATAAAATTTCCTCTGGGGGGAGGGTCTTAAGTGTTATTTCTGTTATTTATTTTTTATTTATTA

TTTTAGAGTCTTTTCTTCTCTTACGGC

>NewWorld1_Panama_DQ130061_DQ130061.1 Bemisia tabaci isolate 12 from Panama cytochrome oxidase subunit I (COI) gene, partial cds; mitochondrial

GAAAATTAGAAGTATTTGGTAGGCTGGGCATAATTTATGCCATGATAACTATTGGTATCTTGGGATTTAT

TGTTTGGGGACATCACATGTTTACTGTTGGAATAGATGTTGACACTCGGGCTTATTTTACTTCAGCTACT

ATAGTTATTGCTGTTCCGACGGGAATTAAAATCTTTAGGTGGCTTGCTACTCTAGGTGGAATAAAGTCTA

ATAAGTTTAGACCCCTAGTTCTTTGATTTACAGGATTTCTATTTTTATTTACTATGGGTGGATTAACTGG

AATTATTCTTGGTAATTCTTCGGTAGATGTGTGTCTTCATGATACTTATTTTGTTGTTGCTCATTTTCAT

TATGTTTTATCTATAGGAATTATCTTTGCTATTATAGGCGGAGTAATTTATTGATTTCCTTTAATTTTAG

GTCTAACTTTAAACAATTATAACCTGGTGTCTCAATTTTATATAATATTTGTGGGAGTAAACCTGACATT

TTTTCCCCAGCATTTTCTTGGTTTAAGTGGGATACCTCGTCGGTATTCTGACTACCCTGATTGTTATTTA

CTATGAAATAAAATTTCCTCTGGGGGGAGGGTCTTAAGTGTTATTTCTGTTATTTATTTTTTATTTATTA

TCTTAGAGTCTTTTCTTCTCTTACGGC

>NewWorld1_Sudan_EU760727_EU760727.1 Bemisia tabaci isolate Sudan2 cytochrome oxidase subunit I (COI) gene, partial cds; mitochondrial

GAAAATTAGAAGTATTTGGTAGGCTGGGTATAATTTATGCTATGATAACTATTGGCATTTTGGGATTTAT

TGTGTGGGGACATCACATGTTTACTGTTGGAATAGATGTTGACACCCGGGCTTATTTTACTTCGGCTACT

ATAGTTATTGCTGTTCCGACGGGAATTAAAATCTTTAGGTGGCTTGCTACTCTAGGTGGGATAAAATCTA

ATAAGTTTAGACCTCTAGTTCTCTGATTTACAGGATTTCTATTTTTATTTACTATGGGCGGATTAACTGG

AATTATTCTTGGTAATTCTTCGGTAGATGTGTGTCTTCACGACACTTATTTTGTTGTTGCTCATTTTCAT

TATGTTTTATCTATAGGAATTATTTTTGCTATTATAGGCGGAGTAATTTATTGATTCCCTTTAATTTTAG

GCTTAACTCTAAACAATTATAATCTGGTGTCTCAATTTTATATAATATTTGTGGGAGTAAACCTGACATT

TTTTCCACAGCATTTTCTTGGTTTAAGTGGAATACCTCGTCGGTATTCTGACTATCCTGATTGTTATTTA

CTATGAAATAAAATTTCCTCTGGGGGGAGGGTCTTAAGTGTTATTTCTGTTATTTATTTTTTATTTATTA

TTTTAGAATCTTTTCTTCTCTTACGGC

>NewWorld_Barbados_1920_ MK386668_MK386668.1 Bemisia tabaci_collected_1920 cytochrome oxidase subunit I (COI) gene, partial cds; mitochondrial

GAAAATTAGAAGTATTTGGTAGGCTGGGCATAATTTATGCCATGATAACTATTGGTATCTTGGGATTTAT

TGTTTGGGGACATCACATGTTTACTGTTGGAATAGATGTTGACACTCGGGCTTATTTTACTTCAGCTACT

ATAGTTATTGCTGTTCCGACGGGAATTAAAATCTTTAGGTGGCTTGCTACTCTAGGTGGAATAAAGTCTA

ATAAGTTTAGACCCCTAGTTCTCTGATTTACAGGATTTCTATTTTTATTTACTATGGGTGGATTAACTGG

AATTATTCTTGGTAATTCTTCGGTAGATGTGTGTCTTCATGATACTTATTTTGTTGTTGCTCATTTTCAT

TATGTTTTATCTATAGGAATTATCTTTGCTATTATAGGCGGAGTAATTTATTGATTTCCTTTAATTTTAG

GTCTAACTTTAAACAATTATAACCTGGTGTCTCAATTTTATATAATATTTGTGGGAGTAAACCTGACATT

TTTTCCACAGCATTTTCTTGGTTTAAGTGGGATACCTCGTCGGTATTCTGACTACCCTGATTGTTATCTA

CTATGAAATAAAATTTCCTCTGGGGGGAGGGTCTTAAGTGTTATTTCTGTTATTTATTTTTTATTTATTA

TCTTAGAGTCTTTTCTTCTCTTACGGC

>SubSahAf1_Uganda_H46_AY057180_AY057180.1

GTAAACTCGAAGTATTTGGAAGTCTAGGTATAATTTATGCTATGTTAACTATTGGTATTCTGGGGTTTAT

TGTTTGGGGTCATCATATATTTACTGTCGGGATAGATGTGGACACTCGGGCGTATTTCACTTCAGCTACC

ATAATTATTGCTGTTCCTACCGGAATCAAGATTTTTAGGTGACTGGCAACATTGGGTGGTATAAAGTCTA

ATAAGTTTAGTCCGCTAGTTCTTTGATTTACGGGGTTTTTATTTTTATTTACTATAGGAGGTTTAACTGG

AATTATTCTTGGCAATTCTTCTGTCGACGTTTGCTTACATGACACTTATTTTGTTGTTGCTCATTTTCAT

TATGTATTATCTATGGGAATTATTTTTGCTATTATGGGTGGGATTATTTATTGATTTCCCTTAATTCTAG

GGTTAACTTTAAATTATTACAATTTAATTTCTCAATTCTATATTATATTTATTGGAGTAAATCTTACATT

TTTTCCCCAGCATTTTCTTGGTTTGAGGGGAATACCGCGTCGGTATTCAGACTATCCAGATTGCTACCTA

GTTTGAAATAAAATTTCTTCTGTGGGTAGGATCTTGAGTATTATTTCTGTTATTTATTTTTTATTTATTG

TTTTAGAGTCTTTTCTTCTGGTTCGTC

>SubSahAf1_Uganda_Masaka_AY903462_AY903462.1

GTAAACTCGAAGTATTTGGAAGTCTAGGTATAATTTATGCTATGTTAACTATTGGTATTCTGGGGTTTAT

TGTTTGGGGTCATCATATATTTACTGTCGGGATAGATGTGGACACTCGGGCGTATTTCACTTCAGCTACC

ATAATTATTGCTGTTCCTACCGGAATCAAGATTTTTAGGTGACTGGCAACATTGGGTGGTATAAAGTCTA

ATAAGTTTAGTCCGCTAGTTCTTTGATTTACGGGGTTTTTATTTTTATTTACTATAGGAGGTTTAACTGG

AATTATTCTTGGCAATTCTTCTGTCGACGTTTGCTTACATGACACTTATTTTGTTGTTGCTCATTTTCAT

TATGTATTATCTATGGGAATTATTTTTGCTATTATGGGTGGGATTATTTATTGATTTCCCTTAATTCTAG

GGTTAACTTTAAATTATTACAATTTAATTTCTCAATTCTATATTATATTTATTGGAGTAAATCTTACATT

TTTTCCCCAGCATTTTCTTGGTTTGAGGGGAATACCGCGTCGGTATTCAGACTATCCAGATTGCTACCTA

GTTTGAAATAAAATTTCTTCTGTGGGTAGGATCTTGAGTATTATTTCTGTTATTTATTTTTTATTTATTG

TTTTAGAGTCTTTTCTTCTGGTTCGTC

>SubSahAf1_Uganda_Namulonge_AY903511_AY903511.1

GTAAACTCGAAGTATTTGGAAGTCTAGGTATAATTTATGCTATGTTAACTATTGGTATTCTGGGGTTTAT

TGTTTGGGGTCATCATATATTTACTGTCGGGATAGATGTGGACACTCGGGCGTATTTCACTTCAGCTACC

ATAATTATTGCTGTTCCTACCGGAATCAAGATTTTTAGGTGACTGGCAACATTGGGTGGTATAAAGTCTA

ATAAGTTTAGTCCGCTAGTTCTNTGATTTACGGGGTTTTTATTTTTATTTACTATAGGAGGTTTAACTGG

AATTATTCTTGGCAATTCTTCTGTCGACGTTTGCTTACATGACACTTATTTTGTTGTTGCTCATTTTCAT

TATGTATTATCTATGGGAATTATTTTTGCTATTATGGGTGGGATTATTTATTGATTTCCCTTAATTCTAG

GGTTAACTTTAAATTATTACAATTTAATTTCTCAATTCTATATTATATTTATTGGAGTAAATCTTACATT

TTTTCCCCAGCATTTTCTTGGTTTGAGGGGAATACCGCGTCGGTATTCAGACTATCCAGATTGCTACCTA

GTTTGAAATAAAATTTCTTCTGTGGGTAGGATCTTGAGTATTATTTCTGTTATTTATTTTTTATTTATTG

TTTTAGAGTCTTTTCTTCTGGTTCGTC

>SubSahAf1_Uganda_Namulonge_AY903470_AY903470.1

GTAAACTCGAAGTATTTGGAAGTCTAGGTATAATTTATGCTATGTTAACTATTGGTATTCTGGGGTTTAT

TGTTTGGGGTCATCATATATTTACTGTCGGGATAGATGTGGACACTCGGGCGTATTTCACTTCAGCTACC

ATAATTATTGCTGTTCCTACCGGAATCAAGATTTTTAGGTGACTGGCAACATTGGGTGGTATAAAGTCTA

ATAAGTTTAGTCCGCTAGTTCTTTGATTTACGGGGTTTTTATTTTTATTTACTATAGGAGGTTTAACTGG

AATTATTCTTGGCAATTCTTCTGTCGACGTTTGCTTACATGACACTTATTTTGTTGTTGCTCATTTTCAT

TATGTATTATCTATGGGAATTATTTTTGCTATTATGGGTGGGATTATTTATTGATTTCCCTTAATTCTAG

GGTTAACTTTAAATTATTACAATTTAATTTCTCAATTCTATATTATATTTATTGGAGTAAATCTTACATT

TTTTCCCCAGCATTTTCTTGGTCTGAGGGGAATACCGCGTCGGTATTCAGACTATCCAGATTGCTACCTA

GTTTGAAATAAAATTTCTTCTGTGGGTAGGATCTTGAGTATTATTTCTGTTATTTATTTTTTATTTATTG

TTTTAGAGTCTTTTCTTCTGGTTCGTC

>SubSahAf1_Uganda_H42_AY057149_AY057149.1

GTAAACTCGAAGTATTTGGAAGTCTAGGTATAATTTATGCTATGTTAACTATTGGTATTCTGGGGTTTAT

TGTTTGGGGTCATCATATATTTACTGTCGGGATAGATGTGGACACTCGGGCGTATTTCACTTCAGCTACC

ATAATTATTGCTGTTCCTACCGGAATCAAGATTTTTAGGTGACTGGCAACATTGGGTGGTATAAAGTCTA

ATAAGTTTAGTCCGCTAGTTCTTTGATTTACGGGGTTTTTATTTTTATTTACTATAGGAGGTTTAACTGG

AATTATTCTTGGCAATTCTTCTGTCGACGTTTGCTTACATGACACTTATTTTGTTGTTGCTCATTTTCAT

TATGTATTATCTATGGGAATTATTTTTGCTATTATGGGTGGGATTATTTATTGATTTCCCTTAATTCTAG

GGTTAACTTTAAATTATTACAATTTAATTTCTCAATTCTATATTATATTTATTGGAGTAAATCTTACATT

TTTTCCCCAGCATTTTCTTGGTTTGAGGGGAATACCGCGTCGGTATTCAGACTATCCAGATTGCTACCTA

GTTTGAAATAAAATTTCTTCTGTGGGTAGGATCTTGAGTATTATTTCTGTTATTTATTTTTTATTTATTG

TTTTAGAGTCTTTTCTTCTAGTTCGTC

>SubSahAf1_Uganda_Busukuma_AY903487_AY903487.1

GTAAACTCGAAGTATTTGGAAGTCTAGGTATAATTTATGCTATGTTAACTATTGGTATTCTGGGGTTTAT

TGTTTGGGGTCATCATATATTTACTGTCGGGATAGATGTGGACACTCGGGCGTATTTCACTTCAGCTACC

ATAATTATTGCTGTTCCTACCGGAATCAAGATTTTTAGGTGACTGGCAACATTGGGTGGTATAAAGTCCA

ATAAGTTTAGTCCGCTAGTTCTTTGATTTACGGGGTTTTTATTTTTATTTACTATAGGAGGTTTAACTGG

AATTATTCTTGGCAATTCTTCTGTCGACGTTTGCTTACATGACACTTATTTTGTTGTTGCTCATTTTCAT

TATGTATTATCTATGGGAATTATTTTTGCTATTATGGGTGGGATTATTTATTGATTTCCCTTAATTCTAG

GGTTAACTTTAAATTATTACAATTTAATTTCTCAATTCTATATTATATTTATTGGAGTAAATCTTACATT

TTTTCCCCAGCATTTTCTTGGTTTGAGGGGAATACCGCGTCGGTATTCAGACTATCCAGATTGCTACCTA

GTTTGAAATAAAATTTCTTCTGTGGGTAGGATCTTGAGTATTATTTCTGTTATTTATTTTTTATTTATTG

TTTTAGAGTCTTTTCTTCTGGTTCGTC

>SubSahAf1_Uganda_Masaka_AY563648_AY563648.1

GTAAACTCGAAGTATTTGGAAGTCTAGGTATAATTTATGCTATGTTAACTATTGGTATTCTGGGGTTTAT

TGTTTGGGGTCATCATATATTTACTGTCGGGATAGATGTGGACACTCGGGCGTATTTCACTTCAGCTACC

ATAATTATTGCTGTTCCTACCGGAATCAAGATTTTTAGGTGACTGGCAACATTAGGTGGTATAAAGTCTA

ATAAGTTTAGTCCGCTAGTTCTTTGATTTACGGGGTTTTTATTTTTATTTACTATAGGAGGTTTAACTGG

AATTATTCTTGGCAATTCTTCTGTCGACGTTTGCTTACATGACACTTATTTTGTTGTTGCTCATTTTCAT

TATGTATTATCTATGGGAATTATTTTTGCTATTATGGGTGGGATTATTTATTGATTTCCCTTAATTCTAG

GGTTAACTTTAAATTATTACAATTTAATTTCTCAATTCTATATTATATTTATTGGAGTAAATCTTACATT

TTTTCCCCAGCATTTTCTTGGTTTGAGGGGAATACCGCGTCGGTATTCAGACTATCCAGATTGCTACCTA

GTTTGAAATAAAATTTCTTCTGTGGGTAGGATCTTGAGTATTATTTCTGTTATTTATTTTTTATTTATTG

TTTTAGAGTCTTTTCTTCTGGTTCGTC

>SubSahAf1_Uganda_Namulonge_AY903512_AY903512.1

GTAAACTCGAAGTATTTGGAAGTCTAGGTATAATTTATGCTATGTTAACTATTGGTATTCTGGGGTTTAT

TGTTTGGGGTCATCATATATTTACTGTCGGGATAGATGTGGACACTCGGGCGTATTTCACTTCAGCTACC

ATAATTATTGCTGTTCCTACCGGAATCAAGATTTTTAGGTGACTGGCAACATTGGGTGGTATAAAGTCTA

ATAAGTTTAGTCCGCTAGTTCTTTGATTTACGGGGTTTTTATTCTTATTTACTATAGGAGGTTTAACTGG

AATTATTCTTGGCAATTCTTCTGTCGACGTTTGCTTACATGACACTTATTTTGTTGTTGCTCATTTTCAT

TATGTATTATCTATGGGAATTATTTTTGCTATTATGGGTGGGATTATTTATTGATTTCCCTTAATTCTAG

GGTTAACTTTAAATTATTACAATTTAATTTCTCAATTCTATATTATATTTATTGGAGTAAATCTTACATT

TTTTCCCCAGCATTTTCTTGGTTTGAGGGGAATACCGCGTCGGTATTCAGACTATCCAGATTGCTACCTA

GTTTGAAATAAAATTTCTTCTGTGGGTAGGATCTTGAGTATTATTTCTGTTATTTATTTTTTATTTATTG

TTTTAGAGTCTTTTCTTCTGGTTCGTC

>SubSahAf1_Uganda_Soroti_AY563679_AY563679.1

GTAAACTCGAAGTATTTGGAAGTCTAGGTATAATTTATGCTATGTTAACTATTGGTATTCTGGGGTTTAT

TGTTTGGGGTCATCATATATTTACTGTCGGGATAGATGTGGACACTCGGGCGTATTTCACTTCAGCTACC

ATAATTATTGCTGTTCCTACCGGAATCAAGATTTTTAGGTGACTGGCAACATTGGGTGGTATAAAGTCTA

ATAAGTTTAGTCCGCTAGTTCTTTGATTTACGGGGTTTTTATTTTTATTTACTATAGGAGGTTTAACTGG

AATTATTCTTGGCAATTCTTCTGTCGACGTTTGCTTACATGACACTTATTTTGTTGTTGCTCATTTTCAT

TATGTATTATCTATGGGAATTATTTTTGCTATTATGGGTGGGATTATTTATTGATTTCCCTTAATTCTAG

GGTTAACTTTAAATTATTACAATTTAATTTCTCAATTCTATATTATATTTATTGGAGTAAATCTTACATT

TTTTCCCCAGCATTTTCTTGGTTTGAGGGGAATACCGCGTCGGTATTCAGACTATCCAGATTGCTACCTA

GTTTGAAATAAAATTTCTTCTGTGGGTAGGATCTTGAGTATTATTTCGGTTATTTATTTTTTATTTATTG

TTTTAGAGTCTTTTCTTCTGGTTCGTC

>SubSahAf1_Uganda_H44_AY057169_AY057169.1

GTAAACTCGAAGTATTTGGAAGTCTAGGTATAATTTATGCTATGTTAACTATTGGTATTCTGGGGTTTAT

TGTTTGGGGTCATCATATATTTACTGTCGGGATAGATGTGGACACTCGGGCGTATTTCACTTCAGCTACC

ATAATTATTGCTGTTCCTACCGGAATCAAGATTTTTAGGTGACTGGCAACATTGGGTGGTATAAAGTCTA

ATAAGTTTAGTCCGCTAGTTCTTTGATTTACGGGGTTTTTATTTTTATTTACTATAGGAGGCTTAACTGG

AATTATTCTTGGCAATTCCTCTGTCGACGTTTGCTTACATGACACTTATTTTGTTGTTGCTCATTTTCAT

TATGTATTATCTATGGGAATTATTTTTGCTATTATGGGTGGGATTATTTATTGATTTCCCTTAATTCTAG

GGTTAACTTTAAATTATTACAATTTAATTTCTCAATTCTATATTATATTTATTGGAGTAAATCTTACATT

TTTTCCCCAGCATTTTCTTGGTTTGAGGGGTATACCGCGTCGGTATTCAGACTATCCAGATTGCTACCTA

GTTTGAAATAAGATTTCTTCTGTGGGTAGGATCTTGAGTATTATTTCTGTTATTTATTTTTTATTTATCG

TTTTAGAGTCTTTTCTTCTGGTTCGTC

>SubSahAf1_Congo_DQ130054_DQ130054.1

GTAAACTCGAAGTATTTGGAAGTCTAGGTATAATTTATGCTATGTTAACTATTGGTATTCTGGGGTTTAT

TGTTTGGGGTCATCATATATTTACTGTTGGGATAGATGTGGACACTCGGGCGTATTTCACTTCAGCCACC

ATAATTATTGCTGTTCCTACCGGAATCAAAATTTTTAGGTGACTGGCAACATTAGGTGGTATAAAGTCTA

ATAAGTTTAGTCCGCTAGTTCTTTGATTTACGGGGTTTTTATTTTTATTTACTATAGGAGGCTTAACTGG

AATTATTCTTGGCAATTCCTCTGTCGACGTTTGCTTACATGACACTTATTTTGTTGTTGCTCATTTTCAT

TATGTATTATCTATGGGAATTATTTTTGCTATTATGGGTGGGATTATTTATTGATTTCCCTTAATTCTAG

GGTTAACTTTAAATTATTACAATTTAATTTCTCAATTCTATATTATATTTATTGGAGTAAATCTTACATT

TTTTCCCCAGCATTTTCTTGGTTTGAGGGGTATACCGCGTCGGTATTCAGACTATCCAGATTGCTACCTA

GTTTGAAATAAGATTTCTTCTGTGGGTAGGATCTTGAGTATTATTTCTGTTATTTATTTTTTATTTATTG

TTTTAGAGTCTTTTCTTCTGGTTCGTC

>SubSahAf1_Uganda_Mukono_AY903479_AY903479.1

GTAAACTCGAAGTATTTGGAAGTCTAGGTATAATTTATGCTATGTTAACTATTGGTATCCTGGGGTTTAT

TGTTTGGGGTCATCATATATTTACTGTTGGGATAGATGTGGACACTCGGGCGTATTTCACTTCAGCTACC

ATAATTATTGCTGTTCCTACCGGAATCAAAATTTTTAGGTGACTGGCAACATTAGGTGGTATAAAGTCTA

ATAAGTTTAGTCCGCTAGTTCTTTGATTTACGGGGTTTTTATTTTTATTTACTATAGGAGGTTTAACTGG

AATTATTCTTGGCAATTCCTCTGTCGACGTTTGCTTACATGACACTTATTTTGTTGTTGCTCATTTTCAT

TATGTATTATCTATGGGAATTATTTTTGCTATTATGGGTGGGATTATTTATTGATTTCCCTTAATTCTAG

GGTTAACTTTAAATTATTACAATTTAATTTCTCAATTCTATATTATATTTATTGGAGTAAATCTTACATT

TTTTCCCCAGCATTTTCTTGGTTTGAGGGGTATACCGCGTCGGTATTCAGACTATCCAGATTGCTACCTA

GTTTGAAATAAAATTTCTTCTGTGGGTAGGATCTTGAGTATTATTTCTGTTATTTATTTTTTATTTATTG

TTTTAGAGTCCTTTCTTCTGGTTCGTC

>SubSahAf1_Tanzania_AF418667_AF418667.2

GTAAACTCGAAGTATTTGGAAGTCTAGGTATAATTTATGCTATGTTAACTATTGGTATTCTGGGGTTTAT

TGTTTGGGGTCATCATATATTTACTGTTGGGATAGATGTGGATACTCGGGCTTATTTCACTTCAGCTACT

ATAATTATTGCTGTTCCTACCGGAATTAAAATTTTTAGGTGACTGGCAACATTGGGTGGTATAAAGTCTA

ATAAGTTTAGTCCGCTAGTTCTTTGATTTACGGGGTTTTTATTTTTATTTACTATAGGAGGTTTAACTGG

AATTATTCTTGGCAATTCCTCTGTCGACGTTTGCTTACATGACACTTATTTTGTTGTTGCTCATTTTCAT

TATGTATTATCTATGGGAATTATTTTTGCTATTATGGGTGGGATTATTTATTGATTTCCCTTAATTCTAG

GGTTAACTTTAAATTATTACAATTTAATTTCTCAATTCTATATTATATTTATTGGAGTAAATCTTACATT

TTTTCCCCAGCATTTTCTTGGTTTGAGGGGTATACCGCGTCGGTATTCAGACTATCCAGATTGCTACCTA

GTTTGAAATAAAATTTCTTCTGTGGGCAGGATCTTGAGTATTATTTCTGTTATTTATTTTTTATTTATTG

TTTTAGAGTCTTTTCTTCTGGTTCGTC

>SubSahAf1_Uganda_Malawi_AY057162_AY057162.1

GTAAACTCGAAGTATTTGGAAGTCTAGGTATAATTTATGCTATGTTAACTATTGGTATTCTGGGGTTTAT

TGTTTGGGGTCATCATATATTTACTGTTGGGATAGATGTGGACACTCGGGCTTATTTCACTTCAGCTACT

ATAATTATTGCTGTTCCTACCGGAATTAAAATTTTTAGGTGACTGGCAACATTAGGTGGTATAAAGTCTA

ATAAGTTTAGTCCGCTAGTTCTTTGATTTACGGGGTTTTTATTTTTATTTACTATAGGAGGTTTAACTGG

AATTATTCTTGGCAATTCCTCTGTCGACGTTTGCTTACATGACACTTATTTTGTTGTTGCTCATTTTCAT

TATGTATTATCTATGGGAATTATTTTTGCTATTATGGGTGGGATTATTTATTGATTTCCCTTAATTCTAG

GGTTAACTTTAAATTATTACAATTTAATTTCTCAATTCTATATTATATTTATTGGAGTAAATCTTACATT

TTTTCCCCAGCATTTTCTTGGTTTGAGGGGTATACCGCGTCGGTATTCAGACTATCCAGATTGCTACCTA

GTTTGAAATAAAATTTCTTCTGTGGGCAGGATCTTGAGTATTATTTCTGTTATTTATTTTTTATTTATTG

TTTTAGAGTCTTTTCTTCTGGTTCGTC

>SubSahAf1_Ghana_AF418668_AF418668.2

GTAAACTCGAAGTATTTGGAAGTCTAGGTATAATTTATGCTATGTTAACTATTGGTATTCTGGGGTTTAT

TGTTTGGGGTCATCATATATTTACTGTTGGGATAGATGTGGACACTCGGGCTTATTTCACTTCAGCTACT

ATAATTATTGCTGTTCCTACCGGAATTAAAATTTTTAGGTGACTGGCAACATTGGGTGGTATAAAGTCTA

ATAAGTTTAGTCCGCTAGTTCTTTGATTTACGGGGTTTTTATTTTTATTTACTATAGGAGGTTTAACTGG

AATTATTCTTGGTAATTCCTCTGTCGACGTTTGCTTACATGACACTTATTTTGTTGTTGCTCATTTTCAT

TATGTATTATCTATGGGAATTATTTTTGCTATTATGGGTGGGATTATTTATTGATTTCCCTTAATTCTAG

GGTTAACTTTAAATTATTACAATTTAATTTCTCAATTCTATATTATATTTATTGGAGTAAATCTTACATT

TTTTCCCCAGCATTTTCTTGGTCTGAGGGGTATACCGCGTCGGTATTCAGACTATCCAGATTGCTACCTA

GTTTGAAATAAAATTTCTTCTGTGGGTAGGATCTTGAGTATTATTTCTGTTATTTATTTTTTATTTATTG

TTTTAGAGTCTTTTCTTCTGGTTCGTC

>SubSahAf2_Uganda_AY057173_AY057173.1 Bemisia tabaci clone 32Namu cytochrome oxidase subunit I (COI) gene, partial sequence; mitochondrial gene for mitochondrial product

GTAAACTTGAAGTGTTTGGAAGACTGGGAATAATTTATGCTATGCTAACTATCGGTATTCTGGGATTTAT

TGTTTGGGGTCATCATATATTTACTGTTGGGATAGATGTGGATACTCGGGCTTATTTTACTTCAGCTACT

ATAATTATTGCTGTTCCTACTGGAATTAAGATCTTTAGGTGACTCGCAACATTGGGTGGTATAAAGTCTA

ATAAGTTCAGTCCTCTAGTTCTTTGATTTACAGGTTTCTTATTTTTATTTACTATAGGGGGTTTAACTGG

AATTATTCTTGGCAACTCTTCTGTTGACGTTTGTCTACATGACACTTACTTTGTTGTTGCTCATTTTCAT

TATGTATTATCTATGGGAATTATTTTTGCTATTATGGGTGGGATTATTTATTGATTTCCTTTAATCCTAG

GGCTTACTTTAAATTATTATAATCTAATTTCTCAATTTTATATTATATTTATTGGAGTAAATCTTACATT

TTTTCCACAACATTTCCTTGGTTTAAGGGGGATGCCTCGCCGGTATTCAGATTATCCAGACTGCTATCTA

GTTTGAAACAAAATTTCTTCTGTGGGGAGTATCTTGAGTATTATTTCTGTTATTTATTTTTTATTTATTG

TTTTAGAGTCTTTTCTTCTGATCCGGC

>SubSahAf2_Uganda_Kumi_AY563646_AY563646.1 Bemisia tabaci clone 52KumK cytochrome oxidase subunit I (COI) gene, partial cds; mitochondrial

GTAAACTTGAAGTGTTTGGAAGACTGGGAATAATTTATGCTATGCTAACTATCGGTATTCTGGGATTTAT

TGTTTGGGGTCATCATATATTTACTGTTGGGATAGATGTGGATACTCGGGCTTATTTTACTTCAGCTACT

ATAATTATTGCTGTTCCTACTGGAATTAAGATCTTTAGGTGACTCGCAACATTGGGTGGTATAAAGTCTA

ATAAGTTCAGTCCTCTAGTTCTTTGATTTACAGGTTTCTTATTTTTATTTACTATAGGGGGTTTAACTGG

AATTATTCTTGGCAACTCTTCTGTTGACGTTTGTCTACATGACACTTATTTTGTTGTTGCTCATTTTCAT

TATGTATTATCTATGGGAATTATTTTTGCTATTATGGGTGGGATTATTTATTGATTTCCTTTAATCCTAG

GGCTTACTTTAAATTATTATAATCTAATTTCTCAATTTTATATTATATTTATTGGAGTAAATCTTACATT

TTTTCCACAACATTTCCTTGGTTTAAGGGGGATGCCTCGCCGGTATTCAGATTATCCAGACTGCTATCTA

GTTTGAAACAAAATTTCTTCTGTGGGGAGTATCTTGAGTATTATTTCTGTTATTTATTTTTTATTTATTG

TTTTAGAGTCTTTTCTTCTGATCCGGC

>SubSahAf2_Uganda_AY057141_AY057141.1 Bemisia tabaci clone 1Buli cytochrome oxidase subunit I (COI) gene, partial cds; mitochondrial gene for mitochondrial product

GTAAACTTGAAGTGTTTGGAAGACTGGGAATAATTTATGCTATGCTAACTATCGGTATTCTGGGATTTAT

TGTTTGGGGTCATCATATATTTACTGTTGGGATAGATGTGGATACTCGGGCTTATTTTACTTCAGCTACT

ATAATTATTGCTGTTCCTACTGGAATTAAGATCTTTAGATGACTCGCAACATTGGGTGGTATAAAGTCTA

ATAAGTTCAGTCCTCTAGTTCTTTGATTTACAGGTTTCTTATTTTTATTTACTATAGGGGGTTTAACTGG

AATTATTCTTGGCAACTCTTCTGTTGACGTTTGTCTACATGACACTTATTTTGTTGTTGCTCATTTTCAT

TATGTATTATCTATGGGAATTATTTTTGCTATTATGGGTGGGATTATTTATTGATTTCCTTTAATCCTAG

GGCTTACTTTAAATTATTATAATCTAATTTCTCAATTTTATATTATATTTATTGGAGTAAATCTTACATT

TTTTCCACAACATTTCCTTGGTTTAAGGGGGATGCCTCGCCGGTATTCAGATTATCCAGACTGCTATCTA

GTTTGAAACAAAATTTCTTCTGTGGGGAGTATCTTGAGTATTATTTCTGTTATTTATTTTTTATTTATTG

TTTTAGAGTCTTTCCTTCTGATCCGGC

>SubSahAf2_Mali_AY827605_AY827605.1 Bemisia tabaci voucher IMIDA-Ma1.2 cytochrome oxidase subunit I (COI) gene, partial cds; mitochondrial

GTAAACTTGAAGTGTTTGGAAGATTGGGAATAATTTATGCTATGCTAACTATCGGTATTCTGGGATTTAT

TGTTTGGGGTCATCATATATTTACTGTTGGGATAGATGTGGATACTCGGGCGTATTTTACTTCAGCTACT

ATAATTATTGCTGTTCCTACTGGAATTAAGATCTTTAGGTGACTCGCAACATTGGGTGGTATAAAGTCTA

ATAAGTTCAGTCCTCTAGTTCTTTGATTTACAGGTTTCTTATTTTTATTTACTATAGGGGGTTTAACTGG

AATTATTCTTGGCAACTCTTCTGTTGACGTTTGTCTACATGACACTTATTTTGTTGTTGCTCATTTTCAT

TATGTATTATCTATGGGAATTATTTTTGCTATTATGGGTGGGATTATTTATTGATTTCCTTTAATCCTAG

GGCTTACTTTAAATTATTATAATCTAATTTCTCAATTTTATATTATATTTATTGGAGTAAATCTTACATT

TTTTCCACAACATTTCCTTGGTTTAAGGGGGATGCCTCGCCGGTATTCAGATTATCCAGACTGCTATTTA

GTTTGAAACAAAATTTCTTCTGTGGGGAGTATCTTGAGTATTATTTCTGTTATTTATTTTTTATTTATTG

TTTTAGAGTCTTTTCTTCTGATCCGGC

>SubSahAf2_Mali_AY827604_AY827604.1 Bemisia tabaci voucher IMIDA-Ma1.1 cytochrome oxidase subunit I (COI) gene, partial cds; mitochondrial

GTAAACTTGAAGTGTTTGGAAGACTGGGAATAATTTATGCTATGCTAACTATCGGTATTCTGGGATTTAT

TGTTTGGGGTCATCATATATTTACTGTTGGGATAGATGTGGATACTCGGGCGTATTTTACTTCAGCTACT

ATAATTATTGCTGTTCCTACTGGAATTAAGATCTTTAGGTGACTCGCAACATTGGGTGGTATAAAGTCTA

ATAAGTTCAGTCCTCTAGTTCTTTGATTTACAGGTTTCTTATTTTTATTTACTATAGGGGGTTTAACTGG

AATTATTCTTGGCAACTCTTCTGTTGACGTTTGTCTACATGACACTTATTTTGTTGTTGCTCATTTTCAT

TATGTATTATCTATAGGAATTATTTTTGCTATTATGGGTGGGATTATTTATTGATTTCCTTTAATTCTAG

GGCTTACTTTAAATTATTATAATCTAATTTCTCAATTTTATATTATATTTATTGGAGTAAATCTTACATT

TTTTCCACAACATTTCCTTGGTTTAAGGGGGATGCCTCGCCGGTATTCAGATTATCCAGACTGCTATCTA

GTTTGAAACAAAATTTCTTCTGTGGGGAGTATCTTGAGTATTATTTCTGTTATTTATTTTTTATTTATTG

TTTTAGAGTCTTTTCTTCTGATCCGGC

>KM377923.1 Bemisia tabaci isolate NG_Owerri-5_Imo_state_SSA3 cytochrome oxidase subunit I gene, partial cds; mitochondrial

GTAAACTTGAAGTATTTGGAAGATTAGGAATAATTTATGCTATGTTGACTATTGGCATTCTGGGATTTAT

CGTTTGGGGTCATCATATATTTACTGTTGGGATAGATGTGGATACTCGGGCGTATTTTACTTCAGCTACT

ATAATTATTGCTGTTCCTACTGGAATTAAGATTTTTAGATGACTCGCAACATTGGGTGGTATAAAGTCTA

ATAAGTTTAGTCCCCTAGTTCTTTGATTTACGGGTTTTTTATTTTTATTTACTATAGGGGGTTTAACTGG

AATTATTCTTGGTAATTCTTCTGTCGACGTTTGTCTACATGATACTTATTTTGTTGTTGCTCATTTTCAT

TATGTATTATCTATGGGAATTATTTTTGCTATCATGGGTGGGATTATTTATTGATTTCCTTTAATTTTAG

GACTAACCTTAAATTATTACAACTTAATTTCTCAATTTTATATTATATTTATTGGAGTAAATCTCACATT

TTTTCCACAACACTTTCTTGGTTTAAGGGGGATACCACGTCGGTATTCAGACTATCCAGACTGTTATCTA

GTTTGAAACAAAATTTCTTCTGTGGGGAGGATCTTGAGTATTATTTCTGTTATTTATTTTTTATTTATTG

TTTTAGAGTCTTTCCTTCTAATTCGGC

>KM377925.1 Bemisia tabaci isolate NG_Umuahia-5_Abia_SSA3 cytochrome oxidase subunit I gene, partial cds; mitochondrial

GTAAACTTGAAGTATTTGGAAGATTAGGAATAATTTATGCTATGTTGACTATTGGCATTCTGGGATTTAT

CGTTTGGGGTCATCATATATTTACTGTTGGGATAGATGTGGATACTCGGGCGTATTTTACTTCAGCTACT

ATAATTATTGCTGTTCCTACTGGAATTAAGATTTTTAGATGACTCGCAACATTGGGTGGTATAAAGTCTA

ATAAGTTTAGTCCCCTAGTTCTTTGATTTACGGGGTTTTTATTTTTATTTACTATAGGGGGTTTAACTGG

AATTATTCTTGGTAATTCTTCTGTCGACGTTTGTCTACATGATACTTATTTTGTTGTTGCTCATTTTCAT

TATGTATTATCTATGGGAATTATTTTTGCTATCATGGGTGGGATTATTTATTGATTTCCTTTAATTTTAG

GACTAACCTTAAATTATTACAACTTAATTTCTCAATTTTATATTATATTTATTGGAGTAAATCTCACATT

TTTTCCACAACACTTTCTTGGTTTAAGGGGGATACCACGTCGGTATTCAGACTATCCAGACTGTTATCTA

GTTTGAAACAAAATTTCTTCTGTGGGGAGGATCTTGAGTATTATTTCTGTTATTTATTTTTTATTTATTG

TTTTAGAGTCTTTCCTTCTAATTCGGC

>KM377934.1 Bemisia tabaci isolate NG_Owerri3_Imo_state_SSA3 cytochrome oxidase subunit I gene, partial cds; mitochondrial

GTAAACTTGAAGTATTTGGAAGATTAGGAATAATTTATGCTATGTTGACTATTGGCATTCTGGGATTTAT

CGTTTGGGGTCATCATATATTTACTGTTGGGATAGATGTGGATACTCGGGCGTATTTTACTTCAGCTACT

ATAATTATTGCTGTTCCTACTGGAATTAAGATTTTTAGATGACTCGCAACATTGGGTGGTATAAAGTCTA

ATAAGTTTAGTCCCCTAGTTCTTTGATTTACGGGGTTTTTATTTTTATTTACTATAGGGGGTTTAACTGG

AATTATTCTTGGTAATTCTTCTGTCGACGTTTGTCTACATGATACTTATTTTGTTGTTGCTCATTTTCAT

TATGTATTATCTATGGGAATTATTTTTGCTATCATGGGTGGGATTATTTATTGATTTCCTTTAATTTTAG

GACTAACCTTAAATTATTACAACTTAATTTCTCAATTTTATATTATATTTATTGGAGTAAATCTCACATT

TTTTCCACAACACTTTCTTGGTTTAAGGGGGATACCACGCCGGTATTCAGACTATCCAGACTGTTATCTA

GTTTGAAACAAAATTTCTTCTGTGGGGAGGATCTTGAGTATTATTTCTGTTATTTATTTTTTATTTATTG

TTTTAGAGTCTTTCCTTCTAATTCGGC

>KF425604.1 Bemisia tabaci isolate DC129 cytochrome oxidase subunit I (COI) gene, partial cds; mitochondrial

GTAAACTTGAAGTGTTTGGAAGATTAGGAATAATTTATGCTATGTTGACTATTGGCATTCTGGGATTTAT

CGTTTGGGGTCATCATATATTTACTGTTGGGATAGATGTGGATACTCGGGCGTATTTTACTTCAGCTACT

ATAATTATTGCTGTTCCTACTGGAATTAAGATTTTTAGATGACTCGCAACATTGGGTGGTATAAAGTCTA

ATAAGTTTAGTCCCCTAGTTCTTTGATTTACGGGTTTTTTATTTTTATTTACTATAGGGGGTTTAACTGG

AATTATTCTTGGTAATTCTTCTGTCGACGTTTGTCTACATGATACTTATTTTGTTGTTGCTCATTTTCAT

TATGTATTATCTATGGGAATTATTTTTGCTATCATGGGTGGGATTATTTATTGATTTCCTTTAATTTTAG

GATTAACCTTAAATTATTACAACTTAATTTCTCAATTTTATATTATATTTATTGGAGTAAATCTCACATT

TTTTCCACAACACTTTCTTGGTTTAAGGGGGATACCACGTCGGTATTCAGACTATCCAGACTGTTATCTA

GTTTGAAACAAAATTTCTTCTGTGGGGAGGATCTTGAGTATTATTTCTGTTATTTATTTTTTATTTATTG

TTTTAGAGTCTTTCCTTCTAATTCGGC

>KF425602.1 Bemisia tabaci isolate DC122 cytochrome oxidase subunit I (COI) gene, partial cds; mitochondrial

GTAAACTTGAAGTGTTTGGAAGATTAGGAATAATTTATGCTATGTTGACTATTGGCATTCTGGGATTTAT

CGTTTGGGGTCATCATATATTTACTGTTGGGATAGATGTGGATACTCGGGCGTATTTTACTTCAGCTACT

ATAATTATTGCTGTTCCTACTGGAATTAAGATTTTTAGATGACTCGCAACATTGGGTGGTATAAAGTCTA

ATAAGTTTAGTCCCCTAGTTCTTTGATTTACGGGTTTTTTATTTTTATTTACTATAGGGGGTTTAACTGG

AATTATTCTTGGTAATTCTTCTGTCGACGTTTGTCTACATGATACTTATTTTGTTGTTGCTCATTTTCAT

TATGTATTATCTATGGGAATTATTTTTGCTATCATGGGTGGGATTATTTATTGATTTCCTTTAATTTTAG

GATTAACCTTAAATTATTACAACTTAATTTCTCAATTTTATATTATATTTATTGGAGTAAATCTCACATT

TTTTCCACAACACTTTCTTGGTTTAAGGGGGATACCACGTCGGTATTCAGATTATCCAGACTGTTATCTA

GTTTGAAACAAAATTTCTTCTGTGGGGAGGATCTTGAGTATTATTTCTGTTATTTATTTTTTATTTATTG

TTTTAGAGTCTTTCCTTCTAATTCGGC

>HQ908651.1 Bemisia tabaci isolate To2-1 cytochrome oxidase subunit I (COI) gene, partial cds; mitochondrial

NNAAACTTGAAGTATTTGGAAGATTAGGAATAATTTATGCTATGTTGACTATTGGCATTCTGGGATTTAT

CGTTTGGGGTCATCATATATTTACTGTTGGAATAGATGTGGATACTCGGGCGTATTTTACTTCAGCTACT

ATAATTATTGCTGTTCCTACTGGAATTAAGATTTTTAGATGACTCGCAACATTGGGTGGTATAAAGTCTA

ATAAGTTTAGTCCCCTAGTTCTTTGATTTACGGGTTTTTTATTTTTATTTACTATAGGGGGTTTAACTGG

AATTATTCTTGGTAATTCTTCTGTCGACGTTTGTCTACATGATACTTATTTTGTTGTTGCTCATTTTCAT

TATGTATTATCTATGGGAATTATTTTTGCTATCATGGGTGGGATTATTTATTGATTTCCTTTAATTTTAG

GACTAACCTTAAATTATTACAACTTAATTTCTCAATTTTATATTATATTTATTGGAGTAAATCTCACATT

TTTTCCACAACACTTTCTTGGTTTAAGGGGGATACCACGTCGGTATTCAGACTATCCAGACTGTTATCTA

GTTTGAAACAAAATTTCTTCTGTGGGGAGGATCTTGAGTATTATTTCTGTTATTTATTTTTTATTTATTG

TTTTAGAGTCTTTCCTTCTAATTCGGC

>MG565972.1 Bemisia tabaci isolate SSA3_JN3 cytochrome oxidase subunit I gene, partial cds; mitochondrial

NNAAACTTGAAGTATTTGGAAGATTAGGAATAATTTATGCTATGTTGACTATTGGCATTCTGGGATTTAT

CGTTTGGGGTCATCATATATTTACTGTTGGGATAGATGTGGATACTCGGGCGTATTTTACTTCAGCTACT

ATAATTATTGCTGTTCCTACTGGAATTAAGATTTTTAGATGACTTGCAACATTGGGTGGTATAAAGTCTA

ATAAGTTTAGTCCCCTAGTTCTTTGATTTACGGGTTTTTTATTTTTATTTACTATAGGGGGTTTAACTGG

AATTATTCTTGGTAATTCTTCTGTCGACGTTTGTCTACATGATACTTATTTTGTTGTTGCTCATTTTCAT

TATGTATTATCTATGGGAATTATTTTTGCTATCATGGGTGGGATTATTTATTGATTTCCTTTAATTTTAG

GACTAACCTTAAATTATTACAACTTAATTTCTCAATTTTATATTATATTTATTGGAGTAAATCTCACATT

TTTTCCACAACACTTTCTTGGTTTAAGGGGGATACCACGTCGGTATTCAGACTATCCAGACTGTTATCTA

GTTTGAAACAAAATTTCTTCTGTGGGGAGGATCTTGAGTATTATTTCTGTTATTTATTTTTTATTTATTG

TTTTAGAGTCTTTCCTTCTAATTCGGN

>Uganda_Namulonge_AY903576_AY903576.1 Bemisia tabaci clone UgSpNm15 cytochrome oxidase subunit I-like gene, complete sequence; mitochondrial

GAAAATTAGAAGTATTTGGTAGACTAGGTATAATCTATGCTATATTGACTATCGGTATCCTTGGCTTTAT

TGTATGAGGTCACCATATGTTCACTGTTGGCATAGATGTAGATACTCGAGCTTATTTCACTTCGGCTACT

ATAATTATCGCTGTTCCTACAGGAATTAAAATTTTTAGATGATTAGCTACTCTTGGTGGAATAAAATCTA

ACAAGTTTAGGCCGTTGGTTCTTTGGTTTACCGGATTTTTATTCCTGTTTACTATGGGAGGTTTAACCGG

AATTATTCTTGGTAATTCTTCTGTTGATATTTGTTTACATGATACATATTTTGTTGTTGCTCATTTTCAT

TATGTACTATCTATAGGAATTATTTTTGCTATTATGGGTGGAATAATTTATTGATTTCCTCTAATCTTGG

GGCTTACTTTAAACAACTATAATTTAGTTTCTCAGTTTTATATTATATTCATTGGAGTAAATTTAACATT

TTTTCCCCAACATTTTCTTGGTTTGAGAGGGATACCTCGGCGATATTCGGATTACCCCGACTGTTATCTA

TTATGAAATAAACTTTCTTCTGTGGGAAGAATTTTAAGTGTTGTTTCGATTGTTTACTTTATATGTATTG

TGCTAGAGTCATTTATCCTTTTACGAT

>KX714968_Bemisia_sp_’JpL’_cytochrome oxidase subunit 1 gene,partial cds; mitochondrial.

GAAAACTTGAAGTTTTTGGTAGACTAGGAATAATTTATGCTATGTTAACTATTGGTATTTTAGGTTTTAT

TGTTTGAGGTCATCACATATTTACTGTTGGGATAGATGTTGATACTCGGGCGTATTTTACTTCAGCTACA

ATAATTATTGCTGTTCCTACGGGGATTAAAATCTTTAGTTGACTTGCTACTTTAGGAGGTATAAAGTCTA

ATAAGTTTAGTCCTCTTGTTCTTTGATTTACAGGGTTTTTATTTTTATTTACTATGGGTGGTTTAACTGG

AATCATTTTGGGTAATTCTTCTGTTGATGTTTGTTTACATGATACTTATTTTGTTGTTGCTCATTTTCAC

TATGTTTTATCTATGGGAATTATTTTTGCTGTTATGGGTGGGATTATTTATTGGTTTCCCCTGGTTTTAG

GTCTGAGTTTAAATAATTATAGTTTAGTGTCTCAATTTTATATGATATTCATGGGAGTAAATTTAACATT

TTTTCCTCAACACTTTCTTGGCTTAGGGGGAATACCTCGTCGATATTCAGATTATCCTGATTGTTATCTT

TTGTGGAACAAAATTTCCTCTGCGGGAAGCATCTTAAGTATTATTTCTGTCATTTATTTTTTATTTATTA

TTCTTGAATCTTTATTACTTCTTCGAT

>HM137313_Asia_II-9 isolate h1 cytochrome oxidase subunit 1 gene,partial cds; mitochondrial.

GAAAGCTTGAAGTATTTGGAAGGCTGGGTATAATTTATGCTATAGTAACT

ATTGGTATTTTAGGTTTTATTGTCTGAGGACATCATATATTTACTGTTGG

GATAGATGTTGATACCCGAGCTTATTTTACCTCAGCCACTATAATTATTG

CTGTTCCGACTGGAATTAAAATCTTTAGATGACTTGCTACTTTAGGTGGA

ATAAAGTCTAACAAGCTCAGTCCGCTTGTTCTTTGGTTTACTGGATTTTT

ATTTTTATTTACTATGGGTGGGTTAACTGGAATTATTCTTGGTAACTCTT

CTGTCGATGTTTGCTTACACGATACTTATTTTGTTGTCGCTCATTTTCAT

TATGTTTTATCTATAGGGATTATTTTTGCTATTGTGGGTGGGGTTATTTA

TTGATTTCCATTAATTTTAGGCTTAACACTGAATAGCCATAGTCTGGTAT

CTCAATTTTACATTATATTCCTAGGAGTAAATTTAACATTTTTTCCACAA

CATTTTCTTGGGCTAAGGGGGATACCTCGCCGGTATTCAGATTATCCTGA

TTGTTATTTGATATGAAATAAAATTTCTTCTGCAGGGAGAATTTTGAGTA

TCATCTCTGTTATTTATTTTTTATTTATTGTTTTAGAGTCTTTACTTCTT

TTGCGAT

>HM137345_Asia_II-9 isolate h33 cytochrome oxidase subunit 1 gene,partial cds; mitochondrial.

GAAAGCTTGAAGTATTTGGAAGGCTGGGTATAATTTATGCTATAGTAACT

ATTGGTATTTTAGGTTTTATTGTCTGAGGACATCATATATTTACTGTTGG

GATAGATGTTGATACCCGAGCTTATTTTACCTCAGCCACTATAATTATTG

CTGTTCCGACTGGAATTAAAATCTTTAGATGACTTGCTACTTTAGGTGGA

ATAAAGTCTAACAAGCTCAGTCCGCTTGTTCTTTGGTTTACTGGATTTTT

ATTTTTATTTACTATGGGCGGGTTAACTGGAATTATTCTTGGTAACTCTT

CTGTCGATGTTTGCTTACACGATACTTATTTTGTTGTTGCTCATTTTCAT

TATGTTTTATCTATAGGGATTATTTTTGCTATTGTGGGTGGGGTTATTTA

TTGATTTCCATTAATTTTAGGCTTAACACTGAATAGCCATAGTCTGGTAT

CTCAATTTTACATTATATTCCTAGGAGTAAATTTAACATTTTTTCCACAA

CATTTTCTTGGGCTAAGGGGGATACCTCGCCGGTATTCAGATTATCCTGA

TTGTTATTTGATATGAAATAAAATTTCTTCTGCAGGGAGAATTTTGAGTA

TCATCTCTGTTATTTATTTTTTATTTATTGTTTTAGAGTCTTTACTTCTT

TTGCGAT

>HM137356_Asia_II-10 isolate g44 cytochrome oxidase subunit 1 gene,partial cds; mitochondrial.

GAAAACTTGAAGTATTTGGAAGGCTAGGTATAATTTATGCTATAGTGACT

ATTGGAATTTTAGGTTTCATTGTCTGGGGTCATCATATATTTACTGTTGG

AATAGATGTTGATACTCGAGCTTATTTTACCTCAGCTACTATGATTATTG

CTGTTCCGACTGGAATTAAAATCTTTAGGTGACTTGCTACTTTAGGTGGA

ATAAAATCTAACAAATTTAGGCCGCTTGTTCTTTGATTCACTGGGTTTTT

GTTTTTATTTACTATGGGGGGGTTAACTGGAATTATTCTTGGTAATTCTT

CTGTTGATGTTTGTTTACACGATACTTATTTTGTTGTTGCTCATTTTCAT

TATGTCTTATCTATAGGGATCATTTTTGCTATTGTGGGTGGAGTTATTTA

CTGATTTCCACTAATTTTAGGGCTAACTCTAAATAGTCACAGCTTGGTAT

CCCAGTTTTATATCATATTCTTGGGAGTAAACTTAACGTTTTTTCCACAA

CATTTTCTTGGACTAAGAGGAATACCTCGCCGGTATTCAGACTATCCTGA

TTGTTATTTGATATGAAATAAAATCTCTTCTGCAGGGAGAATCTTGAGTA

TTATTTCTATTATTTATTTGTTATTTATTGTTTTAGAGTCTTTACTTCTT

TTACGAT

>HM137339_Asia_II-10 isolate g27 cytochrome oxidase subunit 1 gene,partial cds; mitochondrial.

GAAAACTTGAAGTATTTGGAAGGCTAGGTATAATTTATGCTATAGTGACT

ATTGGAATTTTAGGTTTCATTGTCTGGGGTCATCATATATTTACTGTTGG

AATAGATGTTGATACTCGAGCTTATTTTACCTCAGCTACTATGATTATTG

CTGTTCCGACTGGAATTAAAATCTTTAGGTGACTTGCTACTTTAGGTGGA

ATAAAATCTAACAAATTTAGGCCGCTTGTTCTTTGATTCACTGGGTTTTT

GTTTTTATTTACTATGGGGGGGTTAACTGGAATTATTCTTGGTAATTCTT

CTGTTGATGTTTGTTTACACGATACTTATTTTGTTGTTGCTCATTTTCAT

TATGTCTTATCTATAGGGATCATTTTTGCTATTGTGGGTGGAGTTATTTA

CTGATTTCCACTAATTTTAGGGCTAACTCTAAATAGTCACAGCTTGGTAT

CCCAGTTTTATATTATATTTTTGGGAGTAAACTTAACGTTTTTTCCACAA

CATTTTCTTGGACTAAGAGGAATACCTCGCCGGTATTCAGACTATCCTGA

TTGTTATTTGATATGAAATAAAATCTCTTCTGCAGGGAGAATCTTGAGTA

TTATTTCTATTATTTATTTGTTATTTATTGTTTTAGAGTCTTTACTTCTT

TTACGAT

>DQ174527_Asia_III from Mesona chinensis cytochrome oxidase subunit Igene, partial cds; mitochondrial.

GGAAACTTGAGGTATTCGGCAGGTTAGGAATAATTTACGCTATAATAACT

ATTGGCATTCTAGGTTTTATTGTTTGAGGTCATCATATATTTACTGTTGG

GATGGATGTTGATACTCGAGCTTATTTTACCTCAGCTACTATGGTTATTG

CTGTTCCAACTGGGATTAAAATTTTCAGGTGGCTTGCCACTTTAGGTGGA

ATAAAATCTAATAAATTTAGGCCGTTAGTGCTTTGATTTACAGGATTTTT

ATTTTTATTTACTATGGGTGGATTAACGGGGATTATTCTTGGTAATTCTT

CTGTTGATGTTTGCTTACACGATACCTATTTTGTTGTTGCTCACTTTCAT

TATGTTTTATCCATGGGAATTATTTTTGCTATTATGGGAGGTCTTATTTA

TTGATTTCCACTAATTTTAGGTCTAACATTAAATAACCACAATTTGGTAT

CTCAGTTTTATATTATATTTTTGGGAGTTAATCTAACATTTTTCCCACAA

CATTTTCTTGGCTTAAGTGGAATGCCTCGTCGGTATTCAGATTATCCTGA

TTGTTATCTTATGTGGAATAAAATTTCTTCTGCGGGGAGTATCTTGAGTA

TTGTTTCTGTTATCTACTTTTTATTTATTGTTTTAGAATCTCTGCTTCTT

TTACGAT

>DQ174528_Asia_III from Solanaceae cytochrome oxidase subunit I gene,partial cds; mitochondrial.

GGAAACTTGAGGTATTCGGCAGGTTAGGAATAATTTACGCTATAATAACT

ATTGGCATTCTAGGTTTTATTGTTTGAGGTCATCATATATTTACTGTTGG

GATGGATGTTGATACTCGAGCTTATTTTACCTCAGCTACTATGGTTATTG

CTGTTCCAACTGGGATTAAAATTTTCAGGTGGCTTGCCACTTTAGGTGGA

ATAAAATCTAATAAATTTAGGCCGTTAGTGCTTTGATTTACAGGATTTTT

ATTTTTATTTACTATGGGTGGATTAACGGGGATTATTCTTGGCAATTCTT

CTGTTGATGTTTGCTTACACGATACCTATTTTGTTGTTGCTCACTTTCAT

TATGTTTTATCCATGGGAATTATTTTTGCTATTATGGGAGGTCTTATTTA

TTGATTTCCACTAATTTTAGGTCTAACATTAAATAACCACAATTTGGTAT

CTCAGTTTTATATTATATTTTTGGGAGTTAATCTAACATTTTTCCCACAA

CATTTTCTTGGCTTAAGTGGAATGCCTCGTCGGTATTCAGATTATCCTGA

TTGTTATCTTATGTGGAATAAAATTTCTTCTGCGGGGAGTATCTTGAGTA

TTGTTTCTGTTATCTACTTTTTATTTATTGTTTTAGAATCTCTGCTTCTT

TTACGAT

>AB440792_Asia_III mitochondrial COI gene for cytochrome oxidasesubunit I, partial cds, isolate: Taketomi-2008-Wb.

GGAAACTTGAGGTATTCGGCAGGTTAGGAATAATTTACGCTATAATAACT

ATTGGCATTCTAGGTTTTATTGTTTGAGGTCATCATATATTTACTGTTGG

GATGGATGTTGATACTCGAGCTTATTTTACCTCAGCTACTATGGTTATTG

CTGTTCCAACTGGGATTAAAATTTTCAGGTGGCTTGCCACTTTAGGTGGA

ATAAAATCTAATAAATTTAGGCCGTTAGTGCTTTGATTTACAGGATTTTT

ATTTTTATTTACTATGGGTGGATTAACGGGGATTATTCTTGGCAATTCTT

CTGTTGATGTTTGCTTACACGATACCTATTTTGTTGTTGCTCACTTTCAT

TATGTTTTATCCATGGGAATTATTTTTGCTATTATGGGAGGTCTTATTTA

TTGATTTCCACTAATTTTAGGTCTAACATTAAATAACTACAATTTGGTAT

CTCAGTTTTATATTATATTTTTGGGAGTTAATCTAACATTTTTTCCACAA

CATTTTCTTGGCTTAAGTGGAATGCCTCGTCGGTATTCAGATTATCCTGA

TTGTTATCTTATGTGGAATAAAATTTCTTCTGCGGGGAGTATCTTGAGTA

TTGTTTCTGTTATCTACTTTTTATTTATTGTTTTAGAATCTCTGCTTCTT

TTACGAT

>AsiaII_11_India_Andhra_Pradesh_HM590147_HM590147.1 Bemisia tabaci isolate KONDGOL-DDK11 cytochrome oxidase subunit 1 gene, partial cds; mitochondrial

GAAAACTTGAAGTATTTGGAAGATTAGGGATAATTTATGCCATAGTAACTATTGGTATCT

TAGGTTTTATTGTTTGAGGTCATCACATATTTACTGTTGGGATAGATGTTGATACTCGGG

CTTATTTTACTTCAGCTACCATAATCATTGCTGTTCCAACTGGAATTAAAATTTTTAGGT

GACTTGCTACTTTAGGAGGAATAAAATCTAACAAATTTAGGCCTCTTGTTCTTTGATTTA

CTGGATTTTTGTTTTTATTTACTATGGGTGGATTAACTGGAATTATCCTTGGTAATTCCT

CTGTTGATGTCTGCTTACATGATACTTATTTTGTTGTTGCTCATTTTCATTATGTTTTGT

CTATAGGGATTATTTTTGCTATTGTAGGAGGAGTTATTTATTGATTTCCATTAATTTTAG

GTTTGACACTAAATGGTCATAGGCTGGTGTCTCAGTTTTACATTATGTTTCTGGGAGTAA

ATTTGACGTTTTTTCCACAACACTTTCTTGGACTAAGGGGGATGCCTCGCCGGTATTCAG

ATTATCCTGATTGTTACTTAATATGAAATAAGATTTCTTCTGCGGGTAGAATTTTGAGTA

TTATTTCTGTTATTTATTTTTTATTTATTGTTCTAGAGTCTTTTCTACTTTTGCGAT

>EU760739_Africa_Cameroon_EU760739_EU760739.1 Bemisia tabaci isolate Cameroon Ma5 cytochrome oxidase subunit I (COI) gene, partial cds; mitochondrial

GGAAGCTTGAAGTATTTGGGAGTTTAGGAATAATTTACGCTATAATAACTATTGGTATTC

TAGGTTTTATTGTTTGAGGCCATCATATGTTTACTGTTGGAATAGATGTGGACACTCGGG

CTTATTTTACTTCAGCTACAATGATTATCGCTGTTCCAACAGGAATTAAGATTTTTAGAT

GACTCGCTACTTTAGGTGGGATGAAGTCTAATAAGTTTAGTCCGCTTGTTCTTTGATTTA

CAGGATTTTTATTTTTGTTTACCATGGGTGGGCTAACTGGAATTATTCTTGGTAATTCTT

CTGTTGATGTTTGTTTACACGATACTTACTTTGTTGTTGCCCACTTTCATTATGTTTTAT

CTATAGGGATTATTTTTGCCATTGTTGGAGGGTTTATTTACTGATTTCCATTAATTTTAG

GTTTAACATTAAATAGACATAATCTAGTGTCTCAGTTTTATCTTATGTTTATTGGAGTAA

ACTTAACATTCTTTCCCCAGCATTTTCTGGGACTAAGGGGAATACCTCGCCGGTACTCAG

ATTATCCTGATTGTTATTTAATATGAAACAAAATTTCTTCTGCTGGGAGAATTTTAAGTG

TTGTTTCTGTCATTTACTTCTTATTTATTGTTTTAGAATCTCTGCTGCTTTTACGCT

>KX570778_MEAM2_African MEAM2 Mugerwa2018

GAAAATTAGAGGTATTTGGAAGGTTAGGTATAATTTATGCTATATTGACTATTGGTATCTTAGGATTTAT

TGTTTGAGGCCATCATATATTTACAGTTGGAATAGATGTAGATACTCGAGCTTATTTCACTTCAGCTACC

ATGATTATTGCTGTTCCTACAGGAATTAAAATTTTTAGCTGGCTTGCTACTTTGGGTGGAATAAAGTCTA

ATAAATTCAGACCTCTTGGCCTTTGATTTACAGGATTTCTATTTTTATTTACTATAGGCGGATTAACTGG

GATTATTCTTGGTAATTCTTCTGTAGATGTGTGTCTGCATGATACTTATTTTGTTGTTGCACATTTTCAT

TATGTTTTATCAATAGGAATTATTTTTGCTATTGTGGGAGGAGTTATTTATTGATTTCCATTAATCTTGG

GTTTAACTTTAAATAATTATAGATTGGTGTCTCAATTTTATATCATATTTATTGGAGTAAATTTAACTTT

TTTCCCTCAGCATTTTCTTGGTTTGGGGGGAATGCCTCGCCGATATTCAGATTATGCTGATTGTTATCTA

GTATGAAATAAAATTTCTTCTGCGGGAAGGATTTTGAGTATTATTTCTGTTATTTATTTTTTATTTATTG

TTTTAGAATCTTTACTTCTTCTGCGTT

>KX570804_SSA10 voucher UG76 cytochrome oxidase subunit I (COI)gene, partial cds; mitochondrial.

GAAAACTAGAGGTATTTGGAAGGCTGGGGATAATTTATGCTATACTAACT

ATTGGTATTTTAGGTTTTATTGTTTGGGGTCATCATATGTTTACTGTGGG

AATAGACGTTGATACTCGGGCTTATTTTACTTCAGCAACTATAGTTATTG

CTGTTCCGACGGGGATTAAAATTTTTAGGTGACTTGCTACTTTAGGTGGA

ATAAAGTCTAACAAATTTAGTCCTCTTGTTCTTTGATTTACGGGGTTTTT

ATTTCTATTTACCATGGGGGGATTAACTGGAATTATTCTTGGTAATTCCT

CTGTAGATGTTTGTTTGCACGATACATATTTTGTTGTTGCTCATTTTCAT

TATGTTTTATCTATAGGAATTATTTTTGCTATTATGGGAGGGGCTATCTA

TTGATTTCCACTAATTTTAGGTTTAACTTTAAATAATTATAATCTGGTAT

CTCAGTTTTATATTATGTTTATGGGAGTAAACTTAACATTTTTTCCACAG

CATTTTCTTGGTTTGAGTGGAATGCCTCGACGTTACTCAGATTATCCTGA

CTGTTACTTGTTGTGAAACAAGATTTCTTCTGCAGGGAGTATTTTGAGTG

TTATTTCTGTTATCTATTTTTTATTTATTGTTTTAGAATCTTTCCTTCTT

TTACGAT

>KX570811_SSA10 voucher UG304b cytochrome oxidase subunit I (COI)gene, partial cds; mitochondrial.

GAAAACTAGAGGTATTTGGAAGGCTGGGGATAATTTATGCTATACTAACT

ATTGGTATTTTAGGTTTTATTGTTTGGGGTCATCATATGTTTACTGTGGG

AATAGACGTTGATACTCGGGCTTATTTTACTTCAGCAACTATAGTTATTG

CTGTTCCGACGGGGATTAAAATTTTTAGGTGACTTGCTACTTTAGGTGGA

ATAAAGTCTAACAAATTTAGTCCTCTTGTTCTTTGATTTACGGGGTTTTT

ATTTCTATTTACCATGGGGGGATTAACTGGAATTATTCTTGGTAATTCCT

CTGTAGATGTTTGTTTGCACGATACATATTTTGTTGTTGCTCATTTTCAT

TATGTTTTATCTATAGGAATTATTTTTGCTATTATGGGAGGGGCTATTTA

TTGATTTCCACTAATTTTAGGTTTAACTTTAAATAATTATAATCTGGTAT

CTCAGTTTTATATTATGTTTATGGGAGTAAACTTAACATTTTTTCCACAG

CATTTTCTTGGTTTGAGTGGAATGCCTCGACGTTACTCAGATTATCCTGA

CTGTTACTTGTTGTGAAACAAGATTTCTTCTGCAGGGAGTATTTTGAGTG

TTATTTCTGTTATCTATTTTTTATTTATTGTTTTAGAATCTTTCCTTCTT

TTACGAT

>KX570808_SSA10 voucher UG86 cytochrome oxidase subunit I (COI)gene, partial cds; mitochondrial.

GAAAACTAGAGGTATTTGGAAGGCTGGGGATAATTTATGCTATACTAACT

ATTGGTATTTTAGGTTTTATTGTTTGGGGTCATCATATGTTTACTGTGGG

AATAGACGTTGATACTCGGGCTTATTTTACTTCAGCAACTATAGTTATTG

CTGTTCCGACGGGGATTAAAATTTTTAGGTGACTTGCTACTTTAGGTGGA

ATAAAGTCTAACAAATTTAGTCCTCTTGTTCTTTGATTTACGGGGTTTTT

ATTTCTATTTACCATGGGGGGATTAACTGGAATTATTCTTGGTAATTCCT

CTGTAGATGTTTGTTTGCACGATACATATTTTGTTGTTGCTCATTTTCAT

TATGTTTTATCTATAGGAATTATTTTTGCTATTATGGGAGGGGCTATCTA

TTGATTTCCACTAATTTTAGGTTTAACTTTAAATAATTATAATCTGGTAT

CTCAGTTTTATATTATGTTTATGGGAGTAAACTTAACATTTTTTCCACAG

CATTTTCTTGGTTTGAGTGGAATGCCTCGACGTTACTCAGATTATCCTGA

CTGTTACTTGTTGTGAAACAAGATTTCTTCTGCAGGGAGTATTTTGAGTG

TTATTTCTGTTATCTATTTTTTATTTATTGTTTTAGAATCTTTCCTTCTT

TTACGAT

>KX570809_SSA10 voucher UG86bB cytochrome oxidase subunit I (COI)gene, partial cds; mitochondrial.

GAAAACTAGAGGTATTTGGAAGGCTGGGGATAATTTATGCTATACTAACT

ATTGGTATTTTAGGTTTTATTGTTTGGGGTCATCATATGTTTACTGTGGG

AATAGACGTTGATACTCGGGCTTATTTTACTTCAGCAACTATAGTTATTG

CTGTTCCGACGGGGATTAAAATTTTTAGGTGACTTGCTACTTTAGGTGGA

ATAAAGTCTAACAAATTTAGTCCTCTTGTTCTTTGATTTACGGGGTTTTT

ATTTCTATTTACCATGGGGGGATTAACTGGAATTATTCTTGGTAATTCCT

CTGTAGATGTTTGTTTGCACGATACATATTTTGTTGTTGCTCATTTTCAT

TATGTTTTATCTATAGGAATTATTTTTGCTATTATGGGAGGGGCTATTTA

TTGATTTCCACTAATTTTAGGTTTAACTTTAAATAATTATAATCTGGTAT

CTCAGTTTTATATTATGTTTATGGGAGTAAACTTAACATTTTTTCCACAG

CATTTTCTTGGTTTGAGTGGAATGCCTCGCCGTTACTCAGATTATCCTGA

CTGTTACTTGTTGTGAAACAAGATTTCTTCTGCAGGGAGTATTTTGAGTG

TTATTTCTGTTATCTATTTTTTATTTATTGTTTTAGAATCTTTCCTTCTT

TTACGAT

>KX570806_SSA10 voucher UG221 cytochrome oxidase subunit I (COI)gene, partial cds; mitochondrial.

GTAAACTAGAGGTATTTGGAAGGCTGGGGATAATTTATGCTATACTAACT

ATTGGTATTTTAGGTTTTATTGTTTGGGGTCATCATATGTTTACTGTGGG

AATAGACGTTGATACTCGGGCTTATTTTACTTCAGCAACTATAGTTATTG

CTGTTCCTACGGGAATTAAAATTTTTAGGTGACTTGCTACTTTAGGTGGA

ATAAAGTCTAACAAATTTAGTCCTCTTGTTCTTTGATTTACGGGGTTTTT

ATTTCTATTTACCATGGGGGGATTAACTGGAATTATTCTTGGTAATTCCT

CTGTAGATGTTTGTTTGCACGATACATATTTTGTTGTTGCTCATTTTCAT

TATGTTTTATCTATAGGAATTATTTTTGCTATTATGGGAGGGGCTATCTA

TTGATTTCCACTAATTTTAGGTTTAACTTTAAATAATTATAATCTGGTAT

CTCAGTTTTATATTATGTTTATGGGAGTAAACTTAACATTTTTTCCACAG

CATTTTCTTGGTTTGAGTGGAATGCCTCGACGTTACTCAGATTATCCTGA

CTGTTACTTGTTGTGAAACAAGATTTCTTCTGCAGGGAGTATTTTGAGTG

TTATTTCTGTTATCTATTTTTTATTTATTGTTTTAGAATCTTTCCTTCTT

TTACGAT

>KX570805_SSA10 voucher UG76b cytochrome oxidase subunit I (COI)gene, partial cds; mitochondrial.

GTAAACTAGAGGTATTTGGAAGGCTGGGGATAATTTATGCTATACTAACT

ATTGGTATTTTAGGTTTTATTGTTTGGGGTCATCATATGTTTACTGTGGG

AATAGACGTTGATACTCGGGCTTATTTTACTTCAGCAACTATAGTTATTG

CTGTTCCTACGGGAATTAAAATTTTTAGGTGACTTGCTACTTTAGGTGGA

ATAAAGTCTAACAAATTTAGTCCTCTTGTTCTTTGATTTACGGGGTTTTT

ATTTCTATTTACTATGGGGGGATTAACTGGAATTATTCTTGGTAATTCCT

CTGTAGATGTTTGTTTGCACGATACATATTTTGTTGTTGCTCATTTTCAT

TATGTTTTATCTATAGGAATTATTTTTGCTATTATGGGAGGGGCTATCTA

TTGATTTCCACTAATTTTAGGTTTAACTTTAAATAATTATAATCTGGTAT

CTCAATTTTATATTATGTTTATGGGAGTAAACTTAACATTTTTTCCACAG

CATTTTCTTGGTTTGAGTGGAATGCCTCGACGTTACTCAGATTATCCTGA

CTGTTACTTGTTGTGAAACAAGATTTCTTCTGCAGGGAGTATTTTGAGTG

TTATTTCTGTTATCTATTTTTTATTTATTGTTTTAGAATCTTTCCTTCTT

TTACGAT

>KX570818_SSA12 voucher UG324b cytochrome oxidase subunit I (COI)gene, partial cds; mitochondrial.

GAAAATTAGAGGTATTTGGAAGTTTGGGTATAATTTATGCTATATTAACT

ATTGGTATCTTAGGATTTATTGTCTGAGGTCATCACATATTTACAGTTGG

AATAGATGTTGATACTCGAGCTTATTTTACTTCAGCTACTATAATTATTG

CTGTTCCTACAGGAATTAAAATTTTCAGTTGACTTGCTACTTTGGGTGGA

ATAAAGTCTAATAAATTTAGTCCTCTTGGCCTTTGATTTACAGGATTTTT

ATTTTTATTTACTATGGGCGGATTAACTGGAATTATTCTTGGTAATTCTT

CTGTAGATGTATGTTTGCATGACACTTATTTTGTTGTTGCACATTTTCAT

TATGTTTTATCAATAGGAATCATTTTTGCTATTGTAGGAGGAGTTATCTA

CTGATTTCCATTAATCTTGGGTTTAACTTTAAATAATTACAGCTTGGTGT

CTCAATTTTATATCATGTTTATTGGAGTAAATTTAACTTTTTTTCCTCAG

CATTTTCTTGGCTTAGGAGGGATGCCTCGTCGATATTCAGATTATGCTGA

TTGTTATCTATTATGAAACAAAATTTCTTCTGCGGGAAGGATTTTGAGTA

TCATTTCTGTTATTTATTTTTTATTTGTTGTTTTAGAATCTTTTCTTCTT

ATGCGTT

>KX570837_SSA12 voucher UG290c cytochrome oxidase subunit I (COI)gene, partial cds; mitochondrial.

GAAAATTAGAGGTATTTGGAAGTTTGGGTATAATTTATGCTATATTAACT

ATTGGTATCTTAGGATTTATTGTTTGAGGTCATCACATATTTACAGTTGG

AATAGATGTTGATACTCGAGCTTATTTTACTTCAGCTACTATAATTATTG

CTGTTCCTACAGGAATTAAAATTTTCAGTTGACTTGCTACTTTGGGTGGA

ATAAAGTCTAATAAATTTAGTCCTCTTGGCCTTTGATTTACAGGGTTTTT

ATTTTTATTTACTATGGGCGGATTAACTGGAATTATTCTTGGTAATTCTT

CTGTAGATGTATGTTTACATGACACTTATTTTGTTGTTGCACATTTTCAT

TATGTTTTGTCAATAGGAATCATTTTTGCTATTGTAGGAGGAGTTATCTA

CTGATTTCCATTAATCTTGGGTTTAACTTTAAATAATTACAGCTTGGTGT

CTCAATTTTATATCATGTTTATTGGAGTAAATTTAACTTTTTTTCCTCAG

CATTTTCTTGGCTTAGGAGGGATGCCTCGTCGATATTCAGATTATGCTGA

TTGTTATCTATTATGAAACAAAATTTCTTCTGCGGGAAGGATTTTGAGTA

TCATTTCTGTTATTTATTTTTTATTTGTTGTTTTAGAATCTTTTCTTCTT

ATGCGTT

>KX570829_SSA13 voucher UG211c cytochrome oxidase subunit I (COI)gene, partial cds; mitochondrial.

GAAAATTAGAGGTATTTGGAAGTTTGGGTATAATTTATGCTATATTAACT

ATTGGTATTTTAGGGTTTATTGTTTGAGGTCATCATATATTTACAGTTGG

AATAGATGTTGATACTCGAGCTTATTTTACGTCAGCTACTATGATTATCG

CTGTTCCCACAGGAATTAAAATTTTCAGTTGGCTTGCTACTTTGGGTGGG

ATAAAGTCCAATAAATTCAGGCCCCTTGGCCTTTGATTTACAGGATTTTT

ATTTTTATTTACTATAGGTGGGTTAACCGGAATTATTCTTGGTAACTCTT

CTGTAGATGTATGTTTACATGACACTTATTTTGTTGTTGCACATTTTCAT

TATGTATTATCAATAGGAATCATTTTTGCTATTGTAGGAGGAGTTATTTA

TTGATTTCCATTAATTTTGGGTTTAACTTTAAATAATTACAGCTTGGTGT

CTCAGTTTTATATCATGTTTATTGGAGTAAATTTAACTTTTTTCCCTCAG

CATTTTCTTGGTTTAGGGGGAATGCCTCGTCGATACTCAGATTATGCTGA

TTGCTATCTATTATGAAACAAAATTTCTTCTGCGGGAAGGATTTTGAGTA

TTATTTCTGTTATTTATTTTTTATTTATCGTTTTAGAATCTTTTCTTCTT

ATGCGTT

>KX570833_SSA13 voucher UG293 cytochrome oxidase subunit I (COI)gene, partial cds; mitochondrial.

GAAAATTAGAGGTATTTGGAAGTTTGGGTATAATTTATGCTATATTAACT

ATTGGTATTTTAGGGTTTATTGTTTGAGGTCATCATATATTTACAGTTGG

AATAGATGTTGATACTCGAGCTTATTTTACGTCAGCTACTATGATTATCG

CTGTTCCCACAGGAATTAAAATTTTCAGTTGGCTTGCTACTTTGGGTGGG

ATAAAGTCCAATAAATTCAGGCCCCTTGGCCTTTGATTTACAGGATTTTT

ATTTTTATTTACTATAGGTGGGTTAACCGGAATTATTCTTGGTAACTCTT

CTGTAGATGTATGTTTACATGACACTTATTTTGTTGTTGCACATTTTCAT

TATGTATTATCAATAGGAATCATTTTTGCTATTGTAGGAGGAGTTATTTA

TTGATTTCCATTAATTTTGGGTTTAACTTTAAATAATTACAGCTTGGTGT

CTCAGTTTTATATCATGTTTATTGGAGTAAATTTAACTTTTTTCCCTCAG

CATTTTCTTGGTTTAGGGGGAATGCCTCGTCGATACTCAGATTATGCTGA

TTGCTATCTATTATGAAACAAAATTTCTTCTGCGGGAAGGATTTTGAGTA

TTATTTCTGTTATTTATTTTTTATTTATCGTTTTAGAATCTTTTCTTCTT

ATGCGTT

>KX570852_SSA6 voucher UG76c cytochrome oxidase subunit I (COI)gene, partial cds; mitochondrial.

GTAAACTTGAAGTCTTTGGAAGTCTAGGTATAATTTATGCTATATTAACT

ATTGGTATCCTGGGGTTTATTGTTTGGGGGCATCATATATTTACTGTTGG

GATAGATGTAGATACTCGGGCGTATTTTACTTCAGCTACCATGATCATTG

CTGTTCCAACTGGAATTAAGATTTTTAGGTGGCTGGCAACATTGGGTGGT

ATAAAGTCTAATAAGTTTAGTCCTCTAGTTCTTTGATTTACGGGGTTTTT

ATTTTTATTCACTATGGGGGGCTTAACGGGAATTATTCTTGGCAATTCTT

CTGTCGATGTTTGTTTACATGATACTTATTTTGTTGTTGCTCATTTTCAT

TATGTATTATCTATGGGAATTATTTTTGCCATTATGGGTGGGGTTATCTA

TTGATTTCCTTTAATTATGGGATTAACTTTAAATTATTACAATTTAGTTT

CTCAATTTTATATTATATTTATTGGAGTAAATCTTACATTTTTTCCACAA

CATTTCCTTGGTTTAAGGGGGATACCACGTCGGTATTCAGACTATCCCGA

CTGCTATCTAGTTTGAAATAAAATTTCTTCTGTGGGGAGGATCTTGAGTG

TTATCTCTGTTATTTACTTTTTATTTATTGTTTTAGAGTCTTTCCTTCTA

GTTCGTC

>KX570850_SSA6 voucher UG202b cytochrome oxidase subunit I (COI)gene, partial cds; mitochondrial.

GTAAACTTGAAGTCTTTGGAAGTCTAGGTATAATTTATGCTATATTAACT

ATTGGTATCCTGGGGTTTATTGTTTGGGGTCATCATATATTTACTGTTGG

GATAGATGTAGATACTCGGGCGTATTTTACTTCAGCTACCATGATCATTG

CTGTTCCAACTGGAATTAAGATTTTTAGGTGGCTGGCAACATTGGGTGGT

ATAAAGTCTAATAAGTTTAGTCCTCTAGTTCTTTGATTTACGGGGTTTTT

ATTTTTATTCACTATGGGGGGCTTAACGGGAATTATTCTTGGCAATTCTT

CTGTCGATGTTTGTTTACATGATACTTATTTTGTTGTTGCTCATTTTCAT

TATGTATTATCTATGGGAATTATTTTTGCCATTATGGGTGGGGTTATCTA

TTGATTTCCTTTAATTATGGGATTAACTTTAAATTATTACAATTTAGTTT

CTCAATTTTATATTATATTTATTGGAGTAAATCTTACATTTTTTCCACAA

CATTTCCTTGGTTTAAGGGGGATACCACGTCGGTATTCAGACTATCCCGA

CTGCTATCTAGTTTGAAATAAAATTTCTTCTGTGGGGAGGATCTTGAGTG

TTATCTCTGTTATTTACTTTTTATTTATTGTTTTAGAGTCTTTCCTTCTA

GTTCGTC

>KX570856_SSA9 voucher UG152b cytochrome oxidase subunit I (COI)gene, partial cds; mitochondrial.

GTAAACTTGAAGTATTTGGAAGACTGGGAATAATTTATGCTATGTTAACT

ATTGGTATCCTGGGGTTTATTGTTTGGGGCCATCATATATTTACTGTTGG

GATAGATGTGGACACTCGGGCGTATTTTACTTCAGCTACTATAATTATTG

CCGTTCCTACTGGAATTAAGATTTTTAGGTGACTTGCAACATTGGGCGGT

ATAAAGTCTAATAAGTTCAGTCCACTAGTTCTCTGATTTACAGGTTTTTT

ATTTCTATTTACTATGGGAGGTTTAACTGGAATTATTCTTGGTAATTCTT

CTGTCGACGTTTGTTTACATGACACTTATTTTGTTGTTGCTCATTTTCAT

TATGTATTATCTATGGGAATTATTTTTGCTATTATGGGTGGAATTATCTA

TTGATTTCCTTTAATTCTGGGGTTAACCTTAAATTATTACAATTTAATTT

CTCAATTTTATATTATATTTATTGGAGTAAACCTCACATTTTTTCCACAA

CATTTTCTTGGTTTAGGGGGGATACCGCGTCGGTATTCAGATTATCCAGA

CTGCTATCTAGTTTGAAACAAAATTTCCTCTGTGGGGAGTATCTTGAGCA

TTATTTCTGTTATTTATTTTTTATTTATTGTTTTAGAGTCTTTCCTTCTA

GTTCGGC

>KX570855_SSA9 voucher UG152 cytochrome oxidase subunit I (COI)gene, partial cds; mitochondrial.

GTAAACTTGAAGTATTTGGAAGACTGGGAATAATTTATGCTATGTTAACT

ATTGGTATCCTGGGGTTTATTGTTTGGGGTCATCATATATTTACTGTTGG

AATAGATGTGGACACTCGGGCGTATTTTACTTCAGCTACTATAATTATTG

CCGTTCCTACTGGAATTAAGATTTTTAGGTGACTTGCAACATTGGGCGGT

ATAAAGTCTAATAAGTTCAGTCCACTAGTTCTCTGATTTACAGGTTTTTT

ATTTCTATTTACTATGGGAGGTTTAACTGGAATTATTCTTGGTAATTCTT

CTGTCGACGTTTGTTTACATGACACTTATTTTGTTGTTGCTCATTTTCAT

TATGTATTATCTATGGGAATTATTTTTGCTATTATGGGTGGAATTATCTA

TTGATTTCCTTTAATTCTGGGGTTAACTTTAAATTATTACAATTTAATTT

CTCAATTTTATATTATATTTATTGGAGTAAACCTTACATTTTTTCCACAA

CATTTTCTTGGTTTAGGGGGGATACCGCGTCGGTATTCAGATTATCCAGA

CTGCTATCTAGTTTGAAACAAAATTTCCTCTGTGGGGAGTATCTTGAGCA

TTATTTCTGTTATTTATTTTTTATTTATTGTTTTAGAGTCTTTCCTTCTA

GTTCGGC

>KX570857_Mugerwa_Uganda1 voucher UG150 cytochrome oxidase subunit I (COI)gene, partial cds; mitochondrial.

GAAAATTAGAAGTATTTGGTAGACTAGGTATAATCTATGCTATATTGACT

ATCGGTATCCTTGGCTTTATTGTATGAGGTCACCATATGTTCACTGTTGG

CATAGATGTAGATACTCGAGCTTATTTCACTTCGGCTACTATAATTATCG

CTGTTCCTACAGGAATTAAAATTTTTAGATGATTAGCTACTCTTGGTGGA

ATAAAATCTAACAAGTTTAGGCCGTTGGTTCTTTGGTTTACCGGATTTTT

ATTTCTGTTTACTATGGGAGGTTTAACCGGAATTATTCTTGGTAATTCTT

CTGTTGATATTTGTTTACATGATACATATTTTGTTGTTGCTCATTTTCAT

TATGTACTATCTATAGGAATTATTTTTGCTATTATGGGTGGAATAATTTA

TTGATTTCCTCTAATCTTGGGGCTTACTTTAAACAACTATAATTTAGTTT

CTCAGTTTTATATTATATTCGTTGGAGTAAATTTAACATTTTTTCCCCAA

CATTTTCTTGGTTTGAGAGGGATACCTCGGCGATATTCGGATTACCCCGA

TTGTTATCTATTATGAAATAAACTTTCTTCTGTGGGAAGAATTTTAAGTG

TTGTTTCGATTGTTTACTTTATATGTATTGTGCTAGAGTCATTTATCCTT

TTACGAT

>KX570858_Mugerwa_Uganda1 voucher UG166Nc cytochrome oxidase subunit I (COI)gene, partial cds; mitochondrial.

GAAAATTAGAAGTATTTGGTAGACTAGGTATAATCTATGCTATATTGACT

ATCGGTATCCTTGGCTTTATTGTATGAGGTCACCATATGTTCACTGTTGG

CATAGATGTAGATACTCGAGCTTATTTCACTTCGGCTACTATAATTATCG

CTGTTCCTACAGGAATTAAAATTTTTAGATGATTAGCTACTCTTGGTGGA

ATAAAATCTAACAAGTTTAGGCCGTTGGTTCTTTGGTTTACCGGATTTTT

ATTTCTGTTTACTATGGGAGGTTTAACCGGAATTATTCTTGGTAATTCTT

CTGTTGATATTTGTTTACATGATACATATTTTGTTGTTGCTCATTTTCAT

TATGTACTATCTATAGGAATTATTTTTGCTATTATGGGTGGAATAATTTA

TTGATTTCCTCTAATCTTGGGGCTTACTTTAAACAACTATAATTTAGTTT

CTCAGTTTTATATTATATTCGTTGGAGTAAATTTAACATTTTTTCCCCAA

CATTTTCTTGGTTTGAGAGGGATACCTCGGCGATACTCGGATTACCCCGA

TTGTTATCTATTATGAAATAAACTTTCTTCTGTGGGAAGAATTTTAAGTG

TTGTTTCGATTGTTTACTTTATATGTATTGTGCTAGAGTCATTTATCCTT

TTACGAT

>KX570862_Mugerwa_Uganda1 voucher UG193c cytochrome oxidase subunit I (COI)gene, partial cds; mitochondrial.

GAAAATTAGAAGTATTTGGTAGACTAGGTATAATCTATGCTATATTGACT

ATCGGTATCCTTGGCTTTATTGTATGAGGTCACCATATGTTCACTGTTGG

TATAGATGTAGATACTCGAGCTTATTTCACTTCGGCTACTATAATTATCG

CTGTTCCTACAGGAATTAAAATTTTTAGATGATTAGCTACTCTTGGTGGA

ATAAAATCTAACAAGTTTAGGCCGTTGGTTCTTTGGTTTACCGGATTTTT

ATTTCTGTTTACTATGGGAGGTTTAACCGGAATTATTCTTGGTAATTCTT

CTGTTGATATTTGTTTACATGATACATATTTTGTTGTTGCTCATTTTCAT

TATGTACTATCTATAGGAATTATTTTTGCTATTATGGGTGGAATAATTTA

TTGATTTCCTCTAATCTTGGGGCTTACTTTAAACAACTATAATTTAGTTT

CTCAGTTTTATATTATATTCGTTGGAGTAAATTTAACATTTTTTCCCCAA

CATTTTCTTGGTTTGAGAGGGATACCTCGGCGATACTCGGATTACCCCGA

TTGTTATCTATTATGAAATAAACTTTCTTCTGTGGGAAGAATTTTAAGTG

TTGTTTCGATTGTTTACTTTATATGTATTGTGCTAGAGTCATTTATCCTT

TTACGAT

>KX570868_Mugerwa_Uganda1 voucher UG231d cytochrome oxidase subunit I (COI)gene, partial cds; mitochondrial.

GAAAATTAGAAGTATTTGGTAGACTAGGTATAATCTATGCTATATTGACT

ATCGGTATCCTTGGCTTTATTGTATGAGGTCACCATATGTTCACTGTTGG

CATAGATGTAGATACTCGAGCTTATTTCACTTCGGCTACTATAATTATCG

CTGTTCCTACAGGAATTAAAATTTTTAGATGATTAGCTACTCTTGGTGGA

ATAAAATCTAACAAGTTTAGGCCGTTGGTTCTTTGGTTTACCGGATTTTT

ATTTCTGTTTACTATGGGAGGTTTAACCGGAATTATTCTTGGTAATTCTT

CTGTTGATATTTGTTTACATGATACATATTTTGTTGTTGCTCATTTTCAT

TATGTACTATCTATAGGAATTATTTTTGCTATTATGGGTGGAATAATTTA

TTGATTTCCTCTAATCTTGGGGCTTACTTTAAACAACTATAATTTAGTTT

CTCAGTTTTATATTATATTCGTTGGAGTAAATTTAACATTTTTTCCCCAA

CATTTTCTTGGTTTGAGAGGGATACCTCGGCGATATTCGGATTACCCCGA

TTGTTATCTATTATGAAATAAACTTTCTTCTGTGGGAAGAATTTTAAGTG

TTGTTTCGATTGTTTACTTTATATGTATTGTGCTAGAGTCATTTATCCTT

TTACGAT

>KX570863_Mugerwa_Uganda1 voucher UG193d cytochrome oxidase subunit I (COI)gene, partial cds; mitochondrial.

GAAAATTAGAAGTATTTGGTAGACTAGGTATAATCTATGCTATATTGACT

ATCGGTATCCTTGGCTTTATTGTATGAGGTCACCATATGTTCACTGTTGG

TATAGATGTAGATACTCGAGCTTATTTCACTTCGGCTACTATAATTATCG

CTGTTCCTACAGGAATTAAAATTTTTAGATGATTAGCTACTCTTGGTGGA

ATAAAATCTAACAAGTTTAGGCCGTTGGTTCTTTGGTTTACCGGATTTTT

ATTTCTGTTTACTATGGGAGGTTTAACCGGAATTATTCTTGGTAATTCTT

CTGTTGATATTTGTTTACATGATACATATTTTGTTGTTGCTCATTTTCAT

TATGTACTATCTATAGGAATTATTTTTGCTATTATGGGTGGAATAATTTA

TTGATTTCCTCTAATCTTGGGGCTTACTTTAAACAACTATAATTTAGTTT

CTCAGTTTTATATTATATTCGTTGGAGTAAATTTAACATTTTTTCCCCAA

CATTTTCTTGGTTTGAGAGGGATACCTCGGCGATACTCGGATTACCCCGA

TTGTTATCTATTATGAAATAAACTTTCTTCTGTGGGAAGAATTTTAAGTG

TTGTTTCGATTGTTTACTTTATATGTATTGTGCTAGAGTCATTTATCCTT

TTACGAT

>Bemisia_sp_PDB_1_MN056066.1_Bemisia_sp_cytochrome_c_oxidase_subunit_1_gene,_partial_cds;_mitochondrial

GAAAGCTTGAAGTATTTGGAAGATTAGGAATAATTTATGCTATATTAACT

ATTGGTATTTTAGGTTTTATTGTTTGAGGTCATCACATATTTACCGTTGG

AATAGATGTTGATACTCGAGCATATTTTACTTCAGCTACTATAATTATTG

CTGTTCCTACAGGAATTAAAATTTTTAGATGACTTGCTACATTAGGTGGT

ATAAAAACTAATAAATTTAGGCCGGTAGTTCTTTGATTTACAGGATTCTT

ATTTTTATTTACTATGGGTGGGTTGACTGGAATTATTCTTGGTAATTCTT

CTGTTGATATTTGCTTACATGATACTTATTTTGTTGTTGCTCATTTTCAT

TATGTTTTATCTATAGGAATTATTTTTGCTATTATGGGTGGGTTAATTTA

TTGATTTCCTTTAATTTTAGGGTTGACTTTAAATAATTATAATTTGGTTT

CTCAATTTTATATCATATTTATTGGAGTTAATTTAACATTTTTCCCTCAG

CATTTTCTTGGTTTAAGAGGGATACCTCGTCGATATTCAGATTATCCTGA

TTGTTACTTGTTGTGGAATAAACTTTCTTCTGTGGGAAGAATTTTAAGTG

TTATCTCGGTTGTTTATTTTATATTTATTGTTTTAGAATCTTTTATCCTT

TTACGGT

>Bemisia_atriplex_GU086362_GU086362.1_Bemisia_atriplex_isolate_Bemisia_atriplex_cytochrome_c_oxidase_subunit_1_gene,_partial_cds;_mitochondrial

GAAAGCTTGAAGTTTTTGGTAGGCTTGGGATAATTTATGCCATAATAACT

ATCGGTATTTTGGGTTTTATTGTTTGAGGTCATCATATATTTACTGTTGG

TATAGACGTTGACACTCGGGCATATTTTACTTCAGCCACAATAATTATTG

CTGTTCCTACGGGAATTAAGATTTTTAGGTGGCTTGCTACTTTGGGTGGG

ATAAAATCTAATAAGCTAAGACCTCTTGTTCTTTGATTTACAGGATTTTT

ATTCTTATTTACTATGGGAGGATTAACTGGAATTATTCTTGGAAATTCAT

CTGTTGATGTTTGTTTACACGATACTTATTTTGTTGTTGCTCATTTTCAT

TATGTTTTATCTATAGGAATTATTTTTGCTATTATAGGTGGTGTTATTTA

TTGATTTCCTTTAATTTTAGGTCTAAGCTTAAATAATTATAATTTGGTTT

CTCAGTTTTATATAATATTTATAGGAGTAAACTTAACGTTTTTCCCTCAG

CACTTCCTTGGTCTAAGTGGTATACCCCGGCGATATTCAGATTATCCTGA

TTGTTATTTATTATGAAATAAAATTTCCTCTGCTGGAAGAATTTTAAGTG

TTATTTCTGTTATTTATTTTTTATTTATTATTTTAGAATCTTTTTTAGTT

CTTCGTC

>Bemisia_berbericola_HQ457046_HQ457046.1_Bemisia_berbericola_cytochrome oxidase subunit_1_gene,_partial_cds;_mitochondrial

GAAAACTTGAAGTCTTTGGTAGTCTTGGCATGATTTATGCAATACTTACT

ATTGGTGTATTAGGGTTTATTGTATGGGGTCATCATATGTTTACAGTGGG

AATGGATGTAGACACACGTGCTTATTTCACTTCAGCAACTATAGTAATTG

CTGTTCCTACAGGTATTAAAATTTTTAGGTGGCTAGCTACCTTAGGGGGC

ATAAAATCAAACAAGTACAGTCCTTTATTACTTTGATTTACGGGATTTAT

TTTTCTATTTACAATAGGAGGTTTGACTGGAATTATTTTAGGTAATTCAT

CTGTTGATGTTTGTTTGCACGATACTTATTTTGTTGTGGCTCATTTTCAT

TATGTTTTGTCTATAGGAATTATTTTTGCTATTATAGGAGGTTTTATTTA

TTGATTCCCTTTAGTTATAGGGATTAGCCTTAATATGTATAGCTTGGTTT

CTCAATTTTATATAATATTTTTAGGAGTAAATTTAACTTTTTTCCCACAA

CATTTCTTAGGGTTAGGGGGAATACCTCGGCGATATTCTGATTATGGTGA

TTGTTATTTGCTTTGAAATAAAGTTTCATCTGCAGGTAGGATTTTAAGAT

TAATTTCTATTATTTACTTTTTGTTTATCGTTTTGGAATCTCTAATTTTG

CTTCGTN

>Bemisia_sp_PDB_2-1_MN056067.1_Bemisia_sp cytochrome c oxidase subunit 1 gene, partial_cds;_mitochondrial

GTAAGTTAGAAGTCTTTGGTAGGTTGGGCATGATTTATGCTATATTAACT

ATTGGAGTTTTGGGCTTCATCGTCTGAGGACATCACATATTTACTGTAGG

CATGGATGTAGACACCCGGGCGTATTTCACCTCGGCTACCATAATTATTG

CTGTGCCCACAGGAATTAAGATTTTTAGTTGGTTAGCCACTTTGGGGGGT

ATAAAGTCTAACAAGCTAAGACCCTTAGTGCTTTGGTTTACTGGGTTCTT

GTTTTTATTCACTATGGGGGGTCTAACCGGGATCATCCTAGGAAACTCTT

CCGTGGACGTCTGTCTTCATGACACTTATTTCGTGGTCGCTCATTTTCAT

TATGTGTTATCTATGGGTATTATCTTTGCTATCATGGGAGGCTTCATTTA

CTGATTTCCTCTTGTACTTGGTGTTTCACTTAATGCTTATGGGCTAGTTT

CTCAATTTTATATTATGTTTCTTGGTGTTAATCTTACTTTCTTTCCCCAA

CACTTTTTAGGATTGGGGGGTATACCTCGCCGTTACTCTGATTATGGAGA

TTGTTATTTGCTTTGAAACAAGGTGTCTTCCGCCGGGAGCATCATTAGGT

TAATTTCTATCATCTACTTTTTATTTATTGTTTTAGAGTCTTTAGTTTTA

TTGCGAG

>Bemisia_sp_PDB_2-2_MN056068.1_Bemisia_sp cytochrome c oxidase subunit 1 gene,_partial_cds;_mitochondrial

GTAAGTTAGAAGTCTTTGGTAGGTTGGGCATGATTTATGCTATATTAACT

ATTGGAGTTTTGGGCTTCATCGTCTGAGGACATCACATATTTACTGTAGG

CATGGATGTAGACACCCGGGCGTATTTCACCTCGGCTACCATAATTATTG

CTGTGCCCACAGGAATTAAGATTTTTAGTTGGTTAGCCACTTTGGGGGGT

ATAAAGTCTAACAAGCTAAGACCCTTAGTGCTTTGGTTTACTGGGTTCTT

GTTTTTATTCACTATGGGGGGTCTAACCGGGATCATCCTAGGAAACTCTT

CCGTGGACGTCTGTCTTCATGACACTTATTTCGTGGTCGCTCATTTTCAT

TATGTGTTATCTATGGGTATTATCTTTGCTATCATGGGAGGCTTCATTTA

CTGATTTCCTCTTGTACTTGGTGTTTCACTTAATGCTTATGGGCTAGTTT

CTCAATTTTATATTATGTTTCTTGGTGTTAATCTTACTTTCTTTCCCCAA

CACTTTTTAGGATTGGGGGGTATACCTCGCCGTTACTCTGATTATGGAGA

TTGTTATTTGCTTTGAAACAAGGTGTCTTCCGCCGGGAGCATCATTAGGT

TAATTTCTATCATCTACTTTTTATTTATTGTTTTAGAGTCTTTAGTTTTA

TTGCGAG

>AF418673.2_Bemisia_afer

GTAAACTCGAAGTGTTCGGAAGGTTGGGGATAATTTACGCTATACTTACT

ATTGGAGTCCTTGGCTTCATTGTGTGGGGGCATCACATGTTTACTGTGGG

TATGGATGTAGATACCCGGGCATACTTTACATCTGCCACTATAATTATTG

CAGTGCCCACCGGTATCAAGATTTTCAGCTGGTTGGCCACTTTGGGGGGT

ATAAAGTCTAATAAATTTAGCCCACTTGTGCTTTGATTTACTGGGTTTTT

ATTTCTATTCACCATAGGAGGTCTTACCGGGATCATTTTGGGCAATTCTT

CCGTTGATGTCTGTCTTCATGATACTTACTTTGTGGTTGCCCACTTTCAT

TATGTACTATCAATGGGGATTATCTTCGCCATTATGGGGGGGTTTATCTA

CTGGTTTCCGCTGGTTCTTGGCATCACCCTTAACAATTATAGCCTGGTGG

GCCAATTTTATCTTATATTTCTGGGAGTTAACTTGACATTTTTTCCTCAG

CATTTTTTGGGTTTGGGGGGGATGCCTCGGCGCTATTCTGATTACGCAGA

TTGCTACCTGCTTTGGAATAAAATTTCGTCCATTGGTAGTATTATTAGTC

TTGTTTCTATCATTTACTTTCTATTTATTGTCCTAGAGTCTTTAGTTCTT

TTACGGG

>KF734668_Bemisia_afer

GTAAACTCGAAGTGTTCGGAAGGTTGGGGATAATTTACGCTATACTTACT

ATTGGAGTCCTTGGCTTCATTGTGTGGGGGCATCACATGTTTACTGTGGG

TATGGATGTAGATACCCGGGCATACTTTACATCTGCCACTATAATTATTG

CAGTGCCCACCGGTATCAAGATTTTCAGCTGGTTGGCCACTTTGGGGGGT

ATAAAGTCTAATAAATTTAGCCCACTTGTGCTTTGATTTACTGGGTTTTT

ATTTCTATTCACCATAGGAGGTCTTACCGGGATCATTTTGGGCAATTCTT

CCGTTGATGTCTGTCTTCATGATACTTACTTTGTGGTTGCCCACTTTCAT

TATGTACTATCAATGGGGATTATCTTCGCCATTATGGGGGGGTTTATCTA

CTGGTTTCCGCTGGTTCTTGGCATCACCCTTAACAATTATAGCCTGGTGG

GCCAATTTTATCTTATATTTCTGGGAGTTAACTTGACATTTTTTCCTCAG

CATTTTTTGGGTTTGGGGGGGATGCCTCGGCGCTATTCTGATTACGCAGA

TTGCTACCTGCTTTGGAATAAAATTTCGTCCATTGGTAGTATTATCAGTC

TTGTTTCTATCATTTACTTTCTATTTATTGTCCTAGAGTCTTTAGTCCTT

TTACGGG

>KX570815_SSA11_SubSaharan Africa 11

GAAAATTGGAAGTATTTGGAAGATTGGGCATAATCTATGCTATAATAACT

ATTGGTATTCTAGGGTTTATCGTTTGGGGTCATCATATGTTTACCGTTGG

AATAGATGTTGATACTCGAGCCTATTTTACTTCAGCTACTATAATTATTG

CTGTTCCCACAGGGATTAAGATTTTTAGGTGGTTAGCCACTTTAGGTGGG

ATAAAGACTAATAAGTTTAGGCCCCTTAGTCTTTGATTTACGGGGTTTTT

ATTTTTATTTACCATGGGAGGATTAACTGGAATTATTCTTGGTAACTCTT

CCGTAGATGTTTGTTTACACGATACATATTTTGTTGTTGCTCATTTTCAT

TATGTTTTATCTATGGGGATTATTTTTGCTATCATGGGAGGAATTATTTA

TTGATTCCCATTAATTTTGGGCTTGACTCTAAATAATTTTAGTCTAGTTT

CTCAGTTTTATATAATATTCATGGGGGTAAACTTAACCTTTTTTCCACAA

CATTTCCTTGGCTTAAGTGGGATGCCGCGACGTTACTCTGATTATCCTGA

TTGTTATCTAATATGAAATAAGGTTTCTTCTCTGGGGAGAATTTTAAGTG

TCATCTCTGTTATTTATTTTTTATTTATTGTTTTAGAATCTTTACTTCTT

TTGCGAT
