## Supplemental Data 2 for "Take out the rubbish – Removing NUMTs and pseudogenes from the *Bemisia tabaci* cryptic species mtCOI database"

**Suppl. Data 2: HTS reference COI gene set**

>MH186144_Bemisia tabaci 'MED'_cytochrome c subunit I, partial CDS

GAAAATTAGAGGTATTTGGAAGGTTGGGCATAATTTATGCTATATTGACT

ATTGGTATCTTAGGGTTTATTGTTTGAGGACATCATATATTTACAGTTGG

AATAGATGTAGATACTCGAGCTTATTTCACTTCAGCTACTATGATTATTG

CCGTTCCTACAGGAATTAAAATTTTTAGTTGGCTTGCTACTTTGGGTGGA

ATAAAGTCCAATAAATTCAGGCCCCTTGGCCTTTGATTTACAGGATTTTT

ATTTTTATTTACTATAGGTGGATTAACTGGAATTATTCTTGGTAACTCTT

CTGTAGATGTGTGTTTGCATGACACTTATTTTGTTGTTGCGCATTTTCAT

TATGTCTTATCAATAGGAATTATTTTTGCTATTGTAGGAGGAGTTATCTA

TTGATTTCCATTAATCTTGGGCTTAACCTTAAATAATTATAGCTTGGTGT

CTCAATTTTATATCATGTTCATTGGAGTAAATTTAACTTTTTTTCCTCAG

CATTTTCTTGGTTTGGGGGGAATGCCTCGCCGATATTCAGATTATGCTGA

TTGTTATCTAGTATGGAACAAAATTTCTTCTGCGGGAAGGATTTTGAGTA

TCATTTCTGTTATTTATTTTTTATTTATTGTTTTAGAATCTTTTCTTCTT

TTGCGTT

>MH047295_Bemisia tabaci 'MED'_cytochrome c subunit I, partial CDS

GAAAATTAGAGGTATTTGGAAGGTTGGGCATAATTTATGCTATATTGACT

ATTGGTATCTTAGGGTTTATTGTTTGAGGACATCATATATTTACAGTTGG

AATAGATGTAGATACTCGAGCTTATTTCACTTCAGCTACTATGATTATTG

CCGTTCCTACAGGAATTAAAATTTTTAGTTGGCTTGCTACTTTGGGTGGA

ATAAAGTCCAATAAATTCAGGCCCCTTGGCCTTTGATTTACAGGATTTTT

ATTTTTATTTACTATAGGTGGATTAACTGGAATTATTCTTGGTAACTCTT

CTGTAGATGTGTGTTTGCATGACACTTATTTTGTTGTTGCGCATTTTCAT

TATGTCTTATCAATAGGAATTATTTTTGCTATTGTAGGAGGAGTTATCTA

TTGATTTCCATTAATCTTGGGCTTAACCTTAAATAATTATAGCTTGGTGT

CTCAATTTTATATCATGTTCATTGGAGTAAATTTAACTTTTTTTCCTCAG

CATTTTCTTGGTTTGGGGGGAATGCCTCGCCGATATTCAGATTATGCTGA

TTGTTATCTAGTATGGAACAAAATTTCTTCTGCGGGAAGGATTTTGAGTA

TCATTTCTGTTATTTATTTTTTATTTATTGTTTTAGAATCTTTTCTTCTT

TTGCGTT

>MH186143_Bemisia tabaci 'MED'_cytochrome c subunit I, partial CDS

GAAAATTAGAGGTATTTGGAAGGTTGGGCATAATTTATGCTATATTGACT

ATTGGTATCTTAGGGTTTATTGTTTGAGGACATCATATATTTACAGTTGG

AATAGATGTAGATACTCGAGCTTATTTCACTTCAGCTACTATGATTATTG

CCGTTCCTACAGGAATTAAAATTTTTAGTTGGCTTGCTACTTTGGGTGGA

ATAAAGTCCAATAAATTCAGGCCCCTTGGCCTTTGATTTACAGGATTTTT

ATTTTTATTTACTATAGGTGGATTAACTGGAATTATTCTTGGTAACTCTT

CTGTAGATGTGTGTTTGCATGACACTTATTTTGTTGTTGCGCATTTTCAT

TATGTCTTATCAATAGGAATTATTTTTGCTATTGTAGGAGGAGTTATCTA

TTGATTTCCATTAATCTTGGGCTTAACCTTAAATAATTATAGCTTGGTGT

CTCAATTTTATATCATGTTCATTGGAGTAAATTTAACTTTTTTTCCTCAG

CATTTTCTTGGTTTGGGGGGAATGCCTCGCCGATATTCAGATTATGCTGA

TTGTTATCTAGTATGGAACAAAATTTCTTCTGCGGGAAGGATTTTGAGTA

TCATTTCTGTTATTTATTTTTTATTTATTGTTTTAGAATCTTTTCTTCTT

TTGCGTT

>MH205752_Bemisia tabaci 'MED'_cytochrome c subunit I, partial CDS

GAAAATTAGAGGTATTTGGAAGGTTGGGCATAATTTATGCTATATTGACT

ATTGGTATCTTAGGGTTTATTGTTTGAGGACATCATATATTTACAGTTGG

AATAGATGTAGATACTCGAGCTTATTTCACTTCAGCTACTATGATTATTG

CCGTTCCTACAGGAATTAAAATTTTTAGTTGGCTTGCTACTTTGGGTGGA

ATAAAGTCCAATAAATTCAGGCCCCTTGGCCTTTGATTTACAGGATTTTT

ATTTTTATTTACTATAGGTGGATTAACTGGAATTATTCTTGGTAACTCTT

CTGTAGATGTGTGTTTGCATGACACTTATTTTGTTGTTGCGCATTTTCAT

TATGTCTTATCAATAGGAATTATTTTTGCTATTGTAGGAGGAGTTATCTA

TTGATTTCCATTAATCTTGGGCTTAACCTTAAATAATTATAGCTTGGTGT

CTCAATTTTATATCATGTTCATTGGAGTAAATTTAACTTTTTTTCCTCAG

CATTTTCTTGGTTTGGGGGGAATGCCTCGCCGATATTCAGATTATGCTGA

TTGTTATCTAGTATGGAACAAAATTTCTTCTGCGGGAAGGATTTTGAGTA

TCATTTCTGTTATTTATTTTTTATTTATTGTTTTAGAATCTTTTCTTCTT

TTGCGTT

>KY951447_Bemisia tabaci 'MED'_cytochrome c subunit I, partial CDS

GAAAATTAGAGGTATTTGGAAGGTTGGGCATAATTTATGCTATATTGACT

ATTGGTATCTTAGGGTTTATTGTTTGAGGACATCATATATTTACAGTTGG

AATAGATGTAGATACTCGAGCTTATTTCACTTCAGCTACTATGATTATTG

CCGTTCCTACAGGAATTAAAATTTTTAGTTGGCTTGCTACTTTGGGTGGA

ATAAAGTCCAATAAATTCAGGCCCCTTGGCCTTTGATTTACAGGATTTTT

ATTTTTATTTACTATAGGTGGATTAACTGGAATTATTCTTGGTAACTCTT

CTGTAGATGTGTGTTTGCATGACACTTATTTTGTTGTTGCGCATTTTCAT

TATGTCTTATCAATAGGAATTATTTTTGCTATTGTAGGAGGAGTTATCTA

TTGATTTCCATTAATCTTGGGCTTAACCTTAAATAATTATAGCTTGGTGT

CTCAATTTTATATCATGTTCATTGGAGTAAATTTAACTTTTTTTCCTCAG

CATTTTCTTGGTTTGGGGGGAATGCCTCGCCGATATTCAGACTATGCTGA

TTGTTATCTAGTATGGAACAAAATTTCTTCTGCGGGAAGGATTTTGAGTA

TCATTTCTGTTATTTATTTTTTATTTATTGTTTTAGAATCTTTTCTTCTT

TTGCGTT

>JQ906700_Bemisia tabaci 'MED'_cytochrome c subunit I, partial CDS

GAAAATTAGAGGTATTTGGAAGGTTGGGCATAATTTATGCTATATTGACT

ATTGGTATCTTAGGGTTTATTGTTTGAGGACATCATATATTTACAGTTGG

AATAGATGTAGATACTCGAGCTTATTTCACTTCAGCTACTATGATTATTG

CCGTTCCTACAGGAATTAAAATTTTTAGTTGGCTTGCTACTTTGGGTGGA

ATGAAGTCCAATAAATTCAGGCCCCTTGGCCTTTGATTTACAGGATTTTT

ATTTTTATTTACTATAGGTGGATTAACTGGAATTATTCTTGGTAACTCTT

CTGTAGATGTGTGTTTGCATGACACTTATTTTGTTGTTGCGCATTTTCAT

TATGTCTTATCAATAGGAATTATTTTTGCTATTGTAGGAGGAGTTATCTA

TTGATTTCCATTAATCTTGGGCTTAACCTTAAATAATTATAGCTTGGTGT

CTCAATTTTATATCATGTTCATTGGAGTAAATTTAACTTTTTTTCCTCAG

CATTTTCTTGGTTTGGGGGGAATGCCTCGCCGATATTCAGATTATGCTGA

TTGTTATCTAGTATGGAACAAAATTTCTTCTGCGGGAAGGATTTTGAGTA

TCATTTCTGTTATTTATTTTTTATTTATTGTTTTAGAATCTTTTCTTCTT

TTGCGTT

>MH205753_Bemisia tabaci 'MED'_cytochrome c subunit I, partial CDS

GAAAATTAGAGGTATTTGGAAGGTTGGGCATAATTTATGCTATATTGACT

ATTGGTATCTTAGGGTTTATTGTTTGAGGACATCATATATTTACAGTTGG

AATAGATGTAGATACTCGAGCTTATTTCACTTCAGCTACTATGATTATTG

CTGTTCCTACAGGAATTAAAATTTTTAGTTGGCTTGCTACTTTGGGTGGA

ATAAAGTCCAATAAATTCAGGCCCCTTGGCCTTTGATTTACAGGATTTTT

ATTTTTATTTACTATAGGTGGATTAACTGGAATTATTCTTGGTAACTCTT

CTGTAGATGTGTGTTTGCATGACACTTATTTTGTTGTTGCGCATTTTCAT

TATGTCTTATCAATAGGAATTATTTTTGCTATTGTAGGAGGAGTTATCTA

TTGATTTCCATTAATCTTGGGTTTAACCTTAAATAATTATAGTTTGGTGT

CTCAATTTTATATCATGTTCATTGGAGTAAACTTAACTTTTTTTCCTCAG

CATTTTCTTGGTTTGGGGGGAATGCCTCGCCGATATTCAGATTATGCTGA

TTGTTATCTAGTATGGAACAAAATTTCTTCTGCGGGAAGGATTTTGAGTA

TTATTTCTGTTATTTATTTTTTATTTATTGTTTTAGAATCTTTTCTTCTT

CTGCGTT

>MH205754_Bemisia tabaci 'ASL'_cytochrome c oxidase subunit I, partial CDS

GAAAATTAGAGGTATTTGGAAGGTTGGGTATAATTTATGCTATATTGACT

ATTGGTATTTTAGGGTTTATTGTTTGAGGACATCACATATTTACAGTTGG

AATAGATGTAGATACTCGAGCTTACTTCACTTCAGCTACTATGATTATTG

CTGTTCCTACAGGAATTAAAATTTTTAGTTGGCTTGCTACTTTGGGTGGA

ATAAAGTCTAATAAATTTAGGCCTCTTGGCCTTTGATTTACAGGATTTTT

ATTTTTATTTACTATAGGTGGATTAACTGGAATTATTCTTGGTAACTCTT

CTGTAGATGTATGTTTGCATGACACTTATTTTGTTGTTGCGCATTTTCAT

TATGTCTTATCAATAGGAATTATTTTTGCTATTGTAGGAGGAGTTATCTA

TTGATTTCCATTAATCTTGGGTTTAACCTTAAATAACTATAGTTTGGTGT

CTCAATTTTATATCATGTTCATTGGAGTAAACTTAACTTTTTTTCCTCAG

CATTTTCTTGGTTTGGGAGGAATGCCTCGCCGATATTCAGATTATGCTGA

TTGTTATCTAGTATGGAACAAGATTTCTTCTGCGGGAAGGATTTTGAGTA

TCATTTCTGTTATTTATTTTTTATTTATTGTTTTAGAATCCTTTCTTCTT

CTGCGTT

>KY951452_Bemisia tabaci 'MEAM1'_cytochrome c subunit I, partial CDS

GAAAATTAGAGGTATTTGGAAGGTTGGGTATAATTTATGCTATATTGACT

ATTGGTATTCTAGGGTTTATTGTTTGAGGTCATCATATATTCACAGTTGG

AATAGATGTAGATACTCGAGCTTATTTCACTTCAGCCACTATAATTATTG

CTGTTCCCACAGGAATTAAAATTTTTAGTTGACTTGCTACTTTGGGTGGA

ATAAAGTCTAATAAATTAAGGCCTCTTGGCCTTTGATTTACAGGATTTTT

ATTTTTATTTACTATAGGTGGGTTAACTGGAATTATTCTTGGTAATTCTT

CTGTAGATGTGTGTCTGCATGACACTTATTTTGTTGTTGCACATTTTCAT

TATGTTTTATCAATAGGAATTATTTTTGCTATTGTAGGAGGAGTTATCTA

TTGATTTCCACTAATCTTAGGTTTAACCTTAAATAATTATAGATTGGTGT

CTCAATTTTATATCATGTTTATTGGAGTAAATTTAACTTTTTTTCCTCAG

CATTTTCTTGGTTTAGGGGGAATGCCTCGTCGATATTCAGATTATGCTGA

TTGCTATCTAGTATGAAATAAAATTTCTTCTGCGGGAAGGATTCTGAGTA

TTATTTCTGTTATTTATTTTTTATTTATTGTTTTAGAATCCTTTCTTCTT

CTGCGGT

>KY951450_Bemisia tabaci 'MEAM1'_cytochrome c subunit I, partial CDS

GAAAATTAGAGGTATTTGGAAGGTTGGGTATAATTTATGCTATATTGACT

ATTGGTATTCTAGGGTTTATTGTTTGAGGTCATCATATATTCACAGTTGG

AATAGATGTAGATACTCGAGCTTATTTCACTTCAGCCACTATAATTATTG

CTGTTCCCACAGGAATTAAAATTTTTAGTTGGCTTGCTACTTTGGGTGGA

ATAAAGTCTAATAAATTAAGGCCTCTTGGCCTTTGATTTACAGGATTTTT

ATTTTTATTTACTATAGGTGGGTTAACTGGAATTATTCTTGGTAATTCTT

CTGTAGATGTGTGTCTGCATGACACTTATTTTGTTGTTGCACATTTTCAT

TATGTTTTATCAATAGGAATTATTTTTGCTATTGTAGGAGGAGTTATCTA

TTGATTTCCACTAATCTTAGGTTTAACCTTAAATAATTATAGATTGGTGT

CTCAATTTTATATCATGTTTATTGGAGTAAATTTAACTTTTTTTCCTCAG

CATTTTCTTGGTTTAGGGGGAATGCCTCGTCGATATTCAGATTATGCTGA

TTGCTATCTAGTATGAAATAAAATTTCTTCTGCGGGAAGGATTCTGAGTA

TTATTTCTGTTATTTATTTTTTATTTATTGTTTTAGAATCCTTTCTTCTT

CTGCGGT

>MH186145_Bemisia tabaci 'MEAM1'_cytochrome c subunit I, partial CDS

GAAAATTAGAGGTATTTGGAAGGTTGGGTATAATTTATGCTATATTGACT

ATTGGTATTCTAGGGTTTATTGTTTGAGGTCATCATATATTCACAGTTGG

AATAGATGTAGATACTCGAGCTTATTTCACTTCAGCCACTATAATTATTG

CTGTTCCCACAGGAATTAAAATTTTTAGTTGGCTTGCTACTTTGGGTGGA

ATAAAGTCTAATAAATTAAGGCCTCTTGGCCTTTGATTTACAGGATTTTT

ATTTTTATTTACTATAGGTGGGTTAACTGGAATTATTCTTGGTAATTCTT

CTGTAGATGTGTGTCTGCATGACACTTATTTTGTTGTTGCACATTTTCAT

TATGTTTTATCAATAGGAATTATTTTTGCTATTGTAGGAGGAGTTATCTA

TTGATTTCCACTAATCTTAGGTTTAACCTTAAATAATTATAGATTGGTGT

CTCAATTTTATATCATGTTTATTGGAGTAAATTTAACTTTTTTTCCTCAG

CATTTTCTTGGTTTAGGGGGAATGCCTCGTCGATATTCAGATTATGCTGA

TTGCTATCTAGTATGAAATAAAATTTCTTCTGCGGGAAGGATTCTGAGTA

TTATTTCTGTTATTTATTTTTTATTTATTGTTTTAGAATCCTTTCTTCTT

CTGCGGT

>KY951449_Bemisia tabaci 'MEAM1'_cytochrome c subunit I, partial CDS

GAAAATTAGAGGTATTTGGAAGGTTGGGTATAATTTATGCTATATTGACT

ATTGGTATTCTAGGGTTTATTGTTTGAGGTCATCATATATTCACAGTTGG

AATAGATGTAGATACTCGAGCTTATTTCACTTCAGCCACTATAATTATTG

CTGTTCCCACAGGAATTAAAATTTTTAGTTGACTTGCTACTTTGGGTGGA

ATAAAGTCTAATAAATTAAGGCCTCTTGGCCTTTGATTTACAGGATTTTT

ATTTTTATTTACTATAGGTGGGTTAACTGGAATTATTCTTGGTAATTCTT

CTGTAGATGTGTGTCTGCATGACACTTATTTTGTTGTTGCACATTTTCAT

TATGTTTTATCAATAGGAATTATTTTTGCTATTGTAGGAGGAGTTATCTA

TTGATTTCCACTAATCTTAGGTTTAACCTTAAATAATTATAGATTGGTGT

CTCAATTTTATATCATGTTTATTGGAGTAAATTTAACTTTTTTTCCTCAG

CATTTTCTTGGTTTAGGGGGAATGCCTCGTCGATATTCAGATTATGCTGA

TTGCTATCTAGTATGAAATAAAATTTCTTCTGCGGGAAGGATTCTGAGTA

TTATTTCTGTTATTTATTTTTTATTTATTGTTTTAGAATCCTTTCTTCTT

CTGCGGT

>KY951448_Bemisia tabaci 'IO'_cytochrome c subunit I, partial CDS

GGAAATTGGAGGTATTTGGAAGGTTGGGTATAATTTATGCTATATTAACT

ATTGGCATCTTGGGGTTTATTGTTTGAGGTCACCATATATTTACAGTTGG

AATAGATGTAGATACTCGAGCTTATTTTACCTCAGCTACTATGATTATTG

CTGTTCCTACAGGAATTAAAATTTTCAGTTGGCTTGCTACTTTGGGTGGA

ATAAAGTCTAATAAATTCAGGCCTCTTGGCCTTTGGTTCACAGGATTTTT

ATTTTTATTTACTATGGGTGGATTAACTGGAATTATTCTTGGTAACTCTT

CTGTAGATGTATGTTTGCACGATACTTATTTTGTTGTTGCACATTTTCAT

TATGTTTTATCAATAGGAATCATTTTTGCTATTGTAGGAGGAGTTATTTA

TTGATTTCCATTAATCTTGGGTTTAAGTTTAAATAATTATAGTTTGGTAT

CTCAATTTTATATTATGTTCATTGGAGTAAATTTAACTTTTTTTCCTCAG

CACTTTCTTGGTTTAGGGGGAATGCCTCGACGATATTCAGACTATGCTGA

CTGCTATCTAGTATGAAACAAAATTTCCTCTGCGGGAAGAATTTTGAGTA

TCATTTCTGTTATTTATTTTTTATTTATTGTTTTAGAATCTTTCCTTCTT

CTGCGGT

>AY521259_BEmisia tabaci 'NW1'_cytochrome c subunit I, partial CDS

GAAAATTAGAAGTATTTGGTAGCCTGGGTATAATTTATGCCATGATAACT

ATTGGTATCTTGGGATTTATTGTTTGGGGACATCACATGTTTACTGTTGG

AATAGATGTTGACACTCGGGCTTATTTTACTTCAGCTACTATAGTTATTG

CTGTTCCGACGGGAATTAAAATCTTTAGGTGGCTTGCTACTCTAGGTGGA

ATAAAGTCTAATAAGTTTAGACCCCTAGTTCTCTGATTTACAGGATTTCT

ATTTTTATTTACTATGGGTGGATTAACTGGAATTATTCTTGGTAATTCTT

CGGTAGATGTGTGTCTTCATGATACTTATTTTGTTGTTGCTCATTTTCAT

TATGTTTTATCTATAGGAATTATCTTTGCTATTATAGGCGGAGTAATTTA

TTGATTTCCTTTAATTTTAGGTCTAACTTTAAACAATTATAACCTGGTGT

CTCAATTTTATATAATATTTGTGGGAGTAAACCTGACATTTTTTCCACAG

CATTTTCTTGGTTTAAGTGGAATACCTCGTCGGTATTCTGACTACCCTGA

TTGTTATTTACTATGAAATAAAATTTCCTCTGGGGGGAGGGTCTTAAGTG

TTATTTCTGTTATTTATTTTTTATTTATTATTTTAGAGTCTTTTCTTCTC

TTACGGC

>MK386668_Bemisia tabaci 'NW_Barbados1920'_cytochrome c subunit I, partial CDS

GAAAATTAGAAGTATTTGGTAGGCTGGGCATAATTTATGCCATGATAACT

ATTGGTATCTTGGGATTTATTGTTTGGGGACATCACATGTTTACTGTTGG

AATAGATGTTGACACTCGGGCTTATTTTACTTCAGCTACTATAGTTATTG

CTGTTCCGACGGGAATTAAAATCTTTAGGTGGCTTGCTACTCTAGGTGGA

ATAAAGTCTAATAAGTTTAGACCCCTAGTTCTCTGATTTACAGGATTTCT

ATTTTTATTTACTATGGGTGGATTAACTGGAATTATTCTTGGTAATTCTT

CGGTAGATGTGTGTCTTCATGATACTTATTTTGTTGTTGCTCATTTTCAT

TATGTTTTATCTATAGGAATTATCTTTGCTATTATAGGCGGAGTAATTTA

TTGATTTCCTTTAATTTTAGGTCTAACTTTAAACAATTATAACCTGGTGT

CTCAATTTTATATAATATTTGTGGGAGTAAACCTGACATTTTTTCCACAG

CATTTTCTTGGTTTAAGTGGGATACCTCGTCGGTATTCTGACTACCCTGA

TTGTTATCTACTATGAAATAAAATTTCCTCTGGGGGGAGGGTCTTAAGTG

TTATTTCTGTTATTTATTTTTTATTTATTATCTTAGAGTCTTTTCTTCTC

TTACGGC

>MH191394_Bemisia tabaci 'NW2'_cytochrome c subunit I, partial CDS

GAAAATTAGAAGTATTTGGTAGGTTGGGCATAATTTATGCCATGATAACT

ATTGGCATTTTGGGATTTATTGTTTGGGGCCATCACATGTTTACTGTCGG

AATAGATGTTGACACCCGAGCTTATTTTACCTCAGCTACTATAGTTATTG

CTGTTCCGACGGGAATTAAAATTTTTAGGTGGCTTGCTACTCTAGGTGGA

ATAAAGTCTAATAAGTTTAGACCCCTAGTTCTCTGATTTACAGGATTTCT

ATTTTTATTTACTATGGGTGGATTAACTGGAATTATTCTTGGTAATTCCT

CGGTAGATGTGTGTCTTCACGATACCTACTTTGTTGTTGCTCATTTTCAT

TATGTTTTATCGATAGGAATTATCTTTGCTATTATAGGCGGAGTAATTTA

TTGATTTCCTTTAATTTTAGGCTTAACTCTAAACAATTATAATTTGGTGG

CTCAATTTTATATAATATTTGTGGGGGTAAACCTGACATTTTTTCCACAG

CATTTTCTTGGTTTAAGTGGAATGCCTCGTCGGTATTCTGATTATCCTGA

TTGTTATTTACTATGAAATAAAATTTCCTCTGGGGGGAGGGTCTTGAGCG

TTATTTCTGTTATTTATTTTTTATTTATTATTTTAGAATCTTTTCTTCTT

TTACGAC

>MH191393_Bemisia tabaci 'NW2'_cytochrome c subunit I, partial CDS

GAAAATTAGAAGTATTTGGTAGGTTGGGCATAATTTATGCCATGATAACT

ATTGGCATTTTGGGATTTATTGTTTGGGGCCATCACATGTTTACTGTCGG

AATAGATGTTGACACCCGAGCTTATTTTACCTCAGCTACTATAGTTATTG

CTGTTCCGACGGGAATTAAAATTTTTAGGTGGCTTGCTACTCTAGGTGGA

ATAAAGTCTAATAAGTTTAGACCCCTAGTTCTCTGATTTACAGGATTTCT

ATTTTTATTTACTATGGGTGGATTAACTGGAATTATTCTTGGTAATTCCT

CGGTAGATGTGTGTCTTCACGATACCTACTTTGTTGTTGCTCATTTTCAT

TATGTTTTATCGATAGGAATTATCTTTGCTATTATAGGCGGAGTAATTTA

TTGATTTCCTTTAATTTTAGGCTTAACTCTAAACAATTATAATTTGGTGG

CTCAATTTTATATAATATTTGTGGGGGTAAACCTGACATTTTTTCCACAG

CATTTTCTTGGTTTAAGTGGAATGCCTCGTCGGTATTCTGATTATCCTGA

TTGTTATTTACTATGAAATAAAATTTCCTCTGGGGGGAGGGTCTTGAGCG

TTATTTCTGTTATTTATTTTTTATTTATTATTTTAGAATCTTTTCTTCTT

TTACGAC

>KY951451_Bemisia tabaci 'Aus'_cytochrome c subunit I, partial CDS

GAAAACTTGAGGTATTTGGGAGACTCGGAATAATTTACGCTATAATAACT

ATTGGTATTCTTGGTTTTATTGTTTGGGGTCATCATATATTTACTGTTGG

GATAGATGTTGATACTCGAGCTTACTTTACTTCAGCCACTATAATTATTG

CTGTTCCAACTGGAATCAAAATTTTTAGGTGGCTTGCTACCTTAGGTGGA

ATAAGGGCTAACAAATTTAGTCCTCTTGTGCTTTGATTCACAGGATTTTT

ATTCTTATTTACCATGGGTGGGTTAACTGGGATTATTCTTGGTAATTCTT

CTGTTGATGTCTGCTTGCACGATACTTATTTTGTTGTTGCTCATTTTCAT

TATGTTTTATCTATGGGAATTATTTTTGCTATCGTGGGAGGTCTCATTTA

TTGATTTCCATTAATTCTAGGTTTAACATTAAATAGACATAATTTGGTTT

CTCAGTTTTACGTTATATTTTTGGGAGTTAATTTAACGTTTTTTCCACAA

CATTTCCTTGGTTTAAGCGGAATACCTCGCCGGTACTCAGATTATCCCGA

TTGCTATCTAATATGAAATAAAATTTCTTCTGCGGGGAGTATCTTGAGCA

TTATTTCTGTTATCTATTTTCTATTTATTGTTTTAGAATCTTTGCTGCTT

TTGCGGT

>KJ778614_Bemisia tabaci 'Asia_I'_cytochrome c subunit I, partial CDS

GGAAACTTGAGGTATTTGGCAGGTTAGGAATAATTTATGCTATAATAACT

ATTGGCATTTTGGGGTTTATTGTTTGAGGTCATCATATATTTACTGTTGG

TATAGATGTTGATACTCGAGCTTATTTTACTTCAGCTACTATGGTTATTG

CTGTTCCAACTGGGATTAAGATTTTCAGGTGGCTTGCTACTTTAGGTGGA

ATAAAATCCAATAAATTAAGGCCGTTAGTTCTTTGATTTACAGGATTTTT

ATTCTTATTTACCATGGGTGGACTAACTGGGATTATTCTTGGTAATTCTT

CTGTTGATGTTTGTTTGCATGATACTTATTTTGTTGTTGCTCATTTTCAT

TATGTTTTATCCATAGGAATCATTTTTGCTATCATAGGAGGTTTTATTTA

CTGATTTCCATTAATCTTAGGTCTAACATTAAATAACCATAATTTGGTAT

CTCAGTTTTATATTATATTTTTGGGCGTTAACCTAACATTTTTTCCACAA

CACTTTCTTGGATTAAGCGGAATACCTCGTCGGTATTCAGATTATCCTGA

TTGTTATCTCATATGAAATAAAATTTCTTCTGCGGGGAGTATCTTGAGAA

TTATTTCTGTTATCTATTTTTTATTTATTGTTTTAGAATCTTTGCTTCTT

TTACGGC

>KX714967_Bemisia tabaci 'Asia_II-7'_emiliae_cytochrome c subunit I, partial CDS

GAAAACTTGAGGTATTTGGCAGGTTAGGTATAATTTATGCTATAGTAACG

ATTGGTATTTTAGGTTTTATTGTTTGAGGTCATCATATATTTACTGTTGG

GATGGATGTTGATACCCGGGCCTATTTTACCTCAGCTACTATGATTATTG

CTGTTCCGACTGGGATTAAAATTTTTAGGTGACTTGCTACTCTAGGTGGG

ATAAAATCTAATAGATTTAGCCCCCTTGGGCTTTGGTTTACTGGATTTCT

TTTTTTATTTACTATGGGTGGGTTAACTGGGATTATTCTTGGTAATTCTT

CTGTTGATGTCTGCTTACATGATACTTATTTTGTTGTTGCTCATTTTCAT

TATGTTTTATCTATAGGAATTATTTTTGCTATTGTGGGAGGGGTTATTTA

CTGATTTCCGTTAATCTTGGGCTTAACACTGAATAGCCACAGTCTGGTAT

CACAGTTTTATATTATGTTTTTGGGAGTAAACTTAACTTTTTTTCCACAA

CATTTTCTTGGGCTAAGAGGAATACCTCGCCGATATTCAGACTATCCTGA

TTGTTACTTGATATGAAATAAGATTTCTTCTGCGGGGAGAATTTTGAGCA

TTATTTCTGTTATTTATTTTTTATTTATTGTTTTAGAGTCGTTGCTTCTT

TTACGTC

>KX714968_Bemisia sp. 'JpL' cytochrome c subunit I, partial CDS

GAAAACTTGAAGTTTTTGGTAGACTAGGAATAATTTATGCTATGTTAACT

ATTGGTATTTTAGGTTTTATTGTTTGAGGTCATCACATATTTACTGTTGG

GATAGATGTTGATACTCGGGCGTATTTTACTTCAGCTACAATAATTATTG

CTGTTCCTACGGGGATTAAAATCTTTAGTTGACTTGCTACTTTAGGAGGT

ATAAAGTCTAATAAGTTTAGTCCTCTTGTTCTTTGATTTACAGGGTTTTT

ATTTTTATTTACTATGGGTGGTTTAACTGGAATCATTTTGGGTAATTCTT

CTGTTGATGTTTGTTTACATGATACTTATTTTGTTGTTGCTCATTTTCAC

TATGTTTTATCTATGGGAATTATTTTTGCTGTTATGGGTGGGATTATTTA

TTGGTTTCCCCTGGTTTTAGGTCTGAGTTTAAATAATTATAGTTTAGTGT

CTCAATTTTATATGATATTCATGGGAGTAAATTTAACATTTTTTCCTCAA

CACTTTCTTGGCTTAGGGGGAATACCTCGTCGATATTCAGATTATCCTGA

TTGTTATCTTTTGTGGAACAAAATTTCCTCTGCGGGAAGCATCTTAAGTA

TTATTTCTGTCATTTATTTTTTATTTATTATTCTTGAATCTTTATTACTT

CTTCGAT

>MK940754_SSA2_Cytochrome c oxidase subunit I, partial CDS

GTAAACTTGAAGTGTTTGGAAGACTGGGAATAATTTATGCTATGCTAACT

ATCGGTATTCTGGGATTTATTGTTTGGGGTCATCATATATTTACTGTTGG

GATAGATGTGGATACTCGGGCTTATTTTACTTCAGCTACTATAATTATTG

CTGTTCCTACTGGAATTAAGATCTTTAGGTGACTCGCAACATTGGGTGGT

ATAAAGTCTAATAAGTTCAGTCCTCTAGTTCTTTGATTTACAGGTTTCTT

ATTTTTATTTACTATAGGGGGTTTAACTGGAATTATTCTTGGCAACTCTT

CTGTTGACGTTTGTCTACATGACACTTATTTTGTTGTTGCTCATTTTCAT

TATGTATTATCTATGGGAATTATTTTTGCTATTATGGGTGGGATTATTTA

TTGATTTCCTTTAATCCTAGGGCTTACTTTAAATTATTATAATCTAATTT

CTCAATTTTATATTATATTTATTGGAGTAAATCTTACATTTTTTCCACAA

CATTTCCTTGGTTTAAGGGGGATGCCTCGCCGGTATTCAGATTATCCAGA

CTGCTATCTAGTTTGAAACAAAATTTCTTCTGTGGGGAGTATCTTGAGTA

TTATTTCTGTTATTTATTTTTTATTTATTGTTTTAGAGTCTTTTCTTCTG

ATCCGGC

>MK940753_SSA1_Cytochrome c oxidase subunit I, partial CDS

GTAAACTCGAAGTATTTGGAAGTCTAGGTATAATTTATGCTATGTTAACT

ATTGGTATTCTGGGGTTTATTGTTTGGGGTCATCATATATTTACTGTCGG

GATAGATGTGGACACTCGGGCGTATTTCACTTCAGCTACCATAATTATTG

CTGTTCCTACCGGAATCAAGATTTTTAGGTGACTGGCAACATTGGGTGGT

ATAAAGTCTAATAAGTTTAGTCCGCTAGTTCTTTGATTTACGGGGTTTTT

ATTTTTATTTACTATAGGAGGTTTAACTGGAATTATTCTTGGCAATTCTT

CTGTCGACGTTTGCTTACATGACACTTATTTTGTTGTTGCTCATTTTCAT

TATGTATTATCTATGGGAATTATTTTTGCTATTATGGGTGGGATTATTTA

TTGATTTCCCTTAATTCTAGGGTTAACTTTAAATTATTACAATTTAATTT

CTCAATTCTATATTATATTTATTGGAGTAAATCTTACATTTTTTCCCCAG

CATTTTCTTGGTTTGAGGGGAATACCGCGTCGGTATTCAGACTATCCAGA

TTGCTACCTAGTTTGAAATAAAATTTCTTCTGTGGGTAGGATCTTGAGTA

TTATTTCTGTTATTTATTTTTTATTTATTGTTTTAGAGTCTTTTCTTCTG

GTTCGTC

>KF734668_COX1_Bafer_Wang COX1: cytochrome c oxidase subunit I, partial CDS

GTAAACTCGAAGTGTTCGGAAGGTTGGGGATAATTTACGCTATACTTACT

ATTGGAGTCCTTGGCTTCATTGTGTGGGGGCATCACATGTTTACTGTGGG

TATGGATGTAGATACCCGGGCATACTTTACATCTGCCACTATAATTATTG

CAGTGCCCACCGGTATCAAGATTTTCAGCTGGTTGGCCACTTTGGGGGGT

ATAAAGTCTAATAAATTTAGCCCACTTGTGCTTTGATTTACTGGGTTTTT

ATTTCTATTCACCATAGGAGGTCTTACCGGGATCATTTTGGGCAATTCTT

CCGTTGATGTCTGTCTTCATGATACTTACTTTGTGGTTGCCCACTTTCAT

TATGTACTATCAATGGGGATTATCTTCGCCATTATGGGGGGGTTTATCTA

CTGGTTTCCGCTGGTTCTTGGCATCACCCTTAACAATTATAGCCTGGTGG

GCCAATTTTATCTTATATTTCTGGGAGTTAACTTGACATTTTTTCCTCAG

CATTTTTTGGGTTTGGGGGGGATGCCTCGGCGCTATTCTGATTACGCAGA

TTGCTACCTGCTTTGGAATAAAATTTCGTCCATTGGTAGTATTATCAGTC

TTGTTTCTATCATTTACTTTCTATTTATTGTCCTAGAGTCTTTAGTCCTT

TTACGGG
