## Supplemental Tabel 1 for "Take out the rubbish – Removing NUMTs and pseudogenes from the *Bemisia tabaci* cryptic species mtCOI database"

**Supplemental Table 1:** Summary of up-dated CSIRO v2 *Bemisia* cryptic species mtCOI dataset ofcBoykin et al.^1^ (Note: MEAM2†, SSA-6, SSA9, SSA10, SSA11, SSAS12, SSA13 were obtained from Mugerwa et al.^2^. Trimmed (657bp) and clean COI partial mtCOI sequences can be downloaded from the CSIRO data repository <web-link here>. This current version includes also related *Bemisia* species mtCOI dataset. All clean COI partial mtCOI sequences listed were used to infer the phylogeny as shown in Figure 5. Note that for confidence in species status, HTS-derived mitogenomes should be obtained for the following species: Japan 1, China 1, China 2, Asia III, Australia, Indonesia, AsiaII_1, AsiaII_5, AsiaII_6, AsiaII_8, AsiaII_11, AsiaII_3, AsiaII_9, AsiaII_10, Afirca_Cameroon, Italy1, SSA10, African ‘MEAM2’, MED-Burkina-Faso, SSA12, SSA13, SSA11, New World-2-2, Sudan (New World), SSA1, SSA6, SSA2, SSA9, SSA3, *Bemisia*_sp_PDB_1 (MN056066), *B. atriplex*, *Bemisia*_sp_PDB_2-1 (MN056067), Bemisia_sp_PDB_2-1 (MN056068) and *Bemisia berbericola*.

| **CSIRO COI database** (Boykin et al.^1^) | **Clean COI partial sequences** | **NUMTs/potential NUMTs** | **Comments** |
| --- | --- | --- | --- |
| **NW1 & NW2**  JF901839, JF901844, DQ130053, JN969967, JN969968, JN969969, AJ550167, AJ550168, AY057128, AY057129, AY057133, FN821787, AY057126, DQ130058, DQ130059, EU427729, DQ130060, DQ130061, AY057134, EU760727, AY521259, JF901836, JF901837, JF901838, JN689350, JN689351, JN689352, JN689354, JN689355, JN689353, MK386668^#^ | JF901839, JF901844, DQ130053, AY057129, EU427729, DQ130061, EU760727, AY521259, JN689350, JN689351, JN689352, JN689354, JN689355, MK386668^#^ | AY057134, AY057126, DQ130058, DQ130060, JF901837, JF901836, DQ130059, JN689353, JF901838, AY057128, AJ550168, FN821787, AY057133, JN969967†, JN969969†, JN969968†, AJ550167† | [1] JF901844, JF901839 (‘NW2-2’ in Fig. 5): Previous GenBank entries as ‘NW’. Uncorrected *p*-dist estimates above the current revised 2% species boundary and is closer to the NW2 species, but with phylogenetic analysis indicated these sequences being clustered in a separate branch.  † Potential NUMTs  # Partial mtCOI gene from MK386668 obtained from HTS generated historical sample^3^. This sequence was not in the dataset of Boykin et al.^1^. |
| **Asia1**  AJ748388, AJ748398, AJ748383, DQ986959, EU192044, HM137323, HM137330, DQ116641, AJ748359, AJ748360, AJ748361, AJ748365, AJ748369, AJ748370, AJ748371, AM040596, HM590148, HM590156, HM590157, HM590168, HM590169, HM590171, AM408895, HM590178, DQ116646, HM590174, HM590172, HM590154, DQ116655, JN855572, JN703454, JN703455, JX827728, HM590153, HM590150, HM590159, HM590160, HM590161, DQ116663, DQ116664, HM590152, HM590180, HM590162, HM590163, HM590173, AB248260, AB248262, HE653683, HE653704, HE653706, HE653710, AY686093, AJ510061, AJ510078, AY686095, AF164671, EF528552 | AJ748398, HM137323, AJ748359, AJ748361, AJ748365, AJ748369, AJ748371, HM590156, HM590171, HM590154, JN855572, JN703454, HM590159, HM590180, AB248260, HE653683, HE653706, HE653710, AF164671 | DQ116641, DQ116664, DQ116663, DQ116655, HM590148, HM590178, AY686093, HM590150, HM590168, DQ116646, HM590174, HM590163, EF528552, HE653704, JN703455, HM590160, JX827728, HM590172, AM408895, HM590153, HM590161, DQ986959, AJ748370, AJ748388, HM590162, AB248262, AJ510061, AJ510078, AJ748360, HM590152, HM590157, HM137330, AJ748383, AY686095, AM040596, HM590169, HM590173, EU192044 |  |
| **China2**  AY686072, KC113546, KC959606, KC959600, HQ916820, KC959601, JX678666, KC959599 | KC113546, HQ916820, KC959606, KC959601, KC959599, KC959600, JX678666 | AY686072 | Only AY686072 was listed in the Boykin et al.^1^ dataset. Seven additional sequences were retrieved from GenBank (accessed 11-Sept. 2018). |
| **China1**  GQ139495, HM137328, HM137315, HM137351, DQ309078, HM137317, AY686091, AY686089, AY686085 | GQ139495, HM137328, HM137315, HM137351, DQ309078 | HM137317, AY686091, AY686089, AY686085 |  |
| **Australia & Australia II**  GU086328, DQ130052, KC109797, JX416166, GU086325, GU086326, GU086327, HQ457045 | GU086328, KC109797, JX416166, HQ457045 | GU086325, GU086327, DQ130052, GU086326 | GU086326: amino acid substitution at (nucleotide position 119) a highly conserved site across all *Bemisia* species based on mitogenome sequences (Fig. 1).  GU086328 shares 100% sequence identity at this 657bp mtCOI region with the AUS (AY951451) mitogenome COI gene region as reported by Tay et al.^4^. |
| **AsiaII_1**  HM137337, AJ867557, JN855567, HM590132, JN703449, HM590129, HM590138, JN855579, GU585370, AJ510057, AJ510065, JN703450, HM590137, JN703452, HM590133, HM590135, AJ748385, FJ802393, EU192047, JN855578, AJ748394, JX827726, JN855574, JN855576, HM590143, AY686094 EU192065, GU585369, FJ710473, FJ802389, AJ510058, GU585375, GU585373, JX827725, JX827727, DQ309077, JN855569, HM590142, HM590139, HM590134, HM137326, HM590141, HM590140, JN703453, HM590124, HM590126, HM590131 | HM137337, AJ867557, JN855567, HM590132, JN703449^¥^, HM590129^¥^, HM590138^¥^, JN855579^¥^ | GU585370, AJ510057, AJ510065, JN703450, HM590137, JN703452, HM590133, HM590135, AJ748385, FJ802393, EU192047, JN855578, AJ748394, JX827726, JN855574, JN855576, HM590143, AY686094 EU192065, GU585369, FJ710473, FJ802389, AJ510058, GU585375, GU585373, JX827725, JX827727, DQ309077*, JN855569*, HM590142*, HM590139*, HM590134*, HM137326*, HM590141, HM590140, JN703453, HM590124, HM590126, HM590131 | * Amino acid substitutions in these sequences occurred at highly conserved mtCOI gene region when aligned against the mitogenomes of other *B. tabaci* cryptic species.  ^¥^ These four sequences have a Threonine (T) substitution at amino acid residue position 200, while the remaining four sequences possessed an Isoleucine (I) which is present also for the majority of species with HTS-generated mitogenomes, with the exception of *B. afer* (Leucine), and New World species (V). |
| **AsiaII_2**  AY686088 |  | AY686088 | Potential NUMT – This sequence represents the only GenBank entry; substation at amino acid position 210 from V/I (observed in *Bemisia* spp.) to G with negative BLOSUM90 (i.e., for comparing between closely related sequences^5^; values (i.e., alignment of two amino acids found less often then by chance) of -5 for both changes of either I/G, or V/G. |
| **AsiaII_3**  AJ783706.2, DQ309074, EU192045, DQ309075, AJ867556 | AJ783706.2, DQ309074 | DQ309075, AJ867556, EU192045 | Clean sequences of AsiaII-3 from China |
| **AsiaII_4**  AY686083 |  | AY686083 | Potential NUMT. Two glycine residue substitutions detected at inter-species conserved COI gene regions at positions 146 (W in rest of all species) and 161 (V in rest of all species), with BLOSUM90 values of -4 observed between W/G and between V/G. |
| **AsiaII_5**  AJ748391, AJ748393, AJ748395, AF418666.2, AJ748376, AM408901 AJ748379, DQ116673, AJ748384, HM590144, AJ748390 | AJ748391, AJ748393, AJ748395, AF418666.2, AJ748376, AM408901 | AJ748379, DQ116673, AJ748384, HM590144, AJ748390† | †: potential NUMT – significant amino acid substitution at position 103 (highly conserved in all *Bemisia* species) V: hydrophobic to D: hydrophilic (BLOSUM90 value: -5). |
| **ASIAII_6**  AJ784261.2, DQ174520, AB308125, AB308126, AB440788, AB308120, AB308124, HM137338, HM137352, EU192052, EU192054, AB308123 | DQ174520, AB308125, AB308126, AB440788, AB308124 | AB308123, HM137352, AJ784261.2, AB308120*, HM137338*, EU192052†, EU192054† | * potential NUMTs  † intra-species amino acid substitution W/D detected (BLOSUM90 value: -6; i.e., the change is highly unrelated and expected less than by chance). |
| **AsiaII_7/*B. emiliae***  GQ139492, AJ748378, AM408899, DQ174522, DQ116650, AJ748375, DQ174521, DQ116660, DQ116661, AY686075, DQ116662, AY686064, DQ174523, AJ748372 | GQ139492, AJ748378, DQ174522, DQ116650, AJ748375, DQ174521, DQ116660†, DQ116661† | DQ116662, AY686064, DQ174523, AJ748372, AM408899, AY686075 | AY686075: significant aa substitutions *cf*. all other Asia II-7  † DQ116660, DQ116661: These two sequences are bordering the species p-dist against other AsiaII-7 (1.83%). |
| **AsiaII_8**  HM590188.2, AJ748362, AM040593, HM590185, AJ748358, AM408898, HM590182, HM590184, HM590181, AJ748374, AJ748357 | HM590188.2, AJ748362, AJ748357 | AM040593, HM590185, AJ748358, AM408898, HM590182, HM590184, HM590181, AJ748374 | Clean sequences are AsiaII-8 from India |
| **AsiaII_9**  HM137313, HM137345 | HM137313, HM137345 |  | AsiaII_9 sequences from GenBank (accessed 01-Feb-2019) |
| **AsiaII_10**  HM137339, HM137356 | HM137356, HM137339 |  |  |
| **AsiaII_11**  HM590147, HM590146 | HM590147 | HM590146 |  |
| **Italy1**  AY827596, AY827601, AY827598, AY827600, AY827603, AY827602, AY827599 | AY827601, AY827598, AY827600 | AY827603, AY827602, AY827599, AY827596† | †: potential NUMT, 185 F observed but Y conserved in all *Bemisia* spp. |
| **Sub-Saharan Africa 1**  AY057180, AY903462, AY057185, AY057210, AY057149, AY903487, AY563648, AY903470, AY903511, AY903512, AY563679, AF418668, AF418667, AY057162, AF344264, AY903479, DQ130054, AY057169, AY903469, AY903510, AY057179, AF344276, AF344278, AY903483, AY563702, AY903461, AY903492, AY903493, AY903494, AY827591, AY057178, AY903507, AY057181, AY057182, AF344267, AY903475, AY903502, AY563695, AY563666, AY903464, AF344284, AY903490, AY057183, AY057168, AY563652, AY057151, AY563641, AY563674, AY563657 | AY057180, AY903462, AY057149, AY903487, AY563648, AY903470, AY903511, AY903512, AY563679, AF418668, AF418667, AY057162, AY903479, DQ130054, AY057169 | AY903469, AY903510, AY057179, AF344276, AF344278, AY903483, AY563702, AY903461, AY903492, AY903493, AY903494, AY827591, AY057178, AY903507, AY057181, AY057182, AF344267, AY903475, AY903502, AY563695, AY563666, AY903464, AF344284, AY903490, AY057183, AY057168, AY563652, AY057151, AY563641, AY563674, AY563657, AY057185, AY057210, AF344264 |  |
| **Sub-Saharan Africa 2**  AY057173, AY563646, AY057141, AY827605, AY827604, AY563662, AY057146, AY827611, AY563664, AY827607, AY563667, DQ130063, AM040607, EU760745, GU086361, AY057143 | AY057173, AY563646, AY057141, AY827605, AY827604 | AY563662, AY057146, AY827611, AY563664, AY827607, AY563667, DQ130063, AM040607, EU760745, GU086361, AY057143 |  |
| **Sub- Saharan Africa 3**  AF344257†, KM377923, KM377925, KF425602, KF425604, KM377934, HQ908651, MG565972, | KM377923, KM377925, KF425602, KF425604, KM377934, HQ908651, MG565972 | AF344257† | † from Boykin et al.^1^; seven additional sequences were downloaded from GenBank (accessed 01-Feb-2019). |
| **Sub-Saharan Africa 4:**  AF344245, AF344246, AF344247, AF344255, AF344254, AF344249, AF344251, AF344252 |  | AF344245, AF344246, AF344247, AF344255, AF344254, AF344248, AF344249, AF344251, AF344252, AF344250, AF344253 | AF344248, AF344250, AF344253 downloaded from GenBank (accessed 01-Feb-2019). |
| **MEAM1**  HM070414, JN689356, JN689357, JN689358, AY686062, AY686063, EU192068, HE653718, EU263624, EU263625, AY686073, EF398083, EF398086, EF398088, EF398090, EF398091, EF398113, EF398087, DQ989525, AY686078, GQ332577, FN821808, FN821803, FN821798, FN821804, FN821807, FN821802, FN821801, FN821800, FN821806, DQ130055, DQ130056, DQ133373, GU977249, AM180064, AM408896, AF321927, AJ748368, AM040594, HM070410, GU086350, GU086351, GU086352, GU086353, HM070413, AY766373, DQ174536, EF398127, HM070412, AF418671.3, GU086341, AB204577, DQ174537, AB204578, AB204580, AB204581, GU086347, GU086346, GU086354, GU086355, GU086356, AJ550173, AM176570, AJ517768, GU977267, AJ510067, AJ510075, AJ510079, AJ510081, AJ510076, AJ510071, GU977269, GU977268, GU977273, AJ550174, AJ877260, GU086344, GU086345, GU086357, GU086358, DQ174538, GU086348, AB473559, GU086342, DQ989539, DQ174534, DQ174530, FN821790, FN821791, FN821794, FN821797, DQ133382, AY057123, HM070411, GU086340, GU086343, GU086359, GU086360, AF340215, DQ174535 | HM070414, JN689356*, GQ332577*, FN821808*, FN821798, FN821801, DQ133373*, GU977249*, HM070410*, GU086350, GU086351, GU086352, GU086353, HM070413, AB204577*, AB204580, AB204581*, GU086346, GU086354, GU086355, GU086356, GU086344, GU086345, GU086357, GU086358, AB473559, GU086342*, DQ174530, DQ133382, HM070411*, GU086340*, GU086359, GU08636, AM176570 | EF398091, GU977268, AJ517768, FN821803, FN821804, AM180064, DQ174534, FN821800, GU977267, AJ510081, DQ130055, AJ510075, AJ748368, EU263624, AY766373, AY686073, AY686078, FN821794, AJ877260, EU192068, AF321927, AY686063, AM040594, HM070412, AY057123, GU977269, DQ989525, GU086348, AJ550174, GU086341, FN821790, FN821802, DQ174537, AJ550173, EF398083, EU263625, EF398113, EF398090, FN821807, AJ510076, AB204578, FN821797, DQ174538, HE653718, EF398127, GU086343, DQ174536, AM408896, DQ130056, FN821791, AY686062, JN689357, JN689358, DQ989539, AJ510067, GU977273, AF418671 GU086347, EF398086, EF398088, EF398087, AJ510079, AJ510071, FN821806, AF340215, DQ174535 | Clean MEAM1 sequences from: Yemen, USA, Arizona, UAE, Taiwan, Syria, Saudi Arabia, Pakistan, Kuwait, Japan, Iraq, Iran, Indonesia, Egypt, Cuba, China, Brazil, Australia  * Indicates populations within a single highly related branch: China (GQ332577); USA (HM070411, GU086340); Cuba (FN821808); Egypt (DQ133373; GU977249); Indonesia (HM070410); Japan (AB204581; AB204577); Brazil (JN689356); Taiwan (GU086342); Australia (HM070411) (data not shown) |
| **MEAM2**  AJ550177, KX679576†, AB308110†, KY951453†, KX679574†, FJ939600†, FJ939602† |  | AJ550177, KX679576†, AB308110†, KY951453†, KX679574†, FJ939600†, FJ939602† | MEAM2 is NUMTs of MEAM1^4^.  † sequences as identified in Tay et al.^4^ |
| **Indian Ocean**  AJ550171, AY903538, AY903522, AY903523, AY903537, AY903539, AJ550182, AY903525, AJ550178, AY903536, EU760735, AY903527, AJ550180, AJ550179 | AJ550171, AY903538, AY903522, AY903537, AY903539, AJ550182 | AY903525, AJ550178, AY903536, EU760735, AY903527, AJ550180, AJ550179, AY903523 |  |
| **Uganda/Uganda1**  AY903576, AY903553, AY903577, AY903553, KX570857^a^, KX570858^a^, KX570862 ^a^, KX570868 ^a^, KX570863^a^ | AY903576, KX570857^a^, KX570858^a^, KX570862^a^, KX570868^a^, KX570863^a^ | AY903553, AY903577, AY903553 | a: Not in Boykin et al.^1^; named as ‘Uganda1’ in Mugerwa et al.^2^. |
| **SSA5**  AM040598 |  | AM040598 | This sequence is a chimeric PCR product between SSA1 and SSA2, see Suppl. Fig. 1 |
| **MED**  FJ766381, FJ766432, GU086329, DQ365856, DQ133378, GU086333, AM691059, GU086339, AM176575, GQ139500, DQ365857, GU086330, AM691062, FJ025797, AM691067, AM040609, GU086332, EU760721, AM176571, GU086335, DQ302946, AY827617, FJ766385, FJ766400, FJ766405, FJ766408, FJ766387, FJ766391, AY903534, AY827579, AY057136, AY827606, AY903556, AY903531, FJ766383, AY827588, AY827590, AY903540, AY827589, EF694108, GU086337, GU086338, EF694112, EF667476, AY903578, AY057138, AF342773, HE653724, AM176573, DQ174540, EF694111, AJ517769, AM691063, AM180063, AF342775, AY827614, GU086336, AF342769, AY827612, AM691055, EU192061, EU192049, AY827613, AY827615, GU086334, AB297898, GU086331, AB297897, AY827619, DQ174539, AY903529, AY903564, AY903551, AY827581, AY827582, AY827580, GU168793, EU760741, AM691064, FJ766429, FJ766384, FJ766431, AM176574 | FJ766381, FJ766432, GU086329, GU086333, AM691059, GU086339, GU086330, AM691062, AM691067, AM040609, GU086332, EU760721, AM176571, GU086335, DQ302946, FJ766385, FJ766400, FJ766405, FJ766387, FJ766391, AY057136, AY903556, AY903531, FJ766383, AY827588, AY827590, AY903540, AY827589 | EF694108, GU086337, GU086338, EF694112, EF667476, AY903578, AY057138, AF342773, HE653724, AM176573, DQ174540, EF694111, AJ517769^3’^, AM691063, AM180063, AF342775, AY827614, GU086336, AF342769^3’^, AY827612, AM691055, EU192061^5’3’^, EU192049^5’3’^, AY827613, AY827615, GU086334, AB297898, GU086331, AB297897, AY827619, DQ174539, AY903529^3’^, AY903564^3’^, AY903551, AY827581, AY827582^3’^, AY827580, GU168793, EU760741, GQ139500, AM176575, DQ133378^5’^, FJ025797^3’^, FJ766408^3’^_,_ AY903534^3’^, DQ365856, DQ365857, AM691064**, FJ766429^††^, FJ766384^††^, FJ766431^††^, AM176574^ƒ^, AY827617, AY827606, AY827579 | **Note:** locations of INDELs on affected sequences are indicated by 5’ and/or 3’.  Refer to Fig. 3 for alignments of all MED-NUMTs with no evidence of INDELs against corresponding of partial mtCOI amino acid residues of related *B. tabaci* cryptic species as obtained via HTS methods. |
| **AsiaIII**  DQ174527, AB440792, DQ174528 | DQ174527, AB440792, DQ174528 |  |  |
| **Africa_Cameroon**  EU760739 | EU760739 |  |  |
| **Japan2**  AB308111, AB240967, AB308118, AB308119, AB308115, AB308116, AB308117 | AB308111, AB308119, AB308115, AB308116, AB308117, KX714968† | AB240967, AB308118 | † as reported in Tay et al.^6^ |
| **Japan1**  AB440785, AB440786, AB440790 | AB440786 | AB440785, AB440790 |  |
| **China3 (N=2)**  AJ748382, EU192051 |  | AJ748382, EU192051 |  |
| **MEAM2† / (African-MEAM2)**  KX570778 | KX570778 |  | † renamed as ‘African MEAM2’ in Fig. 5. Authenticity of the sequence requires HTS mitogenome confirmation. |
| **Sub-Saharan African 10**  KX570804, KX570811, KX570808, KX570809, KX570806, KX570805 | KX570804, KX570811, KX570808, KX570809, KX570806, KX570805 |  | Authenticity of the SSA10 species status requires HTS mitogenome confirmation. |
| **Sub-Saharan African 12**  KX570818, KX570837 | KX570818, KX570837 |  | Authenticity of the SSA12 species status requires HTS mitogenome confirmation. |
| **Sub-Saharan African 13**  KX570829, KX570833 | KX570829, KX570833 |  | Authenticity of the SSA13 species status requires HTS mitogenome confirmation. |
| **Sub-Saharan African 6**  KX570852, KX570850 | KX570852, KX570850 |  | Authenticity of the SSA6 species status requires HTS mitogenome confirmation. |
| **Sub-Saharan African 9**  KX570856, KX570855 | KX570856, KX570855 |  | Authenticity of the SSA9 species status requires HTS mitogenome confirmation. |
| **Sub-Saharan African 11**  KX570815 | KX570815 |  | Authenticity of the SSA11 species status requires HTS mitogenome confirmation. |
| ***Bemisia afer***  KF734668^GB^, AF418673.2 ^GB^, GQ139515†, GU220055†, | KF734668 ^GB^, AF418673.2^GB^ | GQ139515, GU220055 | GB: From GenBank (accessed 01-Feb-2019).  † From Boykin et al.^1^ |
| ***Bemisia atriplex***  GU086363, GU086362, HQ457047 | GU086362 | GU086363, HQ457047 | Potential NUMTs difficult to predict, HTS-derived mitogenome from this species is needed |
| ***Bemisia berbericola***  HQ457046 | HQ457046 |  |  |
| *Bemisia*_sp_PDB_1 | MN056066 |  | Require HTS-draft mitogenome to confirm species |
| *Bemisia*_sp_PDB_2-1  *Bemisia*_sp_PDB_2-2 | MN056067, MN056068 |  | Require HTS-draft mitogenome to confirm species |

**Note:** sequences excluded from Boykin et al.^1^ dataset were: AB536801, AB536794, AB536800, AB536793, AB558172, AY572538, AJ748380, AY521251, AY648941, AY521262, AY057221, AM179435, AM179436, AM179438, AM179440, AM179429, AM179446, JF693935, AY521265, AM179423, AM179431, DQ989555, GU220056, AY057220, AY572539.

**References**

1.	Boykin, L. M., Savill, A. & De Barro, P. Updated mtCOI reference dataset for the *Bemisia tabaci* species complex *F1000Res*. **6**, 1835. doi:10.12688/f1000research.12858.1. (2017).

2.	Mugerwa, H., et al. African ancestry of New World, *Bemisia tabaci*-whitefly species. *Sci Rep.* **8**, article 2734. doi:10.1038/s41598-018-20956-3. (2018).

3.	Kunz, D., et al. Draft mitochondrial DNA genome of a 1920 Barbados cryptic *Bemisia tabaci* 'New World' species (Hemiptera: Aleyrodidae). *Mitochondrial DNA B*. **4**, 1183-4. doi:10.1080/23802359.2019.1591197. (2019).

4.	Tay, W. T., et al. The Trouble with MEAM2: Implications of Pseudogenes on Species Delimitation in the Globally Invasive *Bemisia tabaci* (Hemiptera: Aleyrodidae) Cryptic Species Complex. *Genome Biol Evol*. **9**, 2732-8. doi:10.1093/gbe/evx173. (2017).

5.	Henikoff, S. & Henikoff, J. G. Amino acid substitution matrices from protein blocks. *Proc Natl Acad Sci U S A.* **89**, 10915-9. doi:10.1073/pnas.89.22.10915. (1992).

6.	Tay, W. T., et al. Novel molecular approach to define pest species status and tritrophic interactions from historical *Bemisia* specimens. *Sci Rep*. **7**, article 429. doi:ARTN 42910.1038/s41598-017-00528-7. (2017).
